# Synapomorphic variations in the THAP domains of the human THAP protein family and its homologs

**DOI:** 10.1101/2021.02.01.429122

**Authors:** Hiral M. Sanghavi, Sharmistha Majumdar

## Abstract

The THAP (<u>Th</u>anatos-<u>a</u>ssociated <u>p</u>rotein) domain is a DNA-binding domain which binds DNA via a zinc coordinating C2CH motif. Although THAP domains share a conserved structural fold, they bind different DNA sequences in different THAP proteins which in turn perform distinct cellular functions. In this study, we investigate (using multiple sequence alignment, *in silico* motif and secondary structure prediction) THAP domain conservation within the homologs of the human THAP (hTHAP) protein family. We report that there is significant variation in sequence and predicted secondary structure elements across hTHAP homologs. Interestingly, we report that the THAP domain can be either longer or shorter than the conventional 90 residues and the amino terminal C2CH motif within the THAP domain serves as a hotspot for insertion or deletion. Our results lay the foundation for future studies which will further our understanding of the evolution of THAP domain and regulation of its function.

## Introduction

The THAP (<u>Th</u>anatos-<u>a</u>ssociated <u>p</u>rotein) domain is a DNA-binding domain which is reported to be 80-90 amino acid residues long and mostly located at the amino terminal end of the corresponding protein. THAP domain containing proteins have been recently reported in diverse groups of animals such as humans, chicken, zebrafish, *C. elegans* and *Drosophila melanogaster* (1–4). No known or predicted proteins containing THAP domains have been found in plants, yeast, fungi or bacteria, suggesting that the THAP domain is a novel protein domain restricted to animals.

THAP domain-containing proteins are involved in diverse cellular functions. For example, Drosophila P element transposase (DmTNP), the cell-cycle transcription factor E2F6 (1) and the transcriptional corepressor CtBP-1 in *C. elegans* (5). The human THAP protein family is a group of twelve proteins (hTHAP0-hTHAP11) which are all characterised by amino-terminal THAP domains (6) and are implicated in cell-cycle regulation, apoptosis, angiogenesis, pluripotency of stem cells (7–11). THAP family members have also been implicated in a variety of human diseases including heart disease (12), torsional dystonia (13) angiogenesis and cancer (14, 15).

Although the THAP domain of THAP proteins share low primary sequence identity (∼10% sequence identity in human THAP proteins) as is typical of large DNA-binding protein families (16), there is strong conservation of the overall protein fold (Suppl. Fig. 1A), as well as secondary structure elements namely the characteristic β;-α-β; fold (1,3, 6, 17, 18), with four loops (L1-L4) flanking and interconnecting the *β;* sheets and the a helix. Recent structural studies illustrate how THAP proteins recognize specific DNA sites through bipartite recognition of adjacent major and minor grooves by specific residues (18): the *β* sheet interacts with the DNA major groove (GC rich sequence in DmTNP) while the carboxy terminal loop 4 (L4) interacts by π-stacking interactions with the DNA minor groove (AT rich sequence in DmTNP) via basic amino acid residues (3, 18, 19).

**Figure 1.**
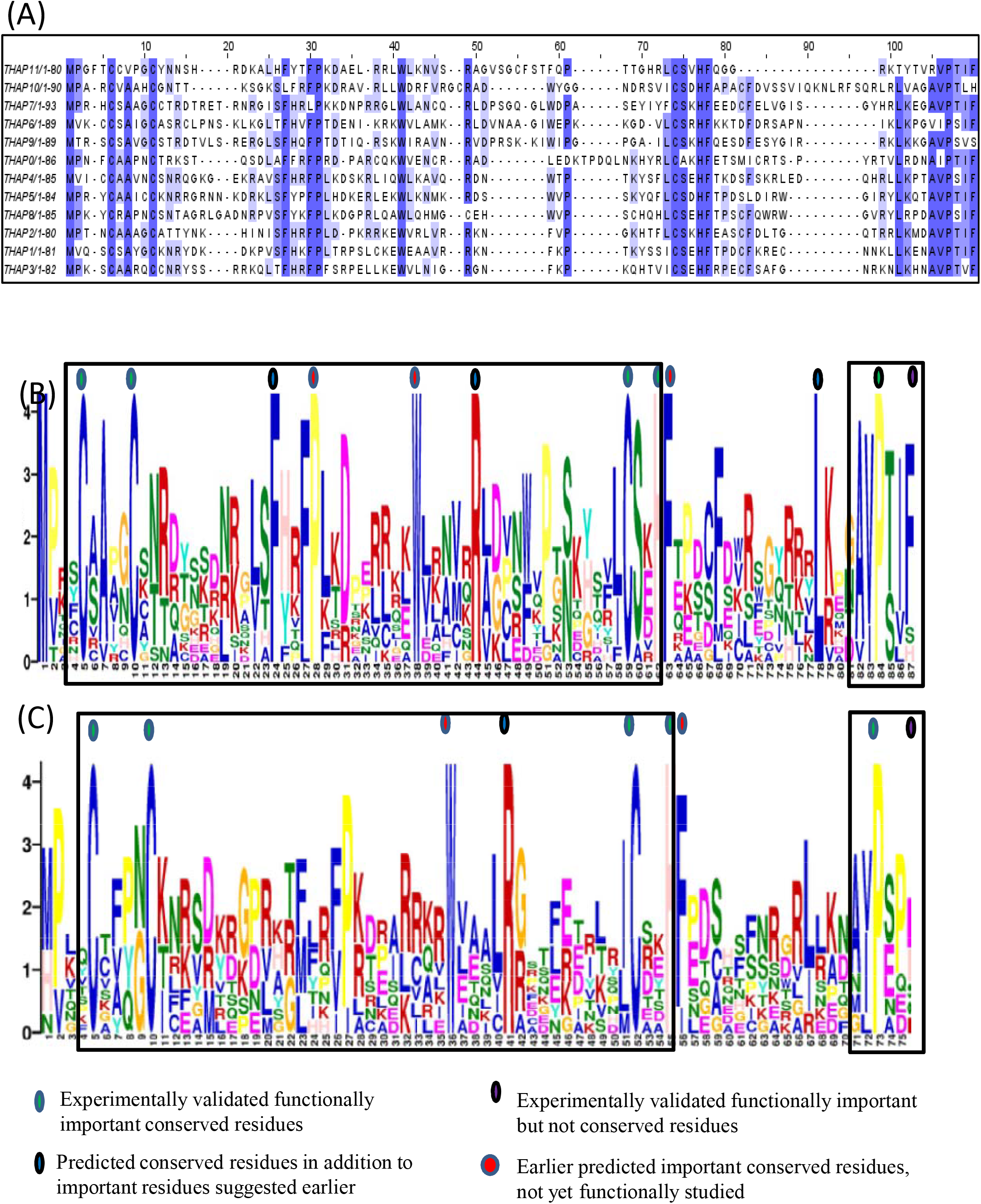
MSA and GLAM2 predicts conserved residues within THAP domain. (A) MSA of all the twelve human THAP proteins highlight conserved residues when viewed in jalview. The conserved residues [functionally important residues (green), function uncharacterised (red), novel conserved residues identified in this study (blue)] within the (B)THAP domain of all twelve hTHAP proteins and (C) other THAP proteins are represented as a position weighted matrix (PWM). Functionally important non-conserved residues (purple) are also highlighted

The conserved sequence signatures in the THAP domain are: (1) C_2_CH (consensus: Cys-X_2– 4_-Cys-X_35–50_-Cys-X_2_-His) zinc-coordinating motif (2) four invariant residues, P, W, F and P, (3) a consensus carboxy terminal AVPTIF box that marks the end of the THAP domain (6, 19). It is to be noted that the consensus C_2_CH motif is significantly different from the classical C2H2 zinc finger motifs (20). THAP domains, with more than 300 identified members, are the second most prevalent zinc-coordinating DNA-binding domain after the classical C_2_H_2_ class of zinc fingers (1, 6, 16,20). The invariant residues are a part of the β;-α-β; secondary structural fold. For example, the first conserved Pro is generally found at the beginning of Loop 2, Trp is found in the centre of helix1, Phe is found at the beginning of loop 3 and the second conserved Pro is a part of the loop 4 that is concluded by AVPTIF (Suppl. figure 1A). It has been experimentally demonstrated that the conserved sequence signatures and the consensus *βαβ* structural fold (3, 18, 19) are indispensable for DNA binding by the THAP domain.

Not much is known of the importance of the four loops (L1-L4) in the β;-α-β; secondary structural fold. L4, which forms the carboxy terminal end of the THAP domain, contains basic residues (Arg65, Arg66 and Arg67 in DmTNP, Arg65 in hTHAP1) involved in binding the minor groove of DNA (17, 18) and the consensus AVPTIF motif. L4 is reported to be flexible unlike the rigid central core of the THAP domain (3, 8, 18, 19) and has been observed to undergo structural changes after binding to DNA in hTHAP1 (17). Interestingly, L4 in different THAP proteins is characterised by the variability in length and primary sequence (3, 8, 18, 19).

The THAP domain is an example of a domain shared between DNA binding proteins and active DNA transposases. Some other examples of such shared domains include the DNA-binding domain of the BED zinc finger, which is shared by both chromatin-boundary element-binding proteins BEAF and DREF (Feschotte and Pritham 2007). It is interesting to note that each THAP protein appears to bind distinct DNA sites. For example, DmTNP (GTTAAGTGGA) (18), hTHAP1 (AGTACGGGCAA) (1), hTHAP5 (GTGTAACTAAC) (21) and hTHAP11 (ACTAYRNNNCCCR) (22) (Suppl. figure 1B) despite similar overall secondary structure (Suppl. figure 1C). This is similar to the basic leucine zipper motif (bZIP) containing proteins where different bZIP proteins bind different DNA sites (23). The difference in DNA binding specificities of THAP proteins is speculated to be due to the variation in the amino acid residues that form *β* sheets (which directly interacts with DNA), the number and sequence of amino acids before the first C of the C2CH motif and the length and composition of loop 4 (18, 19). Till date, there is no comprehensive analysis of the possible diversity in sequence and structural elements in the THAP domains of THAP proteins and their homologs.

In this study, we identify possible synapomorphic (sequence and structure) variations in the hTHAP proteins and their homologs using multiple sequence alignment, *in silico* secondary structure and motif prediction. The predicted secondary structure, presence of basic amino acid residue in L4 in our results agree with the already available structural and biochemical data for the THAP domains of DmTNP, *C. elegans* CtBP-1, hTHAP1. We report conserved amino acid residues in the THAP domain in addition to the ones that are already reported. We identify interesting THAP protein homologs with longer and shorter THAP domains than the conventional ∼90 residue long THAP domain. Identification of a few hotspots for insertions and deletions within the THAP domain challenges the existing understanding about the THAP domain and opens avenues to investigate the evolutionary adaptations of the THAP domain, which is exciting because of the restriction of THAP domain to kingdom animalia.

## Methods

### Curation of THAP protein sequences

There are multiple databases which curate and document protein sequences based on either the families that the protein belongs to (Pfam) (24) or the common patterns in the protein sequence (PROSITE) (25). PROSITE was chosen for this study because the analysis is based on the sequence patterns of various THAP domains. The search string “THAP domain” identified <u>PS50950</u> PROSITE documentation with 45 true positive protein sequences containing THAP type sequence patterns (additional file 1). Of the 45 proteins, *C.elegans* CDC14, CTBP1, Lin36, Lin15B and *Drosophila* P element transposase (DmTNP) were THAP domain containing proteins which are not homologs of any human THAP protein. Thus, they are referred to as “Other THAP proteins” in this study.

### Identification of the THAP domain in the curated protein sequences

Each human THAP protein was aligned with its homologs using Clustal Omega (26). Briefly, Clustal Omega uses HMM models to align multiple sequences and identify identical residues or residues with similar chemical properties, at a position, in the aligned set of sequences. The THAP domain of each human THAP protein homolog was manually identified by using the conserved AVPTIF motif as the carboxy terminal boundary of the domain. These THAP domain sequences for each homolog were stored in a separate word file.

### Identification of conserved residues in the THAP domain

GLAM2 (27) was used to identify a gapped motif in the THAP domain. Briefly, the THAP domain protein sequence from each human THAP protein and its homologs were submitted to GLAM2. It uses an extension of gapless Gibbs sampling algorithm which examines the sequences provided by the user and gives an alignment of different segments of these sequences. This alignment is scored based on position-specific insertion and deletion possibilities. The conserved residues revealed by GLAM2 motifs were validated by multiple sequence alignment (MSA) of the THAP domains of each human THAP protein and its homologs generated using Clustal Omega (26). The GLAM2 and MSA results were carefully analyzed to record the conservation of residues within the THAP domain (C2CH, P,W, F, P, F). The homologs that did not have even one of the above mentioned conserved residues or had replaced the conserved residues with some other amino acid were highlighted in the results.

### THAP domain secondary structure prediction

The secondary structure elements of each THAP domain was predicted using JPRED (28), PSIPRED (29), SPIDER3 (30). Briefly, JPRED constructs a MSA using PSI-BLAST (31) for individual input sequences and uses it to predict local secondary structure using Jnet (32). Additionally, JPRED does a PDB search to identify possible structural homologs of the submitted protein sequences. PSIPRED uses two feed forward neural networks to analyse the PSI-BLAST output. SPIDER3 uses bidirectional recurrent neural networks which capture non-local interactions to accurately predict the secondary structures of the given protein sequences.

### Sequence Curation of human THAP protein homologs

PROTEIN database within the NCBI databank was used to extract the protein sequences of human THAP protein homologs. For each human THAP protein, a keyword search was performed using the protein name (for example, THAP1). Multiple protein sequences of the same protein were available for each organism. Thus, to avoid redundancy among the protein sequences, only the longest sequence was chosen as a THAP protein homolog. The entries which had partial or [PREDICTED] or hypothetical or uncharacterized protein in their names were excluded from the study.

## Results

### Conservation of invariant residues in the THAP domain of hTHAP family proteins and other THAP domain containing proteins

THAP domain-containing proteins (45 proteins identified by PROSITE) were divided into two groups (a) members of human THAP family (hTHAP0-hTHAP11) (b) Other THAP proteins (*C.elegans* CDC14, CTBP1, Lin36, Lin15B, DmTNP). Zebrafish E2F6 and *C. elegans* Him-17, which were earlier reported to contain THAP domains (1, 3) were also added to this group.

The THAP domain sequence in all the group a and b proteins was determined by the presence of the C2CH motif at the amino terminus and an AVPTIF box at the carboxy terminus, as described in the methods. Interestingly, more than one putative THAP domain were reported for *C. elegans* CDC14 (two THAP domains), Lin15B (two THAP domains) and Him-17 (six THAP domains) (1, 3). The two CDC14 THAP domains and six Him-17 THAP domains are very different from each other respectively except the consensus invariant residues within the THAP domain. Thus, only putative THAP domain, which retained the conserved Pro (required for DNA binding ; (3, 18, 19) of the AVPTIF box, was included in this study [one domain each for *C. elegans* CDC14 and Lin15B and four domains for Him-17 (1, 2, 3, 4)].

MSA (ClustalW) was independently performed for THAP domain sequences within each group, after which conserved residues were identified; these included the five residues namely 3 Cys and His of the C2CH motif and Pro of the AVPTIF motif, which have been earlier reported to be functionally indispensable for DNA binding by the THAP domain (3, 18, 19) as seen in figure 1A.

The THAP domain sequences for group a and b were independently submitted to GLAM2 which identifies underlying gapped motifs (as PWMs) in the input sequences after aligning them. (Highlighted by boxes in figure 1A and 1B). The C2CH motif and the AVPTIF box consists of two distinct motifs which constitute the THAP domain. Both of which are functionally distinct where the C2CH motif co-ordinates the zinc ion and the AVPTIF box directly interacts with DNA. These two motifs also form different structural folds, i.e. the C2CH motif folds into the *βαβ* fold whereas the AVPTIF box forms a loop. Different THAP proteins have different inter motif spacers. For example, the intermotif spacer between the His of the C2CH motif and the Ala of the AVPTIF motif is 18 residues in hTHAP1 and 24 residues in DmTNP. These GLAM2 identified gapped motifs for group a and b proteins were then analysed by eye to record the conservation of specific residues within the THAP domain. The five invariant residues (Cys5, Cys10, Cys54, His57, Pro78; green circles in Figure 1B and 1C) as well as some residues of unknown functional significance (Pro26, Trp36, Phe58; residue numbers correspond to hTHAP1; red circles, Figure 1B) which were earlier reported to be conserved in human THAP proteins (6, 19) were found to be conserved. Interestingly, in group b proteins, Trp and Phe (Trp39 and Phe58 in *C.elegans* CtBP -1; red circles in figure 1C) were found to be conserved but the Pro within the C2CH motif is not conserved.

In addition to these, three other residues (Phe22, Arg42, Leu72; residue numbers correspond to hTHAP1; blue circles, Figure 1B) were found to be conserved in group a proteins. The Phe and Arg are a part of the C2CH zinc coordinating motif and Leu is within L4. However, the group b proteins only had a conserved Arg (Arg45 in *C.elegans* CtBP-1) as seen in Figure 1C (blue circle). Interestingly, in the both group a and b proteins, the Phe of the AVPTIF motif was not seen to be strictly conserved (purple circle in figure 1B and C) and was replaced by Ser (hTHAP9), His (hTHAP10), Glu (CtBP-1), Val (DmTNP) or Pro (zE2F6).

### Consensus secondary structural elements amongst many THAP proteins

The THAP domain has been experimentally demonstrated to fold into a *βαβ* (*β*1-L1-*β*2-L2-*α*1-L3-*β*3-L4) secondary structural fold in hTHAP1 (2JTG, 2KO0), hTHAP2 (2D8R), hTHAP11 (2LAU), DmTNP (3KDE) and C. elegans CtBP-1 (2JM3). The secondary structure predictions of the THAP domains of each of the twelve human THAP proteins using JPRED, PSIPRED and SPIDER3 agree with the experimentally identified β; secondary structural fold. Figure 2A and 2B displays results from JPRED. Surprisingly, an additional *β* sheet (*β*4, blue) of length more than or equal to three amino acids residues was predicted within the L4 regions in hTHAP1, hTHAP4, hTHAP9 (Figure 2A) albeit with a very less confidence score. The predicted *β*4 in hTHAP2, hTHAP5, hTHAP6, hTHAP8, hTHAP10 was not considered as it was less than 3 residues long and a typical *β* sheet is made of 3 -10 residues (33).

**Figure 2.**
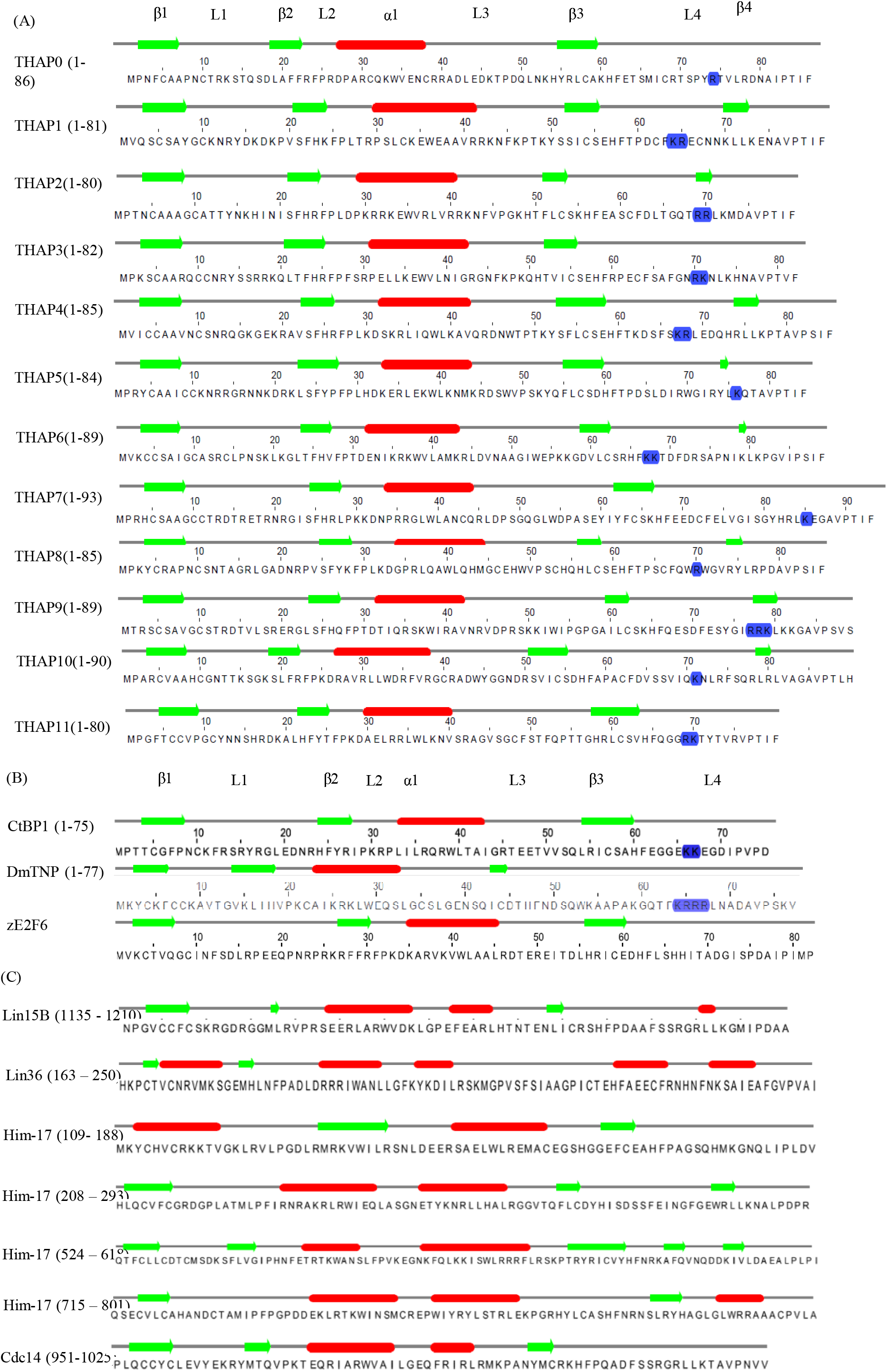
JPRED predicts a consensus structural fold of the THAP domain. Predicted □□□ fold for THAP domains of (A) all twelve hTHAP proteins (B) other THAP proteins (DmTNP, zE2F6, CtBP) (C) predicted structural fold different than consensus □□□ fold for the THAP domains of CDC14, Lin15B, Lin36 and Him-17.

Surprisingly, in the “other THAP protein” group, only C.elegans CtBP-1, DmTNP and zE2F6 had a predicted *βαβ* secondary structural fold in their THAP domains (Figure 2B). Cdc14, Lin-15B and Him17 (3) had an additional short helix between the helix and *β* sheet whereas Lin-36, Him17(1), Him17(2), Him17(4) were predicted to have distinct structural folds with extra helices and sheets (figure 2C).

Interestingly, the length of *β*1(5 residues) and □ 1(10 residues) was conserved in the human THAP family (figure 2A) as well as CtBP-1, DmTNP and zE2F6 (figure 2B). On the other hand, the length of *β*2 and *β*3 varied slightly (figure 2). For example, *β*2 consists of five (hTHAP3, hTHAP5, DmTNP) or four (other hTHAP proteins, CtBP-1, zE2F6) residues while *β*3 consists of two (DmTNP), three (hTHAP2, hTHAP8, hTHAP9), four (zE2F6, hTHAP1, hTHAP3, hTHAP6), five (hTHAP7, hTHAP0, hTHAP 5, hTHAP10) or six (hTHAP11, hTHAP4, CtBP-1) residues.

### Loop 4 length and sequence diversity and its correlation with DNA binding specificity

The length and sequence of L4 has been speculated to be important for different DNA binding specificities for different human THAP proteins (3, 18, 19). This is more significant in the light of structural studies that demonstrate direct interactions between the basic residues in L4 and the pyrimidine ring of a thymine base in their respective DNA binding sites (Arg65, Arg66 and Arg67 and in DmTNP, Arg65 in hTHAP1) (17, 18).

DmTNP has a stretch of four consecutive basic amino acids (Lys64, Arg65, Arg66 and Arg67; figure 2B) in L4. Interestingly, in hTHAP9, three consecutive basic residues (Arg77, Arg78, Lys79) were predicted in L4-*β*4 (figure 2A, highlighted in blue). However, the L4 may also contain two consecutive basic residues as observed in THAP1 (Lys64, Arg65), THAP2 (Lys69, Lys70), THAP3 (Arg70, Lys71), THAP4 (Lys67, Arg68), THAP6 (Lys67, Lys68), THAP11 (Arg69, Lys70), CtBP-1 (Lys66, Lys67) or one basic residue as seen in THAP0, THAP5, THAP7, THAP8, THAP10 (highlighted in blue, figure 2). zE2F6 does not have a single basic residue in the predicted L4 (figure 2B). This raises the possibility that one basic residue in L4 might suffice for minor groove DNA interaction if complemented with another basic residue from another structural fold to form a positively charged surface around the DNA as has been previously suggested (3).

It has been speculated that the length of L4 determines DNA binding affinity: the longer the L4, the tighter the binding (18, 19). However, there are no experimental reports establishing this claim. Thus, it was interesting to observe significantly different lengths of L4 within the human THAP family of proteins (Table 1). It is tempting to hypothesize that hTHAP10 may have the strongest while hTHAP4 may have the weakest interaction with DNA.

**Table 1.**

| Human THAP protein | L4 length (residues) |
| --- | --- |
| hTHAP0 | 26 |
| hTHAP1 | 16 |
| hTHAP2 | 26 |
| hTHAP3 | 26 |
| hTHAP4 | 14 |
| hTHAP5 | 24 |
| hTHAP6 | 27 |
| hTHAP7 | 23 |
| hTHAP8 | 26 |
| hTHAP9 | 15 |
| hTHAP10 | 35 |
| hTHAP11 | 17 |

### Diversity in the THAP domain features in the homologs of hTHAP proteins

The THAP domain is a novel protein domain restricted to animals. The THAP domains of human THAP proteins appear to share structural similarity (Fig.2) and conserved invariant residues. Studying the THAP domain features in the homologs of human THAP proteins may provide insights into the evolution of THAP domains in humans. Thus, protein sequences of homologs of each human THAP protein were extracted from the NCBI PROTEIN database.

#### A. The THAP domain can be longer or shorter than 90 residues

The THAP domain has been reported to be between 80-90 residues long, as demonstrated by structural analysis of the THAP domains of hTHAP1 (2JTG, 2KO0), hTHAP2 (2D8R), hTHAP11 (2LAU), DmTNP (3KDE), CtBP (2JMR. However, analysis of individual hTHAP protein homologs in different organisms revealed proteins with variable THAP domain length. For the purpose of this study, THAP domains of hTHAP protein homologs which are longer than 100 residues have been termed “Long THAP domains” (Table 2) and those with length less than or equal to 50 residues have been termed “Short THAP domains” (Table 3). It was interesting to observe that these homologs with long THAP domains often had high sequence similarity, as noted below:

**Table 2.**
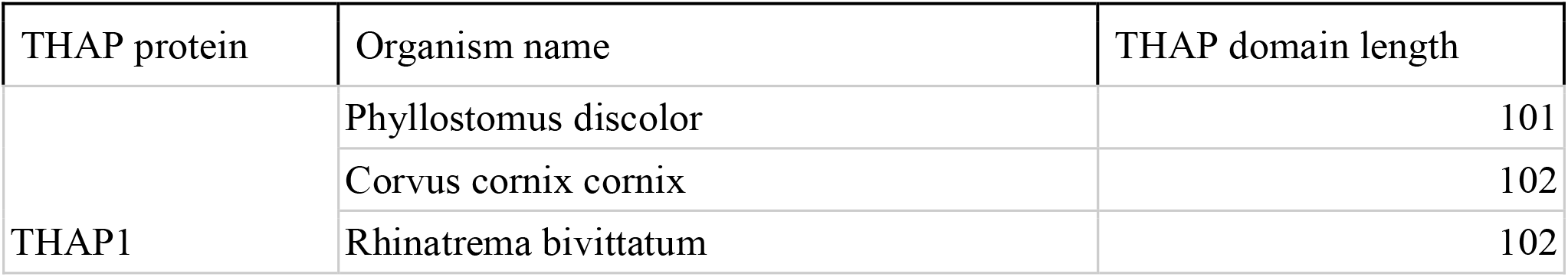
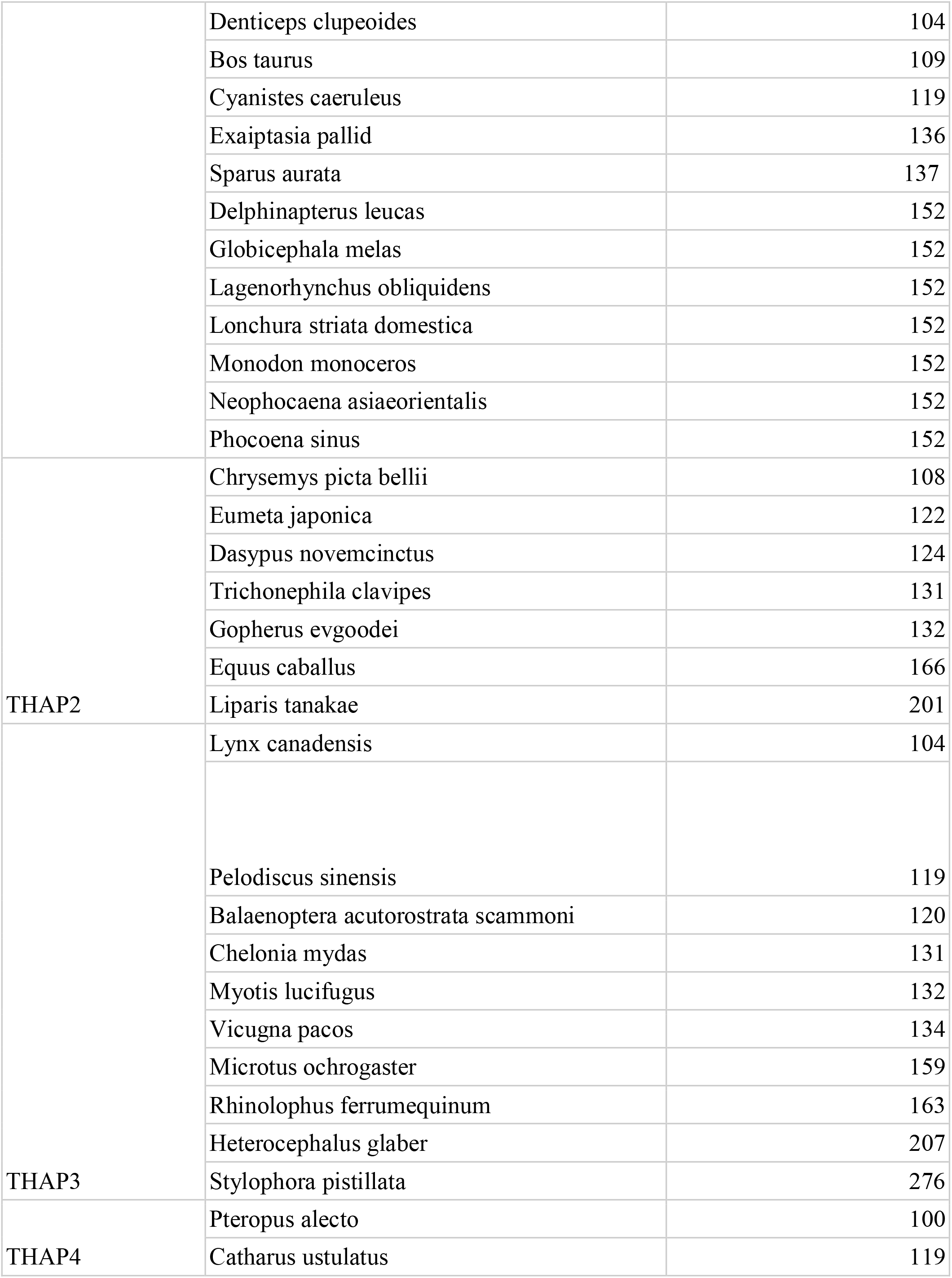
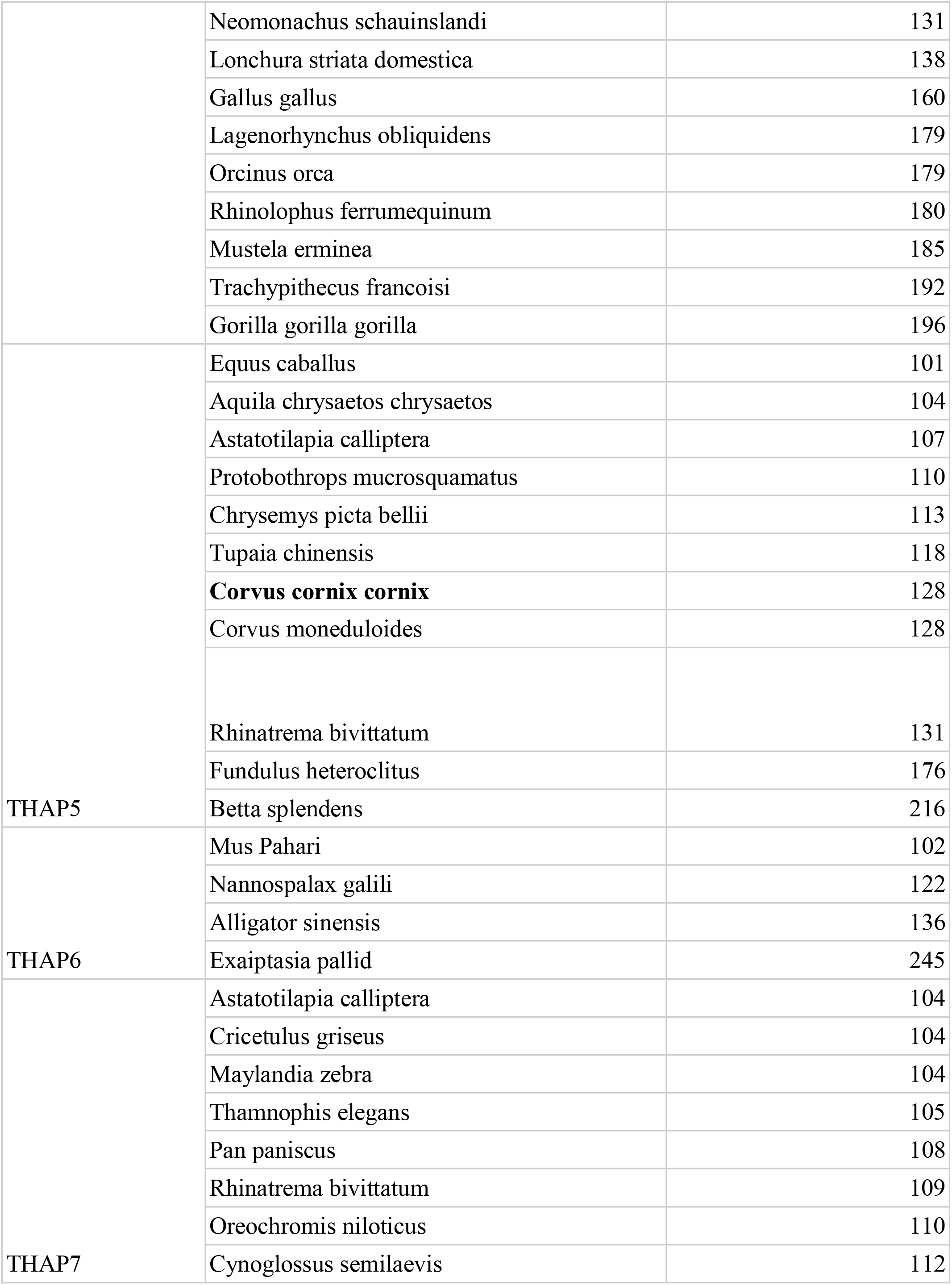
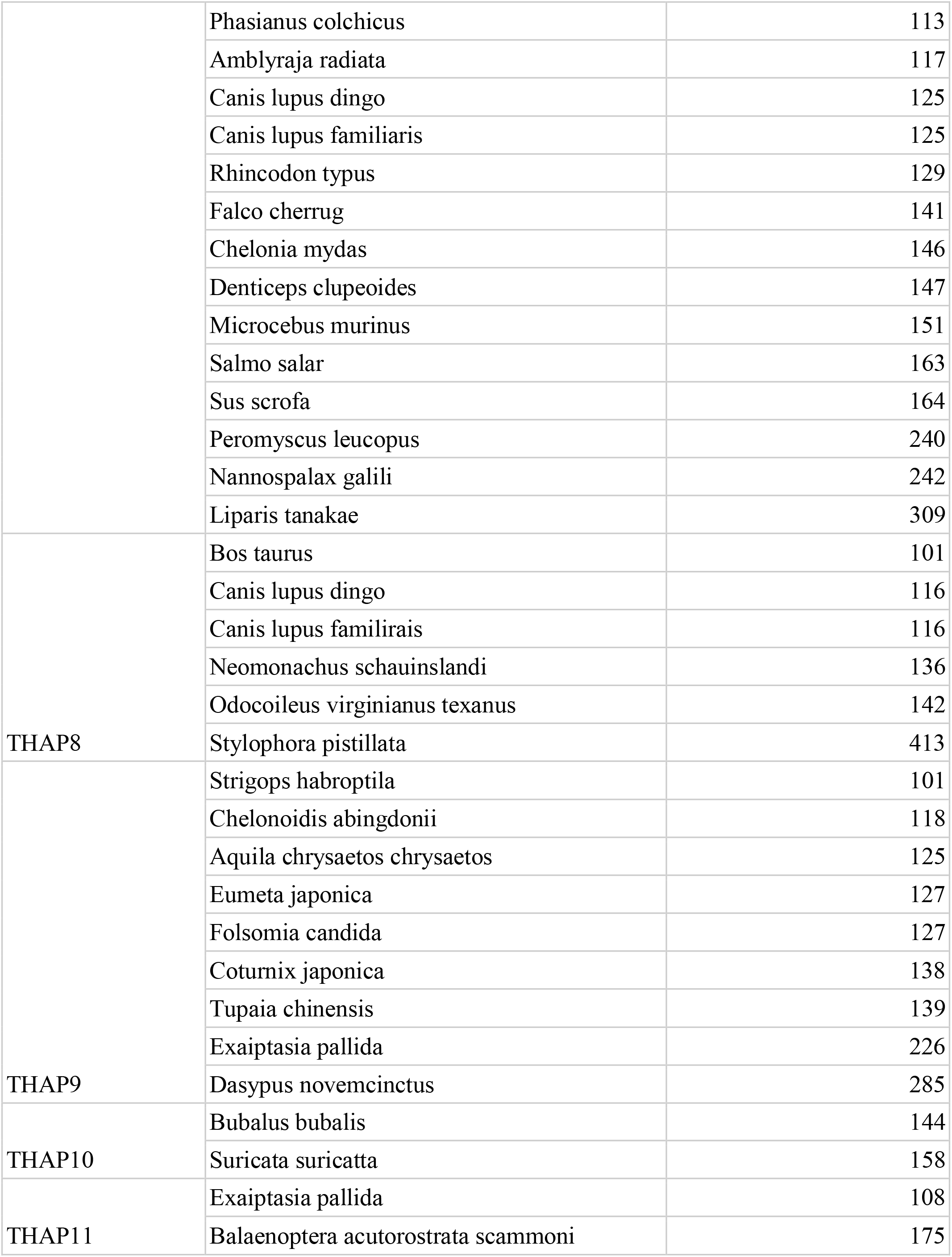

**Table 3.**

| THAP protein | Organism name | THAP domain length |
| --- | --- | --- |
| THAP3 | Alligator sinensis | 39 |
|  | Pteropus alecto | 42 |
| THAP6 | Acinonyx jubatus | 48 |
|  | Myotis lucifugus | 48 |
| THAP4 | Ictidomys tridecemlineatus | 31 |
|  | Bubalus bubalis | 50 |
|  | Odocoileus virginianus texanus | 50 |
| THAP5 | Amblyraja radiata | 42 |
|  | Antrostomus carolinensis | 42 |
|  | Columba livia | 42 |
|  | Falco cherrug | 42 |
|  | Falco peregrinus | 42 |
|  | Geospiza fortis | 42 |
|  | Meleagris gallopavo | 42 |
|  | Oreochromis niloticus | 42 |
|  | Pelodiscus sinensis | 42 |
|  | Tyto alba alba | 42 |
|  | Armadillidium vulgare | 46 |
| THAP8 | Globicephala melas | 42 |
|  | Terrapene carolina triunguis | 42 |
|  | Tyto alba alba | 42 |
|  | Microtus ochrogaster | 45 |

1. hTHAP1: 7 (6 mammalian, 1 aves) out of 16 homologs had <u>152</u> residue long THAP domain (Figure 3A). All 6 mammalian homologs had identical sequences except for two amino acid residues (Suppl. Fig. 2A). However, the aves homolog varied considerably from the mammalian homologs (Suppl Fig. 3).
2. hTHAP4: 2 (mammalian) out of 11 homologs had <u>179</u> residue long identical THAP domains (Figure 3B. Suppl. Fig. 2A).
3. hTHAP5: 2 (avian) out of 11 homologs had <u>128</u> residue long THAP domain (Figure 3C) which are identical except for two residues (Suppl. Fig. 2A).
4. hTHAP7: (a) 2 (1 Actinopterygii, 1 mammalian) out of 22 homologs had <u>104</u> residues long THAP domain. Both these homologs had identical THAP domain sequences (b) 2 (mammalian) out of 22 homologs had <u>125 residue</u> long identical THAP domains (Figure 3D, Suppl. Fig. 2A).
5. hTHAP8: 2 (mammalian) out of 6 homologs had <u>116</u> residue long identical THAP domains (Figure 3E, Suppl. Fig. 2A).
6. hTHAP9: 2 (1 insecta, 1 mammalian) out of 9 homologs had <u>127</u> residue long THAP domains (Figure 3F), both of which were significantly different from each other (Suppl. Fig. 2A, highlighted in grey).

Some interesting observations about hTHAP protein homologs with short THAP domains include

1. hTHAP3:Chinese alligator (Alligator sinensis) has 39 aa long THAP domain and black flying fox (Pteropus alecto) has 42 aa long THAP domain with deletion before the conserved F59.
2. hTHAP4: 2 (mammalian: white-tailed deer (Odocoileus virginianus texanus) and water buffalo (Bubalus bubalis) out of 3 homologs had <u>50</u> residue long THAP domains (Figure 3G, Suppl. Fig. 2B) with identical sequences except for two residues (Suppl. Fig. 2B), deletion before the conserved W38; thirteen-lined ground squirrel (Ictidomys tridecemlineatus) have a 31 aa long THAP domain with deletion before conserved H59.
3. hTHAP5: 10 (7 aves, 1 reptilia, 1 actinopterygii, 1 chondrichthyes: thorny skate (Amblyraja radiata), nile tilapia (Oreochromis niloticus), medium ground finch (Geospiza fortis), barn owl (Tyto alba alba), chinese softshell turtle (Pelodiscus sinensis), rock dove (Columba livia), peregrine falcon (Falco peregrinus), saker falcon (Falco cherrug), chuck-will’s-widow (**Antrostomus carolinensis**) and wild turkey (Meleagris gallopavo)) out of 11 homologs had <u>42</u> residue long THAP domain (Figure 3H, Suppl. Fig. 2B) which had 70% identity and deletion before conserved C57. 46 aa long THAP domain in common pill bug/potato bug (Armadillidium vulgare).
4. hTHAP6: Both mammalian homologs(little brown bat (Myotis Lucifugus) and cheetah (Acinonyx jubatus) had <u>48</u> residue long THAP domain (Figure 3I) which were identical except for two residues (Suppl. Fig. 2B) with deletion before the conserved C62
5. hTHAP8: 3 (mammalian, reptilia, aves) out of 4 homologs had <u>42</u> residue long THAP domain (Figure 3J) which had 74% identity (Suppl. Fig. 2B) with deletion before the conserved C58.The prairie vole (**Microtus ochrogaster**) was also found to have 45 aa long THAP domain with deletion before conserved C58.

**Figure 3.**
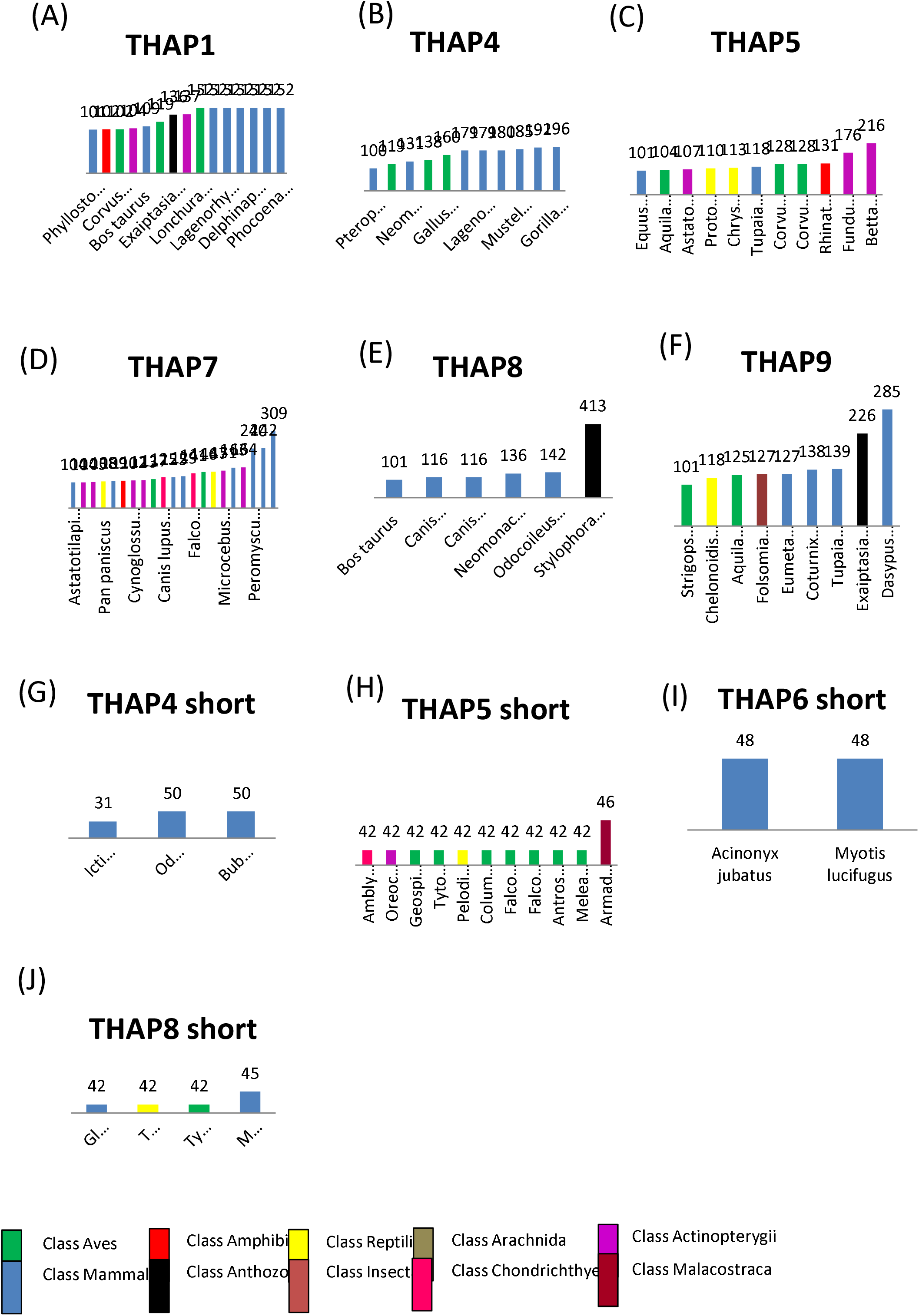
Comparison of THAP domain length among hTHAP homologs. Graphical representation of THAP domains of Long THAP domain containing homologs of (A) THAP1 (B) THAP4 (C) THAP5 (D) THAP7 (E) THAP8 and Short THAP domain containing homologs of (F) THAP4 (G) THAP5 (H) THAP6 (I) THAP8. Y axis represents THAP domain length in residues and X axis represents the scientific name of the organism. The vertical bars are colored according to the taxonomic class of organisms (color key in bottom panel).

These interesting similarities in lengths and sequence of the THAP domain amongst the human THAP protein homologs led us to ask if the homologs with identical or similar THAP domains had sequence similarity beyond the THAP domain. That is, if the homologs with identical THAP domains were identical across the entire length of the protein. We could identify examples of three different possibilities. These are as follows:

1. Sequence similarity across the length of the entire protein. Example: 6 mammalian 152 residue long THAP domain containing THAP1 homologs, 2 mammalian 179 residue long THAP domain containing THAP4 homologs, 2 mammalian 128 residue long THAP5 THAP domain containing homologs, 2 mammalian 125 residue long THAP domain containing THAP7 homologs, both (1 mammalian, 1 actinopterygii) THAP7 homologs with 104 residues long THAP domain, 2 mammalian THAP8 homologs with 116 residue long THAP domain, 2 mammalian THAP4 homologs with 50 residue long THAP domain, 7 avian 42 residues long THAP domain containing THAP5 homologs and 2 mammalian 48 residue long THAP domain containing THAP6 homologs.
2. Sequence similarity only within the THAP domain. Example: 3 THAP8 homologs (1 mammalian, 1 avian and 1 reptilian) with 42 residue long THAP domain.
3. No sequence similarity across the length of the protein, i.e. within THAP domain and beyond THAP domain. Example: The THAP9 homologs with 127 residues (1 mammalian, 1 insecta) long THAP domain.

### Variations in the C2CH zinc coordinating motif of the THAP domain

It was seen that certain homologs did not align with their hTHAP counterpart. Thus, we aligned these homologs separately and surprisingly, found some of these were completely identical to each other. Some such examples of hTHAP protein homologs are mentioned below.

4 types of variations in the C2CH motif.

*2 Cys in amino-terminal region, which do not align with C5 and C10 in hTHAP homolog:* hTHAP3 homologs in desert tortoise (*Gopherus evgoodei*), painted turtle (*Chrysemys picta bellii*), pinta island tortoise (*Chelonoidis abingdonii*), three-toed box turtle (*Terrapene carolina triunguis*), hawaiian monk seal (*Neomonachus schauinslandi*), weddell seal (*Leptonychotes weddellis*).

*Lacks 1st Cys in the first 6 residues:* hTHAP6 homologs in desert tortoise (*Gopherus evgoodei*), pinta island tortoise (*Chelonoidis abingdonii*), which had completely identical THAP domain except for 3 aa.

*Lack 2nd Cys in between 6th to 12th residues:* hTHAP3 homolog in *Terraprene* (three-toed box turtle), in the completely identical long THAP domains (except for two residue). hTHAP1 homologs of narrow-ridged finless porpoise (*Neophocaena asiaeorientalis*), Pacific white-sided dolphin (*Lagenorhynchus obliquidens),* Beluga whale (*Delphinapterus leucas*), Vaquita (*Phocoena sinus*), narwhal (*Monodon monoceros*) and long-finned pilot whale (*Globicephala melas*).

*Lack both Cys in the first 12 residues:* hTHAP6 homologs in wild Bactrian camel (***Camelus ferus***) and alpaca (*Vicugna pacos*); hTHAP4 homologs in chinese softshell turtle (*Pelodiscus sinensis*), golden-collared manakin (*Manacus vitellinus*), cheetah (*Acinonyx jubatus*) and cat (*Felis catus*); hTHAP9 homologs in rohu (Labeo rohita), treeshrew (Tupaia chinensis), Okarito kiwi (**Apteryx rowi**), japanese quail (Coturnix japonica), Kanglang fish (**Anabarilius grahami**), springtail (Folsomia candida), nine-banded armadillo (Dasypus novemcinctus) and bagworm moths (Eumeta japonica).

The GLAM2 predicted gapped motif within the long THAP domains in the homologs of hTHAP0,hTHAP1, hTHAP4,hTHAP5, hTHAP6, hTHAP8 and hTHAP9 did not have a predominant conservation of two Cys residues (Fig. 6A) within the first 40 residues of the C2CH motif. However, one Cys residue (hTHAP3 and hTHAP7) or two Cys residues (hTHAP2) were seen to be predominantly conserved in the long THAP domain homologs (Fig. 6A). GLAM2 predicted motifs for the long THAP domains in hTHAP10 and hTHAP11 homologs cannot be considered because there are only two long THAP domain containing homologs of THAP10 and THAP11.

**(C) Length of the THAP domain has no correlation with total protein length**

We then asked if there was any correlation between the THAP domain length and the total protein length. However, there was no correlation between the THAP domain length to the total protein length. For example, the full length THAP proteins with long THAP domains were not longer than the proteins with either short or canonical ∼90 residue longTHAP domains (THAP3, THAP4, THAP5, THAP6, THAP7, THAP8 in figure 4).

**Figure 4.**
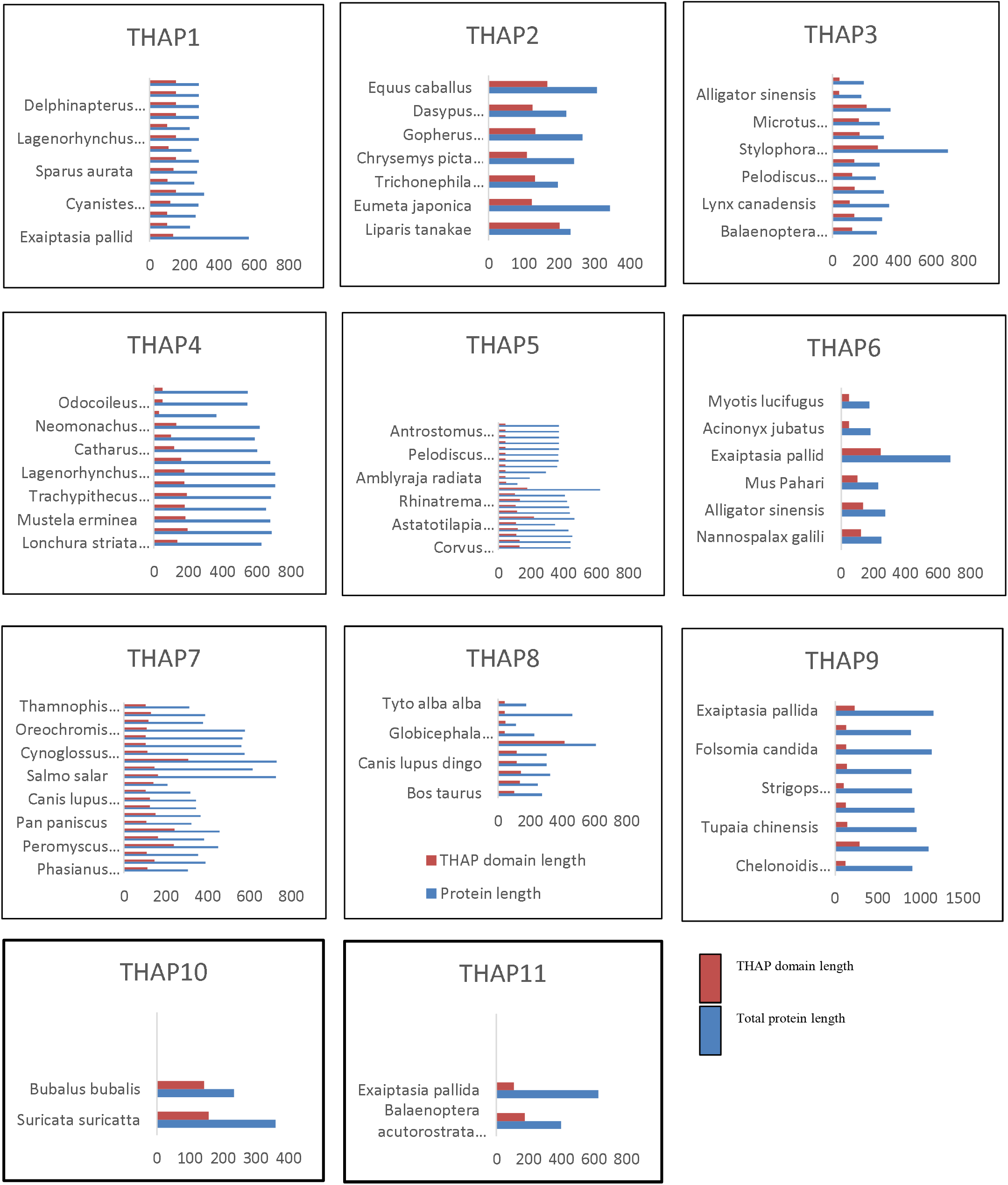
Correlation between THAP domain length and total protein length. Graphical representation of comparison of THAP domain length with total protein length of long and short THAP domains. X axis represent the length of the THAP domain (orange) and length of the total protein (blue) and Y axis represent the scientific name of the organism.

### Amino terminal end of THAP domain is a probable hot spot for insertion or deletion within the THAP domain

The diverse lengths of THAP domains in hTHAP homologs intrigued us to look for a region within the THAP domain which was conserved across the THAP domains of different lengths. Comparison of gapped motifs of all the long THAP domains (Figure 5A) and their canonical ∼90 aa THAP domains for each hTHAP protein homolog (figure 5B) revealed that the carboxy terminal region was found to be more or less similar between the long THAP domains and the ∼90 residues long THAP domains.

**Figure 5.**
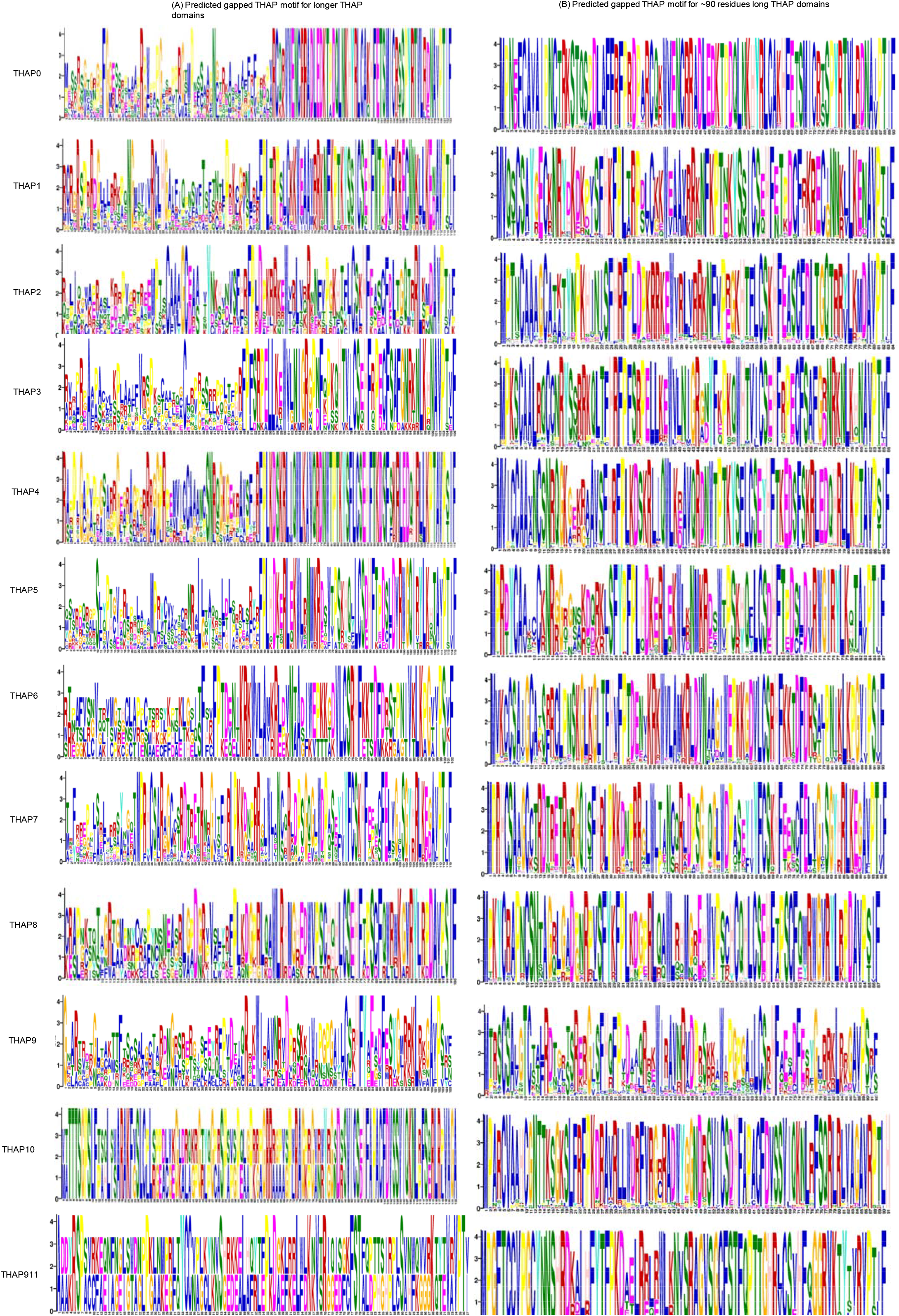
Comparison of GLAM2 predicted motifs of long THAP domains with their conventionally ∼ 90 residues long THAP domain. Different gapped motifs were predicted for (A) long THAP domains and (B) conventional ∼90 residues long THAP domains in each of the twelve human proteins. The GLAM2 predicted gapped motifs are represented as PWMs.

It was interesting to see that the amino terminal region of the THAP domain, i.e the region from 1st aa to the conserved W residue within the C2CH motif, served as a probable hotspot for insertions within the THAP domain. This observation holds true amongst the homologs of hTHAP1, hTHAP2, hTHAP4, hTHAP6, hTHAP10 and hTHAP11 which had THAP domains longer than their protein counterparts with ∼90 residue long THAP domains. All these long THAP domain containing homologs had insertions immediately after the 1st Met or at specific positions such as (hTHAP1 before conserved P26 in loop 2), hTHAP2 (before conserved P26 in loop 2), hTHAP3 (before conserved P27 in loop 2), hTHAP4 (before conserved P28 in loop 2), hTHAP5 (immediately after conserved P27 in loop 2), hTHAP6 (before conserved P29 in loop 2), hTHAP7 (before conserved Loop 2 P29 in Thamnophis elegans, Canis lupus dingo and Canis lupus familiaris but before P52 in loop 3 for other 18 homologs), hTHAP8 (immediately before conserved W40 in *α*1), hTHAP9 (immediately before conserved W38 in *α*1), hTHAP10 (before conserved C54 in *β*3) and hTHAP11 (immediately before conserved W36 in *α*1).

In addition to the amino terminal region being a hot spot for insertions, it also serves as a hot spot for deletions. All the short THAP domain-containing homologs of hTHAP3, hTHAP4, hTHAP5, THAP6 and hTHAP8 proteins were found to have a deletion before the conserved CH of the C2CH motif (loop 4). The only exception was the Ictidomys tridecemlineatus homolog of hTHAP4 which had a THAP domain with a deletion till immediately before the conserved His residue of the C2CH motif (loop 4). These predictions also coincide with the earlier described DM3 domain. It is a truncated THAP domain which has a deletion of the first 20 residues at the amino terminal region (6).

It was also interesting to see that even the homologs of hTHAP proteins with longTHAP domains (figure 5A) and short THAP domains had at least one basic amino acid in the predicted L4 region.

### Insertions in the long THAP domain containing hTHAP protein homologs do not have a consensus sequence and secondary structure fold

GLAM2 was used to separately align the set of long (Figure 5A) or canonical (90 residue; Figure 5B) THAP domains belonging to the homologs of each hTHAP protein. It was observed that the gapped motif generated using the long THAP domain sequences from 15 hTHAP1 homologs shows a lot of variation at the amino terminus (place where insertions are seen within the THAP domain) (Figure 5A) as the insertion sequences differ amongst themselves (suppl figure 4). Moreover, long THAP domains either lack two C residues of the C2CH motif with the exception of THAP2 or have at least one C residue intact at the amino-terminus (Figure 5A) in THAP3, THAP10 and THAP11 .The carboxy terminal region of the long THAP domains i.e. the region from W in the C2CH motif was found to be more conserved than the amino terminal region (Figure 5A).

In addition to these, specific residues were conserved within the insertion sequences of hTHAP protein homologs. For instance, in homologs of hTHAP0 (One Phe, one Gly, one Arg, one Pro one Asn), hTHAP1 (Three Arg, one Asn, one Gly, one His, one Pro), hTHAP2 (One Gly, one Val), hTHAP3 (One Ala), hTHAP4 (Three Arg, three Gly, two Pro, one Ser, one Asn), hTHAP5 (One Ser, one Leu), hTHAP6 (One Phe), hTHAP8 (One Asp), hTHAP9 (One Arg). As only two homologs of hTHAP10 and hTHAP11 had longTHAP domains, we don’t comment on any conserved residues in the insertion sequences of these THAP domains.

Based on the variations in the sequences of insertions in the long THAP domains, it was logical to check if there was a consensus structural fold that was formed by these insertions as that might explain a similar functional role of the long THAP domain containing proteins. However, insertions in each long THAP domain had a different sequence (10% similarity) and secondary structure, despite being the homologs of the same hTHAP protein. For example, the THAP domains of Lonchura and Monodon homologs of hTHAP1 had different predicted secondary structural folds despite being of the same length (Suppl figure 4).

## Discussion

Most proteins contain more than one domain which are usually functionally independent. Typically, a protein domain folds into a core structural motif independent of the rest of the protein and is reported to be more conserved at the tertiary structure level than at the amino acid sequence level (34, 35). Evolutionary divergence or evolutionary conservation of a protein domain is an outcome of the combination of random mutations and selection restrictions imposed on the function of the domain (34, 35). The THAP domain is a C2CH zinc coordinating DNA binding domain. It is an example of the protein domain shared between DNA binding proteins and DNA transposase (DmTNP) which is active in the host organism. Several other such domains which are shared by different proteins are known. For example, BED zinc finger DNA binding domain which is shared by chromatin boundary element binding proteins (BEAF, DREF) and AC1 and Hobo-like transposases in fungi, plants and animals.

The GLAM2 predicted gapped motif within the THAP domain of all the human THAP proteins illustrates the conservation of cys, pro, trp and his residues of the C2CH motif, a phe immediately flanking the C2CH motif and a pro in the AVPTIF motif of the THAP domain. We report three additional conserved residues (Phe, Arg within the C2CH motif and a Leu in the L4 region) in the THAP domains of all the twelve human THAP proteins. We also report that the conserved residues within the THAP domain of human THAP proteins (figure 1A) vary from the conserved residues within the THAP domain of earlier reported other THAP domain containing proteins (Figure 1B).

The THAP domain is reported to fold into a consensus *βαβ* fold despite differences in the amino acid sequence of THAP domains in different proteins. There have been no reports of a fourth beta sheet in the THAP domains of hitherto reported structures. We report the presence of a fourth beta sheet in the predicted secondary structures of hTHAP1, hTHAP4, hTHAP9 proteins. Although, the confidence in the prediction of the fourth beta sheet is very less.

L4 length and sequence variations have been earlier speculated to impose functional diversity in the binding sites of different human THAP proteins. Our sequence and secondary structure analysis provides more confidence to these speculations. We also report that except for hTHAP0, all the twelve human THAP proteins had at least one basic amino acid residue in their respective L4 regions. This is important because the DmTNP structure demonstrates that these basic amino acid residues form direct contact with the nitrogen base in the minor groove of the DNA. However, further experimental studies are required to establish if only one basic residue in the L4 is enough to bind DNA.

We report for the first time that there are THAP domains of lengths different than the conventionally considered 90 residues. Although there is no correlation between the length of the THAP domain and the length of the protein, it was interesting to see that THAP homologs from different taxonomic classes have THAP domains of similar lengths. For example, THAP1 homologs in class aves [Bengalese finch (Lonchura striata domestica)] and in class mammalia [long-finned pilot whale (Globicephala melas)] had 152 residue long THAP domain (Figure 3). It was interesting to observe that even in the same organism, the THAP domain appears to have evolved differently. For example, in two-lined caecilian (Rhinatrema bivittatum), the THAP domain is either 102 residues (THAP1), 131 residues (THAP5) or 109 residues (THAP7) (Figure 3). Similarly, the sea anemone (Exaiptasia pallid) THAP domain is either 136 residues (THAP1), 245 residues (THAP6), 226 residues (THAP9) or 108 residues (THAP11). Interestingly, the long-finned pilot whale (Globicephala melas) has both a 152 residue long THAP domain (THAP1) and a 42 residue short THAP domain (THAP8).

The genus and/or habitat specific THAP domain sub-groups reported in figure 4 suggest the possibility of evolution of THAP domain in a protein specific manner. For example, the marine animal specific THAP domain (length and sequence) in THAP1 homologs may be due to the specific function of THAP1 in these animals. This speculation becomes more interesting because these marine animals specific hTHAP1 homologs are very similar throughout the length of these proteins and not just within the THAP domain. The THAP3 (1) shows a possibility of habitat specific evolution of THAP domain in THAP3 homologs in tortoise where the THAP domain of desert tortoise has an insertion of seven residues as compared to the THAP domain of the island tortoise. The long THAP domain of hTHAP1 homologs can be studied *in silico* by docking it with the hTHAP1 DNA binding site (AGTACGGGCAA). These studies will provide preliminary understanding about the preference of DNA binding site of long THAP domains, whether these domains bind with similar affinity to the DNA sequences and/or if the structural fold changes after binding the DNA. Simulation studies with this complex will provide some understanding about the mode of interaction of the long THAP domain with DNA, if any.

A conserved carboxy terminal region of the longer THAP domains despite varying sequences hints at an evolutionarily important function of the carboxy terminal region of the THAP domain. Counter-intuitively, the variations in the amino terminal region (C2CH zinc coordinating motif) of the longer THAP domains offers a strong possibility of the co-adaptation of the DNA binding function of the zinc finger motif into a different function. This also states that there are other DNA binding motifs that can replace this function of the THAP domain and thus making it redundant and costly to be maintained over the course of evolution of THAP domain in different organisms.

A truncated THAP domain which lacks the first 20 residues including the two C of C2CH motif has been reported by the SMART database earlier as the DM3 domain (6). However, what we report in this study as short THAP domains in THAP3, THAP4, THAP5, THAP6 and THAP8 homologs lack ∼40 residues at the extreme N terminus of the THAP domain. As this deletion includes the first two C of the C2CH motif, the conserved P and W residues within the C2CH motif, it is tempting to speculate that these proteins may not bind DNA. However, the presence of the rest of the THAP domain region which includes the CH of the C2CH motif, the conserved F residue adjacent to the H of C2CH motif and the AVPTIF motif leads to ask questions like do these proteins bind DNA with less affinity or bind to non-specific DNA regions? It would also be very interesting to study if these proteins play any role in the regulation of DNA binding.

The difference in the predicted secondary structural fold of other THAP domains containing proteins leads us to speculate about the possibilities of a different structural fold for THAP domains other than the consensus fold. It will be very interesting to study these exceptions/outliers as the sequences of these THAP domains are also very different from that of the THAP domains of hTHAP proteins and its homologs apart from the conserved amino acid residues. Further biochemical and structural studies are required to study the evolutionary basis of variation in the sequence.

## Figure legends

**Supplementary figure 1.**
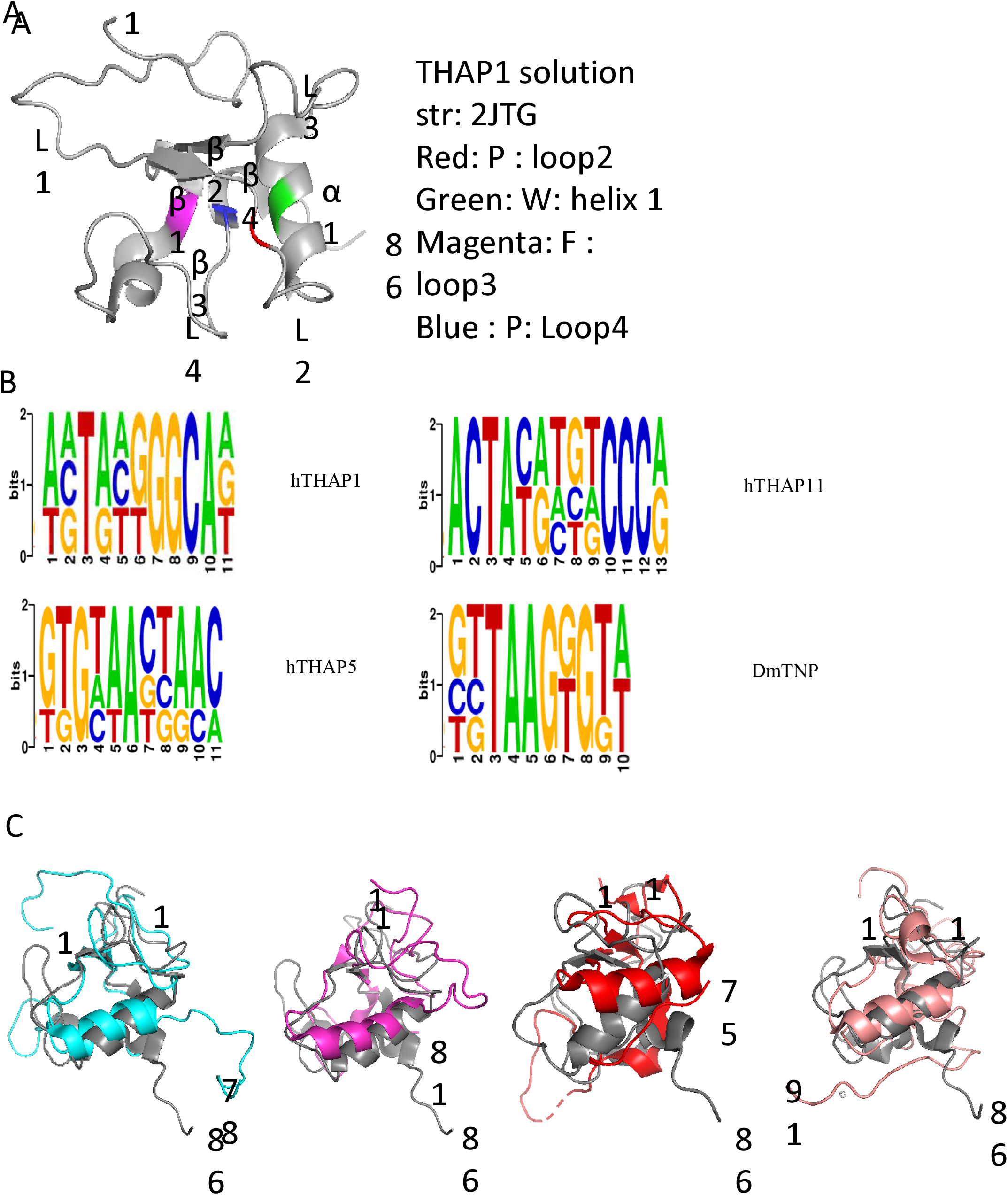
Conserved sequence and structure signatures of THAP domain. (A) The solution structure of THAP domain of human THAP1 (PDB ID: 2JTG) as viewed in Pymol. The conserved P in the C2CH motif is in the loop2 (highlighted in red), the conserved W within the C2CH motif is in the first alpha helix (highlighted in green), the conserved F immediately after the H of C2CH motif is in the beginning of loop3 (highlighted in magenta) and the conserved P of AVPTIF motif is in the loop4 (highlighted in blue). (B) DNA binding sites of different THAP domain containing proteins as created by Weblogo (36). (C) Pymol representations of structural alignment of THAP domains of hTHAP1(gray) with hTHAP2 (blue), hTHAP11 (magenta), DmTNP (red), CtBP-1 (peach).

**Supplementary figure 2.**
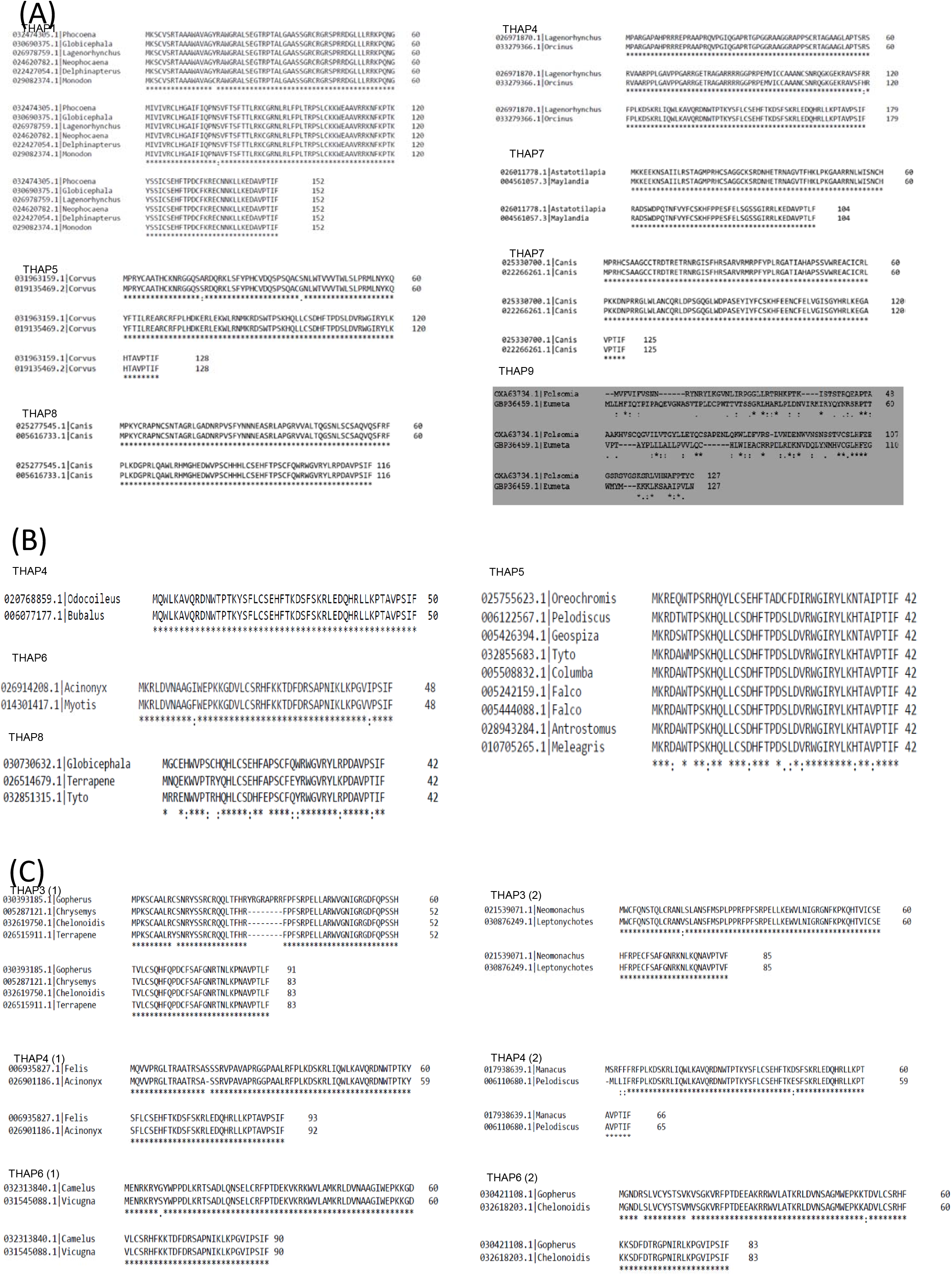
Length based sequence identity of hTHAP homologs. (A) Identical long THAP domain containing homologs of hTHAP1, hTHAP4, hTHAP5, hTHAP7 and hTHAP8. (B) Identical short THAP domain containing homologs of hTHAP4, hTHAP5, hTHAP6 and hTHAP8. (C) ∼90 residues long similar THAP domain containing homologs of hTHAP1, hTHAP3 and hTHAP6 which did not align with their human THAP domain counterpart.

**Supplementary figure 3.**
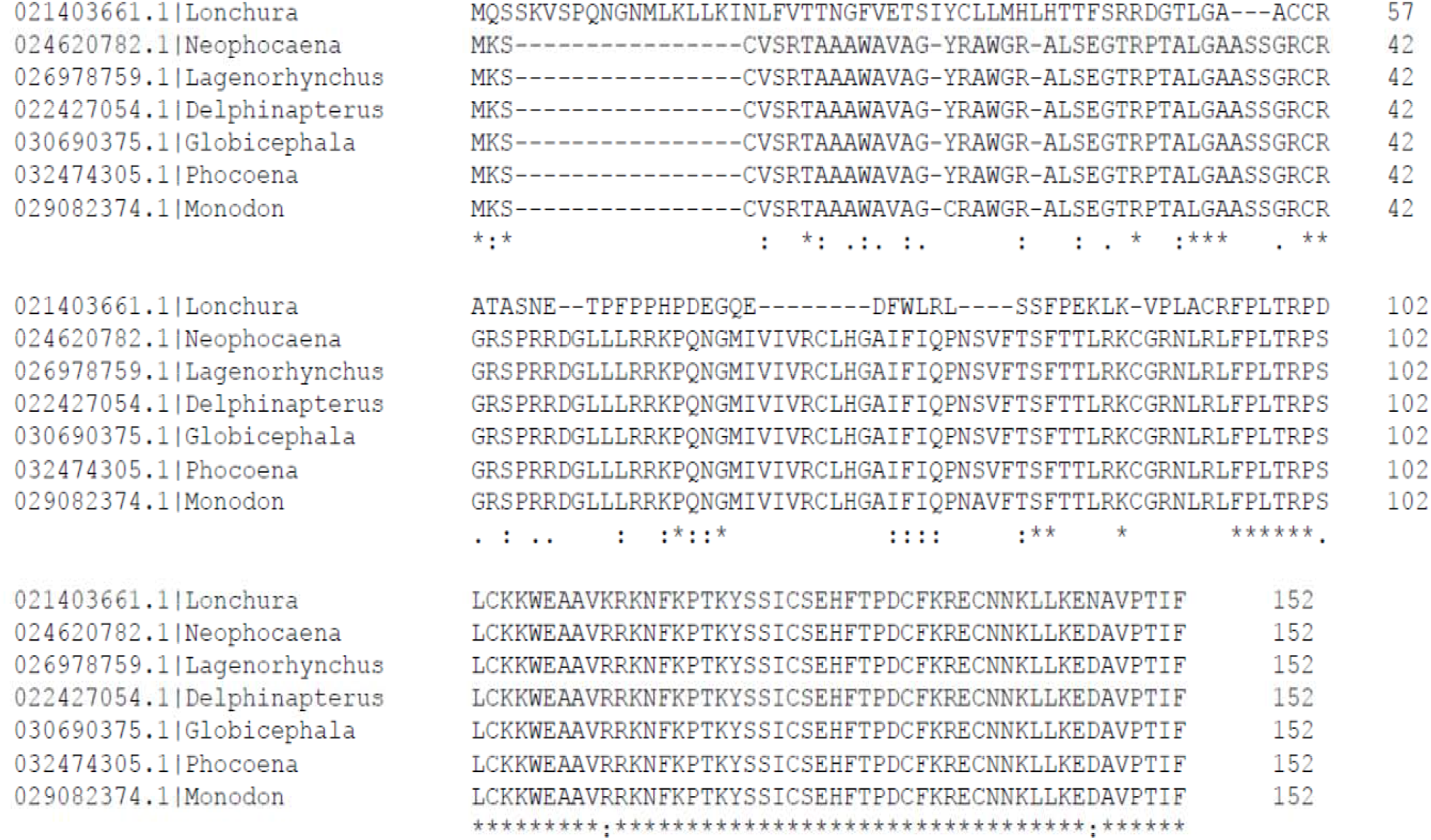
Sequence variation in the 152 residues long THAP domain containing avian and mammalian homologs of hTHAP1.

**Supplementary figure 4.**
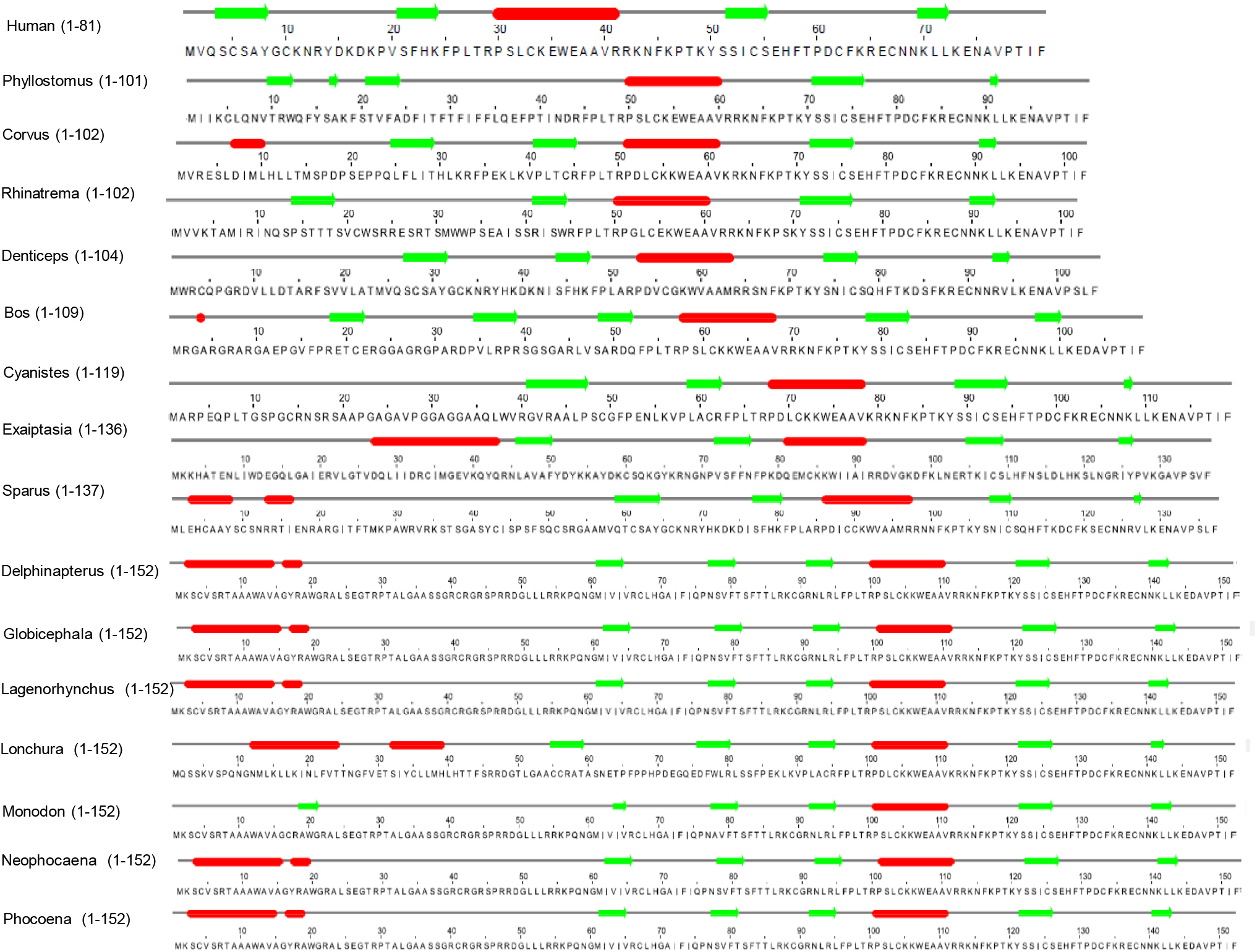
Predicted secondary structural fold of longer THAP domain homologs of hTHAP1. JPRED predicts different secondary structural fold for the insertion regions (regions before W of C2CH motif) of each of the sixteen longer hTHAP1 domain homologs.

## Declarations

### Ethics approval and consent to participate

Not applicable

### Consent for publication

Not applicable

### Availability of data and material

All data generated or analysed during this study are included in this article (and its supplementary files).

## Authors’ contributions

HMS initialized the study and generated the data. HMS & SM analyzed the data. HMS & SM wrote the manuscript. All authors read and approved the final manuscript.

## Acknowledgements

We strongly acknowledge Dr. Sairam S. Mallajosyula, Dr. Althaf Shaik and Richa Rashmi for providing suggestions and reviews about the study.

