## Additional file for "Synapomorphic variations in the THAP domains of the human THAP protein family and its homologs": THAP0.docx

>PNI56822.1 Pan troglodytes

MPNFCAAPNCTRKSTQSDLAFFRFPRDPARCQKWVENCRRADLEDKTPDQLNKHYRLCAKHFETSMICRT

SPYRTVLRDNAIPTIF

>PNJ51200.1 Pongo abelii

MPNFCAAPNCTRKSTQSDLAFFRFPRDPARCQKWVENCRRADLEDKTPDQLNKHYRLCAKHFETSMICRT

SPYRTVLRDNAIPTIF

>XP_020022725.1 Castor canadensis

MPNFCAAPNCTRKSTQSDLAFFRFPRDPARCQKWVENCRRADLEDKTPDQLNKHYRLCAKHFETSMICRT

SPYRTVLRDNAIPTIF

>XP_012600479.1 Microcebusmurinus

MPNFCAAPNCTRKSTQSDLAFFRFPRDPARCQKWVENCRRADLEDKTPDQLNKHYRLCAKHFETSMICRT

SPYRTVLRDNAIPTIF

>XP_020369167.1 Rhincodon typus

MPNFCAAPNCSRGSTNYPDLPFFRFPRDRERCQKWVENCRRADLENRSAEQLHKQYRLCARHFEQSLICT

NSPYRTVLKDNAVPTLF

>XP_010394069.2 Corvuscornixcornix

MTATDHCADYLILLLTCRCQRWVENCRRADLEDKTPDQLNKHYRLCAKHFETSMICRSSPYRTVLRDNAV

PTIF

>XP_020665363.1 Pogonavitticeps

MFPPPKHTFWGCLFLKLLTSGESVSCRCQRWVENCRRADLEDKTPDQLNKHYRLCAKHFETSMICRSSPY

RTVLRDNAVPTIF

>XP_020748437.1 Odocoileusvirginianustexanus

MPNFCAAPNCTRKSTQSDLAFFRFPRDPARCQKWVENCRRADLEDKTPDQLNKHYRLCAKHFETSMICRT

SPYRTVLRDNAIPTIF

>XP_020785855.1 Boleophthalmuspectinirostris

MTDCCAAANCGYQRGEGENTPLFSFPLDPERCKLWLTKCHREDLGSEAPDHLHKTYKLCAKHFEPAMISH

QDDSSTVLKEDAVPTIF

>XP_020845191.1 Phascolarctoscinereus

MPNFCAAPNCTRKSTQSDLAFFRFPRDPARCQKWVENCRRADLEDKTPDQLNKHYRLCAKHFETSMICKS

SPYRTVLRDNAVPTIF

>XP_005074025.1 Mesocricetusauratus

MPNFCAAPNCTRKSTQSDLAFFRFPRDPARCQKWVENCRRADLEDKTPDQLNKHYRLCAKHFETSMICRT

SPYRTVLRDNAIPTIF

>XP_021140792.1 Columba livia

MSFTGVAVQNQSKCAVAQKSIARRCQRWVENCRRADLEDKTPDQLNKHYRLCAKHFETSMICRSSPYRTV

LRDNAVPTIF

>XP_021239113.1 Numidameleagris

MPNFCAAPNCTRKSTQSDLAFFRFPRDPVRCQRWVENCRRADLEDKTPDQLNKHYRLCAKHFETSMICRS

SPYRTVLRDNAVPTIF

>XP_021512236.1 Merionesunguiculatus

MPNFCAAPNCTRKSTQSDLAFFRFPRDPARCQKWVENCRRADLEDKTPDQLNKHYRLCAKHFETSMICKT

SPYRTVLRDNAIPTIF

>XP_012325062.1 Aotusnancymaae

MPNFCAAPNCTRKSTQSDLAFFRFPRDPARCQKWVENCRRADLEDKTPDQLNKHYRLCAKHFETSMICRT

SPYRTVLRDNAIPTIF

>XP_021560352.1 Neomonachusschauinslandi

MPNFCAAPNCTRKSTQSDLAFFRFPRDPARCQKWVENCRRADLEDKTPDQLNKHYRLCAKHFETSMICRT

SPYRTVLRDNAIPTIF

>XP_542300.2 Canis lupus familiaris

MPNFCAAPNCTRKSTQSDLAFFRFPRDPARCQKWVENCRRADLEDKTPDQLNKHYRLCAKHFETSMICRT

SPYRTVLRDNAIPTIF

>XP_023095187.1 Feliscatus

MPNFCAAPNCTRKSTQSDLAFFRFPRDPARCQKWVENCRRADLEDKTPDQLNKHYRLCAKHFETSMICRT

SPYRTVLRDNAIPTIF

>XP_003780933.1 Otolemurgarnettii

MPNFCAAPNCTRKSTQSDLAFFRFPRDPARCQKWVENCRRADLEDKTPDQLNKHYRLCAKHFETSMICRT

SPYRTVLRDNAIPTIF

>XP_003420078.1 Loxodontaafricana

MPNFCAAPNCTRKSTQSDLAFFRFPRDPARCQKWVENCRRADLEDKTPDQLNKHYRLCAKHFETSMICRT

SPYRTVLRDNAIPTIF

>XP_004451033.1 Dasypusnovemcinctus

MPNFCAAPNCTRKSTQSDLAFFRFPRDPARCQKWVENCRRADLEDKTPDQLNKHYRLCAKHFETSMICRT

SPYRTVLRDNAIPTIF

>XP_003468597.1 Caviaporcellus

MPNFCAAPNCTRKSTQSDLAFFRFPRDPARCQKWVENCRRADLEDKTPDQLNKHYRLCAKHFETSMICRT

SPYRTVLRDNAIPTIF

>XP_023501453.1 Equuscaballus

MPNFCAAPNCTRKSTQSDLAFFRFPRDPARCQKWVENCRRADLEDKTPDQLNKHYRLCAKHFETSMICRT

SPYRTVLRDNAIPTIF

>XP_023799812.1 Cyanistescaeruleus

MESYQCQRWVENCRRADLEDKTPDQLNKHYRLCAKHFETSMICRSSPYRTVLRDNAVPTIF

>XP_011482119.1 Oryziaslatipes

MNECCAASNCDYQQRESGKDSQLFSFPLDPERCKQWTNNCQRTDLASQPPETLHKLYRLCSKHFESSMIS

QQDASTFVLKEDAVPTIF

>XP_005290594.1 Chrysemyspictabellii

MPNFCAAPNCTRKSTQSDLAFFRFPRDPVRCQRWVENCRRADLEDKTPDQLNKHYRLCAKHFETSMICRS

SPYRTVLRDNAVPTIF

>XP_024428450.1 Desmodusrotundus

MPNFCAAPNCTRKSTQSDLAFFRFPRDPARCQKWVENCRRADLEDKTPDQLNKHYRLCAKHFETSMICRT

SPYRTVLRDNAIPTIF

>XP_011717425.1 Macacanemestrina

MPNFCAAPNCTRKSTQSDLAFFRFPRDPARCQKWVENCRRADLEDKTPDQLNKHYRLCAKHFETSMICRT

SPYRTVLRDNAIPTIF

>XP_006926697.1 52 Pteropusalecto

MPNFCAAPNCTRKSTQSDLAFFRFPRDPARCQKWVENCRRADLEDKTPDQLNKHYRLCAKHFETSMICRT

SPYRTVLRDNAIPTIF

>XP_425679.4 Gallus gallus

MRQAPCGCLTAQARRPPALSLRREARRARQQPACAPLIWAPHGACTLRAAGLSALWEGGGPCSRRRVAVN

PPGRSQQCGAASPPYPSSLPPPRPPPPPPPPPPPHIPPLRPGGFKGAAPPIRGAMPNFCAAPNCTRKSTQ

SDLAFFRFPRDPVRCQRWVENCRRADLEDKTPDQLNKHYRLCAKHFETSMICRSSPYRTVLRDNAVPTIF

>XP_006022037.1 Alligator sinensis

MRLFPRPCDVSTEIKCQRWVENCRRADLEDKTPDQLNKHYRLCAKHFETSMICRSSPYRTVLRDNAVPTI

F

>XP_025045663.1 Pelodiscussinensis

MAILGQSIDPFSPVPFLLAVANTRCFSGNEQNRCQRWVENCRRADLEDKTPDQLNKHYRLCAKHFETSMI

CRSSPYRTVLRDNAVPTIF

>XP_006045692.1 Bubalusbubalis

MPNFCAAPNCTRKSTQSDLAFFRFPRDPARCQKWVENCRRADLEDKTPDQLNKHYRLCAKHFETSMICRT

SPYRTVLRDNAIPTIF

>XP_025214046.1 Theropithecus gelada

MPNFCAAPNCTRKSTQSDLAFFRFPRDPARCQKWVENCRRADLEDKTPDQLNKHYRLCAKHFETSMICRT

SPYRTVLRDNAIPTIF

>XP_025750257.1 Callorhinusursinus

MPNFCAAPNCTRKSTQSDLAFFRFPRDPARCQKWVENCRRADLEDKTPDQLNKHYRLCAKHFETSMICRT

SPYRTVLRDNAIPTIF

>XP_025866395.1 Vulpesvulpes

MAISATEAYVVATEASKERDAGDTPRKLSPDLQKPAHAKPTEGCAQAHCGSPRQAQPPPRHTRAGALLCA

CPPVEKSAERRGGGGRLREERAPPGARVHCGKEAAASRAGGRCTSGSCGGGGGKSSRPLTALWSRVPSPA

SDRGAPPAPRGFSGPPSSPPASARGGPQALRRGPVGARGPSEHVRQRGLGAAAPPETRGNWGGAQARCQK

WVENCRRADLEDKTPDQLNKHYRLCAKHFETSMICRTSPYRTVLRDNAIPTIF

>XP_025890320.1 Nothoproctaperdicaria

MPNFCAAPNCTRKSTQSDLAFFRFPRDPVRCQRWVENCRRADLEDKTPDQLNKHYRLCAKHFETSMICRS

SPYRTVLRDNAVPTIF

>XP_025930532.1 Apteryx rowi

MIVRRSWCSRQECFMELRKILAAVKAVHLDIETPVLPTLIHGCRCSPVSLDTALQNNLTSLASLGFGRCQ

RWVENCRRADLEDKTPDQLNKHYRLCAKHFETSMICRSSPYRTVLRDNAVPTIF

>XP_025953693.1 Dromaiusnovaehollandiae

MPNFCAAPNCTRKSTQSDLAFFRFPRDPVRCQRWVENCRRADLEDKTPDQLNKHYRLCAKHFETSMICRS

SPYRTVLRDNAVPTIF

>XP_026373914.1 Ursusarctoshorribilis

MPNFCAAPNCTRKSTQSDLAFFRFPRDPARCQKWVENCRRADLEDKTPDQLNKHYRLCAKHFETSMICRT

SPYRTVLRDNAIPTIF

>XP_026579269.1 Pseudonajatextilis

MPNFCAAPNCTRKSTQSDLAFFRFPRDPARCQRWVENCRRVDLEDKTPDQLNKHYRLCAEHFETSMICRS

SPYRTVLRDNAVPTIF

>XP_005357588.1 Microtus ochrogaster

MPNFCAAPNCTRKSTQSDLAFFRFPRDPARCQKWVENCRRADLEDKTPDQLNKHYRLCAKHFETSMICRT

SPYRTVLRDNAIPTIF

>XP_014122611.1 Zonotrichiaalbicollis

MPNFCAAPNCTRKSTQSDLAFFRFPRDPARCQRWVENCRRADLEDKTPDQLNKHYRLCAKHFETSMICRS

SPYRTVLRDNAVPTIF

>XP_026895655.1 Acinonyxjubatus

MPNFCAAPNCTRKSTQSDLAFFRFPRDPARCQKWVENCRRADLEDKTPDQLNKHYRLCAKHFETSMICRT

SPYRTVLRDNAIPTIF

>XP_026939239.1 Lagenorhynchusobliquidens

MPNFCAAPNCTRKSTQSDLAFFRFPRDPARCQKWVENCRRADLEDKTPDQLNKHYRLCAKHFETSMICRT

SPYRTVLRDNAIPTIF

>XP_027434490.1 Zalophuscalifornianus

MPNFCAAPNCTRKSTQSDLAFFRFPRDPARCQKWVENCRRADLEDKTPDQLNKHYRLCAKHFETSMICRT

SPYRTVLRDNAIPTIF

>XP_027555631.1 Neopelmachrysocephalum

MGMVLVPGMKSSGRAGNKNGAAPGRTGDGTGAHPGGGELRDSWEWEWCWSRGWRAPGQPGMVLVPGMESS

RRAGDRNAWFNWMLTVLAGAQAQGSTGYLFPNKQCQRWVENCRRADLEDKTPDQLNKHYRLCAKHFETSM

ICRSSPYRTVLRDNAVPTIF

>XP_027519431.1 Corapipoaltera

MPNFCAAPNCTRKSTQSDLAFFRFPRDPARCQRWVENCRRADLEDKTPDQLNKHYRLCAKHFETSMICRS

SPYRTVLRDNAVPTIF

>XP_027573644.1 Piprafilicauda

MESSGRGNGNGNGDGWRAPGMGTGTGMDGELRPWERERGWMESSGRGNGNGWRAPGMGTGTGMDGELRPL

ERERERGWMESSDRGNGNGNGDGWRAPAVGTGTGMDGELREWERERGWMESSGNGKGPGDGELREWEGSR

GWRAPAVGTGTGMNGELREWERERGWRAPAVGTGRVLGMESSENGNGDGELREWDWCWSWGWRAPGGLGT

GMLGLTGCSQFWKVHRHRAVPVTSFLTNSRCQRWVENCRRADLEDKTPDQLNKHYRLCAKHFETSMICRS

SPYRTVLRDNAVPTIF

>XP_027633677.1 Falco peregrinus

MPNFCAAPNCTRKSTQSDLAFFRSPRDPVRCQRWVENCRRADLEDKTPDQLNKHYRLCAKHFETSMICRS

SPYRTVLRDNAVPTIF

>XP_007061470.2 Cheloniamydas

MPNFCAAPNCTRKSTQSDLAFFRFPRDPVRCQRWVENCRRADLEDKTPDQLNKHYRLCAKHFETSMICRS

SPYRTVLRDNAVPTIF

>XP_027744120.1 Empidonaxtraillii

MESSGSGNGNGPGMESSGSGNGPGMESSGSGNGPGMESSGRGNGNGPGMESSGRGNGNGCQRWVENCRRA

DLEDKTPDQLNKHYRLCAKHFETSMICRSSPYRTVLRDNAVPTIF

>XP_027704153.1 Vombatusursinus

MPNFCAAPNCTRKSTQSDLAFFRFPRDPARCQKWVENCRRADLEDKTPDQLNKHYRLCAKHFETSMICKS

SPYRTVLRDNAVPTIF

>XP_027835356.1 Ovisaries

MPNFCAAPNCTRKSTQSDLAFFRFPRDPARCQKWVENCRRADLEDKTPDQLNKHYRLCAKHFETSMICRT

SPYRTVLRDNAIPTIF

>XP_027971829.1 Eumetopiasjubatus

MPNFCAAPNCTRKSTQSDLAFFRFPRDPARCQKWVENCRRADLEDKTPDQLNKHYRLCAKHFETSMICRT

SPYRTVLRDNAIPTIF

>XP_028017673.1 Balaenopteraacutorostratascammoni

MPNFCAAPNCTRKSTQSDLAFFRFPRDPARCQKWVENCRRADLEDKTPDQLNKHYRLCAKHFETSMICRT

SPYRTVLRDNAIPTIF

>XP_028371184.1 Phyllostomus discolor

MPNFCAAPNCTRKSTQSDLAFFRFPRDPARCQKWVENCRRADLEDKTPDQLNKHYRLCAKHFETSMICRT

SPYRTVLRDNAIPTIF

>XP_028626488.1 Grammomyssurdaster

MPNFCAAPNCTRKSTQSDLAFFRFPRDPARCQKWVENCRRADLEDKTPDQLNKHYRLCAKHFETSMICRT

SPYRTVLRDNAIPTIF

>XP_028581722.1 Podarcismuralis

MPNFCAAPNCTRKSTQSDLAFFRFPRDPARCQRWVENCRRADLEDKTPDQLNKHYRLCAKHFETSMICRS

SPYRTVLRDNAVPTIF

>XP_028655366.1 Erpetoichthyscalabaricus

MPNFCAAPNCTRKSTQSDLAFFRFPRDPIRRCKLWVENCRRADLEDKTSDQLNKHYRLCAKHFEPSMICK

SSPYRTVLKDNAVPTIF

>XP_028733319.1 Peromyscusleucopus

MPNFCAAPNCTRKSTQSDLAFFRFPRDPARCQKWVENCRRADLEDKTPDQLNKHYRLCAKHFETSMICRT

SPYRTVLRDNAIPTIF

>XP_028839571.1 Denticepsclupeoides

MPNFCIAPNCTRRSTQSDLAFFRFPRDTERCNIWVENCRRADLVDKTPDQLNKHYRLCANHFEPSMICKT

SPYRTVLRENAIPTIF

>XP_014970711.2 Macacamulatta

MPNFCAAPNCTRKSTQSDLAFFRFPRDPARCQKWVENCRRADLEDKTPDQLNKHYRLCAKHFETSMICRT

SPYRTVLRDNAIPTIF

>XP_028904329.1 Ornithorhynchusanatinus

MPNFCAAPNCTRKSTQSDLAFFRFPRDPARCQKWVENCRRADLEDKTPDQLNKHYRLCAKHFEASMICRS

SPYRTILRDNAVPTIF

>XP_029064338.1 Monodon monoceros

MSVCHCLLYHTSGPDCEAGWNCRPSFGILPAEYASLGTEREIPRGPAPTFHSYDPLRQNTENKGRGQSQQ

AKVWTVKKKLGKEKKKKKEKSSTSKYQKPTARSSSKLWRTARKMAQESSKECQKWVENCRRADLEDKTPD

QLNKHYRLCAKHFETSMICRTSPYRTVLRDNAIPTIF

>XP_021022504.1 Mus caroli

MPNFCAAPNCTRKSTQSDLAFFRFPRDPARCQKWVENCRRADLEDKTPDQLNKHYRLCAKHFETSMICRT

SPYRTVLRDNAIPTIF

>XP_021049125.1 Mus pahari

MPNFCAAPNCTRKSTQSDLAFFRFPRDPARCQKWVENCRRADLEDKTPDQLNKHYRLCAKHFETSMICRT

SPYRTVLRDNAIPTIF

>XP_008842450.1 Nannospalaxgalili

MPNFCAAPNCTRKSTQSDLAFFRFPRDPARCQKWVENCRRADLEDKTPDQLNKHYRLCAKHFETSMICRT

SPYRTVLRDNAIPTIF

>XP_029458017.1 Rhinatremabivittatum

MLHSRATGARPPKPSSCHLGKPACGRRAGHHGKEEALLGPSHSENWRSGSGRSGTARGGFGAESSRPLRA

TVALRPQHPVSPPTHTLSLTRERGREIIPFKSTGNPPPPILNRKIKNKHYVFIIFLIVKKKQKTKTHTNQ

NNGYRRTTMPNFCAAPNCTRKSTQSDLAFFRFPRDPARCKKWVENCRRADLEDKTADQLNKHYRLCAQHF

ETSMICRSSPYRTVLLENAIPTIF

>XP_029898439.1 Aquila chrysaetoschrysaetos

MPNFCAAPNCTRKSTQSDLAFFRFPRDPVRCQRWVENCRRADLEDKTPDQLNKHYRLCAKHFETSMICRS

SPYRTVLRDNAVPTIF

>XP_030055195.1 Microcaecilia unicolor

MPNYCAALNCSRKSTHSDIGFFRFPRDPARCKQWVENCQRVDLEDKTADHLNKCYRLCARHFDSSMVRRK

SPYRTRLQDNAVPTIF

>XP_018779501.2 Serinuscanaria

APNCTRKSTQSDLAFFRFPRDPARCQRWVENCRRADLEDKTPDQLNKHYRLCAKHFETSMICRSSPYRTV

LRDNAVPTIF

>XP_030187680.1 Lynx canadensis

MPNFCAAPNCTRKSTQSDLAFFRFPRDPARCQKWVENCRRADLEDKTPDQLNKHYRLCAKHFETSMICRT

SPYRTVLRDNAIPTIF

>XP_008493412.2 Calypte anna

MPNFCAAPNCTRKSTQSDLAFFRFPRDPVRCQRWVENCRRADLEDKTPDQLNKHYRLCAKHFETSMICRS

SPYRTVLRDNAVPTIF

>XP_030413541.1 Gopherusevgoodei

MPNFCAAPNCTRKSTQSDLAFFRFPRDPVRCQRWVENCRRADLEDKTPDQLNKHYRLCAKHFETSMICRS

SPYRTVLRDNAVPTIF

>XP_022449566.1 Delphinapterus leucas

MPNFCAAPNCTRKSTQSDLAFFRFPRDPARCQKWVENCRRADLEDKTPDQLNKHYRLCAKHFETSMICRT

SPYRTVLRDNAIPTIF

>XP_030648517.1 Chanoschanos

MPNFCAAPNCTRKSTQSDLAFFRFPRDAQRCRIWVENCRRADLEAKTADQLNKHYRLCAKHFEPHMVCKT

SPFRTVLRDNAIPTIF

>XP_030685502.1 Nomascusleucogenys

MPNFCAAPNCTRKSTQSDLAFFRFPRDPARCQKWVENCRRADLEDKTPDQLNKHYRLCAKHFETSMICRT

SPYRTVLRDNAIPTIF

>XP_030691894.1 Globicephalamelas

MPNFCAAPNCTRKSTQSDLAFFRFPRDPARCQKWVENCRRADLEDKTPDQLNKHYRLCAKHFETSMICRT

SPYRTVLRDNAIPTIF

>XP_010365084.2 Rhinopithecusroxellana

MPNFCAAPNCTRKSTQSDLAFFRFPRDPARCQKWVENCRRADLEDKTPDQLNKHYRLCAKHFETSMICRT

SPYRTVLRDNAIPTIF

>XP_030813770.1 Camarhynchusparvulus

MRCVRGRWDTPATGRNCHRIAGMVAKRHDRQQRCQRWVENCRRADLEDKTPDQLNKHYRLCAKHFETSMI

CRSSPYRTVLRDNAVPTIF

>XP_018892915.1 Gorilla gorillagorilla

MPNFCAAPNCTRKSTQSDLAFFRFPRDPARCQKWVENCRRADLEDKTPDQLNKHYRLCAKHFETSMICRT

SPYRTVLRDNAIPTIF

>XP_031316247.1 Camelusdromedarius

MPNFCAAPNCTRKSTQSDLAFFRFPRDPARCQKWVENCRRADLEDKTPDQLNKHYRLCAKHFETSMICRT

SPYRTVLRDNAIPTIF

>XP_021388106.1 Lonchurastriatadomestica

MPNFCAAPNCTRKSTQSDLAFFRFPRDPARCQRWVENCRRADLEDKTPDQLNKHYRLCAKHFETSMICRS

SPYRTVLRDNAVPTIF

>XP_031456953.1 Phasianuscolchicus

MPNFCAAPNCTRKSTQSDLAFFRFPRDPVRCQRWVENCRRADLEDKTPDQLNKHYRLCAKHFETSMICRS

SPYRTVLRDNAVPTIF

>XP_012686697.2 Clupeaharengus

MPNFCVAPNCTKKSTQTDLAFFRFPRDVKRCHEWVENCRRADLLAKTPDQLNKHYRLCAAHFDPTMICKT

GPFRTVLKDNAVPTIF

>XP_003910488.2 Papioanubis

MPNFCAAPNCTRKSTQSDLAFFRFPRDPARCQKWVENCRRADLEDKTPDQLNKHYRLCAKHFETSMICRT

SPYRTVLRDNAIPTIF

>XP_006203380.1 Vicugnapacos

MPNFCAAPNCTRKSTQSDLAFFRFPRDPARCQKWVENCRRADLEDKTPDQLNKHYRLCAKHFETSMICRT

SPYRTVLRDNAIPTIF

>NP_989053.1 Xenopustropicalis

MPNFCAAPNCTRKSTQSDLAFFRFPRDPDRRQRWVENCRRADLEDKTPDQLNKHYRLCAQHFEDSMICRS

SPYRTVLRENAIPTIF

>XP_023065133.1 Piliocolobustephrosceles

MPNFCAAPNCTRKSTQSDLAFFRFPRDPARCQKWVENCRRADLEDKTPDQLNKHYRLCAKHFETSMICRT

SPYRTVLRDNAIPTIF

>XP_031817214.1 Sarcophilus harrisii

MDRSRSQSPSGFLKNKPTQGSLALLYSQRPFTWPARGSGEAPDECGGFSRPPTPDETYAVCRRERAAILG

GKVHSALSRCQKWVENCRRADLEDKTPDQLNKHYRLCAKHFETSMICKSSPYRTVLRDNAVPTIF

>XP_032063475.1 Aythyafuligula

MPNFCAAPNCTRKSTQSDLAFFRFPRDPVRCQRWVENCRRADLEDKTPDQLNKHYRLCAKHFETSMICRS

SPYRTVLRDNAVPTIF

>XP_032076156.1 Thamnophis elegans

MPNFCAAPNCTRKSTQSDLAFFRFPRDPARCQRWVENCRRVDLEDKTPDQLNKHYRLCAEHFETSMICRS

SPYRTVLRDNAVPTIF

>XP_032216423.1 Mustelaerminea

MPNFCAAPNCTRKSTQSDLAFFRFPRDPARCQKWVENCRRADLEDKTPDQLNKHYRLCAKHFETSMICRT

SPYRTVLRDNAIPTIF

>XP_032283237.1 Phocavitulina

MPNFCAAPNCTRKSTQSDLAFFRFPRDPARCQKWVENCRRADLEDKTPDQLNKHYRLCAKHFETSMICRT

SPYRTVLRDNAIPTIF

>XP_015707969.1 Coturnix japonica

MPNFCAAPNCTRKSTQSDLAFFRFPRDPVRCQRWVENCRRADLEDKTPDQLNKHYRLCAKHFETSMICRS

SPYRTVLRDNAVPTIF

>XP_032345329.1 Camelusferus

MPNFCAAPNCTRKSTQSDLAFFRFPRDPARCQKWVENCRRADLEDKTPDQLNKHYRLCAKHFETSMICRT

SPYRTVLRDNAIPTIF

>XP_032497564.1 Phocoena sinus

MPNFCAAPNCTRKSTQSDLAFFRFPRDPARCQKWVENCRRADLEDKTPDQLNKHYRLCAKHFETSMICRT

SPYRTVLRDNAIPTIF

>XP_032534915.1 Chiroxiphialanceolata

MPNFCAAPNCTRKSTQSDLAFFRFPRDPARCQRWVENCRRADLEDKTPDQLNKHYRLCAKHFETSMICRS

SPYRTVLRDNAVPTIF

>XP_032022852.1 Hylobatesmoloch

MPNFCAAPNCTRKSTQSDLAFFRFPRDPARCQKWVENCRRADLEDKTPDQLNKHYRLCAKHFETSMICRT

SPYRTVLRDNAIPTIF

>XP_030114918.2 Taeniopygiaguttata

MPNFCAAPNCTRKSTQSDLAFFRFPRDPARCQRWVENCRRADLEDKTPDQLNKHYRLCAKHFETSMICRS

SPYRTVLRDNAVPTIF

>XP_032708915.1 Lontracanadensis

MPNFCAAPNCTRKSTQSDLAFFRFPRDPARCQKWVENCRRADLEDKTPDQLNKHYRLCAKHFETSMICRT

SPYRTVLRDNAIPTIF

>XP_032748896.1 Rattusrattus

MPNFCAAPNCTRKSTQSDLAFFRFPRDPARCQKWVENCRRADLEDKTPDQLNKHYRLCAKHFETSMICRT

SPYRTVLRDNAIPTIF

>XP_032878035.1 Amblyrajaradiata

MPNFCAAPNCSRKSTNCPEIPFFRFPKDPTRCQKWVENCRRADLENRSAEQLHKQYRLCARHFEQSLICT

NSPYRTVLKDNAVPTLF

>XP_032846687.1 Tytoalbaalba

LASHNFGRCIGARAPIPIISPNSRCQRWVENCRRADLEDKTPDQLNKHYRLCAKHFETSMICRSSPYRTV

LRDNAVPTIF

>XP_032908331.1 Catharusustulatus

MPNFCAAPNCTRKSTQSDLAFFRFPRDPARCQRWVENCRRADLEDKTPDQLNKHYRLCAKHFETSMICRS

SPYRTVLRDNAVPTIF

>XP_032977322.1 Rhinolophusferrumequinum

MPNFCAAPNCTRKSTQSDLAFFRFPRDPARCQKWVENCRRADLEDKTPDQLNKHYRLCAKHFETSMICRT

SPYRTVLRDNAIPTIF

>XP_030332808.1 Strigopshabroptila

MRAAHPGAAAAARRMGTAPAGGRSRCTVGRRRPLSRRTVAVNPPGRSQQCRAASPPPPRPPPSFLPPPSS

SSSPHTPPPFPSPPLPSAFKGAAAPTRGTMPNFCAAPNCTRKSTQSDLAFFRFPRDPVRCQRWVENCRRA

DLEDKTPDQLNKHYRLCAKHFETSMICRSSPYRTVLRDNAVPTIF

>XP_033001876.1 Lacerta agilis

MPNFCAAPNCTRKSTQSDLAFFRFPRDPARCQRWVENCRRADLEDKTPDQLNKHYRLCAKHFETSMICRS

SPYRTVLRDNAVPTIF

>XP_033063924.1 Trachypithecusfrancoisi

MPNFCAAPNCTRKSTQSDLAFFRFPRDPARCQKWVENCRRADLEDKTPDQLNKHYRLCAKHFETSMICRT

SPYRTVLRDNAIPTIF

>XP_004279862.1 Orcinus orca

MPNFCAAPNCTRKSTQSDLAFFRFPRDPARCQKWVENCRRADLEDKTPDQLNKHYRLCAKHFETSMICRT

SPYRTVLRDNAIPTIF

>XP_015497969.2 Parus major

MPNFCAAPNCTRKSTQSDLAFFRFPRDPARCQRWVENCRRADLEDKTPDQLNKHYRLCAKHFETSMICRS

SPYRTVLRDNAVPTIF

>XP_019804359.2 Tursiopstruncatus

MPNFCAAPNCTRKSTQSDLAFFRFPRDPARCQKWVENCRRADLEDKTPDQLNKHYRLCAKHFETSMICRT

SPYRTVLRDNAIPTIF

>XP_033804978.1 Geotrypetesseraphini

MPTFGNTDGPMEHSILWDRAQCSGLSAPICWLMGISHKFLDSSAATKGMKNIKCKKWVENCRRADLEDKT

ADQLNKHYRLCAQHFETSMICRSSPYRTVLRENAVPTIF

>XP_034362635.1 Arvicanthisniloticus

MPNFCAAPNCTRKSTQSDLAFFRFPRDPARCQKWVENCRRADLEDKTPDQLNKHYRLCAKHFETSMICRT

SPYRTVLRDNAIPTIF

>XP_034282979.1 Pantherophisguttatus

MPNFCAAPNCTRKSTQSDLAFFRFPRDPARCQRWVENCRRVDLEDKTPDQLNKHYRLCAEHFETSMICRS

SPYRTVLRDNAVPTIF

>XP_027798507.1 Marmota flaviventris

MPNFCAAPNCTRKSTQSDLAFFRFPRDPARCQKWVENCRRADLEDKTPDQLNKHYRLCAKHFETSMICRT

SPYRTVLRDNAIPTIF

>XP_034522196.1 Ailuropodamelanoleuca

MPNFCAAPNCTRKSTQSDLAFFRFPRDPARCQKWVENCRRADLEDKTPDQLNKHYRLCAKHFETSMICRT

SPYRTVLRDNAIPTIF

>KAF4789057.1 Turdusrufiventris

MPNFCAAPNCTRKSTQSDLAFFRFPRDPARCQRWVENCRRADLEDKTPDQLNKHYRLCAKHFETSMICRS

SPYRTVLRDNAVPTIF

>XP_034615036.1 Trachemysscriptaelegans

MPNFCAAPNCTRKSTQSDLAFFRFPRDPVRCQRWVENCRRADLEDKTPDQLNKHYRLCAKHFETSMICRS

SPYRTVLRDNAVPTIF

>XP_034789288.1 Pan paniscusMPNFCAAPNCTRKSTQSDLAFFRFPRDPARCQKWVENCRRADLEDKTPDQLNKHYRLCAKHFETSMICRT

SPYRTVLRDNAIPTIF

>XP_034851518.1 Miroungaleonina

MPNFCAAPNCTRKSTQSDLAFFRFPRDPARCQKWVENCRRADLEDKTPDQLNKHYRLCAKHFETSMICRT

SPYRTVLRDNAIPTIF

>XP_011240217.1 Mus musculus

MPNFCAAPNCTRKSTQSDLAFFRFPRDPARCQKWVENCRRADLEDKTPDQLNKHYRLCAKHFETSMICRT

SPYRTVLRDNAIPTIF

>XP_034972129.1 Zootoca vivipara

MPNFCAAPNCTRKSTQSDLAFFRFPRDPARCQRWVENCRRADLEDKTPDQLNKHYRLCAKHFETSMICRS

SPYRTVLRDNAVPTIF

>NP_001179132.2 Bostaurus

MPNFCAAPNCTRKSTQSDLAFFRFPRDPTRCQKWVENCRRADLEDKTPDQLNKHYRLCAKHFETSMICRT

SPYRTVLRDNAIPTIF

>XP_035121271.1 Callithrix jacchus

MPNFCAAPNCTRKSTQSDLAFFRFPRDPARCQKWVENCRRADLEDKTPDQLNKHYRLCAKHFETSMICRT

SPYRTVLRDNAIPTIF

>XP_007645845.1 Cricetulusgriseus

MLNFCAAPNCTRKSTQSDLAFFRFPRDPARCQKWVENCRRADLEDKTPDQLNKHYRLCAKHFETSMICRT

SPYRTVLRDNAIPTIF

>NP_004696.2 Homo sapiens MPNFCAAPNCTRKSTQSDLAFFRFPRDPARCQKWVENCRRADLEDKTPDQLNKHYRLCAKHFETSMICRT

SPYRTVLRDNAIPTIF
