## Additional file for "Synapomorphic variations in the THAP domains of the human THAP protein family and its homologs": THAP1.docx

>PFX25192.1|Stylophora pistillata (Cnidaria)

MPHFCCAGECKNSSDKRPDISFHGLPLDNKALLKTWIAKMRRNPNYFNVNKHVKICSKHFSPEDFINPDA

KKRRLKRNAVPSIF

>KXJ27574.1|Exaiptasia pallid (Cnidaria)

MKKHATENLIWDEGQLGAIERVLGTVDQLIIDRCIMGEVKQYQRNLAVAFYDYKKAYDKCSQKGYKRNGN

PVSFFNFPKDQEMCKKWIIAIRRDVGKDFKLNERTKICSLHFNSLDLHKSLNGRIYPVKGAVPSVF

>RXG53912.1|Armadillidium vulgare (arthropoda)

MLLITDDDLKWVESFFPLTNKKLTKIWGMKMKRGKFYPSKFSKICSDHFTPDCINPGVGFGKPTLKQNAI

PTIF

>JAG83381.1|Fopius arisanus (arthropoda)

MVRQCCIKDCKSEWWPGTELSFHSFPKREESRKKWLEAIPPGKLRSTNLNYSSVCSKHFNESCFSDCINT

KTLRNSLKKDAVPTLF

>PRD35590.1|Trichonephila clavipes (arthropoda)

MVVSCCALGCKERFTKGGAYTFHCFPKDEARRKLWEKQLRRGNFTSTPYSRICSKHFEEKYFQVEKFGGK

WLKSDAVPTIF

>032877350.1|Amblyraja radiate (‎Chordata)

MVLSCSAYGCKNRYDKDRGTSFHRFPLKRPELCKKWLAAVRRRNFIPTKNSNLCSEHFTLDCFRKNCNNK

ILEENAVPSIF

>006036514.2|Alligator sinensis (Chordata)

MHAYMHVCVHVCMFPLTRPDLCKKWEAAVKRKNFKPTKYSSICSEHFTPDCFKRECNNKLLKENAVPTIF

>020368158.1|Rhincodon typus (‎Chordata)

MVLSCSAYGCKNRYDKDRGVSFHRFPLKRPELCKKWLAAVRRRNFTPTKNSNLCSEHFTLDCFRKNCNNK

ILEENAVPSIF

>005278942.1|Chrysemys picta bellii (Chordata)

MVQSCSAYRCRNRYDKEKPISFHKFPLTRPDLCKKWEAAVKRKNFKPTKYSSICSEHFTPDCFKRECNNK

LLKENAVPTIF

>006111180.1|Pelodiscus sinensis (Chordata)

MPTNPECCTRAFCMPTVLLLSVALIVHFLCRFPLTRPDLCKKWEAAVKRKNFKPTKYSSICSEHFTPDCF

KRECNNKLLKENAVPTIF

>007063130.1|Chelonia mydas (Chordata)

MVQSCSAYRCRNRYDKEKPTSFHKFPLTRPDLCKKWEAAVKRKNFKPTKYSSICSEHFTPDCFKRECNNK

LLKENAVPTIF

>032636188.1|Chelonoidis abingdonii (Chordata)

MVQSCSAYRCRNRYDKEKPISFHKFPLTRPDLCKKWEAAVKRKNFKPTKYSSICSEHFTPDCFKRECNNK

LLKENAVPTIF

>020657039.1|Pogona vitticep (Chordata)

MAVPEEQVTEGNEKISLKRSEEGLEKKRFPLTRPDLCKKWEAAVRRKNFKPTKYSSICSEHFTPDCFKRE

CNNKLLKDNAVPTIF

>007425019.1|Python bivittatus (Chordata)

MVQSCSAYRCRNRYDKEKPISFHKFPLTRPDLCKKWEAAVRRKNFKPTKYSSICSEHFTPDCFKRECNNK

LLKDNAVPTIF

>026528788.1|Notechis scutatus (Chordata)

MVQSCSAYRCRNRYDKEKPISFHKFPLTRPDLCKKWEAAVRRKNFKPTKYSSICSEHFTPDCFKRECNNK

LLKDNAVPTIF

>026568547.1|Pseudonaja textilis (Chordata)

MVQSCSAYRCRNRYDKEKPISFHKFPLTRPDLCKKWEAAVRRKNFKPTKYSSICSEHFTPDCFKRECNNK

LLKDNAVPTIF

>028567406.1|Podarcis muralis

MVQSCSAYRCRNRYDKEKPISFHKFPLTRPDLCKKWEAAVRRKNFKPTKYSSICSEHFTPDCFKRECNNK

LLKDNAVPTIF

>015679598.1|Protobothrops mucrosquamatus

MVQSCSAYRCRNRYDKEKPISFHKFPLTRPDLCKKWEAAVRRKNFKPTKYSSICSEHFTPDCFKRECNNK

LLKDNAVPTIF

>032070603.1|Thamnophis elegans

MVQSCSAYRCRNRYDKEKPISFHKFPLTRPDLCKKWEAAVRRKNFKPTKYSSICSEHFTPDCFKRECNNK

LLKDNAVPTIF

>033029477.1|Lacerta agilis

MVQSCSAYRCRNRYDKEKPISFHKFPLTRPDLCKKWEAAVRRKNFKPTKYSSICSEHFTPDCFKRECNNK

LLKDNAVPTIF

>AAH75499.1|Xenopus tropicalis

MVQSCSAYGCKNRYDKDKPISFHKFPLKRPLLCRKWEAAVRRADFKPTKYSSICSDHFTADCFKRECNNK

LLKDNAVPTIF

>001088492.1|Xenopus laevis

MVQSCSAYGCKNRYDKDKPISFHKFPLKRPLLCKKWEAAVRRAEFKPTKYSSICSDHFSADCFKRECNNK

LLKDNAVPTIF

>029458296.1|Rhinatrema bivittatum

MVVKTAMIRINQSPSTTTSVCWSRRESRTSMWWPSEAISSRISWRFPLTRPGLCEKWEAAVRRKNFKPSK

YSSICSEHFTPDCFKRECNNKLLKENAVPTIF

>030050327.1|Microcaecilia unicolor

MVQSCSVYGCKNRYDKDKPVSFHKFPLTRPGLCEKWEAAVRRKNFKPSKYSSICSEHFTPDCFKRACNNK

LLKENAVPTIF

>OPJ61978.1|Patagioenas fasciata monilis

MVQSCSAYRCRNRYDKEKPISFHKFPLTRPDLCKKWEAAVKRKNFKPTKYSSICSEHFTPDCFKRECNNK

LLKENAVPTIF

>010398445.1|Corvus cornix cornix

MVRESLDIMLHLLTMSPDPSEPPQLFLITHLKRFPEKLKVPLTCRFPLTRPDLCKKWEAAVKRKNFKPTK

YSSICSEHFTPDCFKRECNNKLLKENAVPTIF

>005514221.1|Columba livia

MVQSCSAYRCRNRYDKEKPISFHKFPLTRPDLCKKWEAAVKRKNFKPTKYSSICSEHFTPDCFKRECNNK

LLKENAVPTIF

>021236325.1|Numida meleagris

MVQSCSAYRCRNRYDKEKPISFHKFPLTRPDLCKKWEAAVKRKNFKPTKYSSICSEHFTPDCFKRECNNK

LLKENAVPTIF

>023799542.1|Cyanistes caeruleus

MARPEQPLTGSPGCRNSRSAAPGAGAVPGGAGGAAQLWVRGVRAALPSCGFPENLKVPLACRFPLTRPDL

CKKWEAAVKRKNFKPTKYSSICSEHFTPDCFKRECNNKLLKENAVPTIF

>004949398.1|Gallus gallus

MVQSCSAYRCRNRYDKEKPISFHKFPLTRPDLCKKWEAAVKRKNFKPTKYSSICSEHFTPDCFKRECNNK

LLKENAVPTIF

>025905758.1|Nothoprocta perdicaria

MVQSCSAYRCRNRYDKEKPISFHKFPLTRPDLCKKWEAAVKRKNFKPTKYSSICSEHFTPDCFKRECNNK

LLKENAVPTIF

>025934482.1|Apteryx rowi

MVQSCSAYRCRNRYDKEKPISFHKFPLTRPDLCKKWEAAVKRKNFKPTKYSSICSEHFTPDCFKRECNNK

LLKENAVPTIF

>014121990.1|Zonotrichia albicollis

MVQSCSAYRCRNRYDKEKPISFHKFPLTRPDLCKKWEAAVKRKNFKPTKYSSICSEHFTPDCFKRECNNK

LLKENAVPTIF

>027564280.1|Neopelma chrysocephalum

MVQSCSAYRCRNRYDKEKPISFHKFPLTRPDLCKKWEAAVKRKNFKPTKYSSICSEHFTPDCFKRECNNK

LLKENAVPTIF

>027508790.1|Corapipo altera

MHTDIHTYIHTGGSVRPRPLSFPERLKVPLTCRFPLTRPDLCKKWEAAVKRKNFKPTKYSSICSEHFTPD

CFKRECNNKLLKENAVPTIF

>027600947.1|Pipra filicauda

MVQSCSAYRCRNRYDKEKPISFHKFPLTRPDLCKKWEAAVKRKNFKPTKYSSICSEHFTPDCFKRECNNK

LLKENAVPTIF

>027756156.1|Empidonax traillii

MVQSCSAYRCRNRYDKEKPISFHKFPLTRPDLCKKWEAAVKRKNFKPTKYSSICSEHFTPDCFKRECNNK

LLKENAVPTIF

>029861377.1|Aquila chrysaetos

MVQSCSAYRCRNRYDKEKPISFHKFPLTRPDLCKKWEAAVKRKNFKPTKYSSICSEHFTPDCFKRECNNK

LLKENAVPTIF

>030093188.1|Serinus canaria

MVQSCSAYRCRNRYDKEKPISFHKFPLTRPDLCKKWEAAVKRKNFKPTKYSSICSEHFTPDCFKRECNNK

LLKENAVPTIF

>030323812.1|Calypte anna

MVQSCSAYRCRNRYDKEKPISFHKFPLTRPDLCKKWEAAVKRKNFKPTKYSSICSEHFTPDCFKRECNNK

LLKENAVPTIF

>030824511.1|Camarhynchus parvulus

MVQSCSAYRCRNRYDKEKPISFHKFPLTRPDLCKKWEAAVKRKNFKPTKYSSICSEHFTPDCFKRECNNK

LLKENAVPTIF

>021403661.1|Lonchura striata domestica

MQSSKVSPQNGNMLKLLKINLFVTTNGFVETSIYCLLMHLHTTFSRRDGTLGAACCRATASNETPFPPHP

DEGQEDFWLRLSSFPEKLKVPLACRFPLTRPDLCKKWEAAVKRKNFKPTKYSSICSEHFTPDCFKRECNN

KLLKENAVPTIF

>031465242.1|Phasianus colchicus

MVQSCSAYRCRNRYDKEKPISFHKFPLTRPDLCKKWEAAVKRKNFKPTKYSSICSEHFTPDCFKRECNNK

LLKENAVPTIF

>031952746.1|Corvus moneduloides

MVQSCSAYRCRNRYDKEKPISFHKFPLTRPDLCKKWEAAVKRKNFKPTKYSSICSEHFTPDCFKRECNNK

LLKENAVPTIF

>032062388.1|Aythya fuligula

MVQSCSAYRCRNRYDKEKPISFHKFPLTRPDLCKKWEAAVKRKNFKPTKYSSICSEHFTPDCFKRECNNK

LLKENAVPTIF

>015705063.1|Coturnix japonica

MVQSCSAYRCRNRYDKEKPISFHKFPLTRPDLCKKWEAAVKRKNFKPTKYSSICSEHFTPDCFKRECNNK

LLKENAVPTIF

>032532424.1|Chiroxiphia lanceolata

MVQSCSAYRCRNRYDKEKPISFHKFPLTRPDLCKKWEAAVKRKNFKPTKYSSICSEHFTPDCFKRECNNK

LLKENAVPTIF

>030113921.2|Taeniopygia guttata

MVQSCSAYRCRNRYDKEKPISFHKFPLTRPDLCKKWEAAVKRKNFKPTKYSSICSEHFTPDCFKRECNNK

LLKENAVPTIF

>032940902.1|Catharus ustulatus

MVQSCSAYRCRNRYDKEKPISFHKFPLTRPDLCKKWEAAVKRKNFKPTKYSSICSEHFTPDCFKRECNNK

LLKENAVPTIF

>030326314.1|Strigops habroptila

MVQSCSAYRCRNRYDKEKPISFHKFPLTRPDLCKKWEAAVKRKNFKPTKYSSICSEHFTPDCFKRECNNK

LLKENAVPTIF

>ACI68488.1|Salmo salar

MVQSCSAYGCKNRYHKDKNISFHKFPLARPDVCGKWVAAMRRNNFKPTRYSNICSQHFTKDCFKPECNNR

VLKENAVPSLF

>020456631.1|Monopterus albus

MVQTCSAYGCKNRYHKDKDISFHKFPLARPDVCGKWVAAMRRNNFKPTKYSNICSQHFTKDCFKRECNNR

VLKENAVPSLF

>020780191.1|Boleophthalmus pectinirostris

MVQTCSAYGCKNRYHKDKEISFHKFPLARPDVCDKWVSAMRRDNFRPTKYSSLCSQHFTQDSFKRECNNR

VLKENAVPSVF

>021457200.1|Oncorhynchus mykiss

MVQSCSAYGCKNRYHKDRNISFHKFPLARPDVCGKWVAAMRRNNFKPTRYSNICSQHFTKDCFKPECNNR

VLKENAVPSLF

>022063420.1|Acanthochromis polyacanthus

MVQTCSAYGCKNRYHKDKDISFHKFPLARPDVCGKWVSAMRRNNFKPTKYSNICSQHFTKDCFKRECNNR

VLKENAVPSLF

>007229747.2|Astyanax mexicanus

MVQSCSAYGCKNRYHKDKNVSFHKFPLARPDICGKWVAAMRRTNFKPTKYSNICSQHFTKDCFKRECNNR

VLKENAVPSLF

>SBR96341.1|Nothobranchius pienaari

MVQTCSAYGCKNRYHKDKDVSFHKFPLARPDVCGKWVAAMRRNNFKPTKYSNICSQHFTKDCFKRECNNR

VLKENAVPSLF

>SBS46605.1|Nothobranchius furzeri

MVQTCSAYGCKNRYHKDKDVSFHKFPLARPDVCGKWVAAMRRNNFKPTKYSNICSQHFTKDCFKRECNNR

VLKENAVPSLF

>SBP76845.1|Nothobranchius kadleci

MVQTCSAYGCKNRYHKDKDVSFHKFPLARPDVCGKWVAAMRRNNFKPTKYSNICSQHFTKDCFKRECNNR

VLKENAVPSLF

>SBP05728.1|Iconisemion striatum

MVQTCSAYGCKNRYHKDKDVSFHKFPLARPDVCGKWVAAMRRNNFKPTKYSNICSQHFTQDCFKRECNNR

VLKENAVPSLF

>SBQ63219.1|Nothobranchius korthausae

MVQTCSAYGCKNRYHKDKDVSFHKFPLARPDVCGKWVAAMRRNNFKPTKYSNICSQHFTKDCFKRECNNR

VLKENAVPSLF

>SBS08066.1|Nothobranchius rachovii

MVQTCSAYGCKNRYHKDKDVSFHKFPLARPDVCGKWVAAMRRNNFKPTKYSNICSQHFTKDCFKRECNNR

VLKENAVPSLF

>022600186.1|Seriola dumerili

MVQTCSAYGCKNRYHKDKDISFHKFPLARPDVCGKWVAAMRRNNFKPTKYSNICSQHFTKDCFKRECNNR

VLKENAVPSLF

>023119765.1|Amphiprion ocellaris

MVQTCSAYGCKNRYHKDKDISFHKFPLARPDVCGKWVSAMRRNNFKPTKYSNICSQHFTKDCFKRECNNR

VLKENAVPSLF

>023193567.1|Xiphophorus maculatus

MVQTCSAYGCKNRYHKDKDISFHKFPLARPDVCRKWVAAMRRSNFKPTKYSNICSQHFSQDCFKRECNNR

VLKDNAVPSLF

>023253598.1|Seriola lalandi dorsalis

MVQTCSAYGCKNRYHKDKDISFHKFPLARPDVCGKWVAAMRRNNFKPTKYSNICSQHFTKDCFKRECNNR

VLKENAVPSLF

>023664031.1|Paramormyrops kingsleyae

MVQSCSAYGCKNRYHKDKNISFHKFPLARPDICGKWVAAMRRNNFKPTKYSNICSQHFTKDCFKQECNNR

VLKENAVPSLF

>004084658.1|Oryzias latipes

MVQTCSAYGCKNRYHKDKDISFHKFPLARPDVCGKWVAAMRRNNFKPTKYSNICSQHFTSDCFKRECNNR

VLKENAVPSLF

>023996389.1|Salvelinus alpinus

MVQSCSAYGCKNRYHKDKNISFHKFPLARPDVCVKWVAAMRRNNFKPTRYSNICSQHFTKDCFKPECNNR

VLKENAVPSLF

>024124236.1|Oryzias melastigma

MVQTCSAYGCKNRYHKDKDVSFHKFPLARPDVCGKWVAAMRRNNFKPTKYSNICSQHFTKDCFKRECNNR

VLKENAVPSLF

>024237220.1|Oncorhynchus tshawytscha

MVQSCSAYGCKNRYHKDRNISFHKFPLARPDVCGKWVTAMRRNNFKPTRYSNICSQHFTKDCFKPECNNR

VLKENAVPSLF

>012777525.1|Maylandia zebra

MVQTCSAYGCKNRYHKDKEISFHKFPLARPDICCKWVAAMRRNNFKPTKYSNICSQHFTEDSFKRECNNR

VLKESAVPSLF

>008322677.1|Cynoglossus semilaevis

MVQTCSAYGCKNRYSKEKDISFHKFPLTRPDVCEKWVMAMGRDNFKPTKYSSICSQHFTKICFKDCNNRV

LKDHAVPSIF

>005470585.1|Oreochromis niloticus

MVQTCSAYGCKNRYHKDKDISFHKFPLARPDICCKWVAAMRRNNFKPTKYSNICSQHFTEDSFKRECNNR

VLKESAVPSLF

>026030340.1|Astatotilapia calliptera

MVQTCSAYGCKNRYHKDKEISFHKFPLARPDICCKWVAAMRRNNFKPTKYSNICSQHFTEDSFKRECNNR

VLKESAVPSLF

>026099822.1|Carassius auratus

MVQSCSAYGCKNRYHKDKNISFHKFPLARPDICGRWLAAMKRRNFKPTKYSNICSQHFTRDCFKHECNNR

VLKDNAVPSLF

>026153346.1|Mastacembelus armatus

MVQTCSAYGCKNRYHKDKDISFHKFPLARPDVCGKWVAAMRRNNFKPTKYSNICSQHFTKDCFKRECNNR

VLKENAVPSLF

>026196303.1|Anabas testudineus

MVQTCSAYGCKNRYHKDKDISFHKFPLARPDVCGKWVAAMRRNNFKPTKYSNICSQHFTKDCFKRECNNR

VLKESAVPSLF

>026777775.1|Pangasianodon hypophthalmus

MVQSCSAYGCKNRYHKDKDISFHKFPLARPDLCGKWVAAMKRRNFKPTKYSNICSQHFTKDCFKRECNNR

VLKENAVPSLF

>026874629.1|Electrophorus electricus

MVQSCSAYGCKNRYHKDKNISFHKFPLARPDVCGKWVAAMKRSNFKPTKYSNICSQHFTKDCFKRECNNR

VLKENAVPSLF

>026999612.1|Tachysurus fulvidraco

MVQSCSAYGCKNRYHKDKDVSFHKFPLARPDLCGKWVVAMKRRNFKPTKYSNICSQHFTKDCFKRECNNR

VLKENAVPSLF

>010751447.2|Larimichthys crocea

MVQTCSAYGCKNRYHKDKDISFHKFPLARPDVCGKWVAAMRRNNFKPTKYSNICSQHFTKDCFKRECNNR

VLKESAVPSLF

>028273835.1|Parambassis ranga

MVQTCSAYGCKNRYHKDKDISFHKFPLARPDVCGKWVAAMRRNNFKPTKYSNICSQHFTKDCFKRECNNR

VLKENAVPSLF

>028458130.1|Perca flavescens

MVQTCSAYGCKNRYNKDKDISFHKFPLARPDVCGKWVAAMRRNNFKPTKYSNICSHHFTKDCFKRDCNNR

VLKENAVPSLF

>028660878.1|Erpetoichthys calabaricus

MVQSCSAYGCKNRYHKDKNISFHKFPLARPDVCDKWVAAMRRNNFKPTKYSNICSQHFTKDCFKRECNNR

VLKESAVPSLF

>028827787.1|Denticeps clupeoides

MWRCQPGRDVLLDTARFSVVLATMVQSCSAYGCKNRYHKDKNISFHKFPLARPDVCGKWVAAMRRSNFKP

TKYSNICSQHFTKDSFKRECNNRVLKENAVPSLF

>010885922.1|Esox lucius

MVQSCSAYGCKNRYHKDKNISFHKFPLARPDVCGKWVAAMRRNNFKPTKYSNICSQHFTKDCFKRECNNR

VLKENAVPSLF

>029025762.1|Betta splendens

MVQTCSAYGCKNRYHKNNDISFHKFPLARPDICGKWVAAMRRNNFKPTKYSNICSQHFTKDSFKRECNNR

VLKENAVPSLF

>020495712.1|Labrus bergylta

MVQTCSAYGCKNRYNKDKDISFHKFPLARPDVCGKWVAAMRRNNFKPTKYSNICSQHFTKDCFKMECNNR

VLKDNAVPSLF

>029300536.1|Cottoperca gobio

MVQTCSAYGCKNRYHKDKDISFHKFPLARPDVCGKWVAAMRRNNFKPTKYSNICSQHFTKDCFKRECNNR

VLNENAVPLLF

>029372102.1|Echeneis naucrates

MVQTCSAYGCKNRYHKDRDISFHKFPLARPDVCGKWVAAMRRNNFKPTKYSNICSQHFTKDCFKRECNNR

VLKENAVPSLF

>029513109.1|Oncorhynchus nerka

MVQSCSAYGCKNRYHKDRNISFHKFPLARPDVCGKWVEAMRRNNFKPTRYSNICSQHFTKDSFKPECNNR

VLKENAVPSLF

>029566409.1|Salmo trutta

MVQSCSAYGCKNRYHKDKNISFHKFPLARPDVCGKWVAAMRRNNFKPTRYSNICSQHFTKDCFKPECNNR

VLKENAVPSLF

>029932319.1|Myripristis murdjan

MVQSCSAYGCKNRYNKDRNISFHKFPLARPDICGKWIAAMRRNNFKPTKYSNICSQHFTKDSFKMECNNR

VLKENAVPSLF

>029986140.1|Sphaeramia orbicularis

MVQTCSAYGCKNRYHKDKDISFHKFPLARPDVCGKWVAAMRRNNFKPTKYSNICSQHFTKDCFKRECNNR

VLKDNAVPSLF

>030292676.1|Sparus aurata

MLEHCAAYSCSNRRTIENRARGITFTMKPAWRVRKSTSGASYCISPSFSQCSRGAAMVQTCSAYGCKNRY

HKDKDISFHKFPLARPDICCKWVAAMRRNNFKPTKYSNICSQHFTKDCFKSECNNRVLKENAVPSLF

>030598472.1|Archocentrus centrarchus

MVQTCSAYGCKNRYHKDKDVSFHKFPLARPDVCCKWVAAMRRNNFKPTKYSNICSQHFTEDSFKRECNNR

VLKENAVPSLF

>030640807.1|Chanos chanos

MVQSCSAYGCKNRYHKDKNISFHKFPLARPDVCGKWVAAMRRKNFKPTKYSNICSQHFTKDCFKRECNNR

VLKENAVPSLF

>031143175.1|Sander lucioperca

MVQTCSAYGCKNRYNKDKDISFHKFPLARPDVCGKWVAAMRRNNFKPTKYSNICSHHFTKDCFKRDCNNR

VLKENAVPSLF

>012685494.2|Clupea harengus

MVQSCSAYGCKNRYHKDKNISFHKFPLARPDVCGKWVAAMRRHNFKPTKYSNICSQHFAKDCFKRECNNR

VLKENAVPSLF

>031613320.1|Oreochromis aureus

MVQTCSAYGCKNRYHKDKDISFHKFPLARPDICCKWVAAMRRNNFKPTKYSNICSQHFTEDSFKRECNNR

VLKESAVPSLF

>020329047.2|Oncorhynchus kisutch

MVQSCSAYGCKNRYHKDRNISFHKFPLARPDVCGKWVAAMRRNNFKPTRYSNICSQHFTKDCFKPECNNR

VLKENAVPSLF

>031731626.1|Anarrhichthys ocellatus

MVQTCSAYGCKNRYHKDKDISFHKFPLARPDVSAKWVAAMRRNNFKPTKYSNICSQHFTKDCFKRECNNR

VLKENAVPSLF

>032394995.1|Etheostoma spectabile

MVQTCSAYGCKNRYNKDKDISFHKFPLARPDVCGKWVAAMRRNNFKPTKYSNICSHHFTRDCFKRDCNNR

VLKENAVPSLF

>032425466.1|Xiphophorus hellerii

MVQTCSAYGCKNRYHKDKDISFHKFPLARPDVCRKWVAAMRRSNFKPTKYSNICSQHFSQDCFKRECNNR

VLKDNAVPSLF

>JAO42054.1|Poeciliopsis prolifica

MGRYKCAYNCETSTDPDVKFFKFPLYNPRKLKKWLSNMKLKDWAPTRFSVLCINHFEERHIDRTGKCVTL

RDDAVPTIF

>020825295.1|Phascolarctos cinereus

MVQSCSAYGCRNRYDKDKPVSFHKFPLTRPDLCKKWEAAVRRKNFKPSKYSSICSEHFTPDCFKRECNNK

LLKENAVPTIF

>027721249.1|Vombatus ursinus

MVQSCSAYGCRNRYDKDKPVSFHKFPLTRPDLCKKWEAAVRRKNFKPSKYSSICSEHFTPDCFKRECNNK

LLKENAVPTIF

>031806645.1|Sarcophilus harrisii

MVQSCSAYGCRNRYDKDKPVSFHKFPLTRPDLCKKWEAAVRRKNFKPSKYSSICSEHFTPDCFKRECNNK

LLKENAVPTIF

>PNI67599.1|Pan troglodytes

MVQSCSAYGCKNRYDKDKPVSFHKFPLTRPSLCKEWEAAVRRKNFKPTKYSSICSEHFTPDCFKRECNNK

LLKENAVPTIF

>PNJ34608.1|Pongo abelii

MVQSCSAYGCKNRYDKDKPVSFHKFPLTRPSLCKEWEAAVRRKNFKPTKYSSICSEHFTPDCFKRECNNK

LLKENAVPTIF

>AAH86347.1|Rattus norvegicus

MVQSCSAYGCKNRYDKDKPVSFHKFPLTRPSLCKQWEAAVRRKNFKPTKYSSICSEHFTPDCFKRECNNK

LLKENAVPTIF

>JAB42735.1|Callithrix jacchus

MVQSCSAYGCKNRYDKDKPVSFHKFPLTRPSLCKEWEAAVRRKNFKPTKYSSICSEHFTPDCFKRECNNK

LLKENAVPTIF

>AAH38639.1|Mus musculus

MVQSCSAYGCKNRYDKDKPVSFHKFPLTRPSLCKQWEAAVKRKNFKPTKYSSICSEHFTPDCFKRECNNK

LLKENAVPTIF

>AFE78994.1|Macaca mulatta

MVQSCSAYGCKNRYDKDKPVSFHKFPLTRPSLCKEWEAAVRRKNFKPTKYSSICSEHFTPDCFKRECNNK

LLKENAVPTIF

>JAN99934.1|Heterocephalus glaber

MVQSCSAYGCKNRYDKDKPVSFHKFPLTRPSLCKQWEAAVRRKNFKPTKYSSICSEHFTPDCFKRECNNK

LLKENAVPTIF

>CCP74018.1|Neovison vison

MVQSCSAYGCKNRYDKDKPVSFHKFPLTRPSLCKKWEAAVRRKNFKPTKYSSICSEHFTPDCFKRECNNK

LLKENAVPTIF

>020040280.1|Castor canadensis

MVQSCSAYGCKNRYDKDKPVSFHKFPLTRPSLCKKWEEAVKRKNFKPTKYSSICSEHFTPDCFKRECNNK

LLKENAVPTIF

>012632832.1|Microcebus murinus

MVQSCSAYGCKNRYDKDKPVSFHKFPLTRPSLCKKWEAAVRRKNFKPTKYSSICSEHFTPDCFKRECNNK

LLKENAVPTIF

>020743984.1|Odocoileus virginianus texanus

MVQSCSAYGCKNRYDKDKPVSFHKFPLTRPSLCKKWEAAVRRKNFKPTKYSSICSEHFTPDCFKRECNNK

LLKEDAVPTIF

>003359917.1|Sus scrofa

MVQSCSAYGCKNRYDKDKPVSFHKFPLTRPSLCKKWEAAVRRKNFKPTKYSSICSEHFTPDCFKRECNNK

LLKEDAVPTIF

>005066636.1|Mesocricetus auratus

MVQSCSAYGCKNRYDKDKPVSFHKFPLTRPSLCKQWEAAVRRKNFKPTKYSSICSEHFTPDCFKRECNNK

LLKENAVPTIF

>021092391.1|Heterocephalus glaber

MVQSCSAYGCKNRYDKDKPVSFHKFPLTRPSLCKQWEAAVRRKNFKPTKYSSICSEHFTPDCFKRECNNK

LLKENAVPTIF

>021495595.1|Meriones unguiculatus

MVQSCSAYGCKNRYDKDKPVSFHKFPLTRPGLCKQWEAAVRRKNFKPTKYSSICSEHFTPDCFKRECNNK

LLKENAVPTIF

>012331514.1|Aotus nancymaae

MVQSCSAYGCKNRYDKDKPVSFHKFPLTRPSLCKEWEAAVRRKNFKPTKYSSICSEHFTPDCFKRECNNK

LLKENAVPTIF

>008058071.1|Carlito syrichta

MVQSCSAYGCKNRYDKDKPVSFHKFPLTRPSLCKKWEAAVRRKNFKPTKYSSICSEHFTPDCFKRECNNK

LLKENAVPTIF

>005342564.1|Ictidomys tridecemlineatus

MVQSCSAYGCKNRYDKDKPVSFHKFPLTRPSLCEKWEAAVRRKNFKPTKYSSICSEHFTPDCFKRECNNK

LLKENAVPTIF

>848435.1|Canis lupus familiaris

MVQSCSAYGCKNRYDKDKPVSFHKFPLTRPSLCKKWEAAVRRKNFKPTKYSSICSEHFTPDCFKRECNNK

LLKENAVPTIF

>003984790.1|Felis catus

MVQSCSAYGCKNRYDKDKPVSFHKFPLTRPSLCKKWEAAVRRKNFKPTKYSSICSEHFTPDCFKRECNNK

LLKENAVPTIF

>003793957.1|Otolemur garnettii

MVQSCSAYGCKNRYDKDKPVSFHKFPLTRPNLCKKWEAAVRRKNFKPTKYSSICSEHFTPDCFKRECNNK

LLKENAVPTIF

>003412546.1|Loxodonta africana

MVQSCSAYGCKNRYDKDKPVSFHKFPLTRPSLCKKWEAAVRRKNFKPTKYSSICSEHFTPDCFKRECNNK

LLKENAVPTIF

>004478332.1|Dasypus novemcinctus

MVQSCSAYGCKNRYDKDKPVSFHKFPLTRPSLCKKWEAAVRRKNFKPTKYSSICSEHFTPDCFKRECNNK

LLKENAVPTIF

>003474674.1|Cavia porcellus

MVQSCSAYGCKNRYDKDKPVSFHKFPLTRPSLCKQWEAAVRRKNFKPTKYSSICSEHFTPDCFKRECNNK

LLKENAVPTIF

>011371229.1|Pteropus vampyrus

MVQSCSAYGCKNRYDKDKPVSFHKFPLTRPSLCKKWEEAVRRKNFKPTKYSSICSEHFTPDCFKRECNNK

LLKENAVPTIF

>023486531.1|Equus caballus

MVQSCSAYGCKNRYDKDKPVSFHKFPLTRPSLCKKWEAAVRRKNFKPTKYSSICSEHFTPDCFKRECNNK

LLKENAVPTIF

>004626723.1|Octodon degus

MVQSCSAYGCKNRYDKDKPVSFHKFPLTRPSLCKQWEAAVRRKNFKPTKYSSICSEHFTPDCFKRECNNK

LLKENAVPTIF

>006097091.1|Myotis lucifugus

MVQSCSAYGCKNRYHKDKPVSFHKFPLTRPNLCKEWEAAIRRKNFKPTKYSSICSEHFTPDCFKRECNNK

LLKENAVPTIF

>024434635.1|Desmodus rotundus

MVQSCSAYGCKNRYDKDKPVSFHKFPLTRPSLCKEWEAAVRRKNFKPTKYSSICSEHFTPDCFKRECNNK

LLKENAVPTIF

>024620782.1|Neophocaena asiaeorientalis

MKSCVSRTAAAWAVAGYRAWGRALSEGTRPTALGAASSGRCRGRSPRRDGLLLRRKPQNGMIVIVRCLHG

AIFIQPNSVFTSFTTLRKCGRNLRLFPLTRPSLCKKWEAAVRRKNFKPTKYSSICSEHFTPDCFKRECNN

KLLKEDAVPTIF

>011730789.1|Macaca nemestrina

MVQSCSAYGCKNRYDKDKPVSFHKFPLTRPSLCKEWEAAVRRKNFKPTKYSSICSEHFTPDCFKRECNNK

LLKENAVPTIF

>003823323.1|Pan paniscus

MVQSCSAYGCKNRYDKDKPVSFHKFPLTRPSLCKEWEAAVRRKNFKPTKYSSICSEHFTPDCFKRECNNK

LLKENAVPTIF

>006916323.1|Pteropus alecto

MVQSCSAYGCKNRYDKDKPVSFHKFPLTRPSLCKKWEEAVRRKNFKPTKYSSICSEHFTPDCFKRECNNK

LLKENAVPTIF

>015316367.1|Bos taurus

MRGARGRARGAEPGVFPRETCERGGAGRGPARDPVLRPRSGSGARLVSARDQFPLTRPSLCKKWEAAVRR

KNFKPTKYSSICSEHFTPDCFKRECNNKLLKEDAVPTIF

>006075782.1|Bubalus bubalis

MVQSCSAYGCKNRYDKDKPVSFHKFPLTRPSLCKKWEAAVRRKNFKPTKYSSICSEHFTPDCFKRECNNK

LLKEDAVPTIF

>025249042.1|Theropithecus gelada

MVQSCSAYGCKNRYDKDKPVSFHKFPLTRPSLCKEWEAAVRRKNFKPTKYSSICSEHFTPDCFKRECNNK

LLKENAVPTIF

>025299426.1|Canis lupus dingo

MVQSCSAYGCKNRYDKDKPVSFHKFPLTRPSLCKKWEAAVRRKNFKPTKYSSICSEHFTPDCFKRECNNK

LLKENAVPTIF

>025777369.1|Puma concolor

MVQSCSAYGCKNRYDKDKPVSFHKFPLTRPSLCKKWEAAVRRKNFKPTKYSSICSEHFTPDCFKRECNNK

LLKENAVPTIF

>025849469.1|Vulpes vulpes

MVQSCSAYGCKNRYDKDKPVSFHKFPLTRPSLCKKWEAAVRRKNFKPTKYSSICSEHFTPDCFKRECNNK

LLKENAVPTIF

>026236065.1|Urocitellus parryii

MVQSCSAYGCKNRYDKDKPVSFHKFPLTRPSLCEKWEAAVRRKNFKPTKYSSICSEHFTPDCFKRECNNK

LLKENAVPTIF

>026341519.1|Ursus arctos horribilis

MVQSCSAYGCKNRYDKDKPVSFHKFPLTRPSLCKKWEAAVRRKNFKPTKYSSICSEHFTPDCFKRECNNK

LLKENAVPTIF

>005362528.1|Microtus ochrogaster

MVQSCSAYGCKNRYDKDKPVSFHKFPLTRPSLCKQWEAAVRRKNFKPTKYSSICSEHFTPDCFKRECNNK

LLKENAVPTIF

>014943988.1|Acinonyx jubatus

MVQSCSAYGCKNRYDKDKPVSFHKFPLTRPSLCKKWEAAVRRKNFKPTKYSSICSEHFTPDCFKRECNNK

LLKENAVPTIF

>026978759.1|Lagenorhynchus obliquidens

MKSCVSRTAAAWAVAGYRAWGRALSEGTRPTALGAASSGRCRGRSPRRDGLLLRRKPQNGMIVIVRCLHG

AIFIQPNSVFTSFTTLRKCGRNLRLFPLTRPSLCKKWEAAVRRKNFKPTKYSSICSEHFTPDCFKRECNN

KLLKEDAVPTIF

>007622693.1|Cricetulus griseus

MVQSCSAYGCKNRYDKDKPVSFHKFPLTRPGLCKQWEAAVRRKNFKPTKYSSICSEHFTPDCFKRECNNK

LLKENAVPTIF

>027453850.1|Zalophus californianus

MVQSCSAYGCKNRYDKDKPVSFHKFPLTRPSLCKKWEAAVRRKNFKPTKYSSICSEHFTPDCFKRECNNK

LLKENAVPTIF

>027779542.1|Marmota flaviventris

MVQSCSAYGCKNRYDKDKPVSFHKFPLTRPSLCEKWEAAVRRKNFKPTKYSSICSEHFTPDCFKRECNNK

LLKENAVPTIF

>027818357.1|Ovis aries

MVQSCSAYGCKNRYDKDKPVSFHKFPLTRPSLCKKWEAAVRRKNFKPTKYSSICSDHFTPDCFKRECNNK

LLKEDAVPTIF

>027979823.1|Eumetopias jubatus

MVQSCSAYGCKNRYDKDKPVSFHKFPLTRPSLCKKWEAAVRRKNFKPTKYSSICSEHFTPDCFKRECNNK

LLKENAVPTIF

>007179057.1|Balaenoptera acutorostrata scammoni

MVQSCSAYGCKNRYDKDKPVSFHKFPLTRPSLCKKWEAAVRRKNFKPTKYSSICSEHFTPDCFKRECNNK

LLKEDAVPTIF

>028004529.1|Eptesicus fuscus

MVQSCSAYGCKNRYHKDKPVSFHKFPLTRPNLCKEWEAAIRRKNFKPTKYSSICSEHFTPDCFKRECNNK

LLKENAVPTIF

>001271632.1|Macaca fascicularis

MVQPCSAYGCKNRYDKDKPVSFHKFPLTRPSLCKEWEAAVRRKNFKPTKYSSICSEHFTPDCFKRECNNK

LLKENAVPTIF

>028382553.1|Phyllostomus discolor

MIIKCLQNVTRWQFYSAKFSTVFADFITFTFIFFLQEFPTINDRFPLTRPSLCKEWEAAVRRKNFKPTKY

SSICSEHFTPDCFKRECNNKLLKENAVPTIF

>028624164.1|Grammomys surdaster

MVQSCSAYGCKNRYDKDKPVSFHKFPLTRPSLCKQWEAAVKRKNFKPTKYSSICSEHFTPDCFKRECNNK

LLKENAVPTIF

>028710674.1|Peromyscus leucopus

MVQSCSAYGCKNRYDKDKPVSFHKFPLTRPSLCKQWEAAVRRKNFKPTKYSSICSEHFTPDCFKRECNNK

LLKENAVPTIF

>029082374.1|Monodon monoceros

MKSCVSRTAAAWAVAGCRAWGRALSEGTRPTALGAASSGRCRGRSPRRDGLLLRRKPQNGMIVIVRCLHG

AIFIQPNAVFTSFTTLRKCGRNLRLFPLTRPSLCKKWEAAVRRKNFKPTKYSSICSEHFTPDCFKRECNN

KLLKEDAVPTIF

>021025914.1|Mus caroli

MVQSCSAYGCKNRYDKDKPVSFHKFPLTRPSLCKQWEAAVKRKNFKPTKYSSICSEHFTPDCFKRECNNK

LLKENAVPTIF

>021075213.1|Mus pahari

MVQSCSAYGCKNRYDKDKPVSFHKFPLTRPSLCKQWEAAVKRKNFKPTKYSSICSEHFTPDCFKRECNNK

LLKENAVPTIF

>008847477.1|Nannospalax galili

MVQSCSAYGCKNRYDKDKPVSFHKFPLTRPSLCKQWEAAVRRKNFKPTKYSSICSEHFTPDCFKRECNNK

LLKENAVPTIF

>029796494.1|Suricata suricatta28382553.1

MVQSCSAYGCKNRYDKDKPVSFHKFPLTRPSLCKKWEAAVRRKNFKPTKYSSICSEHFTPDCFKRECNNK

LLKENAVPTIF

>022427054.1|Delphinapterus leucas

MKSCVSRTAAAWAVAGYRAWGRALSEGTRPTALGAASSGRCRGRSPRRDGLLLRRKPQNGMIVIVRCLHG

AIFIQPNSVFTSFTTLRKCGRNLRLFPLTRPSLCKKWEAAVRRKNFKPTKYSSICSEHFTPDCFKRECNN

KLLKEDAVPTIF

>003269699.1|Nomascus leucogenys

MVQSCSAYGCKNRYDKDKPVSFHKFPLTRPSLCKEWEAAVRRKNFKPTKYSSICSEHFTPDCFKRECNNK

LLKENAVPTIF

>012861339.1|Echinops telfairi

MVQSCSAYGCKNRYDKDKPVSFHKFPLTRPSLCKEWEAAVRRKNFKPTKYSSICSEHFTPDCFKRECNNK

LLKENAVPTIF

>030690375.1|Globicephala melas

MKSCVSRTAAAWAVAGYRAWGRALSEGTRPTALGAASSGRCRGRSPRRDGLLLRRKPQNGMIVIVRCLHG

AIFIQPNSVFTSFTTLRKCGRNLRLFPLTRPSLCKKWEAAVRRKNFKPTKYSSICSEHFTPDCFKRECNN

KLLKEDAVPTIF

>010354594.1|Rhinopithecus roxellana

MVQSCSAYGCKNRYDKDKPVSFHKFPLTRPSLCKEWEAAIRRKNFKPTKYSSICSEHFTPDCFKRECNNK

LLKENAVPTIF

>004047016.1|Gorilla gorilla

MVQSCSAYGCKNRYDKDKPVSFHKFPLTRPSLCKEWEAAVRRKNFKPTKYSSICSEHFTPDCFKRECNNK

LLKENAVPTIF

>006735410.1|Leptonychotes weddellii

MVQSCSAYGCKNRYDKDKPVSFHKFPLTRPSLCKKWEAAVRRKNFKPTKYSSICSEHFTPDCFKRECNNK

LLKENAVPTIF

>031195281.1|Mastomys coucha

MVQSCSAYGCKNRYDKDKPVSFHKFPLTRPSLCKQWEAAVKRKNFKPTKYSSICSEHFTPDCFKRECNNK

LLKENAVPTIF

>010984608.1|Camelus dromedarius

MVQSCSAYGCKNRYDKDKPVSFHKFPLTRPSLCKKWEAAVRRKNFKPTKYSSICSEHFTPDCFKRECNNK

LLKENAVPTIF

>003902757.1|Papio anubis

MVQSCSAYGCKNRYDKDKPVSFHKFPLTRPSLCKEWEAAVRRKNFKPTKYSSICSEHFTPDCFKRECNNK

LLKENAVPTIF

>015104124.1|Vicugna pacos

MVQSCSAYGCKNRYDKDKPVSFHKFPLTRPSLCKKWEAAVRRKNFKPTKYSSICSEHFTPDCFKRECNNK

LLKENAVPTIF

>023070206.1|Piliocolobus tephrosceles

MVQSCSAYGCKNRYDKDKPVSFHKFPLTRPSLCKEWEAAVRRKNFKPTKYSSICSEHFTPDCFKRECNNK

LLKENAVPTIF

>032143938.1|Sapajus apella

MVQSCSAYGCKNRYDKDKPVSFHKFPLTRPSLCKEWEAAVRRKNFKPTKYSSICSEHFTPDCFKRECNNK

LLKENAVPTIF

>032185099.1|Mustela erminea

MVQSCSAYGCKNRYDKDKPVSFHKFPLTRPSLCKKWEAAVRRKNFKPTKYSSICSEHFTPDCFKRECNNK

LLKENAVPTIF

>032269580.1|Phoca vitulina

MVQSCSAYGCKNRYDKDKPVSFHKFPLTRPSLCKKWEAAVRRKNFKPTKYSSICSEHFTPDCFKRECNNK

LLKENAVPTIF

>006188462.1|Camelus ferus

MVQSCSAYGCKNRYDKDKPVSFHKFPLTRPSLCKKWEAAVRRKNFKPTKYSSICSEHFTPDCFKRECNNK

LLKENAVPTIF

>032474305.1|Phocoena sinus

MKSCVSRTAAAWAVAGYRAWGRALSEGTRPTALGAASSGRCRGRSPRRDGLLLRRKPQNGMIVIVRCLHG

AIFIQPNSVFTSFTTLRKCGRNLRLFPLTRPSLCKKWEAAVRRKNFKPTKYSSICSEHFTPDCFKRECNN

KLLKEDAVPTIF

>031999962.1|Hylobates moloch

MVQSCSAYGCKNRYDKDKPVSFHKFPLTRPSLCKEWEAAVRRKNFKPTKYSSICSEHFTPDCFKRECNNK

LLKENAVPTIF

>032693882.1|Lontra canadensis

MVQSCSAYGCKNRYDKDKPVSFHKFPLTRPSLCKKWEAAVRRKNFKPTKYSSICSEHFTPDCFKRECNNK

LLKENAVPTIF

>032774333.1|Rattus rattus

MVQSCSAYGCKNRYDKDKPVSFHKFPLTRPSLCKQWEAAVRRKNFKPTKYSSICSEHFTPDCFKRECNNK

LLKENAVPTIF

>032960414.1|Rhinolophus ferrumequinum

MVQSCSAYGCKNRYDKDKPVSFHKFPLTRPSLCKKWEAAVRRKNFKPTKYSSICSEHFTPDCFKRECNNK

LLKENAVPTIF

>OWK00335.1|Cervus elaphus hippelaphus

MVQSCSAYGCKNRYDKDKPVSFHKFPLTRPSLCKKWEAAVRRKNFKPTKYSSICSEHFTPDCFKRECNNK

LLKEDAVPTIF

>RLQ76179.1|Cricetulus griseus

MVQSCSAYGCKNRYDKDKPVSFHKFPLTRPGLCKQWEAAVRRKNFKPTKYSSICSEHFTPDCFKRECNNK

LLKENAVPTIF

>48146629|Homo sapiens

MVQSCSAYGCKNRYDKDKPVSFHKFPLTRPSLCKEWEAAVRRKNFKPTKYSSICSEHFTPDCFKRECNNK

LLKENAVPTIF

>012732948.1|Fundulus heteroclitus

MVQTCSAYGCKNRYHKDKDISFHKFPLARPDICRKWVAAMRRSNFKPTKYSNICSQHFTQDCFKRECNNR

VLKESAVPSLF
