## Additional file for "Synapomorphic variations in the THAP domains of the human THAP protein family and its homologs": THAP2.docx

>AAH08358.1|Homo sapiens

MPTNCAAAGCATTYNKHINISFHRFPLDPKRRKEWVRLVRRKNFVPGKHTFLCSKHFEASCFDLTGQTRR

LKMDAVPTIF

>PRD25347.1|Trichonephila clavipes (Arthropoda)

MQQKKGVPLCLILKSLNCNHLKSFWYCHYHKTYSWMPSKCFVLGCRSNYCSELKVRVFSFPRDEILKNQW

LQAIQREDKRQPSKASKICALHFKAEDIITEASGFNSKTQEFITVPLQKHRLKPNAVPSVF

>GBN14036.1|Araneus ventricosus (Arthropoda)

MIKSCCAYECKEKFIKGGFVSFHRFPKNQERRYIWEKNIRRENFKATNSSLICSKHFTTDCFDTGKFGGT

WLKSDAVPTIF

>ACO14780.1|Caligus clemensi (Arthropoda)

MSCAAFGCKNQCSPKSDVSFHRFPRDPELRQEWVRNLKKADFTPKEHSRICSEHFTPECFNRTLNVVRLR

DNALPTIF

>CDW29737.1|Latimeria chalumnae (Chordata)

MGRKCSVPNCRSGYIGSNTLKGISFHALPRDPERKTHWLNVIPRKNWNPSLYSSICSLHFRASDYHENSL

DASRPSKHNTPLNLRILNKDAIPSLF

>JAB67011.1|Anoplophora glabripennis (Arthropoda)

MVSCLLCKNQMRHTRHKSFHRFPRNPKRYEKWIECINMVNWVPKKYDRICSDHFEKTDFYFSKNKKRLKI

DAVPNPN

>JAG79401.1|Fopius arisanus (Arthropoda)

MPTTCIVSSCKAGYRTNPERVSFFSIPKDKSLRDQWARAIRRDKGDLKSHHRVCSKHFRACDIILHKKFM

KPDGEIIAETSLNKSQLREGAVPMIF

>KOB78480.1|Operophtera brumata (Arthropoda)

MRANCSVYGCKSATYNKIGYMLNFHRFPKDPERQKIWKDLCKNPRIMDKHHTCIWMCSLHFEKTCYVFDF

LKHDAIPTLN

>GBP52097.1|Eumeta japonica (Arthropoda)

METEMKISTVYLSENRPKQTPIAKVAPELKQHREQNDSTSYEISKACAVLGCEESKNLDSDSFFRFPEDS

KHRQIWTELTGRSNWTPTDFSYICALHFSDDAFVLGEDSKMYLSNNAIPCLK

>PFX19131.1|Stylophora pistillata (Cnidaria)

MVYCTAYNCNNDGKKMKSLSFFQFPSDEKYRQIWIEKVRRVNWEPNKYSRLCSAHFEPHCFVHNFKLFES

LGLPKPKKATLKHDAIPTIF

>KXJ24883.1|Exaiptasia pallida (Cnidaria)

MPPTRCVVQGCSNKADREKGISIHSSPANSQMRAKWKKFVDIHRKNFSPEGQFGVCSDHFHGSCFTRGIH

MPGVKRVLIPXSVPTIW

>029469107.1|Rhinatrema bivittatum (Chordata)

MPTSCAAFDCTSVWKKDSNISFHRFPLQAERRKRWIQQVNRQNFSPSLHTFLCSKHFEDSCFDRTGQTVR

LRHDAVPTLF

>030042052.1|Microcaecilia unicolor (Chordata)

MPTSCAAFDCTSAWKKDSNISFHRFPLQAERRKQWIQQVNRQNFTPSLHTFLCSKHFEASCFDRTGQTVR

LRHDAVPTLF

>002935845.1|Xenopus tropicalis (Chordata)

MPTTCAASECKNNSKKDSQVTFHRFPADPKRRAEWLKHLNRENFFPTLHTFLCSKHFEESCFDRTGQTVR

LRANAVPTIF

>025039660.1|Pelodiscus sinensis (Chordata)

MPTSCAAAGCAAVYNKSVNVSFHRFPLDPKRRNEWIRLVKRNNFVPGKHTFLCSKHFESSCFDLTGQTRR

LRMDAVPTIF

>024064871.1|Terrapene carolina triunguis

MEQPGSRHLIKGKERVSPDHPVCRLREEMPTSCAAAGCAAVYNKSVNVSFHRFPLDPKRRKEWIRLVKRN

NFVPGKHTFLCSKHFESSCFDLTGQTRRLRMDAVPTIF

>030404593.1|Gopherus evgoodei

MQQPGCRHLIKGKERASLDRPVGRPREEMPTSCAAAGCAAVYNKSVNVSFHRYRPGWAGGGLGRLSRRFS

APEEAGFPLDPKRRKEWIRLVKRNNFVPGKHTFLCSKHFESSCFDLTGQTRRLRMDAVPTIF

>005280123.1|Chrysemys picta bellii

MEPPGRRHLIKGKERVSPDHPVCRLREKMPTSCAAAGCAAVYNKSVNVSFHRFPLDPKRRKEWIRLVKRN

NFVPGKHTFLCSKHFESSCFDLTGQTRRLRVDAVPTIF

>032659714.1|Chelonoidis abingdonii

MPTSCAAAGCAAVYNKSVNVSFHRFPLDPKRRKEWICLVKRNNFMPGKHTFLCSKHFESSCFDLTGQTRR

LRMDAVPTIF

>028674362.1|Erpetoichthys calabaricus

MPTSCSAMGCSSVYSKNGSVSFHRFPVDPKRKEIWIQNVRRENFIPGRYHFLCSKHFQPSCFDLTGQTKR

LRGDAVPTVF

>TNN59395.1|Liparis tanakae

MPSFDLLNRVELYMRAGTFPLESSKSSKKVTKAASKHFIYKDGCLLRSYRGRLLRVVRSDEEVREILARY

HDNNSHAGRVRVVKEIMLMYYWVGVTEAVKAWIQDCAVCQSRTPVEPPAPPVQFCLAYGCDASSYVHPQL

SFHRFPKDAERRRRWLVLSQRDEGSLRTNSCLCSRHFEPACFSPSEAGQTTLSPDAVPTVP

>JAR77966.1|Fundulus heteroclitus

MPTCVAFGCNNKQFSGCGRTFHRFPHGNAERMKQWVLNVRRKKWQPSKTSVLCSEHFEEQCFDRTGQTVR

LREGALPTLF

>027726596.1|Vombatus ursinus

MRGVREMPTNCAAAGCAATYNKHINVSFHRFPLDPQRRKEWVRLVRRKNFVPGKHTFLCSKHFEASCFDL

TGQTRRLKMDAVPTIF

>028934293.1|Ornithorhynchus anatinus

MPTNCAAAGCAATYDKHINVSFHRFPLDPKRRKEWVRLVRRKNFVPGKHAFLCSKHFEASCFDLTGQTRR

LKMDAVPTIF

>003770984.1|Sarcophilus harrisii

MPTNCAAAGCAATYNKHINISFHRFPLDPQRRKEWVRLVRRKNFVPGKHTFLCSKHFEASCFDLTGQTRR

LKMDAVPTIF

>JAA35220.1|Pan troglodytes

MPTNCAAAGCATTYNKHINISFHRFPLDPKRRKEWVRLVRRKNFVPGKHTFLCSKHFEASCFDLTGQTRR

LKMDAVPTIF

>JAB46299.1|Callithrix jacchus

MPTNCAAAGCATTYNKHINISFHRFPLDPKRRKEWVRLVRRKNFVPGKHTFLCSKHFEASCFDLTGQTRR

LKMDAVPTIF

>012594307.1|Microcebus murinus

MPTNCAAAGCATTYNKHINISFHRFPLDPKRRKEWVRLVRRKNFVPGKHTFLCSKHFEASCFDLTGQTRR

LKMDAVPTIF

>012296733.1|Aotus nancymaae

MPTNCAAAGCATTYNKHINISFHRFPLDPKRRKEWVRLVRRKNFVPGKHTFLCSKHFEASCFDLTGQTRR

LKMDAVPTIF

>008053425.1|Carlito syrichta

MPTNCAAAGCATTYNKHINISFHRFPLDPKRRKEWVRLVRRKNFVPGKHTFLCSKHFEASCFDLTGQTRR

LKMDAVPTIF

>003798992.2|Otolemur garnettii

MPTNCAAAGCATTYNKHINISFHRFPLDPKRRKEWVRLVRRKNFVPGKHTFLCSKHFEASCFDLTGQTRR

LKMDAVPTIF

>011725460.1|Macaca nemestrina

MPTNCAAAGCATTYNKHINISFHRFPLDPKRRKEWVRLVRRKNFVPGKHTFLCSKHFEASCFDLTGQTRR

LKMDAVPTIF

>003832962.1|Pan paniscus

MPTNCAAAGCATTYNKHINISFHRFPLDPKRRKEWVRLVRRKNFVPGKHTFLCSKHFEASCFDLTGQTRR

LKMDAVPTIF

>025257522.1|Theropithecus gelada

MPTNCAAAGCATTYNKHINISFHRFPLDPKRRKEWVRLVRRKNFVPGKHTFLCSKHFEASCFDLTGQTRR

LKMDAVPTIF

>001153224.1|Pongo abelii

MPTNCAAAGCATTYNKHINISFHRFPLDPKRRKEWVRLVRRKNFVPGKHTFLCSKHFEASCFDLTGQTRR

LKMDAVPTIF

>003259595.1|Nomascus leucogenys

MPTNCAAAGCATTYNKHINISFHRFPLDPKRRKEWVRLVRRKNFVPGKHTFLCSKHFEASCFDLTGQTRR

LKMDAVPTIF

>010369030.1|Rhinopithecus roxellana

MPTNCAAAGCATTYNKHINISFHRFPLDPKRRKEWVRLVRRKNFVPGKHTFLCSKHFEASCFDLTGQTRR

LKMDAVPTIF

>004053623.2|Gorilla gorilla gorilla

MPTNCAAAGCATTYNKHINISFHRFPLDPKRRKEWVRLVRRKNFVPGKHTFLCSKHFEASCFDLTGQTRR

LKMDAVPTIF

>021778294.1|Papio anubis

MPTNCAAAGCATTYNKHINISFHRFPLDPKRRKEWVRLVRRKNFVPGKHTFLCSKHFEASCFDLTGQTRR

LKMDAVPTIF

>001180430.1|Macaca mulatta

MPTNCAAAGCATTYNKHINISFHRFPLDPKRRKEWVRLVRRKNFVPGKHTFLCSKHFEASCFDLTGQTRR

LKMDAVPTIF

>023082280.1|Piliocolobus tephrosceles

MPTNCAAAGCATTYNKHINISFHRFPLDPKRRKEWVRLVRRKNFVPGKHTFLCSKHFEASCFDLTGQTRR

LKMDAVPTIF

>032151615.1|Sapajus apella

MPTNCAAAGCATTYNKHINISFHRFPLDPKRRKEWVRLVRRKNFVPGKHTFLCSKHFEASCFDLTGQTRR

LKMDAVPTIF

>032026636.1|Hylobates moloch

MPTNCAAAGCATTYNKHINISFHRFPLDPKRRKEWVRLVRRKNFVPGKHTFLCSKHFEASCFDLTGQTRR

LKMDAVPTIF

>033077156.1|Trachypithecus francoisi

MPTNCAAAGCATTYNKHINISFHRFPLDPKRRKEWVRLVRRKNFVPGKHTFLCSKHFEASCFDLTGQTRR

LKMDAVPTIF

>AAH27502.1|Mus musculus

MPTNCAAAGCAATYNKHINISFHRFPLDPKRRKEWVRLVRRKNFVPGKHTFLCSKHFEASCFDLTGQTRR

LKMDAVPTIF

>020034574.1|Castor canadensis

MPTNCAAAGCATTYNKHINISFHRFPLDPKRRKEWVRLVRRKNFVPGKHTFLCSKHFEASCFDLTGQTRR

LKMDAVPTIF

>020753140.1|Odocoileus virginianus texanus

MPTNCAAAGCATTYNKHINISFHRFPLDPKRRKEWVRLVRRKNFVPGKHTFLCSKHFEASCFDLTGQTRR

LKMDAVPTIF

>020824527.1|Phascolarctos cinereus

MPTNCAAAGCAATYNKHINVSFHRFPLDPQRRKEWVRLVRRKNFVPGKHTFLCSKHFEASCFDLTGQTRR

LKMDAVPTIF

>003126431.3|Sus scrofa

MPTNCAAAGCATTYNKHINISFHRFPLDPKRRKEWVRLVRRKNFVPGKHTFLCSKHFEASCFDLTGQTRR

LKMDAVPTIF

>004845090.1|Heterocephalus glaber

MPTNCAAAGCATTYNKHINISFHRFPMDPKRRKEWVRLVRRKNFVPGKHTFLCSKHFEASCFDLTGQTRR

LKMDAVPTIF

>021500514.1|Meriones unguiculatus

MPTNCAAAGCAATYNKHINISFHRFPLDPKRRKEWVRLVRRKNFVPGKHTFLCSKHFEASCFDLTGQTRR

LKMDAVPTIF

>021534331.1|Neomonachus schauinslandi

MPTNCAAAGCATTYNKHINISFHRFPLDPKRRKEWVRLLRRKNFVPGKHTFLCSKHFEASCFDLTGQTRR

LKMDAVPTIF

>005330473.1|Ictidomys tridecemlineatus

MPTNCAAAGCATTYNKHINISFHRFPLDPKRRKEWVRLVRRKNFVPGKHTFLCSKHFEASCFDLTGQTRR

LKMDAVPTIF

>003431484.1|Canis lupus familiaris

MPTNCAAAGCATTYDKHINISFHRFPLDPKRRKEWIRLLRRKNFVPGKHTFLCSKHFEASCFDLTGQTRR

LKMDAVPTIF

>011282427.1|Felis catus

MPTNCAAAGCATTYNKHINISFHRFPLDPKRRKEWVRLLRRKNFVPGKHTFLCSKHFEASCFDLTGQTRR

LKMDAVPTIF

>023414708.1|Loxodonta africana

MPTNCAAAGCAATYNKHINISFHRFPLDPKRRKEWVRLVRRKNFVPGKHTFLCSKHFEASCFDLTGQTRR

LKMDAVPTIF

>023445582.1|Dasypus novemcinctus

MLNAPTTQQGKDPLPRLPSLCQQETQQPAPRNGLKTAPNEGEREMPTNCAAAGCAATYNKHINISFHRFP

LDPKRRKEWVRLVRRKNFVPGKHTFLCSKHFEASCFDLTGQTRRLKMDAVPTIF

>011372622.1|Pteropus vampyrus

MPTNCAAAGCATTYNKHINISFHRFPLDPKRRKEWVRLVRRKNFVPGKHTFLCSKHFEASCFDLTGQTRR

LKMDAVPTIF

>023499789.1|Equus caballus

MAPPPLGCLGAWRGRVRSSSLPLAVDTNGRGEGRQLCGWTAPKTAARPRRGGPAGAGGREGWVRPPCWWR

RRAATERRERGRRAGEMPTHCAAAGCAATYDKHADVSFHRFPLDPKRRKEWVRLVRRKNFVPGKHTFLCS

KHFEASCFDLTGQTRRLKMDAVPTIF

>004632227.1|Octodon degus

MPTNCAAAGCAATYNKHINISFHRFPLDPKRRKEWVRLVRRKNFVPGKHTFLCSKHFEASCFDLTGQTRR

LKMDAVPTIF

>014316967.1|Myotis lucifugus

MPTNCAAAGCATTYNKHINISFHRFPLDPKRRKEWVRLVRRKNFVPGKHTFLCSKHFEASCFDLTGQTRR

LKMDAVPTIF

>024432224.1|Desmodus rotundus

MPTNCAAAGCATTYNKHINISFHRFPLDPKRRKEWVRLTRRKNFVPGKHTFLCSKHFEASCFDLTGQTRR

LKMDAVPTIF

>024592227.1|Neophocaena asiaeorientalis

MPTNCAAAGCATTYNKHINISFHRFPLDPKRRKEWVRLVRRKNFVPGKHTFLCSKHFEASCFDLTGQTRR

LKMDAVPTIF

>003586080.2|Bos taurus

MPTNCAAAGCATTYNKHINISFHRFPLDPKRRKEWVRLVRRKNFVPGKHTFLCSKHFEASCFDLTGQTRR

LKMDAVPTIF

>006923811.2|Pteropus alecto

MPTNCAAAGCATTYNKHINISFHRFPLDPKRRKEWVRLVRRKNFVPGKHTFLCSKHFEASCFDLTGQTRR

LKMDAVPTIF

>006062203.2|Bubalus bubalis

MPTNCAAAGCATTYNKHINISFHRFPLDPKRRKEWVRLVRRKNFVPGKHTFLCSKHFEASCFDLTGQTRR

LKMDAVPTIF

>025738484.1|Callorhinus ursinus

MPTNCAAAGCATTYNKHINISFHRFPLDPKRRKEWVRLLRRKNFVPGKHTFLCSKHFEASCFDLTGQTRR

LKMDAVPTIF

>025782638.1|Puma concolor

MPTNCAAAGCATTYNKHINISFHRFPLDPKRRKEWVRLLRRKNFVPGKHTFLCSKHFEASCFDLTGQTRR

LKMDAVPTIF

>025857498.1|Vulpes vulpes

MPTNCAAAGCATTYDKHINISFHRFPLDPKRRKEWIRLLRRKNFVPGKHTFLCSKHFEASCFDLTGQTRR

LKMDAVPTIF

>026242386.1|Urocitellus parryii

MPTNCAAAGCATTYNKHINISFHRFPLDPKRRKEWVRLVRRKNFVPGKHTFLCSKHFEASCFDLTGQTRR

LKMDAVPTIF

>026355806.1|Ursus arctos horribilis

MPTNCAAAGCATTYNKHINISFHRFPLDPKRRKEWVRLLRRKNFVPGKHTFLCSKHFEASCFDLTGQTRR

LKMDAVPTIF

>005358108.1|Microtus ochrogaster

MPTNCAAAGCAATYNKHINISFHRFPLDPKRRKEWVRLVRRKNFVPGKHTFLCSKHFEASCFDLTGQTRR

LKMDAVPTIF

>014919107.1|Acinonyx jubatus

MPTNCAAAGCATTYNKHINISFHRFPLDPKRRKEWVRLLRRKNFVPGKHTFLCSKHFEASCFDLTGQTRR

LKMDAVPTIF

>026962838.1|Lagenorhynchus obliquidens

MPTNCAAAGCATTYNKHINISFHRFPLDPKRRKEWVRLVRRKNFVPGKHTFLCSKHFEASCFDLTGQTRR

LKMDAVPTIF

>027448867.1|Zalophus californianus

MPTNCAAAGCATTYNKHINISFHRFPLDPKRRKEWVRLLRRKNFVPGKHTFLCSKHFEASCFDLTGQTRR

LKMDAVPTIF

>006146251.1|Tupaia chinensis

MPTNCAAAGCATTYNKHINISFHRFPLDPKRRKEWVRLVKRKNFVPGKHTFLCSKHFEASCFDLTGQTRR

LKIDAVPTIF

>027778806.1|Marmota flaviventris

MPTNCAAAGCATTYNKHINISFHRFPLDPKRRKEWVRLVRRKNFVPGKHTFLCSKHFEASCFDLTGQTRR

LKMDAVPTIF

>027822936.1|Ovis aries

MPTNCAAAGCATTYNKHINISFHRFPLDPKRRKEWVRLVRRKNFVPGKHTFLCSKHFEASCFDLTGQTRR

LKMDAVPTIF

>027966591.1|Eumetopias jubatus

MPTNCAAAGCATTYNKHINISFHRFPLDPKRRKEWVRLLRRKNFVPGKHTFLCSKHFEASCFDLTGQTRR

LKMDAVPTIF

>008153769.1|Eptesicus fuscus

MPTNCAAAGCATTYNKHINISFHRFPLDPKRRKEWVRLVRRKNFVPGKHTFLCSKHFEASCFDLTGQTRR

LKMDAVPTIF

>007195075.2|Balaenoptera acutorostrata scammoni

MPTNCAAAGCATTYNKHINISFHRFPLDPKRRKEWVRLVRRKNFVPGKHTFLCSKHFEASCFDLTGQTRR

LKMDAVPTIF

>007111635.2|Physeter catodon

MPTNCAAAGCATTYNKHINISFHRFPLDPKRRKEWVRLVRRKNFVPGKHTFLCSKHFEASCFDLTGQTRR

LKMDAVPTIF

>028388605.1|Phyllostomus discolor

MPTNCAAAGCATTYNKHINISFHRFPLDPKRRKEWVRLTRRKNFVPGKHTFLCSKHFEASCFDLTGQTRR

LKMDAVPTIF

>028614920.1|Grammomys surdaster

MPTNCAAAGCAATYNKHINISFHRFPLDPTRRKEWVRLVRRKNFVPGKHTFLCSKHFEASCFDLTGQTRR

LKMDAVPTIF

>028733805.1|Peromyscus leucopus

MPTNCAAAGCAATYDKHINVSFHRFPLDPKRRKEWVRLVRRKNFVPGKHTFLCSKHFEASCFDLTGQTRR

LKMDAVPTIF

>029062229.1|Monodon monoceros

MPTNCAAAGCATTYNKHINISFHRFPLDPKRRKEWVRLVRRKNFVPGKHTFLCSKHFEASCFDLTGQTRR

LKMDAVPTIF

>021030026.1|Mus caroli

MPTNCAAAGCAATYNKHINISFHRFPLDPKRRKEWVRLVRRKNFVPGKHTFLCSKHFEASCFDLTGQTRR

LKMDAVPTIF

>021060924.1|Mus pahari

MPTNCAAAGCAATYNKHINISFHRFPLDPKRRKEWVRLVRRKNFVPGKHTFLCSKHFEASCFDLTGQTRR

LKMDAVPTIF

>029420111.1|Nannospalax galili

MPTNCAAAGCAATYNKHINISFHRFPLDPKRRKEWIRLVRRKNFVPGKHTFLCSKHFEASCFDLTGQTRR

LKMDAVPTIF

>029809077.1|Suricata suricatta

MPTNCAAAGCATTYNKHINISFHRFPLDPKRRKEWVRLLRRKNFVPGKHTFLCSKHFEASCFDLTGQTRR

LKMDAVPTIF

>030179100.1|Lynx canadensis

MPTNCAAAGCATTYNKHINISFHRFPLDPKRRKEWVRLLRRKNFVPGKHTFLCSKHFEASCFDLTGQTRR

LKMDAVPTIF

>022427222.1|Delphinapterus leucas

MPTNCAAAGCATTYNKHINISFHRFPLDPKRRKEWVRLVRRKNFVPGKHTFLCSKHFEASCFDLTGQTRR

LKMDAVPTIF

>004718231.|Echinops telfairi

MPTNCAAAGCAATYNKHINISFHRFPLDPKRRKEWVRLVRRKNFVPGKHTFLCSKHFEASCFDLTGQTRR

LKMDAVPTIF

>030722134.1|Globicephala melas

MPTNCAAAGCATTYNKHINISFHRFPLDPKRRKEWVRLVRRKNFVPGKHTFLCSKHFEASCFDLTGQTRR

LKMDAVPTIF

>006744583.1|Leptonychotes weddellii

MPTNCAAAGCATTYNKHINISFHRFPLDPKRRKEWVRLLRRKNFVPGKHTFLCSKHFEASCFDLTGQTRR

LKMDAVPTIF

>031206205.1|Mastomys coucha

MPTNCAAAGCAATYNKHINISFHRFPLDPKRRKEWVRLVRRKNFVPGKHTFLCSKHFEASCFDLTGQTRR

LKMDAVPTIF

>010986523.1|Camelus dromedarius

MPTNCAAAGCATTYNKHINISFHRFPLDPKRRKEWVRLVRRKNFVPGKHTFLCSKHFEASCFDLTGQTRR

LKMDAVPTIF

>006197892.2|Vicugna pacos

MPTNCAAAGCATTYNKHINISFHRFPLDPKRRKEWVRLVRRKNFVPGKHTFLCSKHFEASCFDLTGQTRR

LKMDAVPTIF

>032972237.1|Rhinolophus ferrumequinum

MPTNCAAAGCATTYNKHINISFHRFPLDPKRRKEWVRLVRRKNFVPGKHTFLCSKHFEASCFDLTGQTRR

LKMDAVPTIF

>032768435.1|Rattus rattus

MPTNCAAAGCAATYNKHINISFHRFPLDPKRRKEWVRLVRRKNFVPGKHTFLCSKHFEASCFDLTGQTRR

LKMDAVPTIF

>032710684.1|Lontra canadensis

MPTNCAAAGCATTYNKHINISFHRFPLDPKRRKEWIRLLRRKNFVPGKHTFLCSKHFEASCFDLTGQTRR

LKMDAVPTIF

>032500555.1|Phocoena sinus

MPTNCAAAGCATTYNKHINISFHRFPLDPKRRKEWVRLVRRKNFVPGKHTFLCSKHFEASCFDLTGQTRR

LKMDAVPTIF

>006191167.2|Camelus ferus

MPTNCAAAGCATTYNKHINISFHRFPLDPKRRKEWVRLVRRKNFVPGKHTFLCSKHFEASCFDLTGQTRR

LKMDAVPTIF

>032276406.1|Phoca vitulina

MPTNCAAVGCATTYNKHINISFHRFPLDPKRRKEWVRLLRRKNFVPGKHTFLCSKHFEASCFDLTGQTRR

LKMDAVPTIF

>032202918.1|Mustela erminea

MPTNCAAAGCATTYNKHINISFHRFPLDPKRRKEWIRLLRRKNFVPGKHTFLCSKHFEASCFDLTGQTRR

LKMDAVPTIF
