## Additional file for "Synapomorphic variations in the THAP domains of the human THAP protein family and its homologs": THAP3.docx

>JAG74305.1|Fopius arisanus (Arthropoda)

MRCVYCGKAKEKNKEISFHRFPKDEKLVKTWVMNMGRLDLVPKQWHMLCAEHFSKTCIDFRDLRARLRKG

SVPTIF

>ACO14806.1|Caligus clemensi (Arthrooda)

MPNCCVAPKCRTGFPDVPRDPMVSLYRFAFSNPELLREWLRAILGSNWEPTINSQLCGAHFRNQAFKRNP

SGSNPYIQRLSVRRSLKDDAIPSIF

>PRD29195.1|Trichonephila clavipes (Arthropoda)

MSHCAAYGCSNTKRKESCRNKTFFKFPLKKPILLKTWILMIRRDKFKPSSSSLLCSDHFEDECFAYQNFT

NRRQLKPGSIPTIF

>KXJ14245.1|Exaiptasia pallid (Cnidaria)

MSKFTYCVADCHNRGITGVTFHRFPADQELRXKWLIKIRQGDFKITANTRICSSHFHESDFKPFKDNLGR

RLLVYNAIPTQF

>PFX15097.1|Stylophora pistillata (Cnidaria)

MRSALFLYPESDSDKARRLMSYYQGKLGKIAVVRGIKNILAREQDFGKQLRLSECVVLVGTRQASFLIRT

KQQEKDDDFITFDGKVIHEEFTENKEIVKNRLLIVHFTERNEDDWIPKGFDERRIFHVEDGIVPPDGSPT

LTHLEYRMRKVLLGDDFIRNRLTQERRSRDLKWVSKSKQTHFGSHVEKFILKMPHFCCAGECENSSDKRP

DISFHGLPLDNKALLKTWIAKMRRNPNYFNVNKHVKICSKHFSPEDFINPDAKKRRLKRNAVPSIF

>025062204.1|Alligator sinensis (Chordata)

MDFQPSCHAVLCSQHFQPDCFSPCGNRANLRPDAVPTLF

>029434045.1|Rhinatremabivittatum (Chordata)

MPKSCAVPYCTHRYSSKNKQLTFHRFPFSKPELFKEWLGNVGRLNFKPKQHTVICSEHFTPDCFSAFGNR

KNLIPNAIPTIF

>030041702.1|Microcaecilia unicolor (Chordata)

MPKSCAVPYCTRRYSSKNKQLTFHRFPFSKPDLFKEWVGNVGRLDFKPKQHTVICSEHFTPDCFSAIGNR

KNLMPNAIPTIF

>032904038.1|Amblyraja radiate (Chordata)

MPKSCAAFNCTNRYSSKNKELTFHRFPFSKPDLLIEWMNNVGRAEFKPNQHTVICSDHFKPECFNTWGNR

KNLKHNAVPTVF

>005287121.1|Chrysemys pictabellii (Chordata)

MPKSCAALRCSNRYSSRCRQQLTFHRFPFSRPELLARWVGNIGRGDFQPSSHTVLCSQHFQPDCFSAFGN

RTNLKPNAVPTLF

>014429141.1|Pelodiscus sinensis (Chordata)

MDPAVCPRFAWPATVTSPPSFHVLSTGSKVLPPPTSATGTRVPPLPLLLWLRPFSTLPLPCRFPFSRPEL

LARWVGNIGRGDFQPSSHTVLCSQHFQPDCFSAFGNRTNLKPNAVPTLF

>027678536.1|Chelonia mydas

MIHYLPEAKWNQEFAKDLPLRAVFCSQINNQSSQVLLFSLQHKRLWKKILFTVCVPGPGPSPGHSLPHPA

VSGRFPLSRPELLARWVGNIGRGNFQPSSHTVLCSQHFQPDCFSAFGNRTNLKPNAVPTLF

>026515911.1|Terrapene carolinatriunguis

MPKSCAALRYSNRYSSRCRQQLTFHRFPFSRPELLARWVGNIGRGDFQPSSHTVLCSQHFQPDCFSAFGN

RTNLKPNAVPTLF

>030393185.1|Gopherus evgoodei

MPKSCAALRCSNRYSSRCRQQLTFHRYRGRAPRRFPFSRPELLARWVGNIGRGDFQPSSHTVLCSQHFQP

DCFSAFGNRTNLKPNAVPTLF

>032619750.1|Chelonoidis abingdonii

MPKSCAALRCSNRYSSRCRQQLTFHRFPFSRPELLARWVGNIGRGDFQPSSHTVLCSQHFQPDCFSAFGN

RTNLKPNAVPTLF

>AAH91927.1|Danio rerio

MPKSCSASNCTNRYNNKNPEITFHRFPFSKPSVLKQWLDNIGRDDFQPRKHMVICSLHFTPDCFSGLGNR

KNLLWNAVPTLF

>023861884.1|Salvelinus alpinus

MPKSCSAYNCTNRYSKHTEITFHRFPFSKPTVFRQWLDNICRDDFYPKKHMVICSLHFTTDCFSGLGNRK

NLLWNAVPTLF

>028663222.1|Erpetoichthys calabaricus

MPKSCAAYNCTNRYSNKKGKLTFHRFPLSKPHILKEWLDNIGRENFQPNQHMVLCSEHFTPDCFSGFGNR

KNLIWNAVPTIF

>020830574.1|Phascolarctos cinereus

MPKSCAARQCCNRYSSRRKQLTFHRFPFSRPELLKEWVLNIGRGNFKPKQHTVICSEHFQPDCFSAFGNR

KNLKQNAVPTVF

>028921210.1|Ornithorhynchus anatinus

MDEVLMKRMFSRFFGSALSTSRDSGTPFSQALGPWPPGRFPFSRPELLKEWVLNIGRSNFKPKQHTVICS

EHFKPECFSAFGNRKNLKQNAVPTVF

>031818801.1|Sarcophilus harrisii

MFPFSRPELLKEWVLNIGRGNFKPKQHTVICSEHFRPDCFSAFGNRKNLKQNAVPTVF

>PNI39894.1|Pan troglodytes

MPKSCAARQCCNRYSSRRKQLTFHRFPFSRPELLKEWVLNIGRGNFKPKQHTVICSEHFRPECFSAFGNR

KNLKHNAVPTVF

>PNJ48066.1|Pongo abelii

MPKSCAARQCCNRYSSRRKQLTFHRFPFSRPELLKEWVLNIGRGNFKPKQHTVICSEHFRPECFSAFGNR

KNLKHNAVPTVF

>JAB52274.1|Callithrix jacchus

MPKSCAARQCCNRYSSRRKQLTFHRFPFSRPELLKEWVLNIGRGNFKPKQHTVICSEHFRPECFSAFGNR

KNLKHNAVPTVF

>012646974.1|Microcebus murinus

MPKSCAARQCCNRYNSRRKQLTFHRFPFSRPELLKEWVLNIGRGNFKPKQHTVICSEHFKPECFSAFGNR

KNLKHNAVPTVF

>012329066.1|Aotus nancymaae

MPKSCAARQCCNRYSSRRKQLTFHRFPFSRPELLKEWVLNIGRGNFKPKQHTVICSEHFRPECFSAFGNR

KNLKHNAVPTVF

>008053875.1|Carlito syrichta

MRGLHGPAVLGQPRWSPVSPAFVCRFPFSRPELLKEWVLKIGRGNFKPKQHTVICSEHFRPECFSAFGNR

KNLKHNAVPTVF

>012662778.1|Otolemur garnettii

MPKSCAARQCCNRYSSRRKQLTFHRFPFSRPELLKEWVLNIGRGNFKPKQHTVICSEHFKPECFSAFGNR

KNLKHNAVPTVF

>011735733.1|Macaca nemestrina

MPKSCAARQCCNRYSSRRKQLTFHRFPFSRPELLKEWVLNIGRGNFQPKQHTVICSEHFRPECFSAFGNR

KNLKHNAVPTVF

>025232824.1|Theropithecus gelada

MPKSCAARQCCNRYSSRRKQLTFHRFPFSRPELLKEWVLNIGRGNFQPKQHTVICSEHFRPECFSAFGNR

KNLKHNAVPTVF

>001094480.1|Macaca mulatta

MPKSCAARQCCNRYSSRRKQLTFHRFPFSRPELLKEWVLNIGRGNFQPKQHTVICSEHFRPECFSAFGNR

KNLKHNAVPTVF

>030661318.1|Nomascus leucogenys

MPKSCAARQCCNRYSSRRKQLTFHRFPFSRPELLKEWVLNIGRGNFKPKQHTVICSEHFRPECFSAFGNR

KNLKHNAVPTVF

>010355688.2|Rhinopithecus roxellana

MPKSCAARQCCNRYSSRRKQLTFHRFPFSRPELLKEWVLNIGRANFQPKQHTVICSEHFRPECFSAFGNR

KNLKHNAVPTVF

>004024618.1|Gorilla gorilla gorilla

MPKSCAARQCCNRYSSRRKQLTFHRFPFSRPELLKAWVLNIGRGNFKPKQHTVICSEHFRPECFSAFGNR

KNLKHNAVPTVF

>031519963.1|Papio anubis

MPKSCAARQCCNRYSSRRKQLTFHRFPFSRPELLKEWVLNIGRGNFQPKQHTVICSEHFRPECFSAFGNR

KNLKHNAVPTVF

>023070992.1|Piliocolobus tephrosceles

MPKSCAARQCCNRYSSRRKQLTFHRFPFSRPELLKEWVLNIGRGNFQPKQHTVICSEHFRPECFSAFGNR

KNLKHNAVPTVF

>032156151.1|Sapajus apella

MPKSCAARQCCNRYSSRRKQLTFHRFPFSRPELLKEWVLNIGRGNFKPKQHTVICSEHFRPECFSAFGNR

KNLKHNAVPTVF

>032614658.1|Hylobates moloch

MPKSCAARQCCNRYSSRRKQLTFHRFPFSRPELLKEWVLNIGRGNFKPKQHTVICSEHFRPECFSAFGNR

KNLKHNAVPTVF

>033084449.1|Trachypithecus francoisi

MPKSCAARQCCNRYSSRRKQLTFHRFPFSRPELLKEWVLNIGRANFQPKQHTVICSEHFRPECFSAFGNR

KNLKHNAVPTVF

>AAI20271.1|Bos taurus

MPKSCAARQCCNRYSNRRKQLTFHRFPFSRPELLKEWVLNIGRGDFEPKQHTVICSEHFRPECFSAFGNR

KNLKHNAVPTVF

>AAI59428.1|Rattus norvegicus

MPKSCAARQCCNRYSSRRKQLTFHRFPFSRPELLREWVLNIGRADFKPKQHTVICSEHFRPECFSAFGNR

KNLKHNAVPTVF

>AAH94395.1|Mus musculus

MPKSCAARQCCNRYSSRRKQLTFHRFPFSRPELLREWVLNIGRADFKPKQHTVICSEHFRPECFSAFGNR

KNLKHNAVPTVF

>020017503.1|Castor canadensis

MPKSCAARQCCNRYSNRRKQLTFHRFPFSRPELLKEWVLNLGRANFKPKQHTVICSEHFRPECFSAFGNR

KNLKHNAVPTVF

>020747401.1|Odocoileus virginianustexanus

MPKSCAARQCCNRYSNRRKQLTFHRFPFSRPELLKEWVLNIGRGDFEPKQHTVICSEHFRPECFSAFGNR

KNLKHNAVPTVF

>020950927.1|Sus scrofa

MSVFCHKCLQQPSELPLPSWSALSPAWGAARGPSWFPFSRPELLKEWVLNIGRGDFEPKQHTVICSEHFR

PECFSAFGNRKNLKHNAVPTEF

>005079342.1|Mesocricetus auratus

MPKSCAARQCCNRYSSRRKQLTFHRFPFSRPELLKKWVLSIGRANFKPKQHTVICSEHFRPECFSAFGNR

KNLKHNAVPTVF

>004863757.2|Heterocephalus glaber

MRAQSCASAFGVPGLQVCGCHPRAEVTGVFRRPGAGTPSVTSGGAHTYSAAAERAGVRLGRARTGKPGAD

SARVGAGTPLTRPPVTGLTPVPSPASPPPALARAGPAPALARRPRAAIFVRGRPKMPKSCAARQCCNRYS

SRRKQLTFHRFPFSRPELLKEWVLNIGRADFKPKQHTVICSEHFRPECFSAFGNRKNLKHNAVPTVF

>021499943.1|Meriones unguiculatus

MPKSCAARQCCNRYSSRRKQLTFHRFPFSRPELLKEWLLNIGRANFKPKQHTVICSEHFRPECFSAFGNR

KNLKQNAVPTVF

>021539071.1|Neomonachus schauinslandi

MWCFQNSTQLCRANLSLANSFMSPLPPRFPFSRPELLKEWVLNIGRGNFKPKQHTVICSEHFRPECFSAF

GNRKNLKQNAVPTVF

>849390.2|Canis lupus familiaris

MPKSCAARQCCNRYSSRRKQLTFHRFPFSRPELLKEWVLNIGRANFKPKQHTVICSEHFRPECFSAFGNR

KNLKQNAVPTVF

>023113816.1|Felis catus

MPKSCAARQCCNRYSSRRKQLTFHRFPFSRPELLKEWVLNIGRGNFKPKQHTVICSEHFRPECFSAFGNR

KNLKQNAVPTVF

>003413535.1|Loxodonta africana

MPKSCAARQCCNRYSNRRKQLTFHRFPFSRPELLKEWVLNIGRGNFKPKQHTVICSEHFRPECFSAFGNR

KNLKHNAVPTVF

>003471278.2|Cavia porcellus

MPKSCAARQCCNRYSSLRKQLTFHRFPFSRPELLKEWVLNIGRAGFRPKQHTVICSEHFRPECFSAFGNR

TNLKHNAVPTVF

>011366761.1|Pteropus vampyrus

MPKSCAARQCCNRYSNRRKQLTFHRFPFSRPELLKEWVLNMGRANFKPKQHTVICSEHFRPECFSAFGNR

KNLKHNAVPTLF

>023491638.1|Equus caballus

MPKSCAARQCCNRYSNRRKQLTFHRFPFSRPELLKEWVLNIGRGNFKPKQHTVICSEHFRPECFSAFGNR

KNLKHNAVPTEF

>023555119.1|Octodon degus

MPKSCAARQCCNRYSSRRKQLTFHRFPFSRPELLKAWVLNMGRAGFKPKQHTVICSEHFRPECFSAFGNR

KNLKHNAVPTVF

>023603049.1|Myotis lucifugus

MRATEQRQARQLLPLAPPLQPPAGGRAGGRRRRPAASPGLPAALRRAGQPGPRGGAEPEERRVPPPRSEG

AEEPWFPFSRPELLKEWVLNIGRGNFQPKQHTVICSEHFRPECFSAFGNRKNLKHNAVPTEF

>024410465.1|Desmodus rotundus

MPKSCAARQCCNRYSNRRKQLTFHRFPFSRPELLKEWVLNIGRGNFEPKQHTVICSEHFRPECFSAFGNR

KNLKHNAVPTVF

>024611897.1|Neophocaena asiaeorientalisasiaeorientalis

MLTSLRFPFSRPELLKEWVLNIGRGDFEPKQHTVICSEHFRPECFSAFGNRKNLKHNAVPTVF

>024904393.1|Pteropus alecto

MGRANFKPKQHTVICSEHFRPECFSAFGNRKNLKHNAVPTLF

>006044304.1|Bubalus bubalis

MPKSCAARQCCNRYSNRRKQLTFHRFPFSRPELLKEWVLNIGRGDFEPKQHTVICSEHFRPECFSAFGNR

KNLKHNAVPTVF

>025283079.1|Canis lupus dingo

MPKSCAARQCCNRYSSRRKQLTFHRFPFSRPELLKEWVLNIGRANFKPKQHTVICSEHFRPECFSAFGNR

KNLKQNAVPTVF

>025712422.1|Callorhinus ursinus

MPKSCAARQCCNRYSSRRKQLTFHRCSLERSRFPFSRPELLKEWVLNIGRGNFKPKQHTVICSEHFRPEC

FSAFGNRKNLKQNAVPTVFAF

>025768379.1|Puma concolor

MPFPFSRPELLKEWVLNIGRGNFKPKQHTVICSEHFRPECFSAFGNRKNLKQNAVPTVF

>025860892.1|Vulpes vulpes

MPKSCAARQCCNRYSSRRKQLTFHRFPFSRPELLKEWVLNIGRANFKPKQHTVICSEHFRPECFSAFGNR

KNLKQNAVPTVF

>026241843.1|Urocitellus parryii

MPKSCAARQCCNRYSNRRKQLTFHRFPFSRPELLKEWVLNIGRAGFKPKQHTVICSEHFRPECFSAFGNR

KNLKHNAVPTVF

>026375727.1|Ursus arctoshorribilis

MPKSCAARQCCNRYSSRRKQLTFHRFPFSRPELLKEWVLNIGRGNFKPKQHTVICSEHFRPECFSAFGNR

KNLKLNAVPTVF

>026638047.1|Microtus ochrogaster

MPKSCAARQCCNRYSSRRKQLTFHREAPFKAPASSSVCASGAVHRGLAACAAPGAPHPPRGLSTQFQCFY

HGKMHTSVGSPQTTLHWHPLPVSSHLPTGRARFPFSRPELLKEWVLNIGRVNFKPKQHTVICSEHFRPEC

FSAFGNRKNLKHNAVPTVF

>026916349.1|Acinonyx jubatus

MPKSCAARQCCNRYSSRRKQLTFHRFPFSRPELLKEWVLNIGRGNFKPKQHTVICSEHFRPECFSTFGNR

KNLKQNAVPTVF

>026966132.1|Lagenorhynchus obliquidens

MPKSCAARQCCNRYSNRRKQLTFHRFPFSRPELLKEWVLNIGRGDFEPKQHTVICSEHFRPECFSAFGNR

KNLKHNAVPTVF

>003510263.1|Cricetulus griseus

MPKSCAARQCCNRYSSRRKQLTFHRFPFSRPELLKEWVLNIGRANFKPKQHTVICSEHFRPECFSAFGNR

KNLKHNAVPTVF

>027463634.1|Zalophus californianus

MPKSCAARQCCNRYSSRRKQLTFHRCSLERSRFPFSRPELLKEWVLNIGRGNFKPKQHTVICSEHFRPEC

FSAFGNRKNLKQNAVPTVF

>006145454.1|Tupaia chinensis

MPKSCAARQCCNRYSNRRKQLTFHRFPFSRPELLKEWVLNIGRGNFKPKQHTVICSEHFRPECFSAFGNR

KNLKHNAVPTVF

>012042770.2|Ovis aries

MPKSCAARQCCNRYSNRRKQLTFHRFPFSRPELLKEWVLNIGRGDFEPKQHTVICSEHFRPECFSAFGNR

KNLKHNAVPTVF

>027964274.1|Eumetopias jubatus

MPKSCAARQCCNRYSSRRKQLTFHRCSLERSRFPFSRPELLKEWVLNIGRGNFKPKQHTVICSEHFRPEC

FSAFGNRKNLKQNAVPTVF

>007166348.1|Balaenoptera acutorostratascammoni

MEAHLAAYLADVARGAACLGSLATFTVSGCRRQPRRKLPKAPGTLKGCVQPRESLAPPGCAPRFPFSRPE

LLKEWVLNIGRGDFEPKQHTVICSEHFRPECFSAFGNRKNLKHNAVPTVF

>028016747.1|Eptesicus fuscus

MPKSCAARQCCNRYSSRRKQLTFHRFPFSRPELLKEWVLNIGRGNFEPKQHTVICSEHFRPECFSAFGNR

KNLKHNAVPTEF

>028367361.1|Phyllostomus discolor

MPKSCAARQCCNRYSNRRKQLTFHRFPFSRPELLKEWVLNIGRGDFEPKQHTVICSEHFRPECFSAFGNR

KNLKHNAVPTVF

>028613759.1|Grammomys surdaster

MPKSCAARQCCNRYSSRRKQLTFHRFPFSRPELLRKWVLNIGRADFKPKQHTVICSEHFRPECFSAFGNR

KNLKHNAVPTVF

>028738933.1|Peromyscus leucopus

MPKSCAAKQCCNRYSSRRKQLTFHRFPFSRPELLKEWVLNIGRANFKPKQHTVICSEHFRPECFSAFGNR

KNLKHNAVPTVF

>029086368.1|Monodon monoceros

MPKSCAARQCCNRYSNRRKQLTFHRFPFSRPELLKEWVLNIGRGDFEPKQHTVICSEHFRPECFSAFGNR

KNLKHNAVPTVF

>021016511.1|Mus caroli

MPKSCAARQCCNRYSSLRKQLTFHRFPFSRPELLREWVLNIGRADFKPKQHTVICSEHFRPECFSAFGNR

KNLKHNAVPTVF

>021056597.1|Mus pahari

MPKSCAARQCCNRYSSRRKQLTFHRFPLSRPELLREWVLNMGRADFKPKQHTVICSEHFRPECFSAFGNR

KNLKHNAVPTVF

>008820657.1|Nannospalax galili

MPKSCAARQCCNRYSSRRKQLTFHRFPFSRPELLKEWVLNIGRIGFKPKQHTVICSEHFRPECFSAFGNR

KNLKHNAVPTEF

>029801786.1|Suricata suricatta

MPKSCAARQCCNRYSSRRKQLTFHRFPFSRPELLKEWVLNMGRGDFEPKQHTVICSEHFRPECFSAFGNR

KNLKQNAVPTVF

>030180367.1|Lynx canadensis

MWFNRVRLRDFPSCLLKYLVGFFRVQKSTGPTSALQVPSCLPPLPPRFPFSRPELLKEWVLNIGRGNFKP

KQHTVICSEHFRPECFSAFGNRKNLKQNAVPTVF

>022453973.1|Delphinapterus leucas

MPKSCAARQCCNRYSNRRKQLTFHRFPFSRPELLKEWVLNIGRGDFEPKQHTVICSEHFRPECFSAFGNR

KNLKHNAVPTVF

>012860781.1|Echinops telfairi

MPKSCAARQCCNRYSNRRKQLTFHRFPFSRPELLKEWVLNIGRGNFKPKQHTVICSEHFRPECFSAFGNR

KNLKHNAVPTVF

>030735768.1|Globicephala melas

MPKSCAARQCCNRYSNRRKQLTFHRFPFSRPELLKEWVLNIGRGDFEPKQHTVICSEHFRPECFSAFGNR

KNLKHNAVPTVF

>030876249.1|Leptonychotesweddellis

MWCFQNSTQLCRANVSLANSFMSPLPPRFPFSRPELLKEWVLNIGRGNFKPKQHTVICSEHFRPECFSAF

GNRKNLKQNAVPTVF

>031235454.1|Mastomys coucha

MPKSCAARQCCNRYSSRRKQLTFHRFPFSRPELLREWVLNIGRADFKPKQHTVICSEHFRPECFSAFGNR

KNLKHNAVPTVF

>031319822.1|Camelus dromedarius

MPKSCAARQCCNRYSNRRKQLTFHRFPFSRPELLKEWVLNIGRGNFEPKQHTVICSEHFRPECFSAFGNR

KNLKYNAVPTVF

>031539439.1|Vicugna pacos

MVEPLESVPPRPEFRAWPSKWVPAREISRSVSLVGDVGALWGKSHAPVSSMSTLNPQGTYLTQAYGPATL

YVGTAPPFPFSRPELLKEWVLNIGRGNFEPKQHTVICSEHFRPECFSAFGNRKNLKHNAVPTVF

>032157962.1|Mustela erminea

MPKSCAARQCCNRYSSRRKQLTFHRFPFSRPELLKEWVLNIGRGNFKPKQHTVICSEHFRPECFSAFGNR

KNLKQNAVPTVF

>032250554.1|Phoca vitulina

MPKSCAARQCCNRYSSRRKQLTFHRCSLERSRFPFSRPELLKEWVLNIGRGNFKPKQHTVICSEHFRPEC

FSAFGNRKNLKQNAVPTVF

>032350985.1|Camelus ferus

MPKSCAARQCCNRYSNRRKQLTFHRFPFSRPELLKEWVLNIGRGNFEPKQHTVICSEHFRPECFSAFGNR

KNLKHNAVPTVF

>032719186.1|Lontra canadensis

MPKSCAARQCCNRYSSRRKQLTFHRFPFSRPELLKEWVLNIGRGNFKPKQHTVICSEHFRPECFSAFGNR

KNLKQNAVPTVF

>032749987.1|Rattus rattus

MPKSCAARQCCNRYSSRRKQLTFHRFPFSRPELLRKWVLNIGRADFKPKQHTVICSEHFRPECFSAFGNR

KNLKHNAVPTVF

>032969493.1|Rhinolophus ferrumequinum

MAIEQSQGRLHLPAVPPLTQPRQAAGPDSPRRARRALQTTPTGPAPAASPALWPCRAHRPCRAPQAPAAI

FVRGSRAWGLEMPKSCAARQCCNRYSNRRKQLTFHRFPFSRPELLKAWVLNIGRGNFQPKQHTVICSEHF

RPECFSAFGNRKNLKHNAVPTVF

>OWK08622.1|Cervus elaphushippelaphus

MPKSCAARQCCNRYSNRRKQLTFHRFPFSRPELLKEWVLNIGRGDFEPKHHTVICSEHFRPECFSAFGNR

KNLKHNAVPTVF

>RLQ75012.1|Cricetulus griseus

MPKSCAARQCCNRYSSRRKQLTFHRFPFSRPELLKEWVLNIGRANFKPKQHTVICSEHFRPECFSAFGNR

KNLKHNAVPTVF

>AAH92427.1|Homo sapiens

MPKSCAARQCCNRYSSRRKQLTFHRFPFSRPELLKEWVLNIGRGNFKPKQHTVICSEHFRPECFSAFGNR

KNLKHNAVPTVF
