## Additional file for "Synapomorphic variations in the THAP domains of the human THAP protein family and its homologs": THAP4.docx

>PRD32361.1|Trichonephila clavipes

MVVCAAVKCSNSSRIKQMRFFKFPSEETRRKVWMTNSGLLNEPGPYAKLCEIHFEENQFELHRQDGWRKL

KPNAIPTVF

>ACO10667.1|Caligus rogercresseyi

MPACSALNCKNRSGTPGISLFQFPMDTLLRNEWIQSMKRDRWMPTIHSRLCSKHFEENLLIETQHAKKPK

KLVPGAIPTLF

>ACO14867.1|Caligus clemensi

MVFKCSAPGCKMGYTSSVKDARVTFHRFPMRDPNMLDIWISAIPRDCTKWRPTASSRLCSRHFVEDDFVL

KSRDTSRDVGKTLKFRRLADDAVPSKW

>ACO12088.1|epeophtheirus salmonis

MVGCCAPGCSRRTEHGVRFFGYPKDPKRRKQWEIMTNRPLWTPTEHSKLCQYHFEDDMFKRKEPNWRLKP

NGIPTLF

>025072747.1|Alligator sinensis

MLFFLFPHTSFNLHPSRREILIAISLGYWFPLKDSKRLIQWLKAVQRDNWTPTKYSFLCSEHFTKDSFSK

RLEDQHRLLKPTAVPTIF

>029472197.1|Rhinatrema bivittatum

MVICCAALNCSNRQGKVKKGAISFHRFPLKDSKRVTQWLKAVQRNNWTPTKYSFLCSEHFSKDCFFKRFE

SQHRSLKPTAVPSIF

>030071844.1|Microcaecilia unicolor

MVICCAAFNCSNRQGKGKKGAISFHRFPLKDSKRVTQWLKAVQRNNWTPTKYSFLCSEHFSKDCFSKRFE

DQHRFLKPTAVPSIF

>001072821.1|Xenopus tropicalis

MVICCAAPSCTNRQGKGKKGDVSFHRFPLKDHVRLSLWTDALQRDNWTPGPYSFLCSDHFSPESFVRRMP

DQTPLLTSNAVPSLF

>005295826.1|Chrysemys pictabellii

MVICCAAVNCSNRQGKVHRGAVSFHRFPLKDSKRLIQWLKAVQRDNWTPTKYSFLCSEHFTKDSFSKRLE

DQHRLLKPTAVPTIF

>006110680.1|Pelodiscus sinensis

MLLIFRFPLKDSKRLIQWLKAVQRDNWTPTKYSFLCSEHFTKESFSKRLEDQHRLLKPTAVPTIF

>007067438.1|Chelonia mydas

MVICCAAVNCSNRQGKVHRGAVSFYTFPLKDSKRLIQWLKAVQRDNWTPTKYSFLCSEHFTKDSFSKRLE

DQHRLLKPTAVPTIF

>026516571.1|Terrapene carolinatriunguis

MVICCAAVNCSNRQGKVHRGAVSFHRFPLKDSKRLIQWLKAVQRDNWTPTKYSFLCSEHFTKDSFSKRLE

DQHRLLKPTAVPTIF

>030432480.1|Gopherus evgoodei

MVICCAAVNCSNRQGKVHRGAVSFHRFPLKDSKRLIQWLKAVQRDNWTPTKYSFLCSEHFTKDSFSKRLE

DQHRLLKPTAVPTIF

>032651767.1|Chelonoidis abingdonii

MVICCAAVNCSNRQGKVHRGAVSFHRFPLKDSKRLIQWLKAVQRDNWTPTKYSFLCSEHFTKDSFSKRLE

DQHRLLKPTAVPTIF

>JAI11077.1|Crotalus adamanteus

MVICCAALNCSNRQGKAARGRAAVSFHRFPLKDSKRLIQWLKAVQRDNWIPTKYSFLCSEHFTKDSFSKR

LEDQHRLLKPTAIPTIF

>020641985.1|Pogona vitticeps

MVICCAALNCSNRQGKAPRGRAAVSFHRFPLKDSKRLIQWLKAVQRDNWTPTKYSFLCSEHFTKDSFSKR

LEDQHRLLKPTAVPTIF

>007440507.1|Python bivittatus

MVICCAALNCSNRQGKDPRGRAAVSFHRFPLKDSKRLIQWLKAVQRDNWIPTKYSFLCSEHFTKDSFSKR

LEDQHRLLKPTAIPTIF

>026550378.1|Notechis scutatus

MVICCAALNCSNRQGKGPRDRVAVSFHRFPLKDSKRLIQWLKAVQRDNWIPTKYSFLCSEHFTKDSFSKR

LEDQHRFLKPTAIPSIF

>026557228.1|Pseudonaja textilis

MVICCAALNCSNRQGKGPRDRVAVSFHRFPLKDSKRLIQWLKAVQRDNWIPTKYSFLCSEHFTKDSFSKR

LEDQHRFLKPTAIPSIF

>028586795.1|Podarcis muralis

MVICCAALNCSNRQGKAPRGRAAVSFHRFPLKDSKRLIQWLKAVQRDNWIPTKYSFLCSEHFTKDSFSKR

HEDQHRLLKPTAVPTIF

>015679977.1|Protobothrops mucrosquamatus

MVICCAALNCSNRQGKAARGRAAVSFHRFPLKDSKRLIQWLKAVQRDNWIPTKYSFLCSEHFTKDSFSKR

LEDQHRLLKPTAIPTIF

>032081003.1|Thamnophis elegans

MVICCAALNCSNRQGKGPRGRVAVSFHRFPLKDSKRLIQWLKAVQRDNWIPTKYSFLCSEHFTKDSFSKR

LEDQHRLLKPTAIPTIF

>033006070.1|Lacerta agilis

MVICCAALNCSNRQGKAPRGRAAVSFHRFPLKDSKRLIQWLKAVQRDNWIPTKYSFLCSEHFTKDSFSKR

HEDQHRLLKPTAVPTIF

>ACQ59049.1|Anoplopoma fimbria

MPDSCCAVGCTNRRGNKPGLCFYRIPSEREHPERRALWICAIKRGKGKQWQPSKYTRICSEHFVRGAKSD

IPISPDWVPSVF

>ACM08898.1|Salmo salar

MPVWCSVPYCANYKQAQKQGVIFHSLPTGDIPRCRKWLAAIKNNIYNVNTPVAKYQNIRVCSQHFRPEDY

LRDYQSELMGKPNRRQLKRDAIPSVF

>023678053.1|Paramormyrops kingsleyae

MVISCAAVNCTNRQGKVDKTEVSFHRFPIKNASRLAKWKRAVRRENWMPNKYSFLCSRHFTPDSFLPGCE

DQHRQLKAGAVPSIF

>028650660.1|Erpetoichthyscalabaricus

MVISCAAVNCTNRQGKVDKASVSFHRFPMKNAERLTKWKEAVRRESWTPNKYSFLCSNHFTPDCFESRHD

DQHRQLKATAVPTIF

>001139961.1|Salmo salar

MPVWCSVPYCANYKQAQKQGVIFHSLPTGDIPRCRKWLAAIKNNIYNVNTPVAKYQNIRVCSQHFRPEDY

LRDYQSELMGKPNRRQLKRDAIPSVF

>029883690.1|Aquila chrysaetoschrysaetos

MVICCAAANCSNRQGKAHRGAVSFHRFPLKDSKRLIQWLKAVQRDNWTPTKYSFLCSEHFTKDSFSKRLE

DQHRLLKPTAVPTIF

>OWK51124.1|Lonchura striatadomestica

MLCRTETPSPPARGAGVTRLRHLAVRRRGGAGAGAGAGAGAGPAGRGRRRRSEMVICCAAANCSNRQGKA

RHGAVSFHRFPLKDSKRLIQWLKAVQRDNWTPTKYSFLCSEHFTKDSFFKRLEDHHRLLKPTAVPTIF

>015132590.2|Gallus gallus

MLSYGTLRRSAERRRVGGGKAGPWEAERRRRASGPAAPPMGAARREYAPRNSGGRARPSCCTEAAGPRPS

APGKAAPDRLGQAGGGPSRSSGGGRPAARKRFPLKDSKRLIQWLKAVQRDNWTPTKYSFLCSEHFTKDSF

SRRLEDQHRLLKPTAVPTIF

>025892136.1|Nothoprocta perdicaria

MVICCAAANCSNRQGKALRGAVSFHRFPLKDSKRLIQWLKAVQRDNWTPTKYSFLCSEHFTKDSFSKRLE

DQHRLLKPTAVPTIF

>025934293.1|Apteryx rowi

MVICCAAVNCSNRQGKAHRGTVSFHRFPLKDSKRLIQWLKAVQRDNWTPTKYSFLCSEHFTKDSFSKRLE

DQHRLLKPTAVPTIF

>025966446.1|Dromaius novaehollandiae

MVICCAAVNCSNRQGKAHRGTVSFHRFPLKDSKRLIQWLKAVQRDNWTPTKYSFLCSEHFTKDSFSKRLE

DQHRLLKPTAVPTIF

>027320651.1|Anas platyrhynchos

MVICCAAANCSNRQGKAHRGAVSFHRFPLKDSKRLIQWLKAVQRDNWTPTKYSFLCSEHFTKDSFSKRLE

DQHRLLKPTAVPTIF

>027559752.1|Neopelma chrysocephalum

MVICCAAANCSNRQGKARRGAVSFHRFPLKDSKRLIQWLKAVQRDNWTPTKYSFLCSEHFTKDSFSKRLE

DQHRLLKPTAVPTIF

>027517673.1|Corapipo altera

MVICCAAANCSNRQGKARRGAVSFHRFPLKDSKRLIQWLKAVQRDNWTPTKYSFLCSEHFTKDSFSKRLE

DQHRLLKPTAVPTIF

>027580703.1|Pipra filicauda

MVICCAAANCSNRQGKARRGAVSFHRFPLKDSKRLIQWLKAVQRDNWTPTKYSFLCSEHFTKDSFSKRLE

DQHRLLKPTAVPTIF

>027761925.1|Empidonax traillii

MVICCAAANCSNRQGKARRGAVSFHRFPLKDSKRLIQWLKAVQRDNWTPTKYSFLCSEHFTKDSFSKRLE

DQHRLLKPTAVPTIF

>017938639.1|Manacus vitellinus

MSRFFFRFPLKDSKRLIQWLKAVQRDNWTPTKYSFLCSEHFTKDSFSKRLEDQHRLLKPTAVPTIF

>030083238.1|Serinus canaria

MVICCAAANCSNRQGKARHGAVSFHRFPLKDSKRLIQWLKAVQRDNWTPTKYSFLCSEHFTKDSFFKRLE

DHHRLLKPTAVPTIF

>030312078.1|Calypte anna

MVICCAAANCSNRQGKAHRGAVSFHRFPLKDSKRLIQWLKAVQRDNWTPTKYSFLCSEHFTKDSFSKRLE

DQHRLLKPTAVPTIF

>030810655.1|Camarhynchus parvulus

MVICCAAANCSNRQGKARHGAVSFHRFPLKDSKRLIQWLKAVQRDNWTPTKYSFLCSEHFTKDSFFKRLE

DHHRLLKPTAVPTIF

>021401104.1|Lonchura striatadomestica

MVICCAAANCSNRQGKARHGAVSFHRFPLKDSKRLIQWLKAVQRDNWTPTKYSFLCSEHFTKDSFFKRLE

DHHRLLKPTAVPTIF

>031452365.1|Phasianus colchicus

MVICCAAANCSNRQGKALRGAVSFHRFPLKDSKRLIQWLKAVQRDNWTPTKYSFLCSEHFTKDSFSRRLE

DQHRLLKPTAVPTIF

>031974860.1|Corvus moneduloides

MVICCAAANCSNRQGKACRGAVSFHRFPLKDSKRLIQWLKAVQRDNWTPTKYSFLCSEHFTKDSFSKRLE

DQHRLLKPTAVPTIF

>032049473.1|Aythya fuligula

MVICCAAANCSNRQGKAHRGAVSFHRFPLKDSKRLIQWLKAVQRDNWTPTKYSFLCSEHFTKDSFSKRLE

DQHRLLKPTAVPTIF

>032553760.1|Chiroxiphia lanceolata

MVICCAAANCSNRQGKARRGAVSFHRFPLKDSKRLIQWLKAVQRDNWTPTKYSFLCSEHFTKDSFSKRLE

DQHRLLKPTAVPTIF

>030136130.2|Taeniopygia guttata

MVICCAAANCSNRQGKARHGAVSFHRFPLKDSKRLIQWLKAVQRDNWTPTKYSFLCSEHFTKDSFFKRLE

DHHRLLKPTAVPTIF

>032924784.1|Catharus ustulatus

MGAAIMLYGAGRAPPLVPRESNPLHHARPTRRRLSRLGCHGRTGSSGGGHPAAPTRFPLKDSKRLIQWLK

AVQRDNWTPTKYSFLCSEHFTKDSFFKRLEDRHRSLKPTAVPTIF

>030351978.1|Strigops habroptila

MVICCAAANCSNRQGKAHRGAVSFHRFPLKDSKRLIQWLKAVQRDNWTPTKYSFLCSEHFTKDSFSKRLE

DQHRLLKPTAVPTIF

>015493202.1|Parus major

MVICCAAANCFNRQGKARRGAVSFHRFPLKDSKRLIQWLKAVQRDNWTPTKYSFLCSEHFTKDSFSKRLE

DKHRLLKPTAVPTIF

>020844672.1|Phascolarctos cinereus

MVICCAAVNCSNRQGKGEKRAVSFHRFPLKDSKRLIQWLKAVQRDNWTPTKYSFLCSEHFTKDSFSKRLE

DQHRLLKPTAVPSIF

>027731840.1|Vombatus ursinus

MVICCAAVNCSNRQGKGEKRAVSFHRFPLKDSKRLIQWLKAVQRDNWTPTKYSFLCSEHFTKDSFSKRLE

DQHRLLKPTAVPSIF

>028925586.1|Ornithorhynchus anatinus

MVICCAAANCSNRQGKGEKRAVSFHRFPLKDSKRLIQWLKAVQRDNWTPTKYSFLCSEHFTKDSFSKRLE

DQHRLLKPTAVPSIF

>031813314.1|Sarcophilus harrisii

MVICCAAANCSNRQGKGEKRAVSFHRFPLKDSKRLIQWLKAVQRENWTPTKYSFLCSEHFTKDSFSKRLE

DQHRLLKPTAVPSIF

>PNI12327.1|Pan troglodytes

MVICCAAVNCSNRQGKGEKRAVSFHRFPLKDSKRLIQWLKAVQRDNWTPTKYSFLCSEHFTKDSFSKRLE

DQHRLLKPTAVPSIF

>PNJ10518.1|Pongo abelii

MVICCAAVNCSNRQGKGEKRAVSFHRFPLKDSKRLIQWLKAVQRDNWTPTKYSFLCSEHFTKDSFSKRLE

DQHRLLKPTAVPSIF

>AAH81882.1|Rattus norvegicus

MVICCAAVNCSNRQGKGEKRAVSFHRFPLKDSKRLIQWLKAVQRDNWTPTKYSFLCSEHFTKDSFSKRLE

DQHRLLKPTAVPSIF

>AAI10169.1|Bos taurus

MVICCAAANCSNRQGKGEKRAVSFHRFPLKDSKRLMQWLKAVQRDNWTPTKYSFLCSEHFTKDSFSKRLE

DQHRLLKPTAVPSIF

>AAH63758.1|Mus musculus

MVICCAAVNCSNRQGKGEKRAVSFHRFPLKDSKRLIQWLKAVQRDNWTPTKYSFLCSEHFTKDSFSKRLE

DQHRLLKPTAVPSIF

>AFE78526.1|Macaca mulatta

MVICCAAVNCSNRQGKGEKRAVSFHRFPLKDSKRLIQWLKAVQRDNWTPTKYSFLCSEHFTKDSFSKRLE

DQHRLLKPTAVPSIF

>012611102.1|Microcebus murinus

MVICCAAVNCSNRQGKGEKRAVSFHRFPLKDSKRLIQWLKAVQRDNWTPTKYSFLCSEHFTKDSFSKRLE

DQHRLLKPTAVPSIF

>020768859.1|Odocoileus virginianustexanus

MQWLKAVQRDNWTPTKYSFLCSEHFTKDSFSKRLEDQHRLLKPTAVPSIF

>005077247.1|Mesocricetus auratus

MVICCAAVNCSNRQGKGEKRAVSFHRFPLKDSKRLIQWLKAVQRDNWTPTKYSFLCSEHFTKDSFSKRLE

DQHRLLKPTAVPSIF

>004868550.1|Heterocephalus glaber

MVICCAAVNCSNRQGKGEKRAVSFHRFPLKDSKRLLQWLKAVQRDNWTPTKYSFLCSEHFTKDSFSKRLE

DQHRLLKPTAVPSIF

>021487753.1|Meriones unguiculatus

MVICCAAVNCSNRQGKGEKRAVSFHRFPLKDSKRLIQWLKAVQRDNWTPTKYSFLCSEHFTKDSFSKRLE

DQHRLLKPTAVPSIF

>021534700.1|Neomonachus schauinslandi

MPPYWAAGPERPRPPPTPRAAGCAPPGIGEPGRAEARPGGRAAGRPGVRRHLAGIYYSLGGVTFILYLPY

HLFPLKDSKRLIQWLKAVQRDNWTPTKYSFLCSEHFTKDSFSKRLEDQHRLLKPTAVPSIF

>021575325.1|Ictidomys tridecemlineatus

MKREHFTKDSFSKRLEDQHRLLKPTAVPSIF

>006935827.1|Felis catus

MQVVPRGLTRAATRSASSSRVPAVAPRGGPAALRFPLKDSKRLIQWLKAVQRDNWTPTKYSFLCSEHFTK

DSFSKRLEDQHRLLKPTAVPSIF

>003474603.1|Cavia porcellus

MVICCAAVNCSNRQGKGEKRAVSFHRFPLKDSKRLIQWLKAVQRDNWTPTKYSFLCSEHFTKDSFSKRLE

DQHRLLKPTAVPSIF

>023376857.1|Pteropus vampyrus

MVICCAAANCSNRQGKGEKRAVSFHRFPLKDSKRLMQWLKAVQRDNWTPTKYSFLCSEHFTKDSFSKRLE

DQHRLLKPTAVPSIF

>023498567.1|Equus caballus

MVICCAAANCSNRQGKGEKRAVSFHRFPLKDSKRLIQWLKAVQRDNWTPTKYSFLCSEHFTKDSFSKRLE

DQHRLLKPTAVPSIF

>004637431.1|Octodon degus

MVICCAAVNCSNRQGKGEKRAVSFHRFPLKDSKRLIQWLKAVQRDNWTPTKYSFLCSEHFTKDSFSKRLE

DQHRLLKPTAVPSIF

>014322698.1|Myotis lucifugus

MVICCAAANCSNRQGKGEKRAVSFHRFPLKDSKRLIQWLKAVQRDNWTPTKYSFLCSEHFTKDSFSKRLE

DQHRLLKPTAVPSIF

>015443351.1|Pteropus alecto

MWGATGMRPVARGLTRLPGAVLLEGSAQPRGKEGSCAPEVRFPLKDSKRLMQWLKAVQRDNWTPTKYSFL

CSEHFTKDSFSKRLEDQHRLLKPTAVPSIF

>024842577.1|Bos taurus

MVICCAAANCSNRQGKGEKRAVSFHRFPLKDSKRLMQWLKAVQRDNWTPTKYSFLCSEHFTKDSFSKRLE

DQHRLLKPTAVPSIF

>006077177.1|Bubalus bubalis

MQWLKAVQRDNWTPTKYSFLCSEHFTKDSFSKRLEDQHRLLKPTAVPSIF

>025259921.1|Theropithecus gelada

MVICCAAVNCSNRQGKGEKRAVSFHRFPLKDSKRLIQWLKAVQRDNWTPTKYSFLCSEHFTKDSFSKRLE

DQHRLLKPTAVPSIF

>025319557.1|Canis lupus dingo

MVICCAAANCSNRQGKGEKRAVSFHRFPLKDSKRLIQWLKAVQRDNWTPTKYSFLCSEHFTKDSFSKRLE

DQHRLLKPTAVPSIF

>025744104.1|Callorhinus ursinus

MVICCAAANCSNRQGKGEKRAVSFHRFPLKDSKRLIQWLKAVQRDNWTPTKYSFLCSEHFTKDSFSKRLE

DQHRLLKPTAVPSIF

>026237773.1|Urocitellus parryii

MVICCAAVNCSNRQGKGEKRAVSFHRFPLKDSKRLIQWLKAVQRENWTPTKYSFLCSEHFTKDSFSKRLE

DQHRLLKPTAVPSIF

>005361425.1|Microtus ochrogaster

MVICCAAVNCSNRQGKGEKRAVSFHRFPLKDSKRLIQWLKAVQRDNWTPTKYSFLCSEHFTKDSFSKRLE

DQHRLLKPTAVPSIF

>026901186.1|Acinonyx jubatus

MQVVPRGLTRAATRSASSRVPAVAPRGGPAALRFPLKDSKRLIQWLKAVQRDNWTPTKYSFLCSEHFTKD

SFSKRLEDQHRLLKPTAVPSIF

>026971870.1|Lagenorhynchus obliquidens

MPARGAPAHPRRREPRAAPRQVPGIQGAPRTGPGGRAAGGRAPPSCRTAGAAGLAPTSRSRVAARPPLGA

VPPGARRGETRAGARRRRGGPRPEMVICCAAANCSNRQGKGEKRAVSFRRFPLKDSKRLIQWLKAVQRDN

WTPTKYSFLCSEHFTKDSFSKRLEDQHRLLKPTAVPSIF

>027253477.1|Cricetulus griseus

MVICCAAVNCSNRQGKGEKRAVSFHRFPLKDSKRLIQWLKAVQRDNWTPTKYSFLCSEHFTKDSFSKRLE

DQHRLLKPTAVPSIF

>027444293.1|Zalophus californianus

MVICCAAANCSNRQGKGEKRAVSFHRFPLKDSKRLIQWLKAVQRDNWTPTKYSFLCSEHFTKDSFSKRLE

DQHRLLKPTAVPSIF

>014438294.1|Tupaia chinensis

MVICCAAVNCSNRQGKGEKRAVSFHRFPLKDSKRLLQWLKAVQRDNWTPTKYSFLCSEHFTKDSFSKRLE

DQHRLLKPTAVPSIF

>027807486.1|Marmota flaviventris

MVICCAAVNCSNRQGKGEKRAVSFHRFPLKDSKRLIQWLKAVQRENWTPTKYSFLCSEHFTKDSFSKRLE

DQHRLLKPTAVPSIF

>027822592.1|Ovis aries

MVICCAAANCSNRQGKGEKRAVSFHRFPLKDSKRLMQWLKAVQRDNWTPTKYSFLCSEHFTKDSFSKRLE

DQHRLLKPTAVPSIF

>027961052.1|Eumetopias jubatus

MVICCAAANCSNRQGKGEKRAVSFHRFPLKDSKRLIQWLKAVQRDNWTPTKYSFLCSEHFTKDSFSKRLE

DQHRLLKPTAVPSIF

>027996657.1|Eptesicus fuscus

MVICCAAANCSNRQGKGEKRAVSFHRFPLKDSKRLIQWLKAVQRDNWTPTKYSFLCSEHFTKDSFSKRLE

DQHRLLKPTAVPSIF

>028339373.1|Physeter catodon

MVICCAAANCSNRQGKGEKRAVSFHRFPLKDSKRLIQWLKAVQRDNWTPTKYSFLCSEHFTKDSFSKRLE

GQHRLLKPTAVPSIF

>028611198.1|Grammomys surdaster

MVICCAAVNCSNRQGKGEKRAVSFHRFPLKDSKRLIQWLKAVQRDNWTPTKYSFLCSEHFTKDSFSKRLE

DQHRLLKPTAVPSIF

>028746468.1|Peromyscusleucopus

MVICCAAVNCSNRQGKGEKRAVSFHRFPLKDSKRLIQWLKAVQRDNWTPTKYSFLCSEHFTKDSFSKRLE

DQHRLLKPTAVPSIF

>029081966.1|Monodon monoceros

MVICCAAANCSNRQGKGEKRAVSFHRFPLKDSKRLIQWLKAVQRDNWTPTKYSFLCSEHFTKDSFSKRLE

DQHRLLKPTAVPSIF

>021015871.1|Mus caroli

MVICCAAVNCSNRQGKGEKRAVSFHRFPLKDSKRLIQWLKAVQRDNWTPTKYSFLCSEHFTKDSFSKRLE

DQHRLLKPTAVPSIF

>021053884.1|Mus pahari

MVICCAAVNCSNRQGKGEKRAVSFHRFPLKDSKRLIQWLKAVQRDNWTPTKYSFLCSEHFTKDSFSKRLE

DQHRLLKPTAVPSIF

>008827381.1|Nannospalax galili

MVICCAAVNCSNRQGKGEKRAVSFHRFPLKDSKRLIQWLKAVQRDNWTPTKYSFLCSEHFTKDSFSKRLE

DQHRLLKPTAVPSIF

>030183759.1|Lynx canadensis

MVICCAAANCSNRQGKGEKRAVSFHRFPLKDSKRLIQWLKAVQRDNWTPTKYSFLCSEHFTKDSFSKRLE

DQHRLLKPTAVPSIF

>022423929.1|Delphinapterus leucas

MVICCAAANCSNRQGKGEKRAVSFHRFPLKDSKRLIQWLKAVQRDNWTPTKYSFLCSEHFTKDSFSKRLE

DQHRLLKPTAVPSIF

>030659848.1|Nomascus leucogenys

MVICCAAVNCSNRQGKGEKRAVSFHRFPLKDSKRLIQWLKAVQRDNWTPTKYSFLCSEHFTKDSFSKRLE

DQHRLLKPTAVPSIF

>030771801.1|Rhinopithecus roxellana

MVICCAAVNCSNRQGKGEKRAVSFHRFPLKDSKRLIQWLKAVQRDNWTPTKYSFLCSEHFTKDSFSKRLE

DQHRLLKPTAVPSIF

>030863034.1|Gorilla gorilla gorilla

MSPYGLAPPGRLPRPSARLPTDAARGLRRPAGPKTPEPAAGEDAGGGGGGGGGGGGPAAGACASILPYGA

DAAGRPTSRAVSPPRSLPRGGPRAGGRLGPGPGCAAGPRPAMVICCAAVNCSNRQGKGEKRAVSFHRFPL

KDSKRLIQWLKAVQRDNWTPTKYSFLCSEHFTKDSFSKRLEDQHRLLKPTAVPSIF

>031224362.1|Mastomys coucha

MVICCAAVNCSNRQGKGEKRAVSFHRFPLKDSKRLIQWLKAVQRDNWTPTKYSFLCSEHFTKDSFSKRLE

DQHRLLKPTAVPSIF

>003908251.1|Papio anubis

MVICCAAVNCSNRQGKGEKRAVSFHRFPLKDSKRLIQWLKAVQRDNWTPTKYSFLCSEHFTKDSFSKRLE

DQHRLLKPTAVPSIF

>023050849.1|Piliocolobus tephrosceles

MVICCAAVNCSNRQGKGEKRAVSFHRFPLKDSKRLIQWLKAVQRDNWTPTKYSFLCSEHFTKDSFSKRLE

DQHRLLKPTAVPSIF

>032125444.1|Sapajus apella

MVICCAAVNCSNRQGKGEKRAVSFHRFPLKDSKRLIQWLKAVQRDNWTPTKYSFLCSEHFTKDSFSKRLE

DQHRLLKPTAVPSIF

>032211369.1|Mustela erminea

MPLYRAAGPGRPRPPPTPRAAGCAQPGTGEPRHSESRPGGRTAGRQGVRRHLAVREPRRPAPTSRSRVAA

RPPLGAVPLAARRGEARAGAWRRRGGLRPEMVICCAAANCSNRQGKGEKRAVSFHRFPLKDSKRLIQWLK

AVQRDNWTPTKYSFLCSEHFTKDSFSKRLEDQHRLLKPTAVPSIF

>032285623.1|Phoca vitulina

MVICCAAANCSNRQGKGEKRAVSFHRFPLKDSKRLIQWLKAVQRDNWTPTKYSFLCSEHFTKDSFSKRLE

DQHRLLKPTAVPSIF

>032336675.1|Camelus ferus

MVICCAAANCSNRQGKGEKRAVSFHRFPLKDSKRLIQWLKAVQRDNWTPTKYSFLCSEHFTKDSFSKRLE

DQHRLLKPTAVPSIF

>032492733.1|Phocoena sinus

MVICCAAANCSNRQGKGEKRAVSFHRFPLKDSKRLIQWLKAVQRDNWTPTKYSFLCSEHFTKDSFSKRLE

DQHRLLKPTAVPSIF

>032617093.1|Hylobates moloch

MVICCAAVNCSNRQGKGEKRAVSFHRFPLKDSKRLIQWLKAVQRDNWTPTKYSFLCSEHFTKDSFSKRLE

DQHRLLKPTAVPSIF

>032713613.1|Lontra canadensis

MVICCAAANCSNRQGKGEKRAVSFHRFPLKDSKRLIQWLKAVQRDNWTPTKYSFLCSEHFTKDSFSKRLE

DQHRLLKPTAVPSIF

>032757575.1|Rattusrattus

MVICCAAVNCSNRQGKGEKRAVSFHRFPLKDSKRLIQWLKAVQRDNWTPTKYSFLCSEHFTKDSFSKRLE

DQHRLLKPTAVPSIF

>032968726.1|Rhinolophus ferrumequinum

MSPYGAAGPGAPAHPRRREPRAAPRRVPETSGEPRADGRRPGGGRRHLAVRRRGGRLPRPAAVYLRASPR

GGPAPGSARRGEGRSLAASRRPRTEMVICCAAANCSNRQGKGEKRAVSFHRFPLKDSKRLIQWLKAVQRD

NWTPTKYSFLCSEHFTKDSFSKRLEDQHRLLKPTAVPSIF

>033085352.1|Trachypithecus francoisi

MSPYGLAPPERLLHPPARLPADAARGRLRPAGPEAARTRAWLGRGGGAGAGPRWASAPPSCRTARTPRAG

PRPEPCRRRAPSLGAVRGRAGGRRGPGPGGAAGPRPAMVICCAAVNCSNRQGKGEKRAVSFHRFPLKDSK

RLIQWLKAVQRDNWTPTKYSFLCSEHFTKDSFSKRLEDQHRLLKPTAVPSIF

>033279366.1|Orcinus orca

MPARGAPAHPRRREPRAAPRQVPGIQGAPRTGPGGRAAGGRAPPSCRTAGAAGLAPTSRSRVAARPPLGA

VPPGARRGETRAGARRRRGGPRPEMVICCAAANCSNRQGKGEKRAVSFHRFPLKDSKRLIQWLKAVQRDN

WTPTKYSFLCSEHFTKDSFSKRLEDQHRLLKPTAVPSIF

>RLQ75781.1|Cricetulus griseus

MVICCAAVNCSNRQGKGEKRAVSFHRFPLKDSKRLIQWLKAVQRDNWTPTKYSFLCSEHFTKDSFSKRLE

DQHRLLKPTAVPSIF

>AAH69235.1| Homo sapiens

MVICCAAVNCSNRQGKGEKRAVSFHRFPLKDSKRLIQWLKAVQRDNWTPTKYSFLCSEHFTKDSFSKRLEDQHRLLKPTAVPSIF
