## Additional file for "Synapomorphic variations in the THAP domains of the human THAP protein family and its homologs": THAP5.docx

>PFX18871.1|Stylophora pistillata

MPTACIAIGCSNVSTLKRTRWVSFYRLPIEDKQLLQRWLVNLKRTTLPSRLDRSYVCSDHFEPECFGRDI

RAELMGKRPKPPLLKGSVPTLF

>JAB99866.1|Ceratitis capitata

MKCAVLNCKSTNLSRASDFSFFRFPSEPTLLEKWMKFCRRKNKFNPSTSVMCMNHFTPDDIENRQKFEMG

FSKRRLLKNGAVPTLY

>RXG50939.1|Armadillidium vulgare

MKMKRWKFYPCKFSKICSDHFTPECINPGVGFGKPTLKQNAIPTIF

>006016023.1|Alligator sinensis

MPRYCAASCCKNRGGQSARDQRKLSFYPFPLHDKERLEKWLRNMKRDAWTPSKHQLLCSDHFTPDSLDVR

WGIRYLKQTAVPTIF

>032894250.1|Amblyraja radiata

MKRESWLPSKHQCICCEHFTADSIEWHCGTRCLKSRAAPTIF

>005297957.1|Chrysemys pictabellii

MGSRPCASTSLYPCWGRDLHYALPRSSSQLPFLNPALLIAQGSLGLAAPLDRGSAWFPLHDKERLEKWLR

NMKRDTWTPSKHQLLCSDHFTPDSLDVRWGIRYLKHTAIPTIF

>006122567.1|Pelodiscus sinensis

MKRDTWTPSKHQLLCSDHFTPDSLDVRWGIRYLKHTAIPTIF

>007052720.2|Chelonia mydas

MPRYCAASCCKNRGGQSARDQRKLSFYPFPLHDKERLEKWLRNMKRDTWTPSKHQLLCSDHFTPDSLDVR

WGIRYLKHTAIPTIF

>024056746.1|Terrapene carolinatriunguis

MPRYCAASCCKNRGGQSARDQRKLSFYPFPLHDKERLEKWLRNMKRDTWTPSKHQLLCSDHFTPDSLDVR

WGIRYLKHTAIPTIF

>030404861.1|Gopherus evgoodei

MPRYCAASCCKNRGGQSARDQRKLSFYPFPLHDKERLEKWLRNMKRDTWTPSKHQLLCSDHFTPDSLDVR

WGIRYLKHTAIPTIF

>032649322.1|Chelonoidis abingdonii

MPRYCAASCCKNRGGQSARDQRKLSFYPFPLHDKERLEKWLRNMKRDTWTPSKHQLLCSDHFTPDSLDVR

WGIRYLKHTAIPTIF

>007425483.1|Python bivittatus

MPRYCAATCCRNRAGQSARDQRKLSFYPFPLHDKERLEKWLRNMKRDTWTPSKHQVLCSDHFTPDSLDVR

WGIRYLKTTAVPTIF

>026524291.1|Notechis scutatus

MPRYCAATCCRNRAGQSARDQRKLSFYPRFPLHDKERLEKWLRNMKRDTWTPSKHQVLCSDHFTSDSLDV

RWGIRYLKTTAVPTIF

>026566115.1|Pseudonaja textilis

MPRYCAATCCRNRAGQSARDQRKLSFYPRFPLHDKERLEKWLRNMKRDTWTPSKHQVLCSDHFTSDSLDV

RWGIRYLKTTAVPTIF

>015677932.1|Protobothrops mucrosquamatus

MSNHRRHRQWGPSGTGIAGRRGVFAMPRYCAATCCRNRAGQSARDQRKLSFYPFPLHDKERLEKWLRNMK

RDTWTPSKHQVLCSDHFTPDSLDVRWGIRYLKTTAVPTIF

>032070435.1|Thamnophis elegans

MPRYCAATCCRNRAGQSARDQRKLSFYPFPLHDKERLEKWLRNMKRDTWTPSKHQVLCSDHFTSDSLDVR

WGIRYLKTTAVPTIF

>001090306.1|Xenopus laevis

MTRYCAATRCKNRGGQAAVIQRKVSFYPFPLHDRGRLQEWLRNMKQYKLYPTKHQVLCSDHFTADSFNIR

WGIQYLKPNAIPTLF

>002932859.2|Xenopus tropicalis

MTRYCAATCCKNRGGQAASTKHKLSFYPFPLHDRGRLKEWLCNMKQNKWYPTKHQVLCSDHFTPDSFSMR

WGIRYLKPNAIPTIF

>029470853.1|Rhinatrema bivittatum

MATLVNGERNQLKEISTPPPPYVWGGTAASIYTAARSKMGNYHPVSCVLWSRRNTKWLIIKGLLHHPFTW

YMKRFPLHDTQRLEKWLRNMKRGSWVPSKHHYLCSDHFTPDSFDVRWGITYLKPNAIPTVF

>030071891.1|Microcaecilia unicolor

MTRYCAAFSCKNRDTGAAREERKPSFYPFPLHDMERLERWLRNMKRGSWIPTKHHYLCSDHFTPDSFDVR

WGIRYLKPNAIPTVF

>033814463.1|Geotrypetes seraphini

MTRYCAAFSCKNRDTGAAREERKPSFYPFPLHDMERLERWLQNMKRGSWVPTKHHYLCSDHFTPDSFDIR

WGIRYLKPNAVPTVF

>JAR00445.1|Fundulus heteroclitus

MTDCALTVPIERGQLAIKARTSRPSRRFSIYXXSCARVLVTRHQPQPHRHEYVDALSSGERRADRAAILG

QTKRKSDRIQVSSVSKLPELRPRMPDFCSACGCANKRALQTRSRGITFHKFPKDTTLRKKWELAVRRDGF

VATDRSMLCSEHFKDEDFDRTGQIVRLRLNVIPSIF

>020470057.1|Monopterus albus

MPRYCAVKTCRNRGGAVSRQDNKRISFYPFPLYDKPRLQKWVDNMKREEWTPSRHQYLCSEHFTEDCFDI

RWGIRYLKNTAIPTIF

>022078408.1|Acanthochromis polyacanthus

MPRYCAVKVCRNRGGTASRQDNKRISFYPFPLQDKPRLQKWVDNMKRAEWTPSRHQYLCSDHFTEDCFDI

RWGIRYLKNKAIPTIF

>SBR32238.1|Nothobranchius kuhntae

MPRYCAVKVCRNRGGTAWKTISNMPENKRISFYPFPLQDKQRLQKWVDNMRREEWTPSRHQYLCSEHFTE

DCFDIRWGIRYLKSTAIPTIF

>SBP36917.1|Iconisemion striatum

MPRYCSVKICRNRGGAAWKTINNMQENKRISFYPFPLQDKPRLQQWMDNMRREEWTPSRHQYLCSEHFTE

DCFDIRWGIRYLKNTAIPTIF

>SBS45651.1|Nothobranchius furzeri

MPRYCAVKVCRNRGGTAWKTISNMPENKRISFYPFPLQDKQRLQKWVDNMRREEWTPSRHQYLCNEHFTE

DCFDIRWGIRYLKSTAIPTIF

>SBR62550.1|Nothobranchius pienaari

MPRYCAVKVCRNRGGTAWKTISNMPENKRISFYPFPLQDKQRLQKWVDNMRREEWTPSRHQYLCNEHFTE

DCFDIRWGIRYLKSTAIPTIF

>SBP77876.1|Nothobranchius kadleci

MPRYCAVKVCRNRGGTAWKTISNMPENKRISFYPFPLQDKQRLQKWVDNMRREEWTPSRHQYLCNEHFTE

DCFDIRWGIRYLKSTAIPTIF

>SBQ63544.1|Nothobranchius korthausae

MPRYCAVKICRNRGGTAWKTISNMPENKRISFYPFPLQDKQRLQKWVDNMRREEWTPSRHQYLCSEHFTE

DCFDIRWGIRYLKSTAIPTIF

>SBQ89974.1|Nothobranchius kuhntae

MPRYCAVKVCRNRGGTAWKTISNMPENKRISFYPFPLQDKQRLQKWVDNMRREEWTPSRHQYLCSEHFTE

DCFDIRWGIRYLKSTAIPTIF

>022604746.1|Seriola dumerili

MPRYCAVKACRNRGGSASRQDNKRISFYPFPLQDKPRLQKWVDNMKREEWTPSRHQCLCSEHFTEDCFDI

RWGIRYLKNTAIPTIF

>023205597.1|Xiphophorus maculatus

MRLHRGDATILRRKALPKPWRDFIKTRQENKLLPDKTRLQKWVNNMRRGEWTPSRHQYLCSKHFTEDCFD

IRWGIRYLKNTAIPTLF

>023273671.1|Seriola lalandi dorsalis

MPRYCAVKACRNRGGSASRQDNKRISFYPFPLQDKPRLQKWVDNMKREEWTPSRHQCLCSEHFTEDCFDI

RWGIRYLKNTAIPTIF

>023666146.1|Paramormyrops kingsleyae

MPRYCAVKLCKNRGGVLSKDNKKISFYPFPLRDEARLQKWVDNMKREQWTPSRHQYLCSDHFTEDSFDLR

WGIRYLKNTAVPTIF

>004560915.1|Maylandia zebra

MPRYCAVRVCRNRGGTASRQDSKRISFYPFPLQDKSRLQKWVSNMKREQWTPSRHQYLCSEHFTADCFDI

RWGIRYLKNTAIPTIF

>017280302.1|Kryptolebias marmoratus

MPRYCAVRVCPNRGGTASRKDNKRISFYPFPLHDKPRLQKWVDNMKREDWTPSRHQYLCSEHFTEDCFDI

RWGIRYLKNTAIPTIF

>025755623.1|Oreochromis niloticus

MKREQWTPSRHQYLCSEHFTADCFDIRWGIRYLKNTAIPTIF

>026002725.1|Astatotilapia calliptera

MLYTSLPLRGPQTGYNHLRADMPRYCAVRVCRNRGGTASRQDSKRISFYPFPLQDKSRLQKWVSNMKREQ

WTPSRHQYLCSEHFTADCFDIRWGIRYLKNTAIPTIF

>019129979.2|Larimichthys crocea

MPRYCAVKVCRNRGGTASRQDNKRISFYPFPLQDKPRLQKWVDNMKREEWTPSRHQYLCSEHFTEDCFDI

RWGIRYLKNTAIPTVF

>028256606.1|Parambassis ranga

MPRYCAVKVCRNRGGTTSKQDNKRISFYPFPLQDKPRLQKWVDNMKRAEWTPSRHQYLCSEHFTEDCFDI

RWGIRYLKNTAIPTIF

>028427003.1|Perca flavescens

MPRYCAVKVCRNRGGTASRQDKRISFYPFPLQDKSRLQKWVDNMKREEWTPSRHQYLCSEHFKEDCFDIR

WGIRYLKNTAIPTLF

>028666940.1|Erpetoichthys calabaricus

MPRYCAADCCQNRGGVSSKDNRKISFYPFPLQDKERLKNWVMNMKRENWVPSKHQYLCSDHFTADSFDIR

WGIRYLKHTAVPTIF

>029018960.1|Betta splendens

MAVPLGPRSLLYLKTACVKAWMSGAAAAGPPLGFGVALHVGALQQAIRAGAGGEKHCQKNPRTTSSGVKL

VFWSSGKTGVGVQWNAVTLLPSVIQMPHQRVHKWLRCGVTDSQQSQGEPRLQYQTPPPLYTTSISNKSSG

VHLLLTTHSHASLSRHSTRFPLHDKPRLQKWVDNMKREDWTPSRHQYLCSEHFTEDCFDIRWGIRYLKNT

AVPTVF

>018585452.1|Scleropages formosus

MRSELLFIVSSFEMPRYCAVKLCKNRGGGLSKDNKKISFYPFPLRDEARLRKWVDNMERKEWSPSRHQYL

CSEHFTEDSFDLRWGIRYLKHTAVPTIF

>029317353.1|Cottoperca gobio

MPRYCAVKGCRNRGGTTSTQDNKRISFYPFPLQDKSRLQKWVDNMKREEWAPSRHQYLCSEHFTEDCFDI

RWGIRYLKNTAIPTTF

>|Echeneisnaucrates

MPRYCAMKGCRNRGGSSFGQEHKRISFYPFPLQDKPRLQKWVDNMKREEWTPSRHQCLCSEHFTEDCFDI

RWGIRYLKSTAIPTIF

>029901272.1|Myripristis murdjan

MPRYCAVKVCRNRGGTASKQEDKRISFYPFPLQDKPRLQKWVDNMKREEWTPSRHQYLCSEHFTDDCFDI

RWGIRYLKSTAIPTIF

>029969565.1|Salarias fasciatus

MPRYCAVKVCRNRGGAASRHDNRRISFYPFPLQDKARLQKWVDNMKREDWTPSRHQYLCSEHFTEDCFDV

RWGIRYLKNTSIPTLF

>030005211.1|Sphaeramia orbicularis

MPRYCAVKVCRNRGVHASRQESKRISFYPFPLQDKPRLQKWVDNMNREKWTPSRHQFLCSEHFTEDCFDI

RWGIRYLKNTAIPTIF

>030295642.1|Sparus aurata

MPRYCAVKVCRNRGGTATRHDNKRISFYPFPLQDQPRLQKWVDNMKREEWTPSRHQYLCSEHFTEDCFDI

RWGIRYLKTTAIPTIF

>030576195.1|Archocentrus centrarchus

MPRYCAVRVCRNRGGTASRQDKRISFYPFPLQDKPRLQEWVNNMKREEWTPSRHQYLCSEHFTEDCFDIR

WGIRYLKNTAIPTIF

>031140782.1|Sander lucioperca

MPRYCAVKVCRNRGGTASRQDKRISFYPFPLQDKSRLQKWVDNMKREKWTPSRHQYLCSEHFKEDCFDIR

WGIRYLKNTAIPTLF

>012685686.2|Clupea harengus

MPRYCAVKLCKNRGGVLSKDNKRISFYPFPLRDQARLQKWVDNMKRQEWTPSRHQYLCSEHFTEDSFDLR

WGIRYLKHTAIPTIF

>031606462.1|Oreochromis aureus

MPRYCAVRVCRNRGGTASRQDNKRISFYPFPLQDKSRLQKWVSNMKREQWTPSRHQYLCSEHFTADCFDI

RWGIRYLKNTAIPTIF

>032361337.1|Etheostoma spectabile

MPRYCAVKNCKNRGGTASRQDKRISFYPFPLQDKSRLQKWVRNMKREEWTPSRHQYLCSEHFKEDCFDIR

WGIRYLKNTAIPTLF

>032445660.1|Xiphophorus hellerii

MPRYCAEKLCRNRGGTSSKQDKKISFYPFPLQDKTRLQKWVNNMRREEWTPSRHQYLCSEHFTEDCFDIR

WGIRYLKNTAIPTLF

>026183270.1|Mastacembelus armatus

MPRYCAVKACRNRGRTAFKKANKRISFYPFPLHDEPRLQKWMGNIKQGEWTPSRHQYVCSEHFTEDCFDI

RWGIRYLKNTAIPTLF

>OPJ68763.1|Patagioenas fasciatamonilis

MPRYCAASCCKNRGGQTARDQRKLSFYPFPLHDKERLEKWLRNMKRDAWTPSKHQLLCSDHFTPDSLDVR

WGIRYLKHTAVPTIF

>019135469.2|Corvuscornixcornix

MPRYCAATHCKNRGGQSSRDQRKLSFYPHCVDQSPSQACGNLWTVVVTWLSLPRMLNYKQYFTILREARC

RFPLHDKERLEKWLRNMKRDSWTPSKHQLLCSDHFTPDSLDVRWGIRYLKHTAVPTIF

>005508832.1|Columba livia

MKRDAWTPSKHQLLCSDHFTPDSLDVRWGIRYLKHTAVPTIF

>021240617.1|Numida meleagris

MPRYCAASCCKNRGGQSARDQRKLSFYPFPLHDKERLEKWLRNMKRDAWTPSKHQLLCSDHFTPDSLDVR

WGIRYLKHTAVPTIF

>023786265.1|Cyanistes caeruleus

MPRYCAATHCKNRGGQNAKDQRKLSFYPFPLHDKERLEKWLRNMKRDSWTPSKHQLLCSDHFTPDSLDVR

WGIRYLKNTAVPTIF

>025894934.1|Nothoprocta perdicaria

MPRYCAASCCKNRGGQSARDRRKISFYPFPLHDKERLEKWLRNMKRDAWMPSKHQLLCSDHFTPDSLEVR

WGIRYLKHTAVPTIF

>025925766.1|Apteryx rowi

MPRYCAASCCKNRGGQSARDQRKLSFYPFPLHDKERLEKWLRNMKRDAWTPSKHQLLCSDHFTPDSLDVR

WGIRYLKHTAVPTIF

>025973541.1|Dromaius novaehollandiae

MPRYCAASRCKNRGGQSARDQRKLSFYPFPLHDKERLEKWLRNMKRDAWTPSKHQLLCSDHFTPDSLDVR

WGIRYLKHTAVPTIF

>005480558.1|Zonotrichia albicollis

MPRYCAATRCKNRGGQSAKDKRKLSFYPFPLHDKERLEKWLRNMKRDSWTPSKHQLLCSDHFTPDSLDVR

WGIRYLKSTAVPTIF

>026724116.1|Athene cunicularia

MPRYCAASCCKNRGGQSARDQRKLSFYPFPLHDKERLEKWLRNMKRDAWTPSKHQLLCSDHFTPDSLDVR

WGIRYLKHTAVPTIF

>027538051.1|Neopelma chrysocephalum

MPRYCAATCCKNRGGQSARDQRKLSFYPFPLHDKERLEKWLRNMKRDSWTPSKHQLLCSDHFTPDSLDVR

WGIRYLKHTAVPTIF

>027513784.1|Corapipo altera

MPRYCAATRCKNRGGQSARDQRKLSFYPFPLHDKERLEKWLRNMKRDSWTPSKHQLLCSDHFTPDSLDVR

WGIRYLKHTAVPTIF

>027575487.1|Pipra filicauda

MPRYCAATRCKNRGGQSARDQRKLSFYPFPLHDKERLEKWLRNMKRDSWTPSKHQLLCSDHFTPDSLDVR

WGIRYLKHTAVPTIF

>001006230.1|Gallus gallus

MPRYCAASYCKNRGGQSARDQRKLSFYPFPLHDKERLEKWLRNMKRDAWTPSKHQLLCSDHFTPDSLDVR

WGIRYLKHTAVPTIF

>005242159.1|Falco peregrinus

MKRDAWTPSKHQLLCSDHFTPDSLDVRWGIRYLKHTAVPTIF

>005444088.1|Falco cherrug

MKRDAWTPSKHQLLCSDHFTPDSLDVRWGIRYLKHTAVPTIF

>027749220.1|Empidonax traillii

MPRYCAATRCKNRGGQSARDQRKLSFYPFPLHDKERLEKWLRNMKRDSWIPSKHQLLCSDHFTPDSLDVR

WGIRYLKHTAVPTIF

>028943284.1|Antrostomus carolinensis

MKRDAWTPSKHQLLCSDHFTPDSLDVRWGIRYLKHTAVPTIF

>029869988.1|Aquila chrysaetoschrysaetos

MVPPRARAAPLRHAAVLRRVLLQEPRGPERQGPAQAELLPVSGRRGGFPLHDKERLEKWLRNMKRDAWTP

SKHQLLCSDHFTPDSLDVRWGIRYLKHTAVPTIF

>008927933.3|Manacus vitellinus

MPRYCAATRCKNRGGQSARDQRKLSFYPFPLHDKERLEKWLRNMKRDSWTPSKHQLLCSDHFTPDSLDVR

WGIRYLKHTAVPTIF

>030094335.1|Serinus canaria

MRYSYLRQRKTTITGVLYIFLFVFSTALKRFPLHDKERLEKWLRNMKRDSWTPSKHQLLCSDHFTPDSLD

VRWGIRYLKNTAVPTIF

>008494384.2|Calypte anna

MPRYCAASCCKNRGGQSAKDQRKLSFYPFPLHDKERLEKWLRNMKRDAWTPSKHQLLCSDHFTPDSLDVR

WGIRYLKHTAVPTIF

>030815938.1|Camarhynchus parvulus

MPRYCAATRCKNRGGQSAKDKRKLSFYPFPLHDKERLEKWLRNMKWDSWTPSKHQLLCSDHFTPDSLDVR

WGIRYLKNTAVPTIF

>005426394.1|Geospiza fortis

MKRDSWTPSKHQLLCSDHFTPDSLDVRWGIRYLKNTAVPTIF

>021400947.1|Lonchura striatadomestica

MPRYCAATHCKNRGGQSAKDQRKLSFYPFPLHDKERLEKWLRNMKRDSWTPSKHQLLCSDHFTPDSLDVR

WGIRYLKNTAVPTIF

>031445451.1|Phasianus colchicus

MPRYCAASCCKNRGGQSARDQRKLSFYPFPLHDKERLEKWLRNMKRDAWTPSKHQLLCSDHFTPDSLDVR

WGIRYLKHTAVPTIF

>010705265.1|Meleagris gallopavo

MKRDAWTPSKHQLLCSDHFTPDSLDVRWGIRYLKHTAVPTIF

>031963159.1|Corvus moneduloides

MPRYCAATHCKNRGGQSARDQRKLSFYPHCVDQSPSQACSNLWTVVVTWLSLPRMLNYKQYFTILREARC

RFPLHDKERLEKWLRNMKRDSWTPSKHQLLCSDHFTPDSLDVRWGIRYLKHTAVPTIF

>032060932.1|Aythya fuligula

MPRYCAAACCKNRGGQSARDRRKLSFYPFPLHDKERLEKWLRNMKRDAWTPSKHQLLCSDHFTPDSLDVR

WGIRYLKHTAVPTIF

>015709662.1|Coturnix japonica

MPRYCAASCCKNRGGQSARDQRKLSFYPFPLHDKERLEKWLRNMKRDAWTPSKHQLLCSDHFTPDSLDVR

WGIRYLKHTAVPTIF

>032543756.1|Chiroxiphia lanceolata

MPRYCAATRCKNRGGQSARDQRKLSFYPFPLHDKERLEKWLRNMKRDSWTPSKHQLLCSDHFTPDSLDVR

WGIRYLKHTAVPTIF

>002195322.3|Taeniopygia guttata

MPRYCAATRCKNRGGQSAKDQRKLSFYPFPLHDKERLEKWLRNMKRDSWTPSKHQLLCSDHFTPDSLDVR

WGIRYLKNTAVPTIF

>032855683.1|Tyto albaalba

MKRDAWMPSKHQLLCSDHFTPDSLDVRWGIRYLKHTAVPTIF

>032913993.1|Catharus ustulatus

MPRYCAATRCKNRGGQSAKDQRKLSFYPFPLHDKERLEKWLRNMKRDSWTPSKHQLLCSDHFTPDSLDVR

WGIRYLKNTAVPTIF

>030334789.1|Strigops habroptila

MPRYCAASCCKNRGGQSARDRSKLSFYPFPLHDKERLEKWLRNMKRDAWTPSKHQLLCSDHFTPDSLDVR

WGIRYLKHTAVPTIF

>015484127.1|Parus major

MPRYCAATRCKNRGGQNAKDQRKLSFYPFPLHDKERLEKWLRNMKRDSWTPSKHQLLCSDHFTPDSLDVR

WGIRYLKNTAVPTIF

>020840289.1|Phascolarctos cinereus

MPRYCAAFSCKNRRGRNNKDRKLSFYPFPLHDKERLEKWLRNMKRDTWVPSKYQFLCSDHFTPDSLDIRW

GIRYLKQTAVPTIF

>027694437.1|Vombatus ursinus

MPRYCAAFSCKNRRGRNNKDRKLSFYPFPLHDKERLEKWLRNMKRDTWVPSKYQFLCSDHFTPDSLDIRW

GIRYLKHTAVPTIF

>028929885.1|Ornithorhynchus anatinus

MPRYCAAFCCKNRRGRGGAERRLSFYPFPLHDKERLEKWLRNMKRDAWVPSKYQFLCSDHFTPDSLDVRW

GIRYLKQTAIPTIF

>003771561.1|Sarcophilus harrisii

MPRYCAAFSCKNRRGRNNKDRKLSFYPFPLHDKERLEKWLRNMKRDTWVPSKYQFLCSDHFTPDSLDIRW

GIRYLKQTAVPTIF

>020041899.1|Castor canadensis

MPRYCAAICCKNRRGRNNKDRKLSFYPFPLHDKERLEKWLKNMKRDSWVPSKYQFLCSDHFTPDSLDIRW

GIRYLKQTAIPTIF

>020745621.1|Odocoileus virginianustexanus

MVGIEADSSKENGMFPLHDKERLEKWLKNMKRDSWVPSKYQFLCSDHFTPDSLDIRWGIRYLKQTAIPTI

F

>003134826.2|Sus scrofa

MPRYCAAICCKNRRGRNNTDRKLSFYPFPLHDKERLEKWLKNMKRDSWVPSKYQFLCSDHFTPDSLDIRW

GIRYLKQTAIPTIF

>004839692.1|Heterocephalus glaber

MPRYCAAICCKNRRGRNNKGRKLSFYPFPLHDKERLEKWLKNMKRDSWVPSKYQFLCSDHFTPDSLDIRW

GIRYLKQTAIPTIF

>005335994.1|Ictidomys tridecemlineatus

MPRYCAAICCKNRRGPNNKDRKLSFYPFPLHDKERLEKWLKNMKRDSWVPSKYQFLCSDHFTPDSLDIRW

GIRYLKQTAIPTIF

>539521.2|Canis lupus familiaris

MPRYCAAICCKNRRGRNSKDRKLSFYPFPLHDKERLEKWLKNMKRDSWVPSKYQFLCSDHFTPDSLDIRW

GIRYLKQTAIPTIF

>003983009.1|Felis catus

MPRYCAAICCKNRRGRNSKDRKLSFYPFPLHDKERLEKWLKNMKRDSWVPSKYQFLCSDHFTPDSLDIRW

GIRYLKQTAIPTIF

>003407261.1|Loxodonta africana

MPRYCAAICCKNRRGRNNKDRKLSFYPFPLHDKERLEKWLKNMKRDSWVPSKYQFLCSDHFTPDSLDIRW

GIRYLKQTAVPTIF

>004472374.1|Dasypusnovemcinctus

MPRYCAAVYCKNRRGRNDKDRKLSFYPFPLHDKERLEKWLKNMKRDSWIPSKYQFLCSDHFTPDSLDIRW

GIRYLKQTAIPTIF

>003469923.1|Cavia porcellus

MPRYCAATCCKNRRGRSNKDRKLSFYPFPLHDKERLEKWLKNMKRDSWVPSKYQFLCSDHFTPDSLDIRW

GIRYLKQTAIPTIF

>011360804.1|Pteropus vampyrus

MPRYCAAICCKNRRGRNNKDRKLSFYPFPLHDKERLEKWLKNMKRDSWVPSKYQFLCSDHFTPDSLDVRW

GIRYLKQTAIPTIF

>023495298.1|Equus caballus

MARRQGLLGCELERLHTSPDLVSLNDVNQRLGTQQTCRVGRCTLFPLHDKERLEKWLKNMKRDSWIPSKY

QFLCSDHFTPDSLDVRWGIRYLKQTAVPTIF

>004641690.1|Octodon degus

MPRYCAATCCKNRRGRNNKDRKLSFYPFPLHDKERLEKWLKNMKRDSWVPSKYQFLCSDHFTPDSLDIRW

GIRYLKQTAIPTIF

>006085382.1|Myotis lucifugus

MPRYCAAICCKNRRGRNSKDRKLSFYPFPLHDKERLEKWLKNMKRDSWMPSKYQFLCSDHFTPDSLDIRW

GIRYLKQTAIPTIF

>024624646.1|Neophocaena asiaeorientalisasiaeorientalis

MPRYCAAICCKNRRGRNSKDRKLSFYPFPLHDKERLEKWLKNMKRDSWVPSKYQFLCSDHFTPDSLDIRW

GIRYLKQTAIPTIF

>005205517.1|Bos taurus

MPRYCAAICCKNRRGRNNKERKLSFYPFPLHDKERLEKWLKNMKRDSWVPSKYQFLCSDHFTPDSLDIRW

GIRYLKQTAIPTIF

>006912134.1|Pteropus alecto

MPRYCAAICCKNRRGRNNKDRKLSFYPFPLHDKERLEKWLKNMKRDSWVPSKYQFLCSDHFTPDSLDVRW

GIRYLKQTAIPTIF

>006050764.1|Bubalus bubalis

MPRYCAAICCKNRRGRNNKERKLSFYPFPLHDKERLEKWLKNMKRDSWVPSKYQFLCSDHFTPDSLDIRW

GIRYLKQTAIPTIF

>025320430.1|Canis lupus dingo

MPRYCAAICCKNRRGRNSKDRKLSFYPFPLHDKERLEKWLKNMKRDSWVPSKYQFLCSDHFTPDSLDIRW

GIRYLKQTAIPTIF

>025728946.1|Callorhinus ursinus

MPRYCAAICCKNRRGRNSKDRKLSFYPFPLHDKERLEKWLKNMKRDSWVPSKYQFLCSDHFTPDSLDIRW

GIRYLKQTAIPTIF

>025785638.1|Puma concolor

MPRYCAAICCKNRRGRNSKDRKLSFYPFPLHDKERLEKWLKNMKRDSWVPSKYQFLCSDHFTPDSLDIRW

GIRYLKQTAIPTIF

>025841602.1|Vulpes vulpes

MPRYCAAICCKNRRGRNSKDRKLSFYPFPLHDKERLEKWLKNMKRDSWVPSKYQFLCSDHFTPDSLDIRW

GIRYLKQTAIPTIF

>026264392.1|Urocitellus parryii

MPRYCAAICCKNRRGPNNKDRKLSFYPFPLHDKERLEKWLKNMKRDSWVPSKYQFLCSDHFTPDSLDIRW

GIRYLKQTAIPTIF

>026365153.1|Ursus arctoshorribilis

MPRYCAAICCKNRRGRSSKDRKLSFYPFPLHDKERLEKWLKNMKRDSWVPSKYQFLCSDHFTPDSLDIRW

GIRYLKQTAIPTIF

>014920502.1|Acinonyx jubatus

MPRYCAAICCKNRRGRNSKDRKLSFYPFPLHDKERLEKWLKNMKRDSWVPSKYQFLCSDHFTPDSLDIRW

GIRYLKQTAIPTIF

>027429330.1|Zalophus californianus

MPRYCAAICCKNRRGRNSKDRKLSFYPFPLHDKERLEKWLKNMKRDSWVPSKYQFLCSDHFTPDSLDIRW

GIRYLKQTAIPTIF

>006161191.1|Tupaia chinensis

MRGGSECLDLLPRLSEKGWSAPSSGPRVPDLTSAMPRYCAAICCKNRRGRNNKDRKLSFYPFPLHDKERL

EKWLKNMKRDSWIPSKYQFLCSDHFTPDSLDIRWGIRYLKQTAIPTIF

>027788485.1|Marmota flaviventris

MPRYCAAICCKNRRGPNNKDRKLSFYPFPLHDKERLEKWLKNMKRDSWVPSKYQFLCSDHFTPDSLDIRW

GIRYLKQTAIPTIF

>004007936.2|Ovis aries

MPRYCAAICCKNRRGRNNKERKLSFYPFPLHDKERLEKWLKNMKRDSWVPSKYQFLCSDHFTPDSLDIRW

GIRYLKQTAIPTIF

>027960655.1|Eumetopias jubatus

MPRYCAAICCKNRRGRNSKDRKLSFYPFPLHDKERLEKWLKNMKRDSWVPSKYQFLCSDHFTPDSLDIRW

GIRYLKQTAIPTIF

>007195148.1|Balaenoptera acutorostratascammoni

MPRYCAAICCKNRRGRNSKDRKLSFYPFPLHDKERLEKWLKNMKRDSWVPSKYQFLCSDHFTPDSLDIRW

GIRYLKQTAIPTIF

>008145446.1|Eptesicus fuscus

MPRYCAAICCKNRRGRNNKDRKLSFYPFPLHDKERLEKWLKNMKRDSWMPSKYQFLCSDHFTPDSLDIRW

GIRYLKQTAIPTIF

>007112115.1|Physeter catodon

MPRYCAAICCKNRRGRNSKDRKLSFYPFPLHDKERLEKWLKNMKRDSWVPSKYQFLCSDHFTPDSLDIRW

GIRYLKQTAIPTIF

>028381607.1|Phyllostomus discolor

MPRYCAAICCKNRRGRNNKDRRLSFYPFPLHDKERLEKWLKNMKRDSWMPSKYQFLCSDHFTPDSLDVRW

GIRYLKHTAVPTIF

>029089125.1|Monodon monoceros

MPRYCAAICCKNRRGRNSKDRKLSFYPFPLHDKERLEKWLKNMKRDSWVPSKYQFLCSDHFTPDSLDIRW

GIRYLKQTAIPTIF

>029418183.1|Nannospalax galili

MQSPFLISPASIKHVLRFPLHDKERLEKWLKNMKRDSWVPSKYQFLCSDHFTPDSLDIRWGIRYLKQTAI

PTIF

>029780731.1|Suricata suricatta

MPRYCAAICCKNRRGRNSKDRKLSFYPFPLHDKERLEKWLKNMKRDSWVPSKYQFLCSDHFTPDSLDIRW

GIRYLKQTAVPTIF

>030164099.1|Lynx canadensis

MPRYCAAICCKNRRGRNSKDRKLSFYPFPLHDKERLEKWLKNMKRDSWVPSKYQFLCSDHFTPDSLDIRW

GIRYLKQTAIPTIF

>022438842.1|Delphinapterus leucas

MPRYCAAICCKNRRGRNSKDRKLSFYPFPLHDKERLEKWLKNMKRDSWVPSKYQFLCSDHFTPDSLDIRW

GIRYLKQTAIPTIF

>004702641.1|Echinops telfairi

MPRYCAAVGCKNRRGRNNKDRKLSFYPFPLHDKERLEKWLRNMKRDSWVPSKYQFLCSDHFTPDSLDIRW

GIRYLKQTAVPTIF

>030713427.1|Globicephala melas

MPRYCAAICCKNRRGRNGKDRKLSFYPFPLHDKERLEKWLKNMKRDSWVPSKYQFLCSDHFTPDSLDIRW

GIRYLKQTAIPTIF

>006732416.1|Leptonychotes weddellii

MPRYCAAICCKNRRGRNSKDRKLSFYPFPLHDKERLEKWLKNMKRDSWVPSKYQFLCSDHFTPDSLDIRW

GIRYLKQTAIPTIF

>010973850.1|Camelus dromedarius

MPRYCAAICCKNRRGRNNKDRKLSFYPFPLHDKERLEKWLKNMKRDSWVPSKYQFLCSDHFTPDSLDIRW

GIRYLKQTAIPTIF

>006201965.1|Vicugna pacos

MPRYCAAICCKNRRGRNNRDRKLSFYPFPLHDKERLEKWLKNMKRDSWVPSKYQFLCSDHFTPDSLDIRW

GIRYLKQTAIPTIF

>032160397.1|Mustela erminea

MPRYCAAICCKNRRGRNSKDRKLSFYPFPLHDKERLEKWLKNMKRDSWVPSKYQFLCSDHFTPDSLDVRW

GIRYLKQTAIPTIF

>032257649.1|Phoca vitulina

MPRYCAAICCKNRRGRNSKDRKLSFYPFPLHDKERLEKWLKNMKRDSWVPSKYQFLCSDHFTPDSLDIRW

GIRYLKQTAIPTIF

>032944107.1|Rhinolophus ferrumequinum

MPRYCAAICCKNRRGRNNKDRKLSFYPFPLHDKERLEKWLKNMKRDSWVPSKYQFLCSDHFTPDSLDIRW

GIRYLKQNAIPTIF

>004269941.1|Orcinus orca

MPRYCAAICCKNRRGRNSKDRKLSFYPFPLHDKERLEKWLKNMKRDSWVPSKYQFLCSDHFTPDSLDIRW

GIRYLKQTAIPTIF

>010612335.1|Fukomys damarensis

MPRYCAAICCKNRRGRNTKDRKLSFYPFPLHDKERLEKWLKNMKRDSWVPSKYQFLCSDHFTPDSLDIRW

GVRYLKQTAIPTIF

>019780942.1|Tursiops truncatus

MPRYCAAICCKNRRGRNSKDRKLSFYPFPLHDKERLEKWLKNMKRDSWVPSKYQFLCSDHFTPDSLDIRW

GIRYLKQTAIPTIF

>JAV42924.1|Castor canadensis

MPRYCAAICCKNRRGRNNKDRKLSFYPFPLHDKERLEKWLKNMKRDSWVPSKYQFLCSDHFTPDSLDIRW

GIRYLKQTAIPTIF

>Q7Z6K1.2|Homo Sapiens

MPRYCAAICCKNRRGRNNKDRKLSFYPFPLHDKERLEKWLKNMKRDSWVPSKYQFLCSDHFTPDSLDIRW

GIRYLKQTAVPTIF
