## Additional file for "Synapomorphic variations in the THAP domains of the human THAP protein family and its homologs": THAP6.docx

>ADD24317.1|Lepeophtheirus salmonis (Arthropoda)

MPHCCAVYGCHTSKKKSNEHISFHRFPKEVNERKKWRIAICRKGWKPTDSSRICSIHFKDEDYIEGLQRR

VLTPGAIPSIF

>ACO15709.1|Caligus clemensi (Arthropoda)

MPSCSAPGCRNRSGTKGLTLFSFPKELSLRNEWVLRLKRGKWSPSQNSRLCCRHFEESQFRIIDPVIGRK

KLFRGSVPSVF

>KXJ06406.1|Exaiptasia pallid (Cnidaria)

MQRARYVYLSSSRVKLANDEWKENIALKNDLATYVRQNLHKYEILGLVEKRYPEYAWSIRTLTRRLQHFD

IKYVDYNIELESVRDAVKIEMEGPGKLLGYRALHKKIRDVHGLNVSRNLVYTVMQDVNPQGLQDRGGVGK

PKRPKRTKAFVSNITLVMPTHCCVPLCTHKGYKDTKTGEKISFFCFPKEQNLKRRWLHAIRREEKNDFKV

TTTTKVCSRHFETTDMKKSLAGIVTLKPGTVPSKF

>PFX19223.1|Stylophora pistillata (Cnidaria)

MNWANFFVETMPTHCCVPGCTKKGYLEEGKKISYFIFPTDKSLRKQWSHAIRRHEGKNFQISKSTKVCSR

HFKTEDFQRSLGGSKINLKPGVVPSVF

>KYO31967.1|Alligator mississippiensis (Chordata)

MVKSCSAAGCASRCLPRSKRRGLTFHVFPTDVETKRKWVLAVKRPGIWEPKKADVLCSRHFKKSDFDTRG

PNIKLKPGVVPSIF

>014372997.2|Alligator sinensis (Chordata)

MAPDHQGKAGPRDYPLSPHQLLSMAGTRLEPARGGGWQGVGRREGSKRGVGRSGSGGLTPPSPAPSFFHL

LIHLISFIKFPTDVETKRKWVLAVKRPGIWEPKKADVLCSRHFKKSDFDTRGPNIKLKPGVVPSIF

>023965889.1|Chrysemys pictabellii (Chordata)

MVKSCSAVGCASRCLPNSKLRGLTFHVFPTDEEAKRRWVLATKRLDVNSAGMWEPKKADVLCSRHFKKSD

FDTRGPNIRLKPGVIPSIF

>006125451.1|Pelodiscus sinensis

MVKCCSAVGCASRCLPNSKLRGLTFHVFPTNEEAKRRWVLAMKRLDVNSAAMWEPKKADVLCSRHFKKSD

FDTRGPNIRLKPGVIPSIF

>007062907.1|Chelonia mydas

MVKSCSAIGCASRCLPNSKLRGLTFHVFPTDEEAKRRWVLAMKRLDVNSAGMWEPKKADVLCSRHFKKSD

FDTRGPNIRLKPGVIPSIF

>026509036.1|Terrapene carolinatriunguis

MVKSCSAIGCASRCLPNSKLRGLTFHVFPTDEEAKRRWVLATKRLDVNSAGMWEPKKADVLCSRHFKKSD

FDTRGPNIRLKPGVIPSIF

>030421108.1|Gopherus evgoodei

MGNDRSLVCYSTSVKVSGKVRFPTDEEAKRRWVLATKRLDVNSAGMWEPKKTDVLCSRHFKKSDFDTRGP

NIRLKPGVIPSIF

>032618203.1|Chelonoidis abingdonii

MGNDLSLVCYSTSVMVSGKVRFPTDEEAKRRWVLATKRLDVNSAGMWEPKKADVLCSRHFKKSDFDTRGP

NIRLKPGVIPSIF

>JAR34649.1|Fundulus heteroclitus

MPVSCSAYGCKNRRTSLSRRRGITFHQFPKNSGLRKAWIAAFRRLDFKPSNKTVLCSDHFTEADFDRTGQ

TVRLREGVIPSIF

>SBQ84836.1|Nothobranchius korthausae

MPVFCAAYGCNNRRTAETRSRGITFHKFPRDTESKREWEIALRRKDFVATEFSKICSDHFEPEDFDRTGQ

TVRLRDGVIPSRF

>SBQ45373.1|Nothobranchius kadleci

MPVFCAAYGCNNRRTAETRSRGITFHKFPRDTESKREWEVALRRKDFVATEFSKICSDHFEPEDFDRTGQ

TVRLRDGVIPSRF

>SBP02869.1|Iconisemion striatum

MPDFCAAYGCSNRRCVESRQRGITFHSFPKDGERRRQWEVALRRENFVVNNRTMICSEHFKAEDFDRTGQ

TVRLKAGVGPTIF

>SBQ47568.1|Nothobranchius korthausae

MPVSCAAYGCKNRRTPLSKGQGVTFHQFPKDVTLRKAWISAVRRRDFKPSRKTVLCSSHFKEKDFDRTGQ

TVRLREAVIPSIF

>028929459.1|Ornithorhynchus anatinus

MVKCCSAVGCASRCLPNSKLKGLTFHVFPTDEKAKRKWVLAMKRLDVNAAGIWEPKKGDVLCSRHFEKAD

FDRSAPNVKLKPGVVPSVF

>023409115.1|Loxodonta africana

MVKCCSAIGCASRCLPNSKLKGLTFHVFPTDENMKRKWVLAMKRLDVNAAGIWEPKKGDVLCSRHFKKTD

FDRSAPNIKLKPGVIPSIF

>004472619.1|Dasypus novemcinctus

MVKCCSAIGCASRCLPNSKLKGLTFHVFPTDENIKRKWVLAMKRLDVNAAGIWEPKKGDVLCSRHFKKTD

FDRSAPNIKLKPGVVPSIF

>006142994.1|Tupaia chinensis

MVKCCSAIGCASRCLPNSKLKGLTFHVFPTDENVKRKWVLAMKRLDVNAAGIWEPKKGDVLCSRHFQKTD

FDRSAPNIKLKPGVIPSIF

>004703258.1|Echinops telfairi

MVKCCSAIGCASRCLPNSKLKGLTFHVFPTDENVKRKWVLAMKRLDVNAAAIWEPKKGDVLCSRHFKKTD

FDRSTPNIKLKPGVIPSIF

>023493730.1|Equus caballus

MVKCCSAIGCASRCLPNSKLKGLTFHVFPTDENVKRKWVLAMKRLDVNAAGIWEPKKGDVLCSRHFKKTD

FDRSAPNIKLKPGVIPSIF

>011358843.1|Pteropus vampyrus

MVKCCSAIGCASRCLPNSKLKGLTFHVFPTDENIKRKWVLAMKRLDVNAAGIWEPKKGDVLCSRHFKKTD

FDRSAPNIKLKPGVIPSIF

>014301417.1|Myotis lucifugus

MKRLDVNAAGFWEPKKGDVLCSRHFKKTDFDRSAPNIKLKPGVVPSIF

>024435232.1|Desmodus rotundus

MVKCCSAIGCASRCLPNSKLKGLTFHVFPTDENVKRKWVLAMKRLDVNAAGIWEPKKGDVLCSRHFKKTD

FDRSAPNIKLKPGVVPSIF

>006907711.1|Pteropus alecto

MVKCCSAIGCASRCLPNSKLKGLTFHVFPTDENIKRKWVLAMKRLDVNAAGIWEPKKGDVLCSRHFKKTD

FDRSAPNIKLKPGVIPSIF

>008141895.1|Eptesicus fuscus

MVKCCSAIGCASRCLPNSKLKGLTFHVFPTDENVKRKWVLAMKRLDVNAAGIWEPKKGDVLCSRHFKKTD

FDRSTPNIKLKPGVIPSIF

>028362200.1|Phyllostomus discolor

MVKCCSAIGCASRCLPNSKLKGLTFHVFPTDENVKRKWVLAMKRLDVNAAGIWEPKKGDVLCSRHFKKTD

FDRSAPNIKLKPGVIPSIF

>021558035.1|Neomonachus schauinslandi

MVKCCSAIGCASRCLPNSKLKGLTFHVFPTDENVKRKWVLAMKRLDVNAAGIWEPKKGDVLCSRHFKKTD

FDRSAPNIKLKPGVIPSIF

>026914208.1|Acinonyx jubatus

MKRLDVNAAGIWEPKKGDVLCSRHFKKTDFDRSAPNIKLKPGVIPSIF

>027454257.1|Zalophus californianus

MVKCCSAIGCASRCLPNSKLKGLTFHVFPTDENVKRKWVLAMKRLDVNAAGIWEPKKGDVLCSRHFKKTD

FDRSAPNIKLKPGVIPSIF

>027956557.1|Eumetopias jubatus

MVKCCSAIGCASRCLPNSKLKGLTFHVFPTDENVKRKWVLAMKRLDVNAAGIWEPKKGDVLCSRHFKKTD

FDRSAPNIKLKPGVIPSIF

>029800718.1|Suricata suricatta

MVKCCSAIGCASRCLPNSKLKGLTFHVFPTDENIKRKWVLAMKRLDVNAAGIWEPKKGDVLCSRHFKKTD

FDRSAPNIKLKPGVIPSIF

>030169535.1|Lynx canadensis

MVKCCSAIGCASRCLPNSKLKGLTFHVFPTDENVKRKWVLAMKRLDVNAAGIWEPKKGDVLCSRHFKKTD

FDRTAPNIKLKPGVIPSIF

>006740359.1|Leptonychotes weddellii

MVKCCSAIGCASRCLPNSKLKGLTFHVFPTDENVKRKWVLAMKRLDVNAAGIWEPKKGDVLCSRHFKKTD

FDRSAPNIKLKPGVIPSIF

>032190483.1|Mustela erminea

MVKCCSAIGCASRCLPNSKLKGLTFHVFPTDENVKRKWVLAMKRLDVNAAGIWEPKKGDVLCSRHFKKTD

FDRSAPNIKLKPGVIPSIF

>032256429.1|Phoca vitulina

MVKCCSAIGCASRCLPNSKLKGLTFHVFPTDENVKRKWVLAMKRLDVNAAGIWEPKKGDVLCSRHFKKTD

FDRSAPNIKLKPGVIPSIF

>032695776.1|Lontra canadensis

MVKCCSAIGCASRCLPNSKLKGLTFHVFPTDENVKRKWVLAMKRFDVNAAGIWEPKKGDVLCSRHFKKTD

FDRSAPNIKLKPGVIPSIF

>005639112.1|Canis lupus familiaris

MVKCCSAIGCASRCLPNSKLKGLTFHVFPTDENVKRKWVLAMKRLDVNAAGIWEPKKGDVLCSRHFKKTD

FDRSTPNIKLKPGVIPSIF

>006931122.1|Felis catus

MVKCCSAIGCASRCLPNSKLKGLTFHVFPTDENVKRKWVLAMKRLDVNAAGIWEPKKGDVLCSRHFKKTD

FDRSAPNIKLKPGVIPSIF

>025283810.1|Canis lupus dingo

MVKCCSAIGCASRCLPNSKLKGLTFHVFPTDENVKRKWVLAMKRLDVNAAGIWEPKKGDVLCSRHFKKTD

FDRSTPNIKLKPGVIPSIF

>025743020.1|Callorhinus ursinus

MVKCCSAIGCASRCLPNSKLKGLTFHVFPTDENVKRKWVLAMKRLDVNAAGIWEPKKGDVLCSRHFKKTD

FDRSAPNIKLKPGVIPSIF

>025778078.1|Puma concolor

MVKCCSAIGCASRCLPNSKLKGLTFHVFPTDENVKRKWVLAMKRLDVNAAGIWEPKKGDVLCSRHFKKTD

FDRSAPNIKLKPGVIPSIF

>014935363.1|Acinonyx jubatus

MVKCCSAIGCASRCLPNSKLKGLTFHVFPTDENVKRKWVLAMKRLDVNAAGIWEPKKGDVLCSRHFKKTD

FDRSAPNIKLKPGVIPSIF

>026365741.1|Ursus arctoshorribilis

MVKCCSAIGCASRCLPNSKLKGLTFHVFPTDENVKRKWVLAMKRLDVNAAGIWEPKKGDVLCSRHFKKTD

FDRSAPNIKLKPGVIPSIF

>025848767.1|Vulpes vulpes

MVKCCSAIGCASRCLPNSKLKGLTFHVFPTDENVKRKWVLAMKRLDVNAAGIWEPKKGDVLCSRHFKKTD

FDRSTPNIKLKPGVIPSIF

>JAN97499.1|Heterocephalus glaber

MVKCCSAIGCASRCLPNSKLKGLTFHAFPTDENVKRKWVLAMKRLDVNSAGMWEPKKGDVLCSRHFKKTD

FDRSAPNLKLKPGVIPSIF

>001100679.1|Rattus norvegicus

MVKCCSAIGCASRCLPNSKLKGLTFHVFPTDENIKRKWVLAMKRLDVNTAGIWEPKKGDVLCSRHFKKTD

FDRSTPNTKLKPGAIPSVF

>020038478.1|Castor canadensis

MVKCCSAIGCASRCLPNSKLKGLTFHVFPTDENIKRKWVLAMKRLDVNAAGIWEPKKGDVLCSRHFKKTD

FDRSAPNIKLKPGVIPSIF

>005068141.1|Mesocricetus auratus

MVKCCSAIGCASRCLPNSKLKGLTFHVFPTDENIKRKWVLAMKRLDVNAAGIWEPKKGDVLCSRHFKKTD

FDRSTPNTKLKPGVIPSVF

>021587317.1|Ictidomys tridecemlineatus

MVKCCSAIGCASRCLPNSKLKGLTFHVFPTDENVKRKWVLAMKRLDVNAAGIWEPKKGDVLCSRHFKKTD

FDRSAPNIKLKPGVIPSIF

>023568756.1|Octodon degus

MVKCCSAIGCASRCLPNSKLRGLTFHAFPTDENVKRKWVLAMKRLDVNTAGIWEPKKGDVLCSRHFKKTD

FDRSAPNLKLKPGVIPSIF

>026264251.1|Urocitellus parryii

MVKCCSAIGCASRCLPNSKLKGLTFHVFPTDENVKRKWVLAMKRLDVNAAGIWEPKKGDVLCSRHFKKTD

FDRSAPNIKLKPGXIPSIF

>005359547.1|Microtus ochrogaster

MVKCCSAIGCASRCLPNSKLKGLTFHVFPTDENIKRKWVLAMKRLDVNAAGIWEPKKGDVLCSRHFKKTD

FDRSTPNTKLKPGVVPSVF

>RLQ79101.1|Cricetulus griseus

MVKCCSAIGCASRCLPNSKLKGLTFHVFPTDENIKRKWVLAMKRLDVNTASIWEPKKGDVLCSRHFRKTD

FDRSTPNTKLKPGVIPSVF

>027795553.1|Marmota flaviventris

MVKCCSAIGCASRCLPNSKLKGLTFHVFPTDENVKRKWVLAMKRLDVNAAGIWEPKKGDVLCSRHFKKTD

FDRSAPNIKLKPGVIPSIF

>028608442.1|Grammomys surdaster

MVKCCSAIGCASRCLPNSKLKGLTFHVFPTDENIKRKWVLAMKRLDVNTAGIWEPKKGDVLCSRHFKKTD

FDRSTPNTKLKPGAIPSVF

>028737735.1|Peromyscus leucopus

MVKCCSAIGCASRCLPNSKLKGLTFHVFPTDENIKRKWVLAMKRLDVNAAGIWEPKKGDVLCSRHFKKTD

FDRSAPNTKLKPGVIPSVF

>021018160.1|Mus caroli

MVKGCSAIGCASRCLPNSKLKGLTFHVFPTDENIKRKWVLAMKRLDVNTASIWKPKKGDVLCSRHFKKTD

FDRSTPNTKLKAGAIPSIF

>029401112.1|Mus pahari

MANSTPAGVSVQQTCGRENLARENMREIKKPAELGSQTSRFPTDENIKRKWVLAMKRLDVNTASIWEPKK

GDVLCSRHFKKTDFDRSTPNTKLKAGAIPSVF

>017656233.1|Nannospalax galili

MPQWNSQIDGGENTAVNTITTWRNTFLLCVAKIMVKCCSAIGCASRCLPNSKLKGLTFHVFPTDENIKRK

WVLAMKRLDVNAADIWEPKKGDVLCSRHFKKTDFDRSTPNIKLKPGVIPSVF

>031246838.1|Mastomys coucha

MVKCCSAIGCASRCLPNSKLKGLTFHVFPTDENIKRKWVLAMKRLDVNTAGIWEPKKGDVLCSRHFKKTD

FDRSTPNTKLKPGAIPSVF

>032771937.1|Rattus rattus]

MVKCCSAIGCASRCLPNSKLKGLTFHVFPTDENIKRKWVLAMKRLDVNTAGIWEPKKGDVLCSRHFKKTD

FDRSTPNTKLKPGAIPSVF

>020759786.1|Odocoileus virginianustexanus

MVKCCSAVGCASRCLPNSKLKGLTFHVFPTDENIKRKWVLAVKRRDVNAAGIWEPKKGDVLCSRHFKKTD

FDRSAPNIKLKPGVIPSIF

>005666813.2|Sus scrofa

MVKCCSAIGCASRCLPNSKLKGLTFHVFPTDEKVKRKWVLAMKRLDVNAAGMWEPKKGDVLCSRHFKKTD

FDRTTPNIKLKPGVIPSIF

>024609853.1|Neophocaena asiaeorientalis

MVKCCSAIGCASRCLPNSKLKGLTFHVFPTDENVKRKWVLAMKRLDVNAVDIWEPKKGDVLCSRHFKKTD

FDRSTPNIKLKPGVIPSIF

>005208173.1|Bos taurus

MVKCCSAVGCASRCLPNSKLKGLTFHVFPTDENIKRKWVLAMKRLDVNAAGIWEPKKGDVLCSRHFKKTD

FDRSTPNIKLKPGVVPSIF

>006059713.1|Bubalus bubalis

MVKCCSAVGCASRCLPNSKLKGLTFHVFPTDENIKRKWVLAMKRLDVNAAGIWEPKKGDVLCSRHFKKTD

FDRSTPNIKLKPGVVPSIF

>026943515.1|Lagenorhynchus obliquidens

MVKCCSAVGCASRCLPNSKLKGLTFHVFPTDENVKRKWVLAMKRLDVNAVDIWEPKKGDVLCSRHFKKTD

FDRSTPNIKLKPGVIPSIF

>004009964.2|Ovis aries

MVKCCSAVGCASRCLPNSKLKGLTFHVFPTDENIKRKWVLAMKRLDVNAAGIWEPKKGDVLCSRHFKKTD

FDRSAPNIKLKPGVVPSIF

>007165608.1|Balaenoptera acutorostratascammoni

MVKCCSAVGCASRCLPNSKLKGLTFHVFPTDENVKRKWVLAMKRLDVNAVDTWEPKKGDVLCSRHFKKTD

FDRSAPNIKLKPGVIPSIF

>023971170.1|Physeter catodon

MVKCCSAIGCASRCLPNSKLKGLTFHVFPTDENVKRKWVLAMKRLDVNAVDIWEPKKGDVLCSRHFKKTD

FDRSAPNIKLKPGVIPSIF

>029091085.1|Monodon monoceros

MVKCCSAIGCASRCLPNSKLKGLTFHVFPTDENVKRKWVLAMKRLDVNAVDIWEPKKGDVLCSRHFKKTD

FDRSTPNIKLKPGVIPSIF

>022455088.1|Delphinapterus leucas

MVKCCSAIGCASRCLPNSKLKGLTFHVFPTDENVKRKWVLAMKRLDVNAVDIWEPKKGDVLCSRHFKKTD

FDRSTPNIKLKPGVIPSIF

>030733733.1|Globicephala melas

MVKCCSAVGCASRCLPNSKLKGLTFHVFPTDENVKRKWVLAMKRLDVNAVDIWEPKKGDVLCSRHFKKTD

FDRSTPNIKLKPGVIPSIF

>010981252.1|Camelus dromedarius

MVKCCSAIGCASRCLPNSKLKGLTFHVFPTDEKVKRKWVLAMKRLDVNAAGIWEPKKGDVLCSRHFKKTD

FDRSAPNIKLKPGVIPSIF

>031545088.1|Vicugna pacos

MENRKRYSYWPPDLKRTSADLQNSELCRFPTDEKVKRKWVLAMKRLDVNAAGIWEPKKGDVLCSRHFKKT

DFDRSAPNIKLKPGVIPSIF

>032313840.1|Camelus ferus

MENRKRYGYWPPDLKRTSADLQNSELCRFPTDEKVKRKWVLAMKRLDVNAAGIWEPKKGDVLCSRHFKKT

DFDRSAPNIKLKPGVIPSIF

>032488336.1|Phocoena sinus

MVKCCSAIGCASRCLPNSKLKGLTFHVFPTDENVKRKWVLAMKRLDVNAVDIWEPKKGDVLCSRHFKKTD

FDRSTPNIKLKPGVIPSIF

>033262413.1|Orcinus orca

MVKCCSAVGCASRCLPNSKLKGLTFHVFPTDENVKRKWVLAMKRLDVNAVDIWEPKKGDVLCSRHFKKTD

FDRSTPNIKLKPGVIPSIF

>PNI82799.1|Pan troglodytes

MVKCCSAIGCASRCLPNSKLKGLTFHVFPTDENIKRKWVLAMKRLDVNAAGIWEPKKGDVLCSRHFKKTD

FDRSAPNIKLKPGVIPSIF

>PNJ48765.1|Pongo abelii

MVKCCSAIGCASRCLPNSKLKGLTFHVFPTDENIKRKWVLAMKRLDVNAAGIWEPKKGDVLCSRHFKKTD

FDRSAPNIKLKPGVIPSIF

>JAB44571.1|Callithrix jacchus

MVKCCSAIGCASRCLPNSKLKGLTFHVFPTDENIKRKWVLAMKRLDVNAAGIWEPKKGDVLCSRHFKKTD

FDRSAPNIKLKPGVIPSIF

>AFH29707.1|Macaca mulatta

MVKCCSAIGCASRCLPNSKLKGLTFHVFPTDENIKRKWVLAMKRLDVNAAGIWEPKKGDVLCSRHFKKTD

FDRSAPNIKLKPGVIPSIF

>012600110.1|Microcebus murinus

MVKCCSAIGCASRCLPNSKLKGLTFHVFPTDENIKREWVLAMKRLDVNAAGIWEPKKGDVLCSRHFKKTD

FDRSAPNIKLKPGVIPSIF

>012302412.1|Aotus nancymaae

MVKCCSAIGCASRCLPNSKLKGLTFHVFPTDENIKRKWVLAMKRLDVNAAGIWEPKKGDVLCSRHFKKTD

FDRSAPNIKLKPGVIPSIF

>008067237.1|Carlito syrichta

MVKCCSAIGCASRCLPNSKLRGLTFHVFPTDENVKRKWVLAMKRLDVNAADMWEPKKGDVLCSRHFKKTD

FDRSAPNIKLKPGVVPSIF

>003790127.1|Otolemur garnettii

MVKCCSAIGCASRCLPNSKLKGLTFHVFPTDENIKREWVLAMKRLDVNAAGIWEPKKGDVLCSRHFKKTD

FDRSAPNIKLKPGVIPSIF

>011732031.1|Macaca nemestrina

MVKCCSAIGCASRCLPNSKLKGLTFHVFPTDENIKRKWVLAMKRLDVNAAGIWEPKKGDVLCSRHFKKTD

FDRSAPNIKLKPGVIPSIF

>008955969.1|Pan paniscus

MVKCCSAIGCASRCLPNSKLKGLTFHVFPTDENIKRKWVLAMKRLDVNAAGIWEPKKGDVLCSRHFKKTD

FDRSAPNIKLKPGVIPSIF

>025240966.1|Theropithecus gelada

MVKCCSAIGCASRCLPNSKLKGLTFHVFPTDENIKRKWVLAMKRLDVNAAGIWEPKKGDVLCSRHFKKTD

FDRSAPNIKLKPGVIPSIF

>003265803.1|Nomascus leucogenys

MVKCCSAVGCASRCLPNSKLKGLTFHVFPTDENIKRKWVLAMKRLDVNAAGIWEPKKGDVLCSRHFKKTD

FDRSAPNIKLKPGVIPSIF

>030771973.1|Rhinopithecus roxellana

MVKCCSAIGCASRCLPNSKLKGLTFHVFPTDENIKRKWVLAMKRLDVNAAGIWEPKKGDVLCSRHFKKTD

FDRSAPNIKLKPGVIPSIF

>018881241.2|Gorilla gorilla gorilla

MVKCCSAIGCASRCLPNSKLKGLTFHVFPTDENIKRKWVLAMKRLDVNAAGIWEPKKGDVLCSRHFKKTD

FDRSAPNIKLKPGVIPSIF

>003898800.1|Papio anubis

MVKCCSAIGCASRCLPNSKLKGLTFHVFPTDENIKRKWVLAMKRLDVNAAGIWEPKKGDVLCSRHFKKTD

FDRSAPNIKLKPGVIPSIF

>023045543.1|Piliocolobus tephrosceles

MVKCCSAIGCASRCLPNSKLKGLTFHVFPTDENIKRKWVLAMKRLDVNAAGIWEPKKGDVLCSRHFKKTD

FDRSAPNIKLKPGVIPSIF

>032116917.1|Sapajus apella

MVKCCSAIGCASRCLPNSKLKGLTFHVFPTDENIKRKWVLAMKRLDVNAAGIWEPKKGDVLCSRHFKKTD

FDRSAPNIKLKPGVIPSIF

>032005955.1|Hylobates moloch

MVKCCSAIGCASRCLPNSKLKGLTFHVFPTDENIKRKWVLAMKRLDVNAAGIWEPKKGDVLCSRHFKKTD

FDRSAPNIKLKPGVIPSIF

>033070314.1|Trachypithecus francoisi

MVKCCSAIGCASRCLPNSKLKGLTFHVFPTDENIKRKWVLAMKRLDVNAAGIWEPKKGDVLCSRHFKKTD

FDRSAPNIKLKPGVIPSIF

>AAH22989.1|Homo sapiens

MVKCCSAIGCASRCLPNSKLKGLTFHVFPTDENIKRKWVLAMKRLDVNAAGIWEPKKGDVLCSRHFKKTD

FDRSAPNIKLKPGVIPSIF
