## Additional file for "Synapomorphic variations in the THAP domains of the human THAP protein family and its homologs": THAP7.docx

>025063580.1|Alligator sinensis

MWLENCRRKDPSGQGLWDPTSKYIYFCSKHFEKSCFEMMGIRWTGLSGSSLIPVKMDWSAWSLFPPVKMY

LPSLWTHLMTNAERAVILWYNSPVKLLVLGYLWMNGSTTLPKENGYHRLKEGAIPTIF

>005311791.1|Chrysemys pictabellii

MPRHCSATGCCTRDTRETRNRGISFHRLPKKDNPRRSVWLENCQRKDPSGQGPWNPASDYIYFCSKHFEK

SCFEMVGISGYHRLKEGAVPTVF

>027685254.1|Chelonia mydas

MGWPLGPPPPTSPSALSRGCRAPSSSRENRARGSAAPAPDLGPQRLWSDGNGEMPRHCSAACCCTRDTRE

TRNRGISFHRLPKKDNPRRSVWLENCQRKDPSGQGPWNPASDYIYFCSKHFEKSCFEMVGISGYHRLKEG

AVPTVF

>026513929.1|Terrapene carolinatriunguis

MPRHCSATGCCTRDTRETRNRGISFHRLPKKDNPRRSVWLENCQRKDPSGQGPWNPASDYIYFCSKHFEK

SCFEMVGISGYHRLKEGAVPTVF

>032648878.1|Chelonoidis abingdonii

MPRHCSATGCCTRDTRETRNRGISFHRLPKKDNPRRSMWLENCQRKDPSGQGPWNPVSDYIYFCSKHFEK

SCFEMVGISGYHRLKEGAVPTVF

>020371691.1|Rhincodon typus

MNRWCGWTQAKSGYRLWGSMWSDSGELFLGRAGVRAMPRHCSAEGCTSRDTKDSRINGVTFHRLPKKGRH

QREIWLLNCCRKNPDGTVPWDPPSQFIYFCSKHFEKHCFELVGLSRYHRLRDDAVPTLF

>032869071.1|Amblyraja radiata

MWNDDHYETEAFCGFSQAAPKVPAMPRHCSAEGCRSRDTKDSRINGITFHRLPKKGKSQREIWLLNCCRR

NPDGTVPWTPSSQFVYFCSKHFEKQCFELVGLSGYHRLRDDAVPTLF

>029440515.1|Rhinatrema bivittatum

MTWGCKPLFSPLPCLQMPRHCSAIGCTTRDTRETRNRGISFHRLPKKDNPRRIAWLENCRRTDPSGEGVW

DPSSEYVYFCSKHFEKICFEVVGLSGYHRLKETAVPTIF

>030051253.1|Microcaecilia unicolor

MHLGSRNPRRCRATAQRLAARPVTPERRETAAFPSTGCQRSITPVVSPGLKTVVGQTRVAKGCGTPPQSI

STSVPNILREAASRCGYHRLKETAVPTIF

>001088733.1|Xenopus laevis

MVRSCSAANCVNRQTALSKRKGITFHRFPKEQARRQLWITAVTHSHAAVGTDWTPSIHSSLCSQHFHNIQ

FDRTGQTVRLRDTAVPTIF

>031760995.1|Xenopus tropicalis

MVRSCSAADCVNRQTALSKRKGITFHRFPKEQARRQLWITAVSHSHAAVGTEWTPSIHSSLCSQHFHDKQ

FDRTGQTVRLWDTAVPTIF

>020663905.1|Pogona vitticeps

MPRHCSATGCCTRDTPETRNRGISFHRLPKDNPRRSTWLENCRRKDMTGQGLWDPASKYVYFCSKHFEKS

CFELVGTSGYHRLKEGAVPTIF

>028558169.1|Podarcis muralis

MPRHCSATGCCTRDTPETRNRGISFHRLPKDNPRRAMWLENCRRKDMSGQGLWDPASKYVYFCSKHFEKS

CFELVGTSGYHRLKEGAVPTIF

>033025683.1|Lacerta agilis

MPRHCSATGCCTRDTPEIRNRGISFHRLPKDNPRRAMWLENCRRKDMSGQGLWDPASKYVYFCSKHFEKS

CFELVGTSGYHRLKEGAVPTIF

>JAG66270.1|Boiga irregularis

MPRHCSATGCCTRDTPETRNRGISFHRLPKDNPRRTTWLENCQRKDMSGQGLWDPASKYVYFCSKHFEKN

CFELVGTSGYHRLKEGAVPTIF

>007443582.1|Python bivittatus

MPRHCSATGCCTRDTPETRNRGISFHRLPKDNPRRTTWLENCRRKDLSGQGLWDPASKYVYFCSKHFEKN

CFELVGTSGYHRLKEGAVPTIF

>015665653.1|Protobothrops mucrosquamatus

MPRHCSATGCCTRDTPETRNRGISFHRLPKDNPRRTTWLENCQRKNMSGQGLWDPASKYVYFCSKHFEKN

CFELVGTSGYHRLKEGAVPTIF

>032077555.1|Thamnophis elegans

MAFRLFVSMPALSLGMAASVTLVLTRLNPAELPDPEKVCTKAGQIILKIECNQLIPKPSEDLPSPPVIYI

PDTGVSEPVTTLICGGCFLCSGYHRLKEGAVPTIF

>021238309.1|Numida meleagris

MPRHCSAAGCCTRDTRETRSRGISFHRLPKKDNPRRALWLENSRRRDASGEGRWDPASKYIYFCSQHFEK

SCFEIVGFSGYHRLKEGAVPTVF

>015128753.1|Gallus gallus

MPRHCSAAGCCTRDTRETRSRGISFHRLPKKDNPRRALWLENSRRRDASGEGRWDPASKYIYFCSQHFEK

SCFEIVGFSGYHRLKEGAVPTVF

>027636291.1|Falco peregrinus

MPRHCSAAGCCTRDTRETRNRGISFHRWVPPKDNPRRALWLENSRRRDASGEGLWDPASKYIYFCSQHFE

KSCFEIVGFSGYHRLKEGAVPTVF

>027657025.1|Falco cherrug

MAAVAAGTARAQAQGHCLRRALEAVGDGAAATARHVAALRARLRCDPLTLSRTVRRMLREEDEDSEDSED

SESSPWTLRELLETPLGLENSRRRDASGEGLWDPASKYIYFCSQHFEKSCFEIVGFSGYHRLKEGAVPTV

F

>031466217.1|Phasianus colchicus

MREREAGTVYDMGGGPISLQMPRHCSAAGCCTRDTRETRSRGISFHRLPKKDNPRRALWLENSRRRDASG

EGRWDPASKYIYFCSQHFEKSCFEIVGFSGYHRLKEGAVPTVF

>010726687.1|Meleagris gallopavo

MSFPPPPYPPISPPRLPKKDNPRRALWLENSRRRDASGEGRWDPASKYIYFCSQHFEKSCFEIVGFSGYH

RLKEGAVPTVF

>032060999.1|Aythya fuligula

MPRHCSAAGCCTRDTRETRNRGISFHRLPKKDNPRRALWLENSRRRDASGEGRWDPASKYIYFCSQHFEK

SCFEIVGFSGYHRLKEGAVPTVF

>032939985.1|Catharus ustulatus

MPRHCSAAGCCTRDTRDTRGRGISFHRLPRRDDPRRAQWLENSRRRDPAGGGRWDPSSKYIYFCSQHFEQ

SCFELVGYSGYHRLKEGAVPTVF

>001158876.1|Salmo salar

MAEQHSDLENAINTLVTQFHAAAANNGPTLQTQEFRGLLSSQLPNLVTAKTQTAFDRILLSEKLSSYPAQ

MPRHCSAGGCKSRDTRENRKAGITFHRLPKRGTPRRDLWIINSHRKGPQGQGPWDPQSNFIYFCSKHFTP

ESFELSGVSGYRRLKDDALPTVF

>020480454.1|Monopterus albus

MPRHCSAGGCKSRDNRETRNAGITFHKLPKGATRRNLWITNSHRAETWDPRTNFVYFCSKHFTPESFELT

GCSGIRRLKEDAFPTVF

>022073184.1|Acanthochromis polyacanthus

MPRHCSAGGCKSRDNRETRNAGITFHKLPKGATRRKLWITNSHRADSWDPQTDFVYFCSKHFTAESFELT

GCSGIKRLKEDAFPTVF

>007254778.2|Astyanax mexicanus

MPRHCSAAGCNSRDTREARKSGLTFHRLPKRGNPRRATWILNSCRKGPEGKGQWDPQSDYIYFCSKHFTP

DSFELSGVSGYRRLKDDAVPTLI

>023151005.1|Amphiprion ocellaris

MPRHCSAGGCKCRDNRETRNAGITFHKLPKGATRRKLWITNSHRADSWDPQTDFVYFCSRHFTPESFELT

GCSGIKRLKEDAFPTVF

>023656915.1|Paramormyrops kingsleyae

MPRHCSAAGCKSRDTRETRKAGITFHRLPKRGSSRRTQWILNSQRKDPQGKGQWDPQSHFIYFCSKHFTP

ESFELSGVSGYRRLKEDALPTVF

>004561057.3|Maylandia zebra

MKKEEKNSAIILRSTAGMPRHCSAGGCKSRDNHETRNAGVTFHKLPKGAARRNLWISNCHRADSWDPQTN

FVYFCSKHFPPESFELSGSSGIRRLKEDAVPTLF

>024920893.1|Cynoglossus semilaevis

MFGIYASHTSVFLSAVFPLLLCHFQMPRHCSAGGCKSRDNRETRNAGVTFHKLPKGATRRNLWITNSHRS

NSWDPQTDFVYFCSKHFTPESFELTNYSGIRRLKDDALPTVF

>025758318.1|Oreochromis niloticus

MRKMKKEEKKISHHSEMPRHCSAGGCKSRDNHETRNAGVTFHKLMLWMTRLPKGAARRNLWISNSHRADS

WDPQTDFVYFCSKHFPPESFELSGSSGIRRLKEDAVPTLF

>026011778.1|Astatotilapia calliptera

MKKEEKNSAIILRSTAGMPRHCSAGGCKSRDNHETRNAGVTFHKLPKGAARRNLWISNCHRADSWDPQTN

FVYFCSKHFPPESFELSGSSGIRRLKEDAVPTLF

>001096668.1|Danio rerio

MPRHCSAVGCKSRDTKDVRKSGITFHRLPKKGNPRRTTWIINSRRKGPEGKGQWDPQSGFIYFCSKHFTP

DSFELSGVSGYHRLKDDAIPTVF

>019109132.1|Larimichthys crocea

MPRHCSAGGCKSRDNRETRDAGITFHKLPKGETRRHLWITNSHRADSWDPQTDFVYFCSKHFTPESFELT

GCSGIRRLREDAFPTVF

>028669385.1|Erpetoichthys calabaricus

MPRHCSAAGCKSRDTSETRKNGVTFHRLPKRGNPRRLLWLANCRRTDPESQAMWDPKSQFIYFCSRHFTK

DSFELVGTSGYHRLKDDALPTIF

>028819490.1|Denticeps clupeoides

MNKTPRHLNSSTRGRTSLPSHPFLVKNHGLRSGGADPHPNRFTLGCEPAQQELKMPRHCSAVGCNSRDTR

EARKSGITFHRLPKRGNPRRALWIINSHRKGPKGQGSWDPQSDYVYFCSRHFTAESFELPGVSGYRRVKE

DAVPTIF

>018621416.2|Scleropages formosus

MPRHCSAAGCKSRDTSETRKAGITFHRLPKRGSPRRTLWIINSQRKDPQGKGQWDPQSHFIYFCSKHFTP

ESFELSGISGYRRLKEDALPTVF

>TNN54458.1|Liparis tanakae

MKEGKEGREERIERKERRKERKEGNRGREGGSEERKKKKGGRKGGRKERKEWMEEGRKEREEGKERKTPS

GKGTQRVYSLGQLSASLAVMLLLQAEQSVVAETVSDVNRRLRAQVPGRGLIGLSCRGPGVTQRDASCLWS

RDTVHQAGGLQPAPGLRHIEEMNSDCECKSSSDVDEGGSDKKGIYNRLRNSVNASSRRLNVGGGLASYSR

PSASLFQSAATEMPRHCSAGGCKSRDNRVTRNAGITFHKLPKGAARRNLWITNSHRADSWDPQTDFVYFC

SKHFSPDSFELTGCSGIRRLKEDAFPTVF

>029919098.1|Myripristis murdjan

MPRHCSAGGCKSRDNRETRSAGITFHKLPKGRTRRNLWITNSHRTDLWDPQTDFVYFCSKHFTPESFELT

GCSGIRRLREDALPTVF

>030003463.1|Sphaeramia orbicularis

MPRHCSAGGCKSRDNRDTRNAGITFHKLPKRAPRRRLWISNSHRASSWDPQTDFVYFCSKHFTPDSFELT

GCSGIRRLKEDAFPTVF

>028669381.1|Erpetoichthys calabaricus

MPRHCSAAGCKSRDTSETRKNGVTFHRLPKRGNPRRLLWLANCRRTDPESQAMWDPKSQFIYFCSRHFTK

DSFELVGTSGYHRLKDDALPTIF

>030202294.1|Gadus morhua

MPRHCSAGGCKSRDNRETRKAGITFHKLPKGASRRNLWITNSHRKGLWDPQTDFVYFCSKHFTPENFELT

GYSGIRRLREDAFPTVF

>030597301.1|Archocentrus centrarchus

MPRHCSAGGCKSRDNHETRNAGVTFHKLPKGATRRTLWISNSHRADSWDPQTDFVYFCSKHFTPESFELS

GIGGIRRLKEDAVPTLF

>030644572.1|Chanos chanos

MPRHCSAAGCKSRDTKESRVAGITFHRLPKRGTPRRTLWIINSRRKGPEGKGQWDPQNDFIYFCSKHFTP

ESFELSGVSGYRRLKDDAVPTVF

>031134696.1|Sander lucioperca

MMPRHCSAGGCKSRDNRETRNAGITFHKLPKGATRRNLWITNSHRADYWDPQTDFVYFCSKHFTPESFEL

TGCSGIRRLKEDAFPTVF

>031612473.1|Oreochromis aureus

MPRHCSAGGCKSRDNHETRNAGVTFHKLPKGAARRNLWISNSHRADSWDPQTDFVYFCSKHFPPESFELS

GSSGIRRLKEDAVPTLF

>031694600.1|Anarrhichthys ocellatus

MDRLPKGATRRNLWITNSHRADSWDPQTDFVYFCSKHFTPDSFELTGCSGIRRLKEDAFPTVF

>026156621.1|Mastacembelus armatus

MPRHCSAGGCKSRDNRDTRNAGITFHKLPKGETRRSLWITNSHRADSWDPQTDFVYFCSKHFTPESFELT

GYSGIRRLKEDAFPTVF

>026206054.1|Anabas testudineus

MPRHCSAGGCKSRDNRETRNAGITFHKLPKGATRRNLWITNSHRADSWDPQTDFVYFCSKHFTPESFELT

GCSLFIRSGIRRLKEDAFPTVF

>33492446.1|Epinephelus lanceolatus

MPRHCSAGGCKSRDNRETRNAGITFHKLPKGAIRRNLWITNSHRADSWDPQTDFVYFCSKHFTPESFELT

GCSGIRRLKEDAFPTVF

>028905070.1|Ornithorhynchus anatinus

MPRHCSAAGCCTRDTRETRNRGISFHRLPKKDNPRRGLWLANCRRTDPSGQGLWDPASEYIYFCSKHFEE

NCFELVGISGYHRLKEGAVPTIF

>021543555.1|Neomonachus schauinslandi

MPRHCSAAGCCTRDTRETRNRGISFHRLPKKDNPRRGLWLANCQRLDPSGQGLWDPASEYIYFCSKHFEE

NCFELVGISGYHRLKEGAVPTIF

>022266261.1|Canis lupus familiaris

MPRHCSAAGCCTRDTRETRNRGISFHRSARVRMRPFYPLRGATIAHAPSSVWREACICRLPKKDNPRRGL

WLANCQRLDPSGQGLWDPASEYIYFCSKHFEENCFELVGISGYHRLKEGAVPTIF

>006938784.1|Felis catus

MPRHCSAAGCCTRDTRETRNRGISFHRLPKKDNPRRGLWLANCQRLDPSGQGLWDPASEYIYFCSKHFEE

NCFELVGISGYHRLKEGAVPTIF

>003419322.1|Loxodonta africana

MPRHCSAAGCCTRDTRETRNRGISFHRLPKKDNPRRGLWLANCQRLDPSGQGLWDPASEYIYFCSKHFEE

NCFELVGISGYHRLKEGAVPTIF

>011354232.1|Pteropus vampyrus

MPRHCSAAGCCTRDTRETRNRGISFHRLPKKDNPRRGLWLANCQRLDPSGQGLWDPASEYIYFCSKHFEE

NCFELVGISGYHRLKEGAVPTIF

>023502542.1|Equus caballus

MPRHCSAAGCCTRDTRETRNRGISFHRLPKKDNPRRGLWLANCQRLDPSGQGLWDPASEYIYFCSKHFEE

NCFELVGIRLLFPHSGYHRLKEGAVPTIF

>006099584.1|Myotis lucifugus

MPRHCSAAGCCTRDTRETRNRGISFHRLPKKDNPRRGLWLANCQRLDPSGQGLWDPASEYIYFCSKHFEE

NCFELVGISGYHRLKEGAVPTIF

>024412989.1|Desmodus rotundus

MPRHCSAAGCCTRDTRETRNRGISFHRLPKKDNPRRGLWLANCQRLDPSGQGLWDPASEYIYFCSKHFEE

NCFELVGISGYHRLKEGAVPTIF

>024900318.1|Pteropus alecto

MPRHCSAAGCCTRDTRETRNRGISFHRLPKKDNPRRGLWLANCQRLDPSGQGLWDPASEYIYFCSKHFEE

NCFELVGISGYHRLKEGAVPTIF

>025330700.1|Canis lupus dingo

MPRHCSAAGCCTRDTRETRNRGISFHRSARVRMRPFYPLRGATIAHAPSSVWREACICRLPKKDNPRRGL

WLANCQRLDPSGQGLWDPASEYIYFCSKHFEENCFELVGISGYHRLKEGAVPTIF

>025713030.1|Callorhinus ursinus

MPRHCSAAGCCTRDTRETRNRGISFHRLPKKDNPRRGLWLANCQRLDPSGQGLWDPASEYIYFCSKHFEE

NCFELVGISGYHRLKEGAVPTIF

>025789276.1|Puma concolor

MPRHCSAAGCCTRDTRETRNRGISFHRLPKKDNPRRGLWLANCQRLDPSGQGLWDPASEYIYFCSKHFEE

NCFELVGISGYHRLKEGAVPTIF

>025838595.1|Vulpes vulpes

MPRHCSAAGCCTRDTRETRNRGISFHRLPKKDNPRRGLWLANCQRLDPSGQGLWDPASEYIYFCSKHFEE

NCFELVGISGYHRLKEGAVPTIF

>026343643.1|Ursus arctoshorribilis

MPRHCSAAGCCTRDTRETRNRGISFHRLPKKDNPRRGLWLANCQRLDPSGQGLWDPASEYIYFCSKHFEE

NCFELVGISGYHRLKEGAVPTIF

>026905539.1|Acinonyx jubatus

MPRHCSAAGCCTRDTRETRNRGISFHRLPKKDNPRRGLWLANCQRLDPSGQGLWDPASEYIYFCSKHFEE

NCFELVGISGYHRLKEGAVPTIF

>027431970.1|Zalophus californianus

MPRHCSAAGCCTRDTRETRNRGISFHRLPKKDNPRRGLWLANCQRLDPSGQGLWDPASEYIYFCSKHFEE

NCFELVGISGYHRLKEGAVPTIF

>006140249.1|Tupaia chinensis

MPRHCSAAGCCTRDTRETRNRGISFHRLPKKDNPRRGLWLANCQRLDPSGQGLWDPASEYIYFCSKHFEE

NCFELVGISGYHRLKEGAVPTIF

>027950742.1|Eumetopias jubatus

MPRHCSAAGCCTRDTRETRNRGISFHRLPKKDNPRRGLWLANCQRLDPSGQGLWDPASEYIYFCSKHFEE

NCFELVGISGYHRLKEGAVPTIF

>028002856.1|Eptesicus fuscus

MPRHCSAAGCCTRDTRETRNRGISFHRLPKKDNPRRGLWLANCQRLDPSGQGLWDPASEYIYFCSKHFEE

NCFELVGISGYHRLKEGAVPTIF

>028384503.1|Phyllostomus discolor

MPRHCSAAGCCTRDTRETRNRGISFHRLPKKDNPRRGLWLANCQRLDPSGQGLWDPASEYIYFCSKHFEE

NCFELVGISGYHRLKEGAVPTIF

>029777249.1|Suricata suricatta

MPRHCSAAGCCTRDTRETRNRGISFHRLPKKDNPRRGLWLANCQRLDPSGQGLWDPASEYIYFCSKHFEE

NCFELVGISGYHRLKEGAVPTIF

>030148195.1|Lynx canadensis

MPRHCSAAGCCTRDTRETRNRGISFHRLPKKDNPRRGLWLANCQRLDPSGQGLWDPASEYIYFCSKHFEE

NCFELVGISGYHRLKEGAVPTIF

>006744563.1|Leptonychotes weddellii

MPRHCSAAGCCTRDTRETRNRGISFHRLPKKDNPRRGLWLANCQRLDPSGQGLWDPASEYIYFCSKHFEE

NCFELVGISGYHRLKEGAVPTIF

>032165493.1|Mustela erminea

MPRHCSAAGCCTRDTRETRNRGISFHRLPKKDNPRRGLWLANCQRLDPSGQGLWDPASEYIYFCSKHFEE

NCFELVGISGYHRLKEGAVPTIF

>032253456.1|Phoca vitulina

MPRHCSAAGCCTRDTRETRNRGISFHRLPKKDNPRRGLWLANCQRLDPSGQGLWDPASEYIYFCSKHFEE

NCFELVGISGYHRLKEGAVPTIF

>032731852.1|Lontra canadensis

MPRHCSAAGCCTRDTRETRNRGISFHRLPKKDNPRRGLWLANCQRLDPSGQGLWDPASEYIYFCSKHFEE

NCFELVGISGYHRLKEGAVPTIF

>032953427.1|Rhinolophus ferrumequinum

MPRHCSAAGCCTRDTRETRNSGISFHRLPKKDNPRRGLWLANCQRLDPSGQGLWDPASEYIYFCSKHFEE

NCFELVGISGYHRLKEGAVPTIF

>027703271.1|Vombatus ursinus

MPRHCSAAGCCTRDTRETRTRGISFHRLPKKDNPRRGLWLANCQRLDPSGQGLWDPASEYIYFCSKHFEE

NCFELVGISGYHRLKEGAVPTIF

>031804891.1|Sarcophilus harrisii

MPRHCSAAGCCTRDTRETRTRGISFHRLPKKDNPRRGLWLANCRRLDPSGQGLWDPASEYIYFCSKHFEE

NCFELVGFSGYHRLKEGAVPTIF

>020759105.1|Odocoileus virginianustexanus

MPRHCSAAGCCTRDTRETRNRGISFHRLPKKDNPRRGLWLANCQRLDPSGQGLWDPASEYIYFCSKHFEE

NCFELVGISGYHRLKEGAVPTIF

>003133046.2|Sus scrofa

MRARPARAPHRTTSPFLSESFRSCARRPAGRPLPGPAPPLSERRALPGLYGTDFRREPHLARLRRRSNGE

TMPRHCSAAGCCTRDTRETRNRGISFHRLPKKDNPRRGLWLANCQRLDPSGQGLWDPASEYIYFCSKHFE

ENCFELVGISGYHRLKEGAVPTIF

>024619494.1|Neophocaena asiaeorientalisasiaeorientalis

MPRHCSAAGCCTRDTRETRNRGISFHRLPKKDNPRRGLWLANCQRLDPSGQGLWDPASEYIYFCSKHFEE

NCFELVGISGYHRLKEGAVPTIF

>024833275.1|Bos taurus

MPRHCSAAGCCTRDTRETRNRGISFHRLPKKDNPRRGLWLANCQRLDPSGQGLWDPASEYIYFCSKHFEE

NCFELVGISGYHRLKEGAVPTIF

>006065630.1|Bubalus bubalis

MPRHCSAAGCCTRDTRETRNRGISFHRLPKKDNPRRGLWLANCQRLDPSGQGLWDPASEYIYFCSKHFEE

NCFELVGISGYHRLKEGAVPTIF

>026944455.1|Lagenorhynchus obliquidens

MPRHCSAAGCCTRDTRETRNRGISFHRLPKKDNPRRGLWLANCQRLDPSGQGLWDPASEYIYFCSKHFEE

NCFELVGISGYHRLKEGAVPTIF

>027812685.1|Ovis aries

MPRHCSAAGCCTRDTRETRNRGISFHRLPKKDNPRRGLWLANCQRLDPSGQGLWDPASEYIYFCSKHFEE

NCFELVGISGYHRLKEGAVPTIF

>007196342.1|Balaenoptera acutorostratascammoni

MPRHCSAAGCCTRDTRETRNRGISFHRLPKKDNPRRGLWLANCQRLDPSGQGLWDPASEYIYFCSKHFEE

NCFELVGISGYHRLKEGAVPTIF

>007130697.1|Physeter catodon

MPRHCSAAGCCTRDTRETRNRGISFHRLPKKDNPRRGLWLANCQRLDPSGQGLWDPASEYIYFCSKHFEE

NCFELVGISGYHRLKEGAVPTIF

>029079836.1|Monodon monoceros

MPRHCSAAGCCTRDTRETRNRGISFHRLPKKDNPRRGLWLANCQRLDPSGQGLWDPASEYIYFCSKHFEE

NCFELVGISGYHRLKEGAVPTIF

>022407232.1|Delphinapterus leucas

MPRHCSAAGCCTRDTRETRNRGISFHRLPKKDNPRRGLWLANCQRLDPSGQGLWDPASEYIYFCSKHFEE

NCFELVGISGYHRLKEGAVPTIF

>030692199.1|Globicephala melas

MPRHCSAAGCCTRDTRETRNRGISFHRLPKKDNPRRGLWLANCQRLDPSGQGLWDPASEYIYFCSKHFEE

NCFELVGISGYHRLKEGAVPTIF

>031299044.1|Camelus dromedarius

MPRHCSAAGCCTRDTRETRNRGISFHRLPKKDNPRRGLWLANCQRLDPSGQGLWDPASEYIYFCSKHFEE

NCFELVGISGYHRLKEGAVPTIF

>015107396.1|Vicugna pacos

MPRHCSAAGCCTRDTRETRNRGISFHRLPKKDNPRRGLWLANCQRLDPSGQGLWDPASEYIYFCSKHFEE

NCFELVGISGYHRLKEGAVPTIF

>032328023.1|Camelus ferus

MPRHCSAAGCCTRDTRETRNRGISFHRLPKKDNPRRGLWLANCQRLDPSGQGLWDPASEYIYFCSKHFEE

NCFELVGISGYHRLKEGAVPTIF

>032459591.1|Phocoena sinus

MPRHCSAAGCCTRDTRETRNRGISFHRLPKKDNPRRGLWLANCQRLDPSGQGLWDPASEYIYFCSKHFEE

NCFELVGISGYHRLKEGAVPTIF

>004276066.1|Orcinus orca

MPRHCSAAGCCTRDTRETRNRGISFHRLPKKDNPRRGLWLANCQRLDPSGQGLWDPASEYIYFCSKHFEE

NCFELVGISGYHRLKEGAVPTIF

>AAH18221.1|Mus musculus

MPRHCSAAGCCTRDTRETRNRGISFHRLPKKDNPRRGLWLANCQRLDPSGQGLWDPTSEYIYFCSKHFEE

NCFELVGISGYHRLKEGAVPTIF

>JAO01097.1|Heterocephalus glaber

MPRHCSAAGCCTRDTRETRNRGISFHRLPKKDNPRRGLWLANCQRLDPSGQGLWDPASEYIYFCSKHFEE

NCFELVGISGYHRLKEGAVPTIF

>EDL77894.1|Rattus norvegicus

MPRHCSAAGCCTRDTRETRNRGISFHRLPKKDNPRRGLWLANCQRLDPSGQGLWDPTSEYIYFCSKHFEE

NCFELVGISGYHRLKEGAVPTIF

>020016233.1|Castor canadensis

MPRHCSAAGCCTRDTRETRNRGISFHRLPKKDNPRRGLWLANCQRLDPSGQGLWDPASEYIYFCSKHFEE

NCFELVGISGYHRLKEGAVPTIF

>005077610.1|Mesocricetus auratus

MPRHCSAAGCCTRDTRETRNRGISFHRLPKKDNPRRGLWLANCQRLDPSGQGLWDPASEYIYFCSKHFEE

NCFELVGISGYHRLKEGAVPTIF

>021510191.1|Meriones unguiculatus

MPRHCSAAGCCTRDTRETRNRGISFHRLPKKDNPRRGLWLANCQRLDPSGQGLWDPASEYIYFCSKHFEE

NCFELVGISGYHRLKEGAVPTIF

>003478414.1|Cavia porcellus

MPRHCSAAGCCTRDTRETRNRGISFHRLPKKDNPRRGLWLANCQRLDPSGQGLWDPASEYIYFCSKHFEE

NCFELVGISGYHRLKEGAVPTIF

>023567624.1|Octodon degus

MPRHCSAAGCCTRDTRETRNRGISFHRLPKKDNPRRGLWLANCQRLDPSGQGLWDPASEYIYFCSKHFEE

NCFELVGISGYHRLKEGAVPTIF

>005370043.1|Microtus ochrogaster

MPRHCSAAGCCTRDTRETRNRGISFHRLPKKDNPRRGLWLANCQRLDPSGQGLWDPASEYIYFCSKHFEE

NCFELVGISGYHRLKEGAVPTIF

>007641656.1|Cricetulus griseus

MPRHCSAAGCCTRDTRETRNRGISFHRLPKKDNPRRGLWLANCQRLDPSGQGLWDPASEYIYFCSKHFEE

NCFELVGISGYHRLKEGAVPTIF

>028635487.1|Grammomys surdaster

MPRHCSAAGCCTRDTRETRNRGISFHRLPKKDNPRRGLWLANCQRLDPSGQGLWDPTSEYIYFCSKHFEE

NCFELVGISGYHRLKEGAVPTIF

>028712512.1|Peromyscus leucopus

MWVRGIKLRSSRLASSAETPAALGGVPSGWSQGFLGGGSFSARASAHEVRGALQTAPRLQRANGERSAVA

LDTQTKPASDRTADPGISSSSPHSQHPVALPPGSLPVPPLEGGRHRRRSPYATNFRIGPPIARLWRRSNG

QTVSARAWWRARAHTRRRRWGRALGRAGDGAFPRLPKKDNPRRGLWLANCQRLDPSGQGLWDPASEYIYF

CSKHFEESCFELVGISGYHRLKEGAVPTIF

>021040580.1|Mus caroli

MPRHCSAAGCCTRDTRETRNRGISFHRLPKKDNPRRGLWLANCQRLDPSGQGLWDPTSEYIYFCSKHFEE

NCFELVGISGYHRLKEGAVPTIF

>021065222.1|Mus pahari

MPRHCSAAGCCTRDTRETRNRGISFHRLPKKDNPRRGLWLANCQRLDPSGQGLWDPTSEYIYFCSKHFEE

NCFELVGISGYHRLKEGAVPTIF

>008840470.2|Nannospalax galili

MGPLQKQLELLTAEPSLQLHTRLKTHPHCPPPSWYVSCFEYWSWVPPCNNQVGFSSQDISLRGPQSPTVP

HTTAPSRDLGQAPPSYTQTSHVDNSITQSALPASGTHPLGALPVPPLGATTARCPCLYATNFRRDPPIAG

LWRRSNGKAMPRHCSAAGCCTRDTRETRNRGISFHRLPKKDNPRRGLWLANCQRLDPSGQGLWDPASEYI

YFCSKHFEENCFELVGISGYHRLKEGAVPTIF

>031219144.1|Mastomys coucha

MPRHCSAAGCCTRDTRETRNRGISFHRLPKKDSPRRVLWLANCQRLDPSGKGLWDPTSEYIYFCSKHFEE

NCFELVGISGYHRLKEGAVPTIF

>032755759.1|Rattus rattus

MPRHCSAAGCCTRDTRETRNRGISFHRLPKKDNPRRGLWLANCQRLDPSGQGLWDPTSEYIYFCSKHFEE

NCFELVGISGYHRLKEGAVPTIF

>RLQ68578.1|Cricetulus griseus

MPRHCSAAGCCTRDTRETRNRGISFHSSLVLTVLWLCRLPKKDNPRRGLWLANCQRLDPSGQGLWDPASE

YIYFCSKHFEENCFELVGISGYHRLKEGAVPTIF

>PNI12694.1|Pan troglodytes

MPRHCSAAGCCTRDTRETRNRGISFHRLPKKDNPRRGLWLANCQRLDPSGQGLWDPASEYIYFCSKHFEE

NCFELVGISGYHRLKEGAVPTIF

>PNJ13911.1|Pongo abelii

MPRHCSAAGCCTRDTRETRNRGISFHRLPKKDNPRRGLWLANCQRLDPSGQGLWDPASEYIYFCSKHFEE

NCFELVGISGYHRLKEGAVPTIF

>JAB16893.1|Callithrix jacchus

MPRHCSAAGCCTRDTRETRSRGISFHRLPKKDNPRRGLWLANCQRLDPSGQGLWDPASEYIYFCSKHFEE

NCFELVGISGYHRLKEGAVPTIF

>AFI38390.1|Macaca mulatta

MPRHCSAAGCCTRDTRETRNRGISFHRLPKKDNPRRGLWLANCQRLDPSGQGLWDPASEYIYFCSKHFEE

NCFELVGISGYHRLKEGAVPTIF

>012632129.1|Microcebus murinus

MWAQGRRPRSRRFRSPPAGAVSCPAPRPEHRPSQSLYGTNFRRPPHLARFWRRSNGETMPRHCSAAGCCT

RDTRETRNRGISFHRLPKKDNPRRGLWLANCRRLDPSGQGLWDPASEYIYFCSKHFEENCFELVGISGYH

RLKEGAVPTIF

>012321945.1|Aotus nancymaae

MPRHCSAAGCCTRDTRETRSRGISFHRLPKKDNPRRGLWLANCQRLDPSGQGLWDPASEYIYFCSKHFEE

NCFELVGISGYHRLKEGAVPTIF

>008067012.1|Carlito syrichta

MPRHCSAAGCCTRDTRETRNRGISFHRLPKKDNPRRGLWLANCQRLDPSGQGLWDPASEYIYFCSKHFEE

NCFELVGISGYHRLKEGAVPTIF

>012668146.1|Otolemur garnettii

MPRHCSAAGCCTRDTRETRNRGISFHRLPKKDNPRRGLWLANCQRLDPSGQGLWDPTSEYIYFCSKHFED

NCFELVGISGYHRLKEGAVPTIF

>011738084.1|Macaca nemestrina

MPRHCSAAGCCTRDTRETRNRGISFHRLPKKDNPRRGLWLANCQRLDPSGQGLWDPASEYIYFCSKHFEE

NCFELVGISGYHRLKEGAVPTIF

>014198717.1|Pan paniscus

MPGEPLATELPPLSQMPRHCSAAGCCTRDTRETRNRGISFHRLPKKDNPRRGLWLANCQRLDPSGQGLWD

PASEYIYFCSKHFEENCFELVGISGYHRLKEGAVPTIF

>025254605.1|Theropithecus gelada

MPRHCSAAGCCTRDTRETRNRGISFHRLPKKDNPRRGLWLANCQRLDPSGQGLWDPASEYIYFCSKHFEE

NCFELVGISGYHRLKEGAVPTIF

>030671772.1|Nomascus leucogenys

MPRHCSAAGCCTRDTRETRNRGISFHRLPKKDNPRRGLWLANCQRLDPSGQGLWDPASEYIYFCSKHFEE

NCFELVGISGYHRLKEGAVPTIF

>010367292.1|Rhinopithecus roxellana

MPRHCSAAGCCTRDTRETRNRGISFHRLPKKDNPRRGLWLANCQRLDPSGQGLWDPASEYIYFCSKHFEE

NCFELVGISGYHRLKEGAVPTIF

>004063122.1|Gorilla gorilla gorilla

MPRHCSAAGCCTRDTRETRNRGISFHRLPKKDNPRRGLWLANCQRLDPSGQGLWDPASEYIYFCSKHFEE

NCFELVGISGYHRLKEGAVPTIF

>003905306.1|Papio anubis

MPRHCSAAGCCTRDTRETRNRGISFHRLPKKDNPRRGLWLANCQRLDPSGQGLWDPASEYIYFCSKHFEE

NCFELVGISGYHRLKEGAVPTIF

>023053729.1|Piliocolobus tephrosceles

MPRHCSAAGCCTRDTRETRNRGISFHRLPKKDNPRRGLWLANCQRLDPSGQGLWDPASEYIYFCSKHFEE

NCFELVGISGYHRLKEGAVPTIF

>032127227.1|Sapajus apella

MPRHCSAAGCCTRDTRETRSRGISFHRLPKKDNPRRGLWLANCQRLDPSGQGLWDPASEYIYFCSKHFEE

NCFELVGISGYHRLKEGAVPTIF

>031994922.1|Hylobates moloch

MPRHCSAAGCCTRDTRETRNRGISFHRLPKKDNPRRGLWLANCQRLDPSGQGLWDPASEYIYFCSKHFEE

NCFELVGISGYHRLKEGAVPTIF

>033089981.1|Trachypithecus francoisi

MPRHCSAAGCCTRDTRETRNRGISFHRLPKKDNPRRGLWLANCQRLDPSGQGLWDPASEYIYFCSKHFEE

NCFELVGISGYHRLKEGAVPTIF

>001008695.1||Homo sapiens

MPRHCSAAGCCTRDTRETRNRGISFHRLPKKDNPRRGLWLANCQRLDPSGQGLWDPASEYIYFCSKHFEE

DCFELVGISGYHRLKEGAVPTIF

>**026798166.1|Pangasianodon hypophthalmus**

MPRHCSATGCKSRDNKEARLAGVTFHRLPKKGNPRRTTWIVNARRKGPGGKGLWEPQSDYIYFCSKHFTP

DSFELSGVSGYRRLKDDAVPMLV

>**027033755.1|Tachysurus fulvidraco**

MPRHCSATGCKSRDNKDARLAGVTFHRLPKKGNPRRTTWIVNARRKGPGEKGLWEPQSDYIYFCSKHFTP

DSFELSGVSGYRRLKDDAVPMLV
