## Additional file for "Synapomorphic variations in the THAP domains of the human THAP protein family and its homologs": THAP8.docx

>PFX17745.1|Stylophora pistillata

MGGQVAPEAAETTSLHPRQRWRNVHDIISPVWMRWLKEFVPVLNSQPKWTSECRDLKFEDVVLVIQLDTP

RGRWPLGRIVEVDANLGVPQGSILGPALFPIFINDLSKILEHSAADLYADDTTISINGDYRSAPGALNQA

DVGKVAQRTTDNKMVLNESKTKTMLAAGKRLPKKISSSSLTVQVNSVELEQVQSHKLLAKIDHCALVYQR

LNGVCPDYTLEVLKRNIEIRSTERQSRKKISFQPMINIFDIDKMSYSLLERKDTPIANMAAPARKRGFSV

VTVAKANSTLFMCSIDSSKLASKEFASEAEVKESLEKIYSYFRGMPDNCCVPLYHKSGYRVGPDGKKVTY

HDLPLRNPKRLKTWLVKIRRDASKDFKLTKRTKICSLHFKDSDFRLTLKGRRYIKDDAIPSIF

>TNN56577.1|Liparis tanakae

MPKYCSVPNCKNDSGNGGDRKSFYKFPLQDPVRLQQWLRNMRRANWAPSRHQYICHEHFAPSCFKVRWGI

RYLESDAVPTVF

>JAO88165.1|Poeciliopsis prolifica

MPKYCSVPNCRSSGERKSFYKFPLQDPARLQQWVRNIGRENWTPSRHQYICHEHFAPSCFKVRWGIRYLD

ADAVPTVF

>005283783.3|Chrysemys pictabellii

MGFPLHNPERLQQWLSQMNQEKWVPTRYQHLCSEHFAPSCFEYRWGVRYLKPDAIPTIF

>027686984.1|Chelonia mydas

MCHVVGVARFPLHNPERLQQWLSQMNQEKWVPTRYQHLCSEHFAPSCFEYRWGVRYLKPDAVPTIF

>026514679.1|Terrapene carolinatriunguis

MNQEKWVPTRYQHLCSEHFAPSCFEYRWGVRYLKPDAVPTIF

>030400415.1|Gopherus evgoodei

MPKYCRAPNCSNSAGQRRPGGERLSFYRFPLHNPERLQQWLSQMNQEKWVPTRYQHLCSEHFAPSCFEYR

WGVRYLKPDAVPTIF

>032632107.1|Chelonoidis abingdonii

MVLSYRFPLHNPERLQQWLSQMNQEKWVPTRYQHLCSEHFAPSCFEYRWGVRYLKPDAVPTIF

>020642719.1|Pogona vitticeps

MTKYCRAPNCSNSAGQPRPDRRRLSFYKFPLHNPERLRQWLSQMNQEKWVPTKHQHLCSEHFAPSCFEYR

WGVRYLKPDAIPTIF

>033014516.1|Lacerta agilis

MTKYCRAPNCCNSAKEPRPDNRRLSFYKFPLHNPERLQQWLSQMNQEKWVPTKHQHLCSDHFAPSCFEYR

WGSRFLKPDAVPTIF

>026544871.1|Notechis scutatus

MTKYCRAPRCSNSAGQPRRDQRRLSFYKFPLHSPERLRQWLSQMNQEKWTPTKHQHLCSDHFAPSCFEYR

WGVRYLKPDAVPTIF

>026571185.1|Pseudonaja textilis

MTKYCRAPRCSNSAGQPRRDQRRLSFYKFPLHSPERLRQWLSQMNQEKWTPTKHQHLCSDHFAPSCFEYR

WGVRYLKPDAVPTIF

>015676517.1|Protobothrops mucrosquamatus

MTKYCRAPRCSNSAGQPRRDQRRLSFYKFPLHSPERLRQWLSQMNQEKWVPTKHQHLCSDHFAPSCFEYR

WGVRYLKPDAVPTIF

>032083104.1|Thamnophis elegans

MTKYCRAPRCSNSAGQPRRDQRRLSFYKFPLHSPERLRQWLSQMNQEKWIPTKHQHLCSDHFAPSCFEYR

WGVRYLKPDAVPTIF

>032851315.1|Tyto albaalba

MRRENWVPTRHQHLCSDHFEPSCFQYRWGVRYLRPDAVPTIF

>030330852.1|Strigops habroptila

MPKSCRAPRCSNAAGQARTAARGVSFYRFPLQDAPRLRQWLERMGQENWVPTRHQHLCSDHFEPSCFQYR

WGVRYLRPDAVPTIF

>029862657.1|Aquila chrysaetoschrysaetos

MPKYCRAPHCSNAAGQARPPARRLSFYKFPLHDAARLRQWLTQMQRENWVPTRHQHLCSDHFEPSCFQYR

WGVRYLRPDAIPTIF

>028920420.1|Ornithorhynchus anatinus

MPKYCRAPNCSNTAGQVSADNRRISFYKFPLQDPARLQEWLQQMRQEQWVPTRHQHLCSEHFAPSCFEWR

WGVRYLKPDAVPTIF

>004464942.1|Dasypus novemcinctus

MPKSCWAPNCSNTAGRLGADNRPVSFYKFPLKNGPRLQAWLRHMGREHWVPSCHQHLCSEHFTPASFQWR

WGVRYLRPDAVPSIF

>006171744.1|Tupaia chinensis

MPKYCRAPNCSNTAGRLGSDNRPVSFYKFPLKDGPRLQAWLQHMGREHWVPSCHQHLCSEHFTPSCFQWR

WGVRYLRPDAVPSIF

>023505376.1|Equus caballus

MPKYCRAPNCSNTAGRLGADNRPVSFYKFPLKDGPRLQAWLRHMGREHWVPSCHQHLCSEHFTPSCFQWR

WGVRYLRPDAVPSIF

>032984426.1|Rhinolophus ferrumequinum

MPKYCWAPNCSNTAGRLGADNRPVSFYKFPLKDGPRLQAWLRHMGCEHWVPSCHQHLCSEHFTPSCFQWR

WGVRYLRPDAVPSIF

>024896394.1|Pteropus alecto

MLKHCCAPNCSNTASRLGADNRPVSFYKFPLKVGLQLQAWLRHMGCEHWVPSCHQHLCSEHFTPSCFQCR

CGVRYLRLDAVPSIF

>012932901.1|Heterocephalus glaber

MPKYCRAPNCSNTAGRLGSDNRPVSFYKFPLKDGSRLQSWLRHMGCEHWVPSCHQHLCSEHFTPSCFQWR

WGVRYLRPDAVPSIF

>013221462.1|Ictidomys tridecemlineatus

MPKYCRAPNCSNTAGQLGSDNRPVSFYKFPLKDGPRLEAWLRCMGREHWVPSCHQHLCSEHFTPSCFQWR

WGVRYLRPDAVPSIF

>026235875.1|Urocitellus parryii

MPKYCRAPNCSNTAGQLGSDNRPVSFYKFPLKDGPRLEAWLRCMGREHWVPSCHQHLCSEHFTPSCFQWR

WGVRYLRPDAVPSIF

>026642108.1|Microtus ochrogaster

MGHEHWVPSCHCCLCCLCSERFTPSCFQGRWGVGHLRPDVLLSIF

>027797132.1|Marmota flaviventris

MPKYCRAPNCSNTAGQLGSDNRPVSFYKFPLKDSPRLQAWLRCMGREHWVPSCHQHLCSEHFTPSCFQWR

WGVRYLRPDAVPSIF

>JAV35979.1|Castor canadensis

MPKYCRAPNCSNTAGRLGADNRPVSFYKFPLKDGPRLQAWLRRMGREHWVPSCHQHLCSEHFTPSCFQWR

WGVRYLRPDAVPSIF

>021556653.1|Neomonachus schauinslandi

MRHKGPSRESGGERRGPKYDCRLGRTQWGRTAMHKYCRAPNCSNTSGRLGADNRPVSFYNMQLPCKMAAK

CPSHHHLVPTEPAQGHGIRDWLRHMGHEDWVPSCRQHLCSEHFTPSCFQWRRAVRYLRPDAVLSIF

>005616733.1|Canis lupus familiaris

MPKYCRAPNCSNTAGRLGADNRPVSFYNNNEASRLAPGRVVALTQGSNLSCSAQVQSFRFPLKDGPRLQA

WLRHMGHEDWVPSCHHHLCSEHFTPSCFQWRWGVRYLRPDAVPSIF

>006941499.1|Felis catus

MPKYCRAPNCSNTAGRLGADNRPVSFYKFPLKDGPRLRAWLRHMGHEDWVPSCHQHLCSEHFTPSCFQWR

WGVRYLRPDAVPSIF

>025277545.1|Canis lupus dingo

MPKYCRAPNCSNTAGRLGADNRPVSFYNNNEASRLAPGRVVALTQGSNLSCSAQVQSFRFPLKDGPRLQA

WLRHMGHEDWVPSCHHHLCSEHFTPSCFQWRWGVRYLRPDAVPSIF

>025770592.1|Puma concolor

MPKYCRAPNCSNTAGRLGADNRPVSFYKFPLKDGPRLRAWLRHMGHEDWVPSCHQHLCSEHFTPSCFQWR

WGVRYLRPDAVPSIF

>025866143.1|Vulpes vulpes

MPKYCRAPNCSNTAGRLGADNRPVSFYKFPLKDGPRLQAWLRHMGHEDWVPSCHHHLCSEHFTPSCFQWR

WGVRYLRPDAVPSIF

>026340289.1|Ursus arctoshorribilis

MSKYCRAPNCSNTAGRLGADNRPLSFYQFPLKDGPRLRAWLQHMGHEDWVPSCHQHLCSEHFTPSCFQWR

WGVRYLRPDAVPSIF

>030154696.1|Lynx canadensis

MPKYCRAPNCSNTAGRLGADNRPVSFYKFPLKDGPRLRAWLRHMGHEDWVPSCHQHLCSEHFTPSCFQWR

WGVRYLRPDAVPSIF

>032179641.1|Mustela erminea

MPKYCRAPNCSNTAGRLGADNRPVSFYKFPLKDGPRLQAWLRHMGHEDWEPSCHQHLCSEHFTPSCFQWR

WGVRYLRPDAVPSIF

>032697815.1|Lontra canadensis

MPKYCRAPNCSNTAGRLGADNRPVSFYKFPLKDGPRLQAWLRHMGHEDWEPSCHQHLCSEHFTPSCFQWR

WGVRYLRPDAVPSIF

>030730632.1|Globicephala melas

MGCEHWVPSCHQHLCSEHFAPSCFQWRWGVRYLRPDAVPSIF

>006217158.1|Vicugna pacos

MPKYCRAPNCSNTAGQLGADNRPVSFYKFPLKDGPRLQAWLRHMGREHWVPSCHQHLCSEHFAPSCFQWR

WGVRYLRPDAVPSIF

>032344007.1|Camelus ferus

MPKYCRAPNCSNTAGQLGADNRPVSFYKFPLKDGPRLQAWLRHMGREHWVPSCHQHLCSEHFAPSCFQWR

WGVRYLRPDAVPSIF

>032470532.1|Phocoena sinus

MPKYCRPPNCSNNAGQLGADNRPVSFYKFPLKDGPRLQAWLRHMGCEHWVPSCHQHLCSEHFAPSCFQWR

WGVRYLRPDAVPSIF

>004284258.1|Orcinus orca

MPKYCRAPNCSNNAGQLGADNRPVSFYKFPLKDGPRLQAWLRHMGCEHWVPSCHQHLCSEHFAPSCFQWR

WGVRYLRPDAVPSIF

>020727363.1|Odocoileus virginianustexanus

MARNTQAQLSFPPHVTRTQGPSREAGDKRAGGAGFGGAWRGSKCGCRPGKTQLGWTAMPKYCRAPNCSNT

AGQLGADNRPVSFYKFPLKDGPRLQAWLRHMGREHWVPSCHQHLCSEHFAPSCFQWRWGVRYLRPDAVPS

IF

>013853822.2|Sus scrofa

MPRYCRAPNCSNTAGRLGADHRPYLPWCAVSLLPRFPLKDGPRLQAWLRHMGLEHWVPSCHQHLCSEHFA

PSCFQWRWGVRYLRPDAVPSIF

>024589724.1|Neophocaena asiaeorientalisasiaeorientalis

MPKYCRAPNCSNNAGQLGADNRPVSFYKFPLKDGPRLQAWLRHMGCEHWVPSCHQHLCSEHFAPSCFQWR

WGVRYLRPDAVPSIF

>010813077.1|Bos taurus

MAKEPRKPSFNRAHYASKKLEFVSKEEAESGYGIILWQSLPRSRFPLKDGPRLQAWLRHMGREHWVPSCH

QHLCSEHFAPSCFQWRWGVRYLRPDAVPSIF

>026934847.1|Lagenorhynchus obliquidens

MPKYCRAPNCSNNAGQLGADNRPVSFYKFPLKDGPRLQAWLRHMGCEHWVPSCHQHLCSEHFAPSCFQWR

WGVRYLRPDAVPSIF

>004015258.2|Ovis aries

MPKYCRAPNCSNTAGQLGADNRPVSFYKFPLKDGPRLQAWLRHMGREHWVPSCHQHLCSEHFAPSCFQWR

WGVRYLRPDAVPSIF

>007165244.1|Balaenoptera acutorostratascammoni

MPKYCRAPNCSNNAGQLGADNRPVSFYKFPLKDGPRLQAWLRHMGREHWVPSCYQHLCSEHFAPSCFQWR

WGVRYLRPDAVPSIF

>028340542.1|Physeter catodon

MPKYCRAPNCSNNAGQLGADNRPVSFYKFPLKDGPRLQAWLRHMGREHWVPSCHQHLCSEHFAPSCFQWR

WGVRYLRPDAVPSVF

>029065340.1|Monodon monoceros

MPKYCRAPNCSNNAGQLGADNRPVSFYKFPLKDGPRLQAWLRHMGCEHWVPSCHQHLCSEHFAPSCFQWR

WGVRYLRPDAVPSIF

>030730626.1|Globicephala melas

MPEYCRAPNCSNNAGQLGADNRPVSFYKFPLKDGPRLQAWLRHMGCEHWVPSCHQHLCSEHFAPSCFQWR

WGVRYLRPDAVPSIF

>PNI95866.1|Pan troglodytes

MPKYCRAPNCSNTAGRLGADNRPVSFYKFPLKDGPRLQAWLQHMGREHWVPSCHQHLCSEHFTPSCFQWR

WGVRYLRPDAVPSIF

>PNJ11281.1|Pongo abelii

MPKYCRAPNCSNTAGRLGADNRPVSFYKFPLKDGPRLQAWLRHMGREHWVPSCHQHLCSEHFTPSCFQWR

WGVRYLRPDAVPSIF

>JAB02532.1|Callithrix jacchus

MPKYCRAPNCSNTAGRLGADNRPVSFYKFPLKDGPRLQAWLQHMGREHWVPSCHQHLCSEHFTPSCFQWR

WGVRYLRPDAVPSIF

>AFJ70865.1|Macaca mulatta

MPKYCRAPNCSNTAGSLGADNRPVSFYKFPLKDGPRLQAWLRHMGREHWVPSCHQHLCSEHFTPSCFQWR

WGVRYLRPDAVPSIF

>012630410.1|Microcebus murinus

MPKYCRAPNCSNTAGRLGADNRPVSFYKFPLKDGPRLQAWLRCMGREHWVPSCHQHLCSEHFTPSCFQWR

WGVRYLRPDAVPSIF

>012292917.2|Aotus nancymaae

MPKYCRAPNCSNTAGRLGADNRPVSFYKFPLKDGPRLQAWLQHMGREHWVPSCHQHLCSEHFTPSCFQWR

WGVRYLRPDAVPSIF

>008071829.2|Carlito syrichta

MPKYCRAPNCSNTAGRLGADNRPVSFYKFPLKDGPRLQAWLRHMGREHWVPSCHQHLCSEHFTPSCFQWR

WGVRYLRPDAVPSIF

>023372674.1|Otolemur garnettii

MPKYCRAPNCSNTAGRLGADNRPVSFYKFPLKDGPRLQAWLQRMGREHWVPSYHQHLCSEHFAPSCFQWR

WGVRYLRPDAVPSIF

>011763406.1|Macaca nemestrina

MPKYCRAPNCSNTAGSLGADNRPVSFYKFPLKDGPRLQAWLRHMGREHWVPSCHQHLCSEHFTPSCFQWR

WGVRYLRPDAVPSIF

>003816196.1|Pan paniscus

MPKYCRAPNCSNTAGRLGADNRPVSFYKFPLKDGPRLQAWLQHMGREHWVPSCHQHLCSEHFTPSCFQWR

WGVRYLRPDAVPSIF

>025224356.1|Theropithecus gelada

MPKYCRAPNCSNTAGSLGADNRPVSFYKFPLKDGPRLQAWLRHMGREHWVPSCHQHLCSEHFTPSCFQWR

WGVRYLRPDAVPSIF

>003280121.1|Nomascus leucogenys

MPKYCRAPNCSNTAGRLGADNRPVSFYKFPLKDGPRLQAWLRHMGREHWVPSCHQHLCSEHFTPSCFQWR

WGVRYLRPDAVPSIF

>010367100.1|Rhinopithecus roxellana

MPKYCRAPNCSNTAGRLGADNRPVSFYKFPLKDGPRLQAWLRHMGREHWVPNCHQHLCSEHFTPSCFQWR

WGVRYLRPDAVPSIF

>004060609.2|Gorilla gorilla gorilla

MPKYCRAPNCSNTAGRLGADNRPVSFYKFPLKDGPRLQAWLQHMGREHWVPSCHQHLCSEHFTSSCFQWR

WGVRYLRPDAVPSIF

>031515468.1|Papio anubis

MPKYCTAPNCSNTAGSLGADNRPVSFYKFPLKDGPRLQAWLRHMGREHWVPSCHQHLCSEHFTPSCFQWR

WGVRYLRPDAVPSIF

>026304540.1|Piliocolobus tephrosceles

MPKYCRAPNCSNTAGRLGADNRPVSFYKFPLKDGPRLQAWLRHMGREHWVPNCHQHLCSEHFTPSCFQWR

WGVRYLRPDAVPSIF

>032105416.1|Sapajus apella

MPKYCRAPNCSNTAGRLGADNRPVSFYKFPLKDGPRLQAWLQHMGREHWVPSCHQHLCSEHFTPSCFQWR

WGVRYLRPDAVPSIF

>032026497.1|Hylobates moloch

MPKYCRAPNCSNTAGRLGADNRPVSFYKFPLKDGPRLQAWLRHMGREHWVPSCHQHLCSEHFAPSCFQWR

WGVRYLRPDAVPSIF

>Q8NA92.1|Homo Sapiens

MPKYCRAPNCSNTAGRLGADNRPVSFYKFPLKDGPRLQAWLQHMGCEHWVPSCHQHLCSEHFTPSCFQWRWGVRYLRPDAVPSIF
