## Additional file for "Synapomorphic variations in the THAP domains of the human THAP protein family and its homologs": THAP9.docx

>KXJ06154.1|Exaiptasia pallida

MTATKQELEAMNAQLIDSLTTSFNKLSHELLQHREQSHTDLAAVTSAMSAEKKSDSAKTPGAIGKLPTFS

GKDSDASEFLQDFCFYADFLNWTDQERVRAFPLALQGNARTWFTNMTAKYTTFEELSKAFSDHFLSREND

WMLRHNLSSRHQLPSETKDDEHKRWRDEWLGELKKTRETDKDFRRQINDDKVYTCEKHFYPEDIEIFHSE

KMTKKKLRFGAIPTLN

>ACO14675.1|Caligus clemensi

MDSGKSRRKTCAVSTCSSPHVDYGYSYHRFPKRSNVRDQWIKACRRIDKINPDTATVCSKHFEQECFEQN

LKAELMANYGRTASGKVRKTLKANAVPTLH

>CDW36669.1|Myotis lucifugus

MVNSCSAVGCKNRGVLTTFQQGITFHRFPKNSELKSEWISALRRKNFTPSASAVLCSVHFEETDFKSHTT

RRVLKDGAIPSII

>OXA63734.1|Folsomia candida

MVFVIFVSNNRYNRYLKGVNLIRPGGLLRTRHKFTKISTSTRQEAPTAAAKHVSCQGVILVTGYLLEYQC

SAPENLQKWLDFVRSLVNDENWVNSNSSTVCSLHFEEGSRSVGSKSRLVHNAFPTYC

>GBP36459.1|Eumeta japonica

MLLHFIQYPIPAQEVGNASVTPLDCPWTTVTSSGRLHARLPLDNVIRKIRYQYNRSRPTTVPTAYPLLLA

LLPVVLQCHLWIEACRRPDLRDKNVDQLYNMHVCGLHFEGWMYMKKKLKSAAIPVLN

>KYO21751.1|Alligator mississippiensis

MPKACSAINCPNRDTRESRARGLSFHSFPKAHELRKRWLLAVRRVEPGSRRLWVPGAGACLCSQHFAREE

FELHGGQRRLKAGVIPSLF

>025027499.1|Python bivittatus

MPKACSAINCPNRDTRQNRAKGLSFHSFPKDHELRKKWMLAVNRVEPGTKKLWIPGSGACLCSQHFRQEE

FEIYGGQKRLKVGVIPSLF

>029471725.1|Rhinatrema bivittatum

MPVSCAAFGCKSRYTLEAREKGITFHRFPKSNPILLEKWRIAVERATSTGELWMPSRYQRLCSLHFEEKC

FDTTGQTKRLRDDVIPSIF

>030072182.1|Microcaecilia unicolor

MPVSCAAYGCKSRYTLEAREKGITFHRFPKSNPALLEKWRIAVERATSTGELWMPSRYQRLCSLHFEEKC

FDTTGQTKRLRDDVIPSIF

>004913090.2|Xenopus tropicalis

MPVSCAASGCKSRYTLDAREKGITFHRFPRSNPALLEKWRLAMRRSTRNGELWMPSRYQRLCSLHFKQCC

FDTTGQTKRLREHVIPTIF

>008163880.1|Chrysemys pictabellii

MTRSCSALGCTTRDTGLSRKRGISFHQFPIDDIQRTKWIHAVNRADPKSKKVWIPGPGAILCSRHFAEAD

FESYGMRRKLKKGAVPSVF

>027683756.1|Chelonia mydas

MTRSCSALGCTTRDTGLSRKRGISFHQFPIDDIQRTKWIHAVNRADPKSKKVWIPGPGAILCSRHFAEAD

FESYGMRRKLRKGAVPSVF

>024068400.1|Terrapene carolinatriunguis

MTRSCSALGCTTRDTGLSRKRGISFHQFPIDDIQRTKWIHAVNRADPKSKKVWIPGPGAILCSRHFAEAD

FESYGMRRKLRKGAVPSVF

>030420885.1|Gopherus evgoodei

MTRSCSALGCTTRDTGLSRKRGISFHQFPIDDIQRTKWIHAVNRADPKSKKVWIPGPGAILCSRHFAEAD

FESYGMRRKLRKGAVPSVF

>032641596.1|Chelonoidis abingdonii

MTRSCSALGCTTRDTGLSRKRGISFHQCRGWGGETGKHMGLLRKRSQAFQFSVIGRFPIDDIQRTKWIHA

VNRADPKSKKIWIPGPGAILCSRHFAEADFESYGMRRKLRKGAVPSVF

>ROL50904.1|Anabarilius grahami

MCSKRQVSSSELSFFRFPFDDKDRLKQWIHYVKRKNWRPKKYSRLCSLHFTQDCFIPVQGRNILKADAVP

TIF

>RXN27901.1|Labeo rohita

MVQNGFPKSRERRRKWEVALRRDGFAATDRTLICSEHFRTEDFDRKGQTVRIKDGVVPSIF

>020774187.1|Boleophthalmus pectinirostris

MPSCSAFNCTNRPDGLQKITFFRFPLGDKKRLAQWVEKVRRKNWVPTANSRLCAAHFEEECFLIKGRKTY

LQPSAVPTIF

>023185528.1|Xiphophorus maculatus

MPQFCSAYSCLNLRTVDVRDRGITFHKFPKDKERRKRWEIALRRDGFTASDSSVLCSEHFKTEDFDKTGQ

IVRLRADVMPSIF

>032446710.1|Xiphophorus hellerii

MPQFCSAYSCLNLRTVDVRDRGITFHKFPKDKERRKRWEIALRRDGFTASDSSVLCSEHFKTEDFDKTGQ

IVRLRADVIPSIF

>KQK80998.1|Amazona aestiva

MTRSCSALGCTARDNGRSRERGISFHQFPVDAAQRREWIRAVNRVDPQSRQAWRPGPGAILCSRHFAEAD

FERYGLRRKLRRGAVPSRF

>OPJ75967.1|Patagioenas fasciatamonilis

MTRSCSALGCTARDTGRSRERGISFHQFPVDAAQRREWIRAVNRVDPRSRQAWRPGPGAILCSRHFAETD

FERYGLRRKLRRGAVPSRF

>OWK60383.1|Lonchura striatadomestica

MTRSCSALGCTARDTGRSRERGISFHQFPVDAAQRREWIRAVNRVDPRSRRAWRPGPGAILCSRHFAEGD

FERYGLRRKLRRGAVPSRF

>420555.4|Gallus gallus

MTRSCSALGCSARDNGRSRERGISFHQFPVDAAQRREWIRAVNRLDPRSRRAWSPGPGAILCSRHFAEAD

FERYGLRRKLRRGAVPSRF

>025897091.1|Nothoprocta perdicaria

MTRSCSALGCTARDNGRSRERGISFHQFPVDAAQRRAWIRAVNRVDPRSRQAWSPGPGAILCSRHFAEGD

FERYGLRRKLRRGAVPSRF

>025924266.1|Apteryx rowi

MRRFPVDAAQRREWIRAVNRVDPRSRQAWSPGPGAILCSRHFAEADFERYGLRRKLRRGAVPSRF

>025958537.1|Dromaius novaehollandiae

MTRSCSALGCTARDNGRSRERGISFHQFPVDAAQRREWIRAVNRVDPRSRQAWSPGPGAILCSRHFAEAD

FERYGLRRKLRRGAVPSRF

>027548604.1|Neopelma chrysocephalum

MTRSCSALGCTARDSARSRERGVSFHQFPVDAAQRRAWIRAVNRVDPQSRRPWRPGPGAILCSRHFAEGD

FERFGLRRKLRRGAVPSRF

>027508509.1|Corapipo altera

MTRSCSALGCTARDSARSRERGVSFHQFPVDAAQRRAWIRAVNRVDPQSRRPWRPGPGAILCSRHFAEGD

FERFGLRRKLRRGAVPSRF

>027594338.1|Pipra filicauda

MTRSCSALGCTARDTARSRERGVSFHQFPVDAAQRRAWIRAVNRVDPQSRRPWRPGPGAILCSRHFAEGD

FERFGLRRKLRRGAVPSRF

>027743021.1|Empidonax traillii

MTRSCSALGCTARDSARSRERGVSFHQFPVDAVQRSAWIRAVNRVDPQSRRPWLPGPGAILCSRHFAEGD

FERFGLRRKLRRGAVPSRF

>029871779.1|Aquila chrysaetoschrysaetos

MGRRTLGGRAVAANGRARSVPGWGVAAGCGRGGGAAMTRSCSALGCTARDNGRSRERGISFHQFPVDAAQ

RREWIRAVNRVDPRSRQAWRPGPGAILCSRHFAEADFERYGLRRKLRRGAVPSRF

>030095319.1|Serinus canaria

MTRSCSALGCTARDTGRSRERGISFHQFPVDAAQRREWIRAVNRVDPRSRRAWRPGPGAILCSRHFAEGD

FERYGLRRKLRRGAVPSRF

>030306697.1|Calypte anna

MTRSCSALGCTARDNGRSRERGISFHQFPVDAAQRREWIRAVNRVDPRSHQPWRPGPGAILCSRHFAKAD

FERYGLRRKLRRGAVPSRF

>031452544.1|Phasianus colchicus

MTRSCSALGCSARDNGRSRERGISFHQFPVDAAQRREWIRAVNRLDPRSRRAWSPGPGAILCSRHFAEAD

FERYGLRRKLRRGAVPSRF

>031963888.1|Corvus moneduloides

MTRSCSALGCTARDTGRSRERGISFHQFPADAAQRRQWIRAVNRVDPRSHRAWRPGPGAILCSRHFAEGD

FERYGLRRKLRRGAVPSRF

>015717046.1|Coturnix japonica

MQRARQRAEPGAGHLLPPVSDRAAGGTAPYTALPPDPPHSALPTPFGPTQPYPPYRPHSALPTPFRPTQP

FPPYRRFPVDAAQRLQWIRAVNRVDPRSRRLWLPGPGAILCSRHFAEADFERYGLRRKLRRGAVPSRF

>032541146.1|Chiroxiphia lanceolata

MTRSCSALGCTARDSARSRERGVSFHQFPVDAAQRRAWIRAVNRVDPQSRRPWRPGPGAILCSRHFAEGD

FERFGLRRKLRRGAVPSRF

>012428772.3|Taeniopygia guttata

MTRSCSALGCTARDTGRSRERGISFHQFPVDAAQRREWIRAVNRVDPRSRRAWRPGPGAILCSRHFAEGD

FERYGLRRKLRRGAVPSRF

>032915883.1|Catharus ustulatus

MTRSCSALGCTARDTGRSRERGISFHQFPVDAAQRREWIRAVNRVDPQSRRAWRPGPGAILCSRHFAEGD

FERYGLRRKLRRGAVPSRF

>030347194.1|Strigops habroptila

MGGRCRCRAAAAMTRSCSALGCTARDNGRSRERGISFHQFPVDAAQRREWIRAVNRVDPQSRQAWRPGPG

AILCSRHFAEADFERYGLRRKLRRGAVPSRF

>004383150.1|Trichechus manatuslatirostris

MTRSCSAVGCSTRDTVLSRERGVSFHQFPTDTIQRSKWIRAVNRVDPRSKKIWIPGPGAILCSRHFQESD

FESYGMRRKLKKGAVPSVS

>021116621.1|Heterocephalus glaber

MTRSCSAVGCSTRDTVLSRERGLSFHQCVFPTDTIQRSKWIRAVNRVDPRSKKIWIPGPGAILCSRHFQK

CDFESYGVRRKLKKGAVPSLS

>005339375.2|Ictidomys tridecemlineatus

MTRSCSAVGCSTRDTVLSRERGLSFHQFPADTIQRSKWIRAVNRVDPRSKKIWIPGPGAILCSKHFQESD

FESYGIRRKLKKGAVPSVS

>023438617.1|Dasypus novemcinctus

MVYDEQFVTEHQSMATYAKPICSEARRSRTGPKRRPRALCPRDRASGRRGPVGTEARVPALGSDSTPRGP

ETREKRLEADSPGGDPLRPVPALPRRSSDSPRIRRRPSSPVRGHLHVPFQRGSRSPARPAGASGSDAAPL

PPRRAACHSTSGSGDRPRSRVDSAALVAVTSTLGKFLWGRARSADVLGQDHRKRKDPARCCFKPIMTAGA

RTPRSTSSYYCSRFPTDTIQRAKWIRAVNRVDPRSKKIWIPGPGAILCSKHFQESDFESYGIRRKLKKGA

VPSVS

>011354654.1|Pteropus vampyrus

MTRSCSAVGCSTRDTVQSRKRGYSFHQFPTDTIQRSKWIRAVNRVDPRSKKIWIPGPSAMLCSKHFQESD

FESYGMRRKLKKGAVPSVS

>001915516.1|Equus caballus

MTRSCSAVGCSTRDTVLSRERGLSFHQFPTDTIQRSKWIRAVNRVDPRSKKIWIPGPGAILCSKHFQESD

FESYGIRRKLKKGAVPSVS

>004640377.1|Octodon degus

MTRSCSTVGCSTRDTGQSREHGLSFHQFPTDTVQRSKWIRAVNRVDPRSKKIWIPGPGAILCSKHFQESD

FESYGVRRKLKKGTVPSLS

>006084617.2|Myotis lucifugus

MTRSCSAVGCSTKDTLESRKQGYSFHQFPTDTILRSKWIRAVNRVDPISKKIWIPGPSAMLCSKHFLESD

FESYGLRRMLKKGAVPSVS

>006907655.1|Pteropus alecto

MTRSCSAVGCSTRDTVQSRKRGYSFHQFPTDTIQRSKWIRAVNRVDPRSKKIWIPGPSAMLCSKHFQESD

FESYGMRRKLKKGAVPSVS

>026264803.1|Urocitellus parryii

MTRSCSAVGCSTRDTVLSRERGLSFHQFPADTIQRSKWIRAVNRVDPRSKKIWIPGPGAILCSKHFQESD

FESYGIRRKLKKGAVPSVS

>027628850.1|Tupaia chinensis

MSVFAFALHGNLLRLQLYFHGYLQHHHQTLKRSGKNVSKGLWCKTNQSDSKLWRLNRENIGALSETFMAK

CEVGLARFPTDTILRSKWISAVNRVDPRSKKIWIPGPGAILCSKHFRESDFESYGIRRKLKKGAVPSVS

>027795633.1|Marmota flaviventris

MTRSCSAVGCSTRDTMLSRERGLSFHQFPTDTIQRSKWIRAVNRVDPRSKKIWIPGPGAILCSKHFQESD

FESYGIRRKLKKGAVPSVS

>028004170.1|Eptesicus fuscus

MARSCSAVGCSTRDTLQSRKHGYSFHSFPTDTILRSKWIRAVNRVDPRSKKIWIPGPSAMLCSKHFLESD

FESYGLRRMLKKGAVPSVS

>010627249.1|Fukomys damarensis

MTRSCSAVGCSTRDTVLSRERGLSFHQFPTDTIQRSKWIRAVNRVDPRSKKIWIPGPGAILCSRHFQKSD

FESYGVRRKLKKGAVPSLS

>078948.3|Homo sapiens

MTRSCSAVGCSTRDTVLSRERGLSFHQFPTDTIQRSKWIRAVNRVDPRSKKIWIPGPGAILCSKHFQESD

FESYGIRRKLKKGAVPSVS
