## Additional file for "Synapomorphic variations in the THAP domains of the human THAP protein family and its homologs": THAP10.docx

>JAG74292.1|Fopius arisanus

MPKICVVSNCGTGTKGVEEKRSLFRVPSNKELRLKWEAAIPGIKILKLGQSVCERHFPAKYINKKWIKYD

SRGRVIAQYDFPRPRLDPRAIPCIF

>KXJ24841.1|Exaiptasia pallida

MPNRCVAGGCSNVPDTSRNIALYKFPKDDQQNKKRRRLXINFVLTKRSQWSPTDSSRLCSEHFTSEDFET

TFVAIPGTSFVQRLSLKKDAVPSIH

>017276780.1|Kryptolebias marmoratus

MPNYCVVGYCNNVANRSLGISTYAFPKDGAIRRAWIRFVQQTRADFREPGSGAVVCSSHLKEDCFEIPCP

SVRSCEEETGFRNRVIKKRKANAVPTIG

>020837702.1|Phascolarctos cinereus

MPARCVAAHCGNTTKSGKSLFRFPKDRAVRLLWDRFVRGHRADWYGGNDRSVICSDHFAPCCFDVSSVIQ

KNLQFSQRLRLVAGAVPTLH

>027707807.1|Vombatus ursinus

MPARCVAAHCGNTTKSGKSLFRFPKDRAVRLLWDRFVRGHRADWYGGNDRSVICSDHFAPCCFDVSSVIQ

KNLHFSQRLRLVAGAVPTLH

>031811156.1|Sarcophilus harrisii

MPARCVAAHCGNTTKSGKSLFRFPKDRAVRLLWDRFVRGRRADWYGGNDRSVICSDHFAPCCFDVSSVIQ

KNLHFSQRLRLVAGAVPTLH

>021549430.1|Neomonachus schauinslandi

MPARCVAAHCGNTTKSGKSLFRFPKDRAVRLLWDRFVRGRRADWYGGNDRSVICSDHFAPACFDVSSVIQ

KNLRFSQRLRLVAGAVPTLH

>853894.1|Canis lupus familiaris

MPARCVAAHCGNTTKCGKSLFRFPKDRAVRLLWDRFVRGRRADWYGGNDRSVICSDHFAPACFDVSSVLQ

KNLRFSQRLRLVAGAVPTLH

>023110870.1|Felis catus

MPARCVAAHCGNTTKSGKSLFRFPKDRAVRLLWDRFVRGRRADWYGGNDRSVICSDHFAPACFDVSSVIQ

KNLRFSQRLRLVAGAVPTLH

>025328087.1|Canis lupus dingo

MPARCVAAHCGNTTKCGKSLFRFPKDRAVRLLWDRFVRGRRADWYGGNDRSVICSDHFAPACFDVSSVLQ

KNLRFSQRLRLVAGAVPTLH

>025740257.1|Callorhinus ursinus

MPARCVAAHCGNTTKSGKSLFRFPKDRAVRLLWDRFVRGRRADWYGGNDRSVICSDHFAPACFDVSSVIQ

KNLRFSQRLRLVAGAVPTLH

>025775767.1|Puma concolor

MPARCVAAHCGNTTKSGKSLFRFPKDRAVRLLWDRFVRGRRADWYGGNDRSVICSDHFAPACFDVSSVIQ

KNLRFSQRLRLVAGAVPTLH

>025851818.1|Vulpes vulpes

MPARCVAAHCGNTTKCGKSLFRFPKDRAVRLLWDRFVRGRRADWYGGNDRSVICSDHFAPACFDASSVLQ

KNLRFSQRLRLVAGAVPTLH

>026334217.1|Ursus arctoshorribilis

MPARCVAAHCGNTTKAGKSLFRFPKDRAVRLLWDRFVRGRRADWYGGNDRSVICSDHFAPACFDVSSVIQ

KNLRFSQRLRLVAGAVPTLH

>026923471.1|Acinonyx jubatus

MPARCVAAHCGNTTKSGKSLFRFPKDRAVRLLWDRFVRGRRADWYGGNDRSVICSDHFAPACFDVSSVIQ

KNLRFSQRLRLVAGAVPTLH

>027427343.1|Zalophus californianus

MPARCVAAHCGNTTKSGKSLFRFPKDRAVRLLWDRFVRGRRADWYGGNDRSVICSDHFAPACFDVSSVIQ

KNLRFSQRLRLVAGAVPTLH

>027956299.1|Eumetopias jubatus

MPARCVAAHCGNTTKSGKSLFRFPKDRAVRLLWDRFVRGRRADWYGGNDRSVICSDHFAPACFDVSSVIQ

KNLRFSQRLRLVAGAVPTLH

>029808125.1|Suricata suricatta

METTISGVVHAERECGQEGAGKRPATSVAVATTTSGPSLFPAQLISARAKFGSLCPRPPALGEGLARAGR

GRYPGPASPSVGPQLGTSGGPRRAGRRRGPASPWAGKRGVRRYGGNDRSVICSDHFAPACFDVSSVIQKN

LRFSQRLRLVAGAVPTLH

>030173654.1|Lynx canadensis

MPARCVAAHCGNTTKSGKSLFRFPKDRAVRLLWDRFVRGRRADWYGGNDRSVICSDHFAPACFDVSSVIQ

KNLRFSQRLRLVAGAVPTLH

>006743336.1|Leptonychotes weddellii

MPARCVAAHCGNTTKSGKSLFRFPKDRAVRLLWDRFVRGRRADWYGGNDRSVICSDHFAPACFDVSSVIQ

KNLRFSQRLRLVAGAVPTLH

>032198497.1|Mustela erminea

MPARCVAAHCGNTTKSGKSLFRFPKDRAVRLLWDRFVRGRRADWYGGNDRSVICSDHFAPACFDVSSVIQ

KNLRFSQRLRLVAGAVPTLH

>032273979.1|Phoca vitulina

MPARCVAAHCGNTTKSGKSLFRFPKDRAVRLLWDRFVRGRRADWYGGNDRSVICSDHFAPACFDVSSVIQ

KNLRFSQRLRLVAGAVPTLH

>032720816.1|Lontra canadensis

MPARCVAAHCGNTTKSGKSLFRFPKDRAVRLLWDRFVRGRRADWYGGNDRSVICSDHFAPACFDVSSVIQ

KNLRFSQRLRLVAGAVPTLH

>001074210.1|Bos taurus

MRCGNTTKSGKLLFRFPKGPGRATAVGPLRAGPPANWYRGSDSSVICSEHFAPACFDVSSVIQKNLPFSE

RLRPVAGAASILH

>025116329.1|Bubalus bubalis

MTTTTSGPSLFTSHLSWARAKFVSLCLLSGEDCWKGKDLQKPGTLQGCRSPAPSPEAAAVPIRCGNTTKS

GKLLFRFPKGPGRAAAVGPLRAGPPANWYRGSDSSVICSEHFAPACFDVSSVIQKNLPFSERLRPVAGAA

SILH

>007197844.1|Balaenoptera acutorostratascammoni

MPARCVAAHCGNTTTSGNSLFRFPKDRAVRLLWGRFVRSRRADWYEGNDISVICSDHFATACFDVSSVIQ

KNLRFSQRLRFVAGAVPTLH

>030730953.1|Globicephala melas

MPASCVAAHCGNTTTSGKSLFRFPKDRAVRLLWDRFVRGRRADWYEGNDGSVICSDHFAPACFDVSSVIR

KNLRFSQRLRLVAGAVPTLH

>PNJ43008.1|Pongo abelii

MPARCVAAHCGNTTKSGKSLFRFPKDRAVRLLWDRFVRGCRADWYGGNDRSVICSDHFAPACFDVSSVIQ

KNLRFSQRLRLVAGAVPTLH

>AFE79282.1|Macaca mulatta

MPARCVAAHCGNTTKSGKSLFRFPKDRAVRLLWDRFVRGCRADWYGGNDRSVICSDHFAPACFDVSSVIQ

KNLRFSQRLRLVAGAVPTLH

>PNI74114.1|Pan troglodytes

MPARCVAAHCGNTTKSGKSLFRFPKDRAVRLLWDRFVRGCRADWYGGNDRSVICSDHFAPACFDVSSVIQ

KNLRFSQRLRLVAGAVPTLH

>012623767.1|Microcebus murinus

MPARCVAAHCGNTTKSGKSLFRFPKDRAVRLLWDRFVQGFRADWYGGNDRSVICSDHFAPACFDVSSVIQ

KKLRFSQRLRLVAGAVPTLH

>012306735.1|Aotus nancymaae

MPARCVAAHCGNTTKSGKSLFRFPKDRAVRLLWDRFVRGCRADWYGGNDRSVICSDHFAPACFDVSSVIQ

KNLHFSQRLRLVAGAVPTLH

>011749438.1|Macaca nemestrina

MPARCVAAHCGNTTKSGKSLFRFPKDRAVRLLWDRFVRGCRADWYGGNDRSVICSDHFAPACFDVSSVIQ

KNLRFSQRLRLVAGAVPTLH

>003818590.1|Pan paniscus

MPARCVAAHCGNTTKSGKSLFRFPKDRAVRLLWDRFVRGCRADWYGGNDRSVICSDHFAPACFDVSSVIQ

KNLRFSQRLRLVAGAVPTLH

>025245731.1|Theropithecus gelada

MPARCVAAHCGNTTKSGKSLFRFPKDRAVRLLWDRFVRGCRADWYGGNDRSVICSDHFAPACFDVSSVIQ

KNLRFSQRLRLVAGAVPTLH

>030670542.1|Nomascus leucogenys

MPARCVAAHCGNTTKSGKSLFRFPKDRAVRLLWDRFVRGCRADWYGGNDRSVICSDHFAPACFDVSSVIQ

KNLRFSQRLRLVAGAVPTLH

>010361971.1|Rhinopithecus roxellana

MPARCVAAHCGNTTKSGKSLFRFPKDRAVRLLWDRFVRGCRADWYGGNDRSVICSDHFAPACFDVSSVIQ

KNLRFSQRLRLVAGAVPTLH

>004056471.1|Gorilla gorilla gorilla

MPARCVAAHCGNTTKSGKSLFRFPKDRAVRLLWDRFVRGCRADWYGGNDRSVICSDHFAPACFDVSSVIQ

KNLRFSQRLRLVAGAVPTLH

>009208790.1|Papio anubis

MPARCVAAHCGNTTKSGKSLFRFPKDRAVRLLWDRFVRGCRADWYGGNDRSVICSDHFAPACFDVSSVIQ

KNLRFSQRLRLVAGAVPTLH

>023076466.1|Piliocolobus tephrosceles

MPARCVAAHCGNTTKSGKSLFRFPKDRAVRLLWDRFVRGCRADWYGGNDRSVICSDHFAPACFDVSSVIQ

KNLRFSQRLRLVAGAVPTLH

>032100737.1|Sapajus apell

MPARCVAAHCGNTTKTGKSLFRFPKDRAARLLWDRFVRGCRADWYEGHDRSVICSDHFAPACFDVSSVIQ

KNLHFSQRLRLVAGAVPTLH

>032031979.1|Hylobates moloch

MPARCVAAHCGNTTKSGKSLFRFPKDRAVRLLWDRFVRGCRADWYGGNDRSVICSDHFAPACFDVSSVIQ

KNLRFSQRLRLVAGAVPTLH

>033045437.1|Trachypithecus francoisi

MPARCVAAHCGNTTKSGKSLFRFPKDRAVRLLWDRFVRGCRADWYGGNDRSVICSDHFAPACFDVSSVIQ

KNLRFSQRLRLVAGAVPTLH

>NP_064532.1| Homo sapiens

MPARCVAAHCGNTTKSGKSLFRFPKDRAVRLLWDRFVRGCRADWYGGNDRSVICSDHFAPACFDVSSVIQKNLRFSQRLRLVAGAVPTLH
