## Additional file for "Synapomorphic variations in the THAP domains of the human THAP protein family and its homologs": THAP11.docx

>JAQ16213.1|Lygus hesperus

MSAKGSKNICCVPGCGNMYTNSNVTLYSFPGKPYELERKYRWVQAVNRLNADGSLWEPSKYSKICSHHFV

GGQMSNIADHPAYVPSIF

>KXJ15407.1|Exaiptasia pallida

MADDKRSPAVCFEEIFNEISTDNLPKLNKERLQYCCNALKLKVSGEKKELAQRLEPLGKFKELFDKKVTR

IQEDFKFSTALDPHSIPPLSANWKVVGKQTEVAVPTVT

>030437815.1|Gopherus evgoodei

MPGFTCCVPGCYNNSHRDKALHFYTFPKDAELRRLWLKNVSRAGVSGCFSTFQPTTGHRVCSEHFPGGRK

SYLVRVPTIF

>032644409.1|Chelonoidis abingdonii

MPGFTCCVPGCYNNSHRDKALHFYTFPKDAELRRLWLKNVSRAGVSGCFSTFQPTTGHRVCSEHFPGGRK

SYLVRVPTIF

>020366050.1|Rhincodon typus

MPGFTCCVPGCYNNSHRDRELRFYTFPKDEEQRRLWLQNISRTGVSGCFSTFYPTTGHRVCSAHFPGGRK

TYSVKVPTIF

>032892153.1|Amblyraja radiata

MPGFTCCVPGCYNNSHRDRDLRFYTFPKDEEQRRVWLQNISRTGVTGCFSTFYPTTGHRVCSAHFPGGRK

TYSINVPTIF

>029464683.1|Rhinatrema bivittatum

MPGFTCCVPGCYSNSHRDKALHFYTFPKDPELRRLWLKNVSRAGVSGCFSTFHPTTGHRVCSLHFPGGRK

TYSVRVPTIF

>030060789.1|Microcaecilia unicolor

MPGFTCCVPGCYSNSHRDKALHFYTFPKDAELRRLWLKNVSRAGVSGCFSTFHPTTGHRVCSLHFPGGRK

TYSVRVPTIF

>002935416.2|Xenopus tropicalis

MPGFTCCVPGCYSNSHRDKGLHFYTFPKDPELRCLWLKNVSRGGVSGCFSTFQPTNGHRVCSLHFQGGRK

SYSIKVPTIF

>020666641.1|Pogona vitticeps

MPGFTCCVPGCYNNSHRDKALHFYTFPKDAELRRLWLKNVSRAGVSGCFSTFQPTTGHRVCSEHFPGGRK

SYLVRVPTIF

>026527806.1|Notechis scutatus

MPGFTCCVPGCYNNSHRDKALHFYTFPKDAELRRLWLKNVSRAGVSGCFSTFQPTTGHRVCSEHFPGGRK

SYLVRVPTIF

>028595562.1|Podarcis muralis

MPGFTCCVPGCYNNSHRDKALHFYTFPKDAELRRLWLKNVSRAGVSGCFSTFQPTTGHRVCSEHFPGGRK

SYLVRVPTIF

>032087246.1|Thamnophis elegans

MPGFTCCVPGCYNNSHRDKALHFYTFPKDAELRRLWLKNVSRAGVSGCFSTFQPTTGHRVCSEHFPGGRK

SYLVRVPTIF

>033014807.1|Lacerta agilis

MPGFTCCVPGCYNNSHRDKALHFYTFPKDAELRRLWLKNVSRAGVSGCFSTFQPTTGHRVCSEHFPGGRK

SYLVRVPTIF

>OWK61738.1|Lonchura striatadomestica

MPGFTCCVPGCYNNSHRDKALHFYTFPKDEELRRLWLKNVSRAGVSGCFSTFQPTTGHRVCSEHFQGGRK

SYLVRVPTIF

>015134639.1|Gallus gallus

MPGFTCCVPGCYNNSHRDKALHFYTFPKDEELRRLWLKNVSRAGVSGCFSTFQPTTGHRVCSEHFQGGRK

SYLVRVPTIF

>025941206.1|Apteryx rowi

MPGFTCCVPGCYNNSHRDKALHFYTFPKDEELRRLWLKNVSRAGVSGCFSTFQPTTGHRVCSEHFQGGRK

SYLVRVPTIF

>026651044.1|Zonotrichia albicollis

MPGFTCCVPGCYNNSHRDKALHFYTFPKDEELRRLWLKNVSRAGVSGCFSTFQPTTGHRVCSEHFQGGRK

SYLVRVPTIF

>027321892.1|Anas platyrhynchos

MPGFTCCVPGCYNNSHRDKALHFYTFPKDEELRRLWLKNVSRAGVSGCFSTFQPTTGHRVCSEHFQGGRK

SYLVRVPTIF

>027541061.1|Neopelma chrysocephalum

MPGFTCCVPGCYNNSHRDKALHFYTFPKDEELRRLWLKNVSRAGVSGCFSTFQPTTGHRVCSEHFQGGRK

SYLVRVPTIF

>027572006.1|Pipra filicauda

MPGFTCCVPGCYNNSHRDKALHFYTFPKDEELRRLWLKNVSRAGVSGCFSTFQPTTGHRVCSEHFQGGRK

SYLVRVPTIF

>027762250.1|Empidonax traillii

MPGFTCCVPGCYNNSHRDKALHFYTFPKDEELRRLWLKNVSRAGVSGCFSTFQPTTGHRVCSEHFQGGRK

SYLVRVPTIF

>029881368.1|Aquila chrysaetoschrysaetos

MPGFTCCVPGCYNNSHRDKALHFYTFPKDEELRRLWLKNVSRAGVSGCFSTFQPTTGHRVCSEHFQGGRK

SYLVRVPTIF

>030083387.1|Serinus canaria

MPGFTCCVPGCYNNSHRDKALHFYTFPKDEELRRLWLKNVSRAGVSGCFSTFQPTTGHRVCSEHFQGGRK

SYLVRVPTIF

>030313656.1|Calypte anna

MPGFTCCVPGCYNNSHRDKALHFYTFPKDEELRRLWLKNVSRAGVSGCFSTFQPTTGHRVCSEHFQGGRK

SYLVRVPTIF

>031471124.1|Phasianus colchicus

MPGFTCCVPGCYNNSHRDKALHFYTFPKDEELRRLWLKNVSRAGVSGCFSTFQPTTGHRVCSEHFQGGRK

SYLVRVPTIF

>031977633.1|Corvus moneduloides

MPGFTCCVPGCYNNSHRDKALHFYTFPKDEELRRLWLKNVSRAGVSGCFSTFQPTTGHRVCSEHFQGGRK

SYLVRVPTIF

>032051428.1|Aythya fuligula

MPGFTCCVPGCYNNSHRDKALHFYTFPKDEELRRLWLKNVSRAGVSGCFSTFQPTTGHRVCSEHFQGGRK

SYLVRVPTIF

>032557449.1|Chiroxiphia lanceolata

MPGFTCCVPGCYNNSHRDKALHFYTFPKDEELRRLWLKNVSRAGVSGCFSTFQPTTGHRVCSEHFQGGRK

SYLVRVPTIF

>030138180.2|Taeniopygia guttata

MPGFTCCVPGCYNNSHRDKALHFYTFPKDEELRRLWLKNVSRAGVSGCFSTFQPTTGHRVCSEHFQGGRK

SYLVRVPTIF

>030327286.1|Strigops habroptila

MPGFTCCVPGCYNNSHRDKALHFYTFPKDEELRRLWLKNVSRAGVSGCFSTFQPTTGHRVCSEHFQGGRK

SYLVRVPTIF

>015495180.1|Parus major

MPGFTCCVPGCYNNSHRDKALHFYTFPKDEELRRLWLKNVSRAGVSGCFSTFQPTTGHRVCSEHFQGGRK

SYLVRVPTIF

>020462136.1|Monopterus albus

MPGFTCCVPGCYNNSHRDRDLRFYTFPKDTALRELWLRNISRAGVSGCFSTFQPPTGHRVCSVHFAGGRK

TYTIRVPSLF

>020797324.1|Boleophthalmus pectinirostris

MPGFTCCVPGCYNNSHRDRELRFYTFPKDAALREQWLRNISRAGVSGCFSTFQPTTGHRVCSVHFAGGRK

TYTIRVPSLF

>021163455.1|Fundulus heteroclitus

MPGFTCCVPGCYNNSHRDKELRFYTFPKDPALREQWLRNISRAGVSGCFSTFQPTTGHRVCSVHFAGGRK

TYTVRVPSLF

>022061617.1|Acanthochromis polyacanthus

MPGFTCCVPGCYNNSHRDRDLRFYTFPKDSTLRELWLRNISRAGVSGCFSTFQPTTGHRVCSVHFAGGRK

TYSIRVPSLF

>SBR82524.1|Nothobranchius pienaari

MPGFTCCVPGCYNNSHRDKELRFYTFPKDSALRELWLRNISRAGVSGCFSTFQPTTGHRVCSVHFAGGRK

TYSVRVPSLF

>SBQ71127.1|Nothobranchius korthausae

MPGFTCCVPGCYNNSHRDKELRFYTFPKDNALRELWLRNISRAGVSGCFSTFQPTTGHRVCSVHFAGGRK

TYSVRVPTLF

>SBQ44582.1|Nothobranchius kadleci

MPGFTCCVPGCYNNSHRDKELRFYTFPKDNALRELWLRNISRAGVSGCFSTFQPTTGHRVCSVHFAGGRK

TYSVRVPSLF

>SBR76867.1|Nothobranchius rachovii

MPGFTCCVPGCYNNSHRDKELRFYTFPKDNALRELWLRNISRAGVSGCFSTFQPTTGHRVCSVHFAGGRK

TYSVRVPSLF

>SBS55550.1|Nothobranchius furzeri

MPGFTCCVPGCYNNSHRDKELRFYTFPKDNALRELWLRNISRAGVSGCFSTFQPTTGHRVCSVHFAGGRK

TYSVRVPSLF

>022601851.1|Seriola dumerili

MPGFTCCVPGCYNNSHRDRDLRFYTFPKDTALRELWLRNISRAGVSGCFSTFQPTTGHRVCSVHFAGGRK

TYSIRVPSLF

>023133175.1|Amphiprion ocellaris

MPGFTCCVPGCYNNSHRDRDLRFYTFPKDSTLRELWLRNISRAGVSGCFSTFQPTTGHRVCSVHFAGGRK

TYSIRVPSLF

>005807775.1|Xiphophorus maculatus

MPGFTCCVPGCYNNSHRDKDLRFYTFPKDPALREQWLRNISRAGVSGCFSTFQPTTGHRVCSVHFAGGRK

TYIVRVPSLF

>023251533.1|Seriola lalandi dorsalis

MPGFTCCVPGCYNNSHRDRDLRFYTFPKDTALRELWLRNISRAGVSGCFSTFQPTTGHRVCSVHFAGGRK

TYSIRVPSLF

>023673894.1|Paramormyrops kingsleyae

MPGFTCCVPGCYNNSHRDRELRFYTFPKDTAQREIWLKNISRAGVSGCFSTFQPTTGHRVCSTHFAGGRK

TYSIRVPTLF

>004067383.1|Oryzias latipes

MPGFTCCVPGCYNNSHRDRELRFYTFPKDTTLREQWLRNISRAGVSGCFSTFQPTTGHRVCSVHFAGGRK

TYSVRVPSLF

>024151422.1|Oryzias melastigma

MPGFTCCVPGCYNNSHRDRELRFYTFPKDTTLREQWLRNISRAGVSGCFSTFQPTTGHRVCSVHFAGGRK

TYSVRVPSLF

>004543100.2|Maylandia zebra

MPGFTCCVPGCYNNSHRDRELRFYTFPKDTALREQWLRNISRAGVSGCFSTFQPTTGHRVCSVHFAGGRK

TYSIRVPSLF

>017296486.1|Kryptolebias marmoratus

MPGFTCCVPGCYNNSHRDRDLRFYTFPKDSALREQWLRNISRAGVSGCFSTFQPTTGHRVCSVHFAGGRK

TYSVRVPSLF

>008308461.1|Cynoglossus semilaevis

MPGFTCCVPGCYNNSHRDRDLRFYTFPKDTELREQWLRNISRAGVSGCFSTFQPTTGHRVCSVHFSGGRK

TYTIRVPSLF

>005467153.1|Oreochromis niloticus

MPGFTCCVPGCYNNSHRDRELRFYTFPKDTALREQWLRNISRAGVSGCFSTFQPTTGHRVCSVHFAGGRK

TYSIRVPSLF

>026025120.1|Astatotilapia calliptera

MPGFTCCVPGCYNNSHRDRELRFYTFPKDTALREQWLRNISRAGVSGCFSTFQPTTGHRVCSVHFAGGRK

TYSIRVPSLF

>026786298.1|Pangasianodon hypophthalmus

MPGFTCCVPGCYNNSHRDRELRFYTFPKDPTQREIWLKNISRAGVSGCFSTFQPTTGHRVCSVHFPGGRK

TYTIRVPTLF

>026868598.1|Electrophorus electricus

MPGFTCCVPGCYNNSHRDRELRFYTFPKDPTQREIWLKNISRAGVSGCFSTFQPTTGHRVCSVHFPGGRK

TYTIRVPTLF

>027007806.1|Tachysurus fulvidraco

MPGFTCCVPGCYNNSHRDRELRFYTFPKDPTQREIWLKNISRAGVSGCFSTFQPTTGHRVCSVHFPGGRK

TYTIRVPTLF

>010739566.1|Larimichthys crocea

MPGFTCCVPGCYNNSHRDRDLRFYTFPKDTTLRELWLRNISRAGVSGCFSTFQPTTGHRVCSVHFAGGRK

TYSIRVPSLF

>027870089.1|Xiphophorus couchianus

MPGFTCCVPGCYNNSHRDKDLRFYTFPKDPALREQWLRNISRAGVSGCFSTFQPTTGHRVCSVHFAGGRK

TYIVRVPSLF

>028257770.1|Parambassis ranga

MPGFTCCVPGCYNNSHRDRDLRFYTFPKDTTLRELWLRNISRAGVSGCFSTFQPTTGHRVCSVHFAGGRK

TYTVRVPSLF

>028299014.1|Gouania willdenowi

MPGFTCCVPGCYNNSHRDRELRFYTFPKDNVLREQWLRNISRAGVSGCFSTFQPTTGHRVCSVHFAGGRK

TYTIRVPSLF

>028431235.1|Perca flavescens

MPGFTCCVPGCYNNSHRDRDLRFYTFPKDTTLRELWLRNISRAGVSGCFSTFQPTTGHRVCSVHFPGGRK

TYSIRVPSLF

>028665454.1|Erpetoichthys calabaricus

MPGFTCCVPGCYNNSHRDRELRFYTFPKDPVQREIWLKNISRAGVSGCFSTFQPTTGHRVCSIHFAGGRK

TYSIRVPTLF

>028814717.1|Denticeps clupeoides

MPGFTCCVPGCYNNSQRDRDLRFYTFPKDTTQREIWLKNISRSGVKGCFSTFQPTTGHRVCSVHFAGGRK

TYTVRIPTLF

>010878148.1|Esox lucius

MPGFTCCVPGCYNNSHRDRELRFYTFPKDTTQREIWLKNISRAGVSGCFSTFQPTTGHRVCSVHFSGGRK

TYTIRVPTLF

>029000483.1|Betta splendens

MPGFTCCVPGCYNNSHRDRDLRFYTFPKDTTLREMWLRNISRAGVSGCFSTFQPTTGHRVCSVHFAGGRK

TYSIRVPSLF

>029112987.1|Scleropages formosus

MPGFTCCVPGCYNNSHRDRDLRFYTFPKDTAQREIWLKNISRAGVSGCFSTFQPTTGHRVCSSHFPGGRK

TYSIRVPTLF

>020481780.1|Labrus bergylta

MPGFTCCVPGCYNNSHRDRELRFYTFPKDTTLREMWLRNISRAGVSGCFSTFQPTTGHRVCSVHFAGGRK

TYSIRVPSLF

>029283941.1|Cottoperca gobio

MPGFTCCVPGCYNNSHRDRDLRFYTFPKDSTLREQWLRNISRAGVSGCFSTFQPTTGHRVCSVHFAGGRK

TYTIRVPSLF

>TNN53102.1|Liparis tanakae

MVHTCVVAGCRNRRTPGTTLSFYRFPRDSERKQRWIAAVNREGWVPNDGSRLCSSHFVSGKQVKNPRSPD

YVPSVF

>029354106.1|Echeneis naucrates

MPGFTCCVPGCYNNSHRDRDLRFYTFPKDTALREMWLRNISRAGVSGCFSTFQPTTGHRVCSVHFAGGRK

TYSIRVPSLF

>003970101.1|Takifugu rubripes

MPGFTCCVPGCYNNSHRDRDLRFYTFPKDTTLREQWLRNISRAGVSGCFSTFEPTTGHRVCSVHFAGGRK

TYSVRVPSLF

>029900941.1|Myripristis murdjan

MPGFTCCVPGCYNNSHRDRELRFYTFPKDTTQREIWLKNISRAGVSGCFSTFQPTTGHRVCSVHFAGGRK

TYNIRVPTLF

>029953682.1|Salarias fasciatus

MPGFTCCVPGCYNNSHRDRDLRFYTFPKDTTLREQWLRNISRAGVSGCFSTFQPTTGHRVCSVHFAGGRK

TYTVRVPSLF

>029981311.1|Sphaeramia orbicularis

MPGFTCCVPGCYNNSHRDRDLRFYTFPKDTTLREMWLRNISRAGVSGCFSTFQPTTGHRVCSVHFAGGRK

TYSIRVPSLF

>030580436.1|Archocentrus centrarchus

MPGFTCCVPGCYNNSHRDRDLRFYTFPKDTALREQWLRNISRAGVSGCFSTFQPTTGHRVCSVHFAGGRK

TYSIRVPSLF

>031170248.1|Sander lucioperca

MPGFTCCVPGCYNNSHRDRDLRFYTFPKDTTLRELWLRNISRAGVSGCFSTFQPTTGHRVCSVHFAGGRK

TYSIRVPSLF

>012687875.1|Clupea harengus

MPGFTCCVPGCYNNSQRDRDLRFYTFPKDTTQREIWLKNISRSGVKGCFSTFQPTTGHRVCSEHFAGGRK

TYTIRIPTIF

>031613866.1|Oreochromis aureus

MPGFTCCVPGCYNNSHRDRELRFYTFPKDTALREQWLRNISRAGVSGCFSTFQPTTGHRVCSVHFAGGRK

TYSIRVPSLF

>031701944.1|Anarrhichthys ocellatus

MPGFTCCVPGCYNNSHRDRDLRFYTFPKDTTLRELWLRNISRAGVSGCFSTFQPTTGHRVCSVHFAGGRK

TYTIRVPSLF

>032373924.1|Etheostoma spectabile

MPGFTCCVPGCYNNSHRDRDLRFYTFPKDTTLRELWLRNISRAGVSGCFSTFQPTTGHRVCSVHFAGGRK

TYSIRVPSLF

>032417271.1|Xiphophorus hellerii

MPGFTCCVPGCYNNSHRDKDLRFYTFPKDPALREQWLRNISRAGVSGCFSTFQPTTGHRVCSVHFAGGRK

TYIVRVPSLF

>026175615.1|Mastacembelus armatus

MPGFTCCVPGCYNNSHRDRDLRFYTFPKDSTLRELWLRNISRAGVSGCFSTFQPTTGHRVCSVHFAGGRK

TYSIRVPSLF

>026233854.1|Anabas testudineus

MPGFTCCVPGCYNNSHRDRDLRFYTFPKDATLRELWLRNISRAGVSGCFSTFQPTTGHRVCSVHFAGGRK

TYSIRVPSLF

>033486091.1|Epinephelus lanceolatus

MPGFTCCVPGCYNNSHRDKDLRFYTFPKDSTLREIWLRNISRAGVSGCFSTFQPTTGHRVCSVHFAGGRK

TYTIRVPSLF

>998267.1|Danio rerio

MPGFTCCVPGCYNNSHRDRDLRFYTFPKDPTQREIWLKNISRAGVSGCFSTFQPTTGHRVCSVHFPGGRK

TYTIRVPTLF

>020844972.1|Phascolarctos cinereus

MPGFTCCVPGCYNNSHRDKALHFYTFPKDAELRRQWLKNVSRAGVSGCFSTFQPTAGHRLCSVHFQGGRK

TYTVRVPTIF

>027701568.1|Vombatus ursinus

MPGFTCCVPGCYNNSHRDKALHFYTFPKDAELRRQWLKNVSRAGVSGCFSTFQPTAGHRLCSVHFQGGRK

TYTVRVPTIF

>007658492.2|Ornithorhynchus anatinus

MPGFTCCVPGCYNNSHRDKALHFYTFPKDAELRRLWLKNVSRAGVSGCFSTFQPTTGHRLCSVHFQGGRK

TYTVRVPTIF

>023350824.2|Sarcophilus harrisii

MPGFTCCVPGCYNNSHRDKALHFYTFPKDAELRRQWLKNVSRAGVSGCFSTFQPTAGHRLCSVHFQGGRK

TYTVRVPTIF

>PNI91157.1|Pan troglodytes

MPGFTCCVPGCYNNSHRDKALHFYTFPKDAELRRLWLKNVSRAGVSGCFSTFQPTTGHRLCSVHFQGGRK

TYTVRVPTIF

>PNJ61365.1|Pongo abelii

MPGFTCCVPGCYNNSHRDKALHFYTFPKDAELRRLWLKNVSRAGVSGCFSTFQPTTGHRLCSVHFQGGRK

TYTVRVPTIF

>JAB12082.1|Callithrix jacchus

MPGFTCCVPGCYNNSHRDKALHFYTFPKDAELRRLWLKNVSRAGVSGCFSTFQPTTGHRLCSVHFQGGRK

TYTVRVPTIF

>AAI38967.1|Mus musculus

MPGFTCCVPGCYNNSHRDKALHFYTFPKDAELRRLWLKNVSRAGVSGCFSTFQPTTGHRLCSVHFQGGRK

TYTVRVPTIF

>020026757.1|Castor canadensis

MPGFTCCVPGCYNNSHRDKALHFYTFPKDAELRRLWLKNVSRAGVSGCFSTFQPTTGHRLCSVHFQGGRK

TYTVRVPTIF

>012597922.1|Microcebus murinus

MPGFTCCVPGCYNNSHRDKALHFYTFPKDAELRRLWLKNVSRAGVSGCFSTFQPTTGHRLCSVHFQGGRK

TYTVRVPTIF

>020740754.1|Odocoileus virginianustexanus

MPGFTCCVPGCYNNSHRDKALHFYTFPKDAELRRLWLKNVSRAGVSGCFSTFQPTTGHRLCSVHFQGGRK

TYTVRVPTIF

>005076362.1|Mesocricetus auratus

MPGFTCCVPGCYNNSHRDKALHFYTFPKDAELRRLWLKNVSRAGVSGCFSTFQPTTGHRLCSVHFQGGRK

TYTVRVPTIF

>021112499.1|Heterocephalus glaber

MPGFTCCVPGCYNNSHRDKALHFYTFPKDAELRRLWLKNVSRAGVSGCFSTFQPTTGHRLCSVHFQGGRK

TYTVRVPTIF

>021487840.1|Meriones unguiculatus

MPGFTCCVPGCYNNSHRDKALHFYTFPKDAELRRLWLKNVSRAGVSGCFSTFQPTTGHRLCSVHFQGGRK

TYTVRVPTIF

>012323606.1|Aotus nancymaae

MPGFTCCVPGCYNNSHRDKALHFYTFPKDAELRRLWLKNVSRAGVSGCFSTFQPTTGHRLCSVHFQGGRK

TYTVRVPTIF

>021559252.1|Neomonachus schauinslandi

MPGFTCCVPGCYNNSHRDKALHFYTFPKDAELRRLWLKNVSRAGVSGCFSTFQPTTGHRLCSVHFQGGRK

TYTVRVPTIF

>008063947.1|Carlito syrichta

MPGFTCCVPGCYNNSHRDKALHFYTFPKDAELRRLWLKNVSRAGVSGCFSTFQPTTGHRLCSVHFQGGRK

TYTVRVPTIF

>005318389.1|Ictidomys tridecemlineatus

MPGFTCCVPGCYNNSHRDKALHFYTFPKDAELRRLWLKNVSRAGVSGCFSTFQPTTGHRLCSVHFQGGRK

TYTVRVPTIF

>005620865.1|Canis lupus familiaris

MPGFTCCVPGCYNNSHRDKALHFYTFPKDAELRRLWLKNVSRAGVSGCFSTFQPTTGHRLCSVHFQGGRK

TYTVRVPTIF

>003998187.2|Felis catus

MPGFTCCVPGCYNNSHRDKALHFYTFPKDAELRRLWLKNVSRAGVSGCFSTFQPTTGHRLCSVHFQGGRK

TYTVRVPTIF

>023374670.1|Otolemur garnettii

MPGFTCCVPGCYNNSHRDKALHFYTFPKDAELRRLWLKNVSRAGVSGCFSTFQPTTGHRLCSVHFQGGRK

TYTVRVPTIF

>010595049.1|Loxodonta africana

MPGFTCCVPGCYNNSHRDKALHFYTFPKDAELRRLWLKNVSRAGVSGCFSTFQPTTGHRLCSVHFQGGRK

TYTVRVPTIF

>004459715.1|Dasypus novemcinctus

MPGFTCCVPGCYNNSHRDKALHFYTFPKDAELRRLWLKNVSRAGVSGCFSTFQPTTGHRLCSVHFQGGRK

TYTVRVPTIF

>005004698.1|Cavia porcellus

MPGFTCCVPGCYNNSHRDKALHFYTFPKDAELRRLWLKNVSRAGVSGCFSTFQPTTGHRLCSVHFQGGRK

TYTVRVPTIF

>011378810.1|Pteropus vampyrus

MPGFTCCVPGCYNNSHRDKALHFYTFPKDAELRRLWLKNVSRAGVSGCFSTFQPTTGHRLCSVHFQGGRK

TYTVRVPTIF

>005614984.2|Equus caballus

MPGFTCCVPGCYNNSHRDKALHFYTFPKDAELRRLWLKNVSRAGVSGCFSTFQPTTGHRLCSVHFQGGRK

TYTVRVPTIF

>004625911.1|Octodon degus

MPGFTCCVPGCYNNSHRDKALHFYTFPKDAELRRLWLKNVSRAGVSGCFSTFQPTTGHRLCSVHFQGGRK

TYTVRVPTIF

>006099500.1|Myotis lucifugus

MPGFTCCVPGCYNNSHRDKALHFYTFPKDAELRRLWLKNVSRAGVSGCFSTFQPTTGHRLCSVHFQGGRK

TYTVRVPTIF

>024411704.1|Desmodus rotundus

MPGFTCCVPGCYNNSHRDKALHFYTFPKDAELRRLWLKNVSRAGVSGCFSTFQPTTGHRLCSVHFQGGRK

TYTVRVPTIF

>024589843.1|Neophocaena asiaeorientalisasiaeorientalis

MPGFTCCVPGCYNNSHRDKALHFYTFPKDAELRRLWLKNVSRAGVSGCFSTFQPTTGHRLCSVHFQGGRK

TYTVRVPTIF

>011756085.1|Macaca nemestrina

MPGFTCCVPGCYNNSHRDKALHFYTFPKDAELRRLWLKNVSRAGVSGCFSTFQPTTGHRLCSVHFQGGRK

TYTVRVPTIF

>003812224.1|Pan paniscus

MPGFTCCVPGCYNNSHRDKALHFYTFPKDAELRRLWLKNVSRAGVSGCFSTFQPTTGHRLCSVHFQGGRK

TYTVRVPTIF

>006909839.1|Pteropus alecto

MPGFTCCVPGCYNNSHRDKALHFYTFPKDAELRRLWLKNVSRAGVSGCFSTFQPTTGHRLCSVHFQGGRK

TYTVRVPTIF

>006072059.1|Bubalus bubalis

MPGFTCCVPGCYNNSHRDKALHFYTFPKDAELRRLWLKNVSRAGVSGCFSTFQPTTGHRLCSVHFQGGRK

TYTVRVPTIF

>025226042.1|Theropithecus gelada

MPGFTCCVPGCYNNSHRDKALHFYTFPKDAELRRLWLKNVSRAGVSGCFSTFQPTTGHRLCSVHFQGGRK

TYTVRVPTIF

>025282362.1|Canis lupus dingo

MPGFTCCVPGCYNNSHRDKALHFYTFPKDAELRRLWLKNVSRAGVSGCFSTFQPTTGHRLCSVHFQGGRK

TYTVRVPTIF

>025744830.1|Callorhinus ursinus

MPGFTCCVPGCYNNSHRDKALHFYTFPKDAELRRLWLKNVSRAGVSGCFSTFQPTTGHRLCSVHFQGGRK

TYTVRVPTIF

>025867308.1|Vulpes vulpes

MPGFTCCVPGCYNNSHRDKALHFYTFPKDAELRRLWLKNVSRAGVSGCFSTFQPTTGHRLCSVHFQGGRK

TYTVRVPTIF

>026269031.1|Urocitellus parryii

MPGFTCCVPGCYNNSHRDKALHFYTFPKDAELRRLWLKNVSRAGVSGCFSTFQPTTGHRLCSVHFQGGRK

TYTVRVPTIF

>026350827.1|Ursus arctoshorribilis

MPGFTCCVPGCYNNSHRDKALHFYTFPKDAELRRLWLKNVSRAGVSGCFSTFQPTTGHRLCSVHFQGGRK

TYTVRVPTIF

>005345506.1|Microtus ochrogaster

MPGFTCCVPGCYNNSHRDKALHFYTFPKDAELRRLWLKNVSRAGVSGCFSTFQPTTGHRLCSVHFQGGRK

TYTVRVPTIF

>026931823.1|Acinonyx jubatus

MPGFTCCVPGCYNNSHRDKALHFYTFPKDAELRRLWLKNVSRAGVSGCFSTFQPTTGHRLCSVHFQGGRK

TYTVRVPTIF

>026975364.1|Lagenorhynchus obliquidens

MPGFTCCVPGCYNNSHRDKALHFYTFPKDAELRRLWLKNVSRAGVSGCFSTFQPTTGHRLCSVHFQGGRK

TYTVRVPTIF

>003504337.1|Cricetulus griseus

MPGFTCCVPGCYNNSHRDKALHFYTFPKDAELRRLWLKNVSRAGVSGCFSTFQPTTGHRLCSVHFQGGRK

TYTVRVPTIF

>027473935.1|Zalophus californianus

MPGFTCCVPGCYNNSHRDKALHFYTFPKDAELRRLWLKNVSRAGVSGCFSTFQPTTGHRLCSVHFQGGRK

TYTVRVPTIF

>027809723.1|Marmota flaviventris

MPGFTCCVPGCYNNSHRDKALHFYTFPKDAELRRLWLKNVSRAGVSGCFSTFQPTTGHRLCSVHFQGGRK

TYTVRVPTIF

>014955983.2|Ovis aries

MPGFTCCVPGCYNNSHRDKALHFYTFPKDAELRRLWLKNVSRAGVSGCFSTFQPTTGHRLCSVHFQGGRK

TYTVRVPTIF

>027953042.1|Eumetopias jubatus

MPGFTCCVPGCYNNSHRDKALHFYTFPKDAELRRLWLKNVSRAGVSGCFSTFQPTTGHRLCSVHFQGGRK

TYTVRVPTIF

>007198203.1|Balaenoptera acutorostratascammoni

MQSTHKALLPYTMAVLRRCMWRRAAGRLARRGPMGGVFWAQGAGPVWLCRAAVVGGCPRPRAVPPVKGRA

AAYRPSCRKAQNAGGLGWAGPPGAAMPGFTCCVPGCYNNSHRDEALHFYTFPKDAELRRLWLKNVSRGGI

SGCFSTFQPTTGHRLCSVHFQGGRKTYTVRVPTIF

>028341398.1|Physeter catodon

MPGFTCCVPGCYNNSHRDKALHFYTFPKDAELRRLWLKNVSRAGVSGCFSTFQPTTGHRLCSVHFQGGRK

TYTVRVPTIF

>028357719.1|Phyllostomus discolor

MPGFTCCVPGCYNNSHRDKALHFYTFPKDAELRRLWLKNVSRAGVSGCFSTFQPTAGHRLCSVHFQGGRK

TYAVRVPTIF

>028624535.1|Grammomys surdaster

MPGFTCCVPGCYNNSHRDKALHFYTFPKDAELRRLWLKNVSRAGVSGCFSTFQPTTGHRLCSVHFQGGRK

TYTVRVPTIF

>028710197.1|Peromyscus leucopus

MPGFTCCVPGCYNNSHRDKALHFYTFPKDAELRRLWLKNVSRAGVSGCFSTFQPTTGHRLCSVHFQGGRK

TYTVRVPTIF

>014981922.1|Macaca mulatta

MPGFTCCVPGCYNNSHRDKALHFYTFPKDAELRRLWLKNVSRAGVSGCFSTFQPTTGHRLCSVHFQGGRK

TYTVRVPTIF

>029092888.1|Monodon monoceros

MPGFTCCVPGCYNNSHRDKALHFYTFPKDAELRRLWLKNVSRAGVSGCFSTFQPTTGHRLCSVHFQGGRK

TYTVRVPTIF

>021025916.1|Mus caroli

MPGFTCCVPGCYNNSHRDKALHFYTFPKDAELRRLWLKNVSRAGVSGCFSTFQPTTGHRLCSVHFQGGRK

TYTVRVPTIF

>021076048.1|Mus pahari

MPGFTCCVPGCYNNSHRDKALHFYTFPKDAELRRLWLKNVSRAGVSGCFSTFQPTTGHRLCSVHFQGGRK

TYTVRVPTIF

>008837286.1|Nannospalax galili

MPGFTCCVPGCYNNSHRDKALHFYTFPKDAELRRLWLKNVSRAGVSGCFSTFQPTTGHRLCSVHFQGGRK

TYTVRVPTIF

>029780035.1|Suricata suricatta

MPGFTCCVPGCYNNSHRDKALHFYTFPKDAELRRLWLKNVSRAGVSGCFSTFQPTTGHRLCSVHFQGGRK

TYTVRVPTIF

>030154855.1|Lynx canadensis

MPGFTCCVPGCYNNSHRDKALHFYTFPKDAELRRLWLKNVSRAGVSGCFSTFQPTTGHRLCSVHFQGGRK

TYTVRVPTIF

>022439139.1|Delphinapterus leucas

MPGFTCCVPGCYNNSHRDKALHFYTFPKDAELRRLWLKNVSRAGVSGCFSTFQPTTGHRLCSVHFQGGRK

TYTVRVPTIF

>012360178.2|Nomascus leucogenys

MPGFTCCVPGCYNNSHRDKALHFYTFPKDAELRRLWLKNVSRAGVSGCFSTFQPTTGHRLCSVHFQGGRK

TYTVRVPTIF

>004704659.1|Echinops telfairi

MPGFTCCVPGCYNNSHRDKALHFYTFPKDAELRRLWLKNVSRAGVSGCFSTFQPTTGHRLCSVHFQGGRK

TYTVRVPTIF

>030719496.1|Globicephala melas

MPGFTCCVPGCYNNSHRDKALHFYTFPKDAELRRLWLKNVSRAGVSGCFSTFQPTTGHRLCSVHFQGGRK

TYTVRVPTIF

>010353382.2|Rhinopithecus roxellana

MPGFTCCVPGCYNNSHRDKALHFYTFPKDAELRRLWLKNVSRAGVSGCFSTFQPTTGHRLCSVHFQGGRK

TYTVRVPTIF

>006749030.1|Leptonychotes weddellii

MPGFTCCVPGCYNNSHRDKALHFYTFPKDAELRRLWLKNVSRAGVSGCFSTFQPTTGHRLCSVHFQGGRK

TYTVRVPTIF

>030858506.1|Gorilla gorilla gorilla

MPGFTCCVPGCYNNSHRDKALHFYTFPKDAELRRLWLKNVSRAGVSGCFSTFQPTTGHRLCSVHFQGGRK

TYTVRVPTIF

>001098464.1|Bos taurus

MPGFTCCVPGCYNNSHRDKALHFYTFPKDAELRRLWLKNVSRAGVSGCFSTFQPTTGHRLCSVHFQGGRK

TYTVRVPTIF

>031197103.1|Mastomys coucha

MPGFTCCVPGCYNNSHRDKALHFYTFPKDAELRRLWLKNVSRAGVSGCFSTFQPTTGHRLCSVHFQGGRK

TYTVRVPTIF

>010977571.2|Camelus dromedarius

MPGFTCCVPGCYNNSHRDKALHFYTFPKDAELRRLWLKNVSRAGVSGCFSTFQPTTGHRLCSVHFQGGRK

TYTVRVPTIF

>009194983.1|Papio anubis

MPGFTCCVPGCYNNSHRDKALHFYTFPKDAELRRLWLKNVSRAGVSGCFSTFQPTTGHRLCSVHFQGGRK

TYTVRVPTIF

>023065971.1|Piliocolobus tephrosceles

MPGFTCCVPGCYNNSHRDKALHFYTFPKDAELRRLWLKNVSRAGVSGCFSTFQPTTGHRLCSVHFQGGRK

TYTVRVPTIF

>032139660.1|Sapajus apella

MPGFTCCVPGCYNNSHRDKALHFYTFPKDAELRRLWLKNVSRAGVSGCFSTFQPTTGHRLCSVHFQGGRK

TYTVRVPTIF

>032179448.1|Mustela erminea

MPGFTCCVPGCYNNSHRDKALHFYTFPKDAELRRLWLKNVSRAGVSGCFSTFQPTTGHRLCSVHFQGGRK

TYTVRVPTIF

>032280189.1|Phoca vitulina

MPGFTCCVPGCYNNSHRDKALHFYTFPKDAELRRLWLKNVSRAGVSGCFSTFQPTTGHRLCSVHFQGGRK

TYTVRVPTIF

>001233111.1|Sus scrofa

MPGFTCCVPGCYNNSHRDKALHFYTFPKDAELRRLWLKNVSRAGVSGCFSTFQPTTGHRLCSVHFQGGRK

TYTVRVPTIF

>032342472.1|Camelus ferus

MPGFTCCVPGCYNNSHRDKALHFYTFPKDAELRRLWLKNVSRAGVSGCFSTFQPTTGHRLCSVHFQGGRK

TYTVRVPTIF

>032470240.1|Phocoena sinus

MPGFTCCVPGCYNNSHRDKALHFYTFPKDAELRRLWLKNVSRAGVSGCFSTFQPTTGHRLCSVHFQGGRK

TYTVRVPTIF

>032710060.1|Lontra canadensis

MPGFTCCVPGCYNNSHRDKALHFYTFPKDAELRRLWLKNVSRAGVSGCFSTFQPTTGHRLCSVHFQGGRK

TYTVRVPTIF

>032743797.1|Rattus rattus

MPGFTCCVPGCYNNSHRDKALHFYTFPKDAELRRLWLKNVSRAGVSGCFSTFQPTTGHRLCSVHFQGGRK

TYTVRVPTIF

>032985223.1|Rhinolophus ferrumequinum

MPGFTCCVPGCYNNSHRDKALHFYTFPKDAELRRLWLKNVSRAGVSGCFSTFQPTTGHRLCSVHFQGGRK

TYTVRVPTIF

>033075621.1|Trachypithecus francoisi

MPGFTCCVPGCYNNSHRDKALHFYTFPKDAELRRLWLKNVSRAGVSGCFSTFQPTTGHRLCSVHFQGGRK

TYTVRVPTIF

>004273222.1|Orcinus orca

MPGFTCCVPGCYNNSHRDKALHFYTFPKDAELRRLWLKNVSRAGVSGCFSTFQPTTGHRLCSVHFQGGRK

TYTVRVPTIF

>001100892.1|Rattus norvegicus

MPGFTCCVPGCYNNSHRDKALHFYTFPKDAELRRLWLKNVSRAGVSGCFSTFQPTTGHRLCSVHFQGGRK

TYTVRVPTIF

>NP_065190.2|Homo Sapiens

MPGFTCCVPGCYNNSHRDKALHFYTFPKDAELRRLWLKNVSRAGVSGCFSTFQPTTGHRLCSVHFQGGRK

TYTVRVPTIF
